# Assembly of 43 diverse human Y chromosomes reveals extensive complexity and variation

**DOI:** 10.1101/2022.12.01.518658

**Authors:** Pille Hallast, Peter Ebert, Mark Loftus, Feyza Yilmaz, Peter A. Audano, Glennis A. Logsdon, Marc Jan Bonder, Weichen Zhou, Wolfram Höps, Kwondo Kim, Chong Li, Savannah J. Hoyt, Philip C. Dishuck, David Porubsky, Fotios Tsetsos, Jee Young Kwon, Qihui Zhu, Katherine M. Munson, Patrick Hasenfeld, William T. Harvey, Alexandra P. Lewis, Jennifer Kordosky, Kendra Hoekzema, Human Genome Structural Variation Consortium (HGSVC), Rachel J. O’Neill, Jan O. Korbel, Chris Tyler-Smith, Evan E. Eichler, Xinghua Shi, Christine R. Beck, Tobias Marschall, Miriam K. Konkel, Charles Lee

**Author notes:** These authors contributed equally to this work (Pille Hallast, Peter Ebert, Mark Loftus, Miriam K. Konkel, Charles Lee). Correspondence to: Charles Lee.

## Abstract

The prevalence of highly repetitive sequences within the human Y chromosome has led to its incomplete assembly and systematic omission from genomic analyses. Here, we present long-read *de novo* assemblies of 43 diverse Y chromosomes spanning 180,000 years of human evolution, including two from deep-rooted African Y lineages, and report remarkable complexity and diversity in chromosome size and structure, in contrast with its low level of base substitution variation. The size of the Y chromosome assemblies varies extensively from 45.2 to 84.9 Mbp and include, on average, 81 kbp of novel sequence per Y chromosome. Half of the male-specific euchromatic region is subject to large inversions with a >2-fold higher recurrence rate compared to inversions in the rest of the human genome. Ampliconic sequences associated with these inversions further show differing mutation rates that are sequence context-dependent and some ampliconic genes show evidence for concerted evolution with the acquisition and purging of lineage-specific pseudogenes. The largest heterochromatic region in the human genome, the Yq12, is composed of alternating arrays of *DYZ1* and *DYZ2* repeat units that show extensive variation in the number, size and distribution of these arrays, but retain a 1:1 copy number ratio of the monomer repeats, consistent with the notion that functional or evolutionary forces are acting on this chromosomal region. Finally, our data suggests that the boundary between the recombining pseudoautosomal region 1 and the non-recombining portions of the X and Y chromosomes lies 500 kbp distal to the currently established boundary. The availability of sequence-resolved Y chromosomes from multiple individuals provides a unique opportunity for identifying new associations of specific traits with Y-chromosomal variants and garnering novel insights into the evolution and function of complex regions of the human genome.

## Introduction

The mammalian sex chromosomes evolved from a pair of autosomes, gradually losing their ability to recombine with each other over increasing lengths, leading to degradation and accumulation of large proportions of repetitive sequences on the Y chromosome^1^. The resulting sequence composition of the human Y chromosome is rich in complex repetitive regions, including highly similar segmental duplications (SDs)^2x,3^. This has made the Y chromosome difficult to assemble, and, paired with reduced gene content, has led to its systematic neglect in genomic analyses.

The first human Y chromosome sequence assembly was generated almost 20 years ago, which provided a high quality but incomplete sequence (53.8% or ∼30.8/57.2 Mbp unresolved in GRCh38 Y)^3^. Less than half (∼25 Mbp) of the GRCh38 Y chromosome is composed of euchromatin, which contains two pseudoautosomal regions, PAR1 and PAR2 (∼3.2 Mbp in total), that actively recombine with homologous regions on the X chromosome and are therefore not considered as part of the male-specific Y region (MSY)^3^. The remainder of the Y-chromosomal euchromatin (∼22 Mbp) has been divided into three main classes according to their sequence composition and evolutionary history^3^: (i) the X-degenerate regions (XDR, ∼8.6 Mbp) are remnants of the ancient autosome from which the X and Y chromosomes evolved, (ii) the X-transposed regions (XTR, ∼3.4 Mbp) resulted from a duplicative transposition event from the X chromosome followed by an inversion, and (iii) the ampliconic regions (∼9.9 Mbp) that contain sequences having up to 99.9% intra-chromosomal identity across tens or hundreds of kilobases (**Fig. 1a**). Besides the euchromatin, the Y contains a large proportion of repetitive and heterochromatic sequences, including the (peri-)centromeric *DYZ3* α-satellite and *DYZ17* arrays, *DYZ18* and *DYZ19* arrays, and the largest contiguous heterochromatic block in the human genome, Yq12, which is known to be highly variable in size^3–5^. All of these heterochromatic regions are thought to be composed predominantly of satellites, simple repeats, and SDs^3,6^.

**Figure 1.**
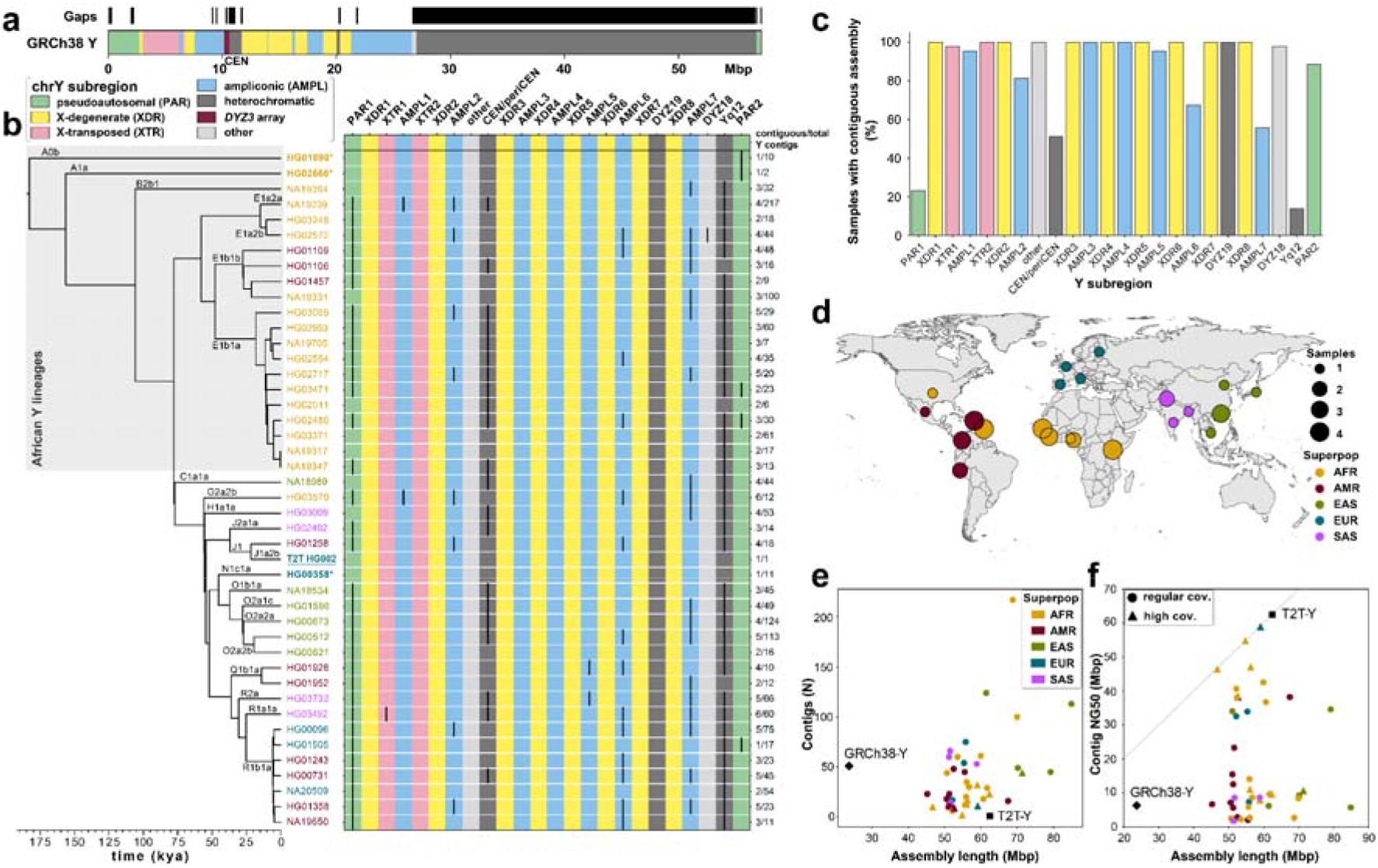
*De novo* assembly outcome. **a.** Human Y chromosome structure based on the GRCh38 Y reference sequence. **b.** Phylogenetic relationships (left) with haplogroup labels of the analysed Y chromosomes with branch lengths drawn proportional to the estimated times between successive splits (see **Fig. S1** and **Table S1** for additional details). Summary of Y chromosome assembly completeness (right) with black lines representing non-contiguous assembly of that region (**Methods**). Numbers on the right indicate the number of Y contigs needed to achieve the indicated contiguity/total number of assembled Y contigs for each sample). CEN - centromere - includes the *DYZ3* α-satellite array and the pericentromeric region. Three contiguously assembled Y chromosomes are in bold and marked with an asterisk (assemblies for HG02666 and HG00358 are contiguous from telomere to telomere, while HG01890 assembly has a break approximately 100 kbp before the end of PAR2) and the T2T Y for HG002 is in bold and underlined. The colour of sample ID corresponds to the superpopulation designation (see panel **d**). Note - GRCh38 Y sequence mostly represents haplogroup R1b. **c.** The proportion of contiguously assembled Y-chromosomal subregions across 43 samples. **d.** Geographic origin and sample size of the included 1000 Genomes Project samples coloured according to the continental groups (AFR, African; AMR, American; EUR, European; SAS, South Asian; EAS, East Asian). **e.** Y-chromosomal assembly length vs. number of Y contigs. Gap sequences (N’s) were excluded from GRCh38. **f.** Y-chromosomal assembly length vs. Y contig NG50. High coverage defined as >5011 genome-wide PacBio HiFi read depth. Gap sequences (N’s) were excluded from GRCh38.

Recent attempts have been made to assemble the human Y chromosome using Illumina short-read^7^ and Oxford Nanopore Technologies (ONT) long-read data^8^, but a contiguous assembly of the ampliconic and heterochromatic regions was not achieved. In April 2022, the first complete *de novo* assembly of a human Y chromosome was reported by the Telomere-to-Telomere (T2T) Consortium^9^ (from individual HG002/NA24385, carrying a rare J1a-L816 Y lineage found among Ashkenazi Jews and Europeans^10^, termed as T2T Y). However, understanding the composition and appreciating the complexity of the Y chromosomes in the human population requires access to assemblies from many diverse individuals. Here, we combined PacBio HiFi and ONT long-read sequence data to assemble the Y chromosomes from 43 males, representing the five continental groups from the 1000 Genomes Project. While both the GRCh38 (mostly R1b-L20 haplogroup) and the T2T Y represent European Y lineages, half of our Y chromosomes constitute African lineages and include most of the deepest-rooted human Y lineages. This newly assembled dataset of 43 Y chromosomes thus provides a more comprehensive view of genetic variation, at the nucleotide level, across over 180,000 years of human Y chromosome evolution.

## Results

### Sample Selection

We selected 43 genetically diverse males from the 1000 Genomes Project, representing 21 largely African haplogroups (A, B and E, including deep-rooted lineages A0b-L1038, A1a-M31 and B2b-M112)^11,12^ (**Figs. 1b,d; Fig. S1; Table S1; Methods**). The time to most recent common ancestor (TMRCA) among our 43 Y chromosomes and the T2T Y was estimated to be approximately 183 thousand years ago (kya) (95% highest posterior density [HPD] interval: 160-209 kya) (**Fig. S1**), consistent with previous reports^13,14^. A pair of closely related African Y chromosomes (NA19317 and NA19347, lineage E1b1a1a1a-CTS8030), was included for assembly validation, as these Y chromosomes are expected to be highly similar (TMRCA 200 years ago (ya) [95% HPD interval: 0 - 500 ya]).

### Constructing De Novo Assemblies

We employed the hybrid assembler Verkko^15^ to generate Y chromosome assemblies, including the ampliconic and heterochromatic regions (**Methods**). Verkko leverages the high accuracy of PacBio HiFi reads (>99.8% base-pair calling accuracy^16,17^) with the length of ONT long/ultra-long reads (median read length N50 134 kbp) to produce highly accurate and contiguous assemblies (**Table S2**). Using this approach, we generated high-quality (median QV 48; **Table S3**) whole-genome (median length 5.9 Gbp; **Table S4**) assemblies for the 43 males studied. The chromosome Y sequences exhibit a high degree of completeness (median length 55.6 Mbp, 79% to 148% assembly length relative to GRCh38 Y; **Fig. 1; Fig. S2; Tables S5-S6**), contiguity (median NG50 9.6 Mbp, median LG50 2, **Table S4**), base-pair quality (median QV 46, **Table S3**), and read-depth profile consistency with the autosomal sequences in the assemblies (**Fig. S3, Table S7**). The Verkko assembly process was robust (sequence identity for NA19317/NA19347 pair of 99.9959%, **Fig. S4**; **Table S8; Supplementary Results ‘*De novo* assembly evaluation’**). We generated a gapless Y chromosome assembly, spanning from PAR1 to PAR2, for three individuals, two of which represent deep-rooted African haplogroups (**Figs. 1b**, **2; Table S9**). These three samples are among nine samples with an increased HiFi coverage of at least 50⨉ (“high-coverage samples”; **Tables S1-S2, S7**).

**Figure 2.**
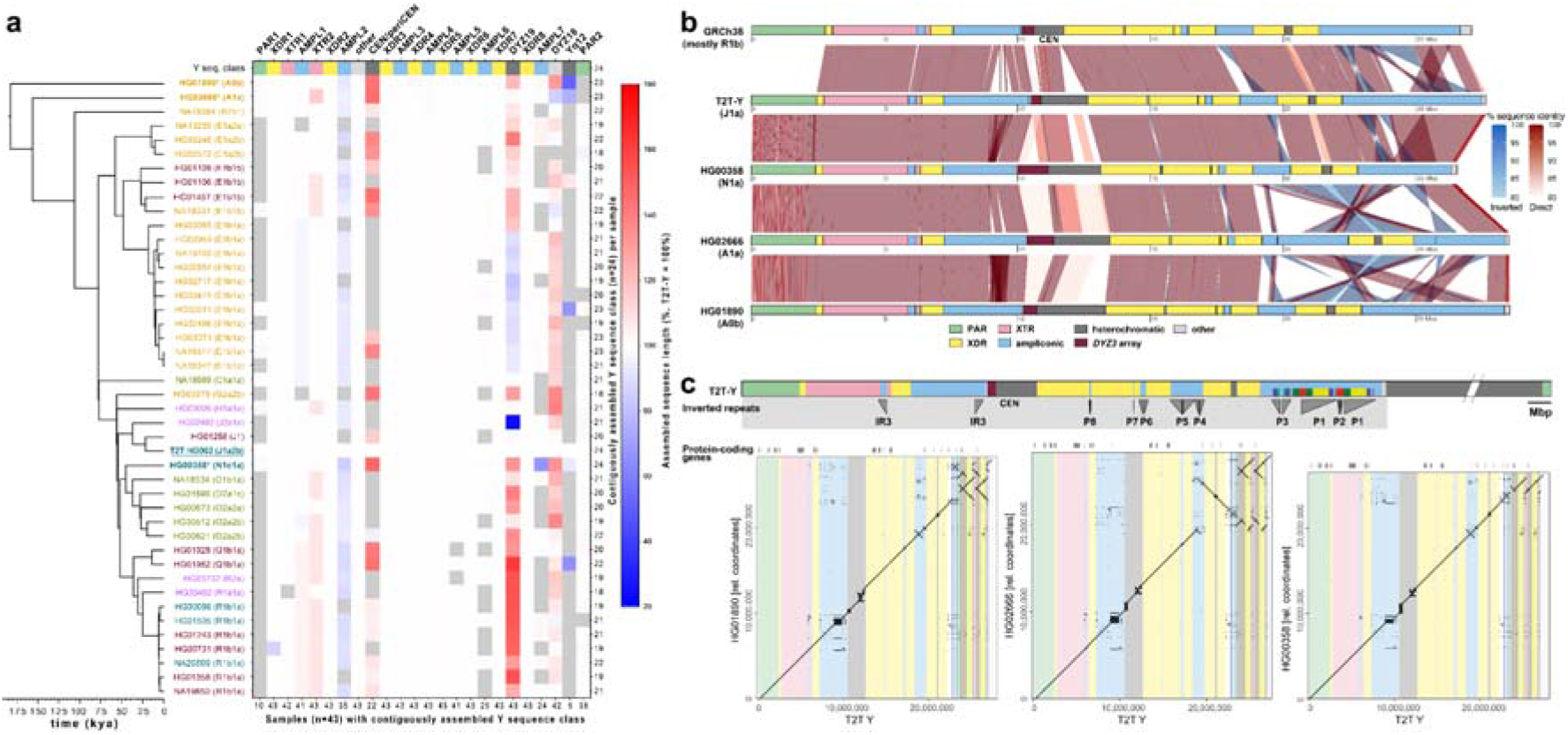
Size and structural variation of Y chromosomes. **a.** Size variation of contiguously assembled Y-chromosomal subregions shown as a heatmap relative to the T2T Y size (as 100%). Boxes in grey indicate regions not contiguously assembled (**Methods**). Numbers on the bottom indicate contiguously assembled samples for each subregion out of a total of 43 samples, and numbers on the right indicate the contiguously assembled Y subregions out of 24 regions for each sample. Samples are coloured as on Fig. 1b. **b.** Comparison of the three contiguously assembled Y chromosomes to GRCh38 and the T2T Y (excluding Yq12 and PAR2 subregions). **c.** Dot plots of three contiguously assembled Y chromosomes vs. the T2T Y (excluding Yq12 and PAR2), annotated with Y subregions and segmental duplications in ampliconic subregion 7 (see **Fig. S39** for details).

Following established procedures^18–20^, we flagged potentially erroneous regions, comprising 0.103% (median; mean 0.31%) up to 0.186% (median; mean 0.467%) of the assembled Y sequence (**Fig. S5, Tables S10-S12**; **Methods**). Although the error rate is increased for the lower-coverage assemblies, increasing the HiFi coverage beyond 50⨉ has limited effect on the error rate (**Fig. S6**).

We further annotated each of the Y-chromosomal assemblies with respect to the 24 Y-chromosomal subregions originally proposed by Skaletsky and colleagues (**Figs. 1a-c; Fig. S2; Table S13; Methods**)^3^. In addition to the three gapless Y chromosomes, we contiguously assembled the MSY excluding Yq12 and the (peri-)centromeric region for 17/43 samples (**Tables S9, S14-S16**). Overall, 17/24 subregions were contiguously assembled across 41/43 samples (**Figs. 1b-c; Fig. S2**).

### Genomic and epigenetic variation of assembled Y chromosomes

*Size variation of the assembled Y chromosomes.* The assembled Y chromosomes showed extensive variation both in size and structure (**Figs. 2a-c**, **3a and 4; Fig. S7-S19; Methods**) with chromosome sizes ranging from 45.2 to 84.9 Mbp (mean 57.6 and median 55.7 Mbp, **Fig. S17**; **Table S14, S16; Methods**). This is, however, a slight underestimate of the true Y-chromosomal size due to assembly gaps. An analysis of the underlying assembly graphs suggest that the paths of complete assemblies would be, on average, 1.15% longer (**Table S6**; **Supplementary Results ‘*De novo* assembly evaluation’**). Among the gaplessly assembled Y-chromosomal subregions (including for the T2T Y), the largest variation in subregion size was seen for the heterochromatic Yq12 (17.6 to 37.2 Mbp, mean 27.6 Mbp), the (peri-)centromeric region (2.0 to 3.1 Mbp, mean 2.6 Mbp) and the *DYZ19* repeat array (63.5 to 428 kbp, mean 307 kbp) (**Figs. 2a**, **4f; Figs. S7, S17-S23; Tables S14-S16**).

**Figure 3.**
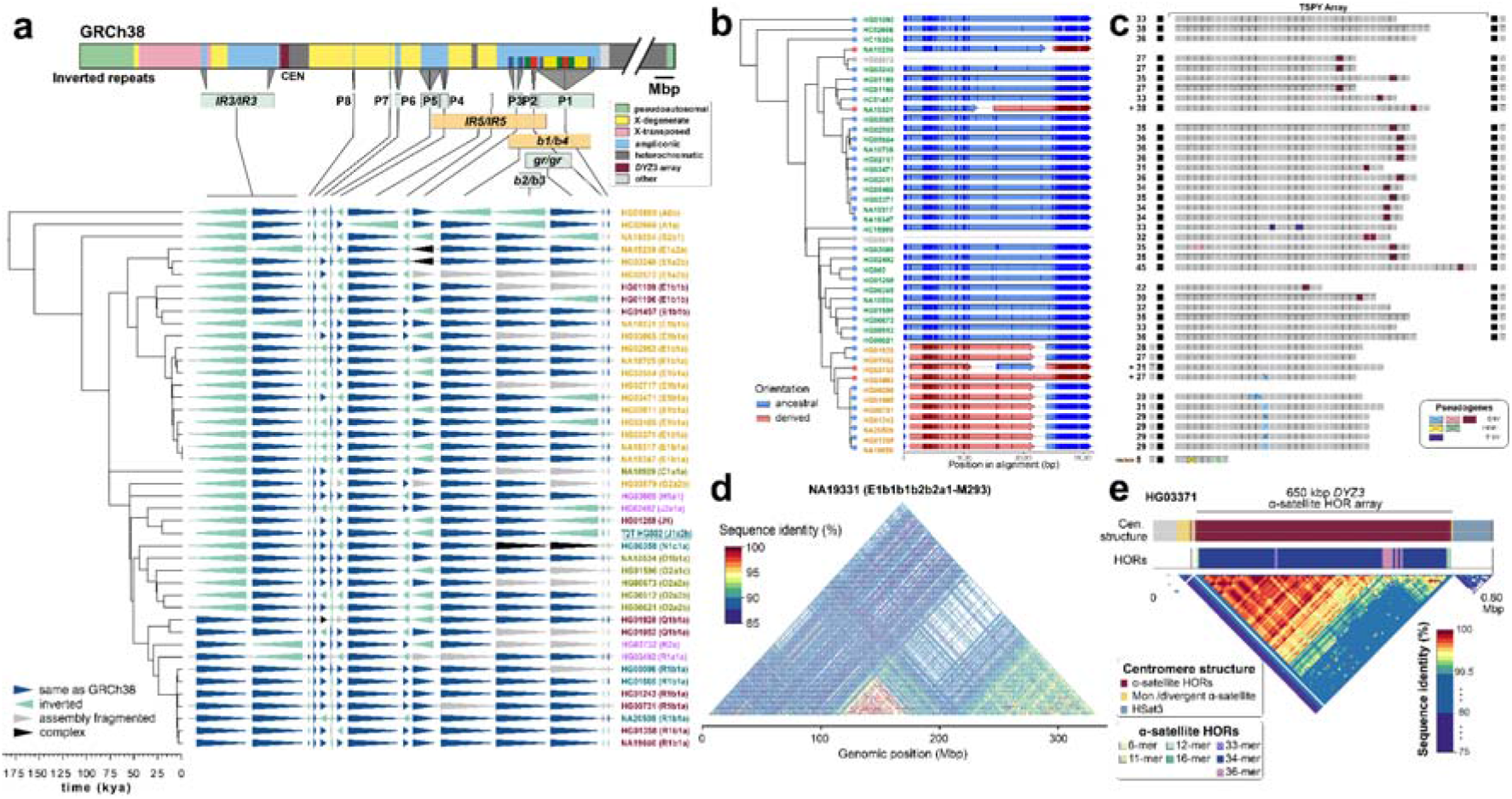
Characterization of large SVs. **a.** Distribution of 14 euchromatic inversions in phylogenetic context, with the schematic of the GRCh38 Y structure shown above, annotated with Y subregions, inverted repeat locations, palindromes (P1-P8), and segmental duplications in ampliconic subregion 7 (see **Fig. S39** for details). Inverted segments are indicated below as green (recurrent) and orange (singleton events) boxes. Samples are coloured the same as in Fig. 1b. **b.** Inversion breakpoint identification in the IR3 repeats. Sample names in orange have undergone two inversions (**Fig. S49, Supplementary Results ‘Y- chromosomal Inversions’**). The red tip colours in the phylogenetic tree indicate samples that have undergone an additional inversion and therefore carry the region between IR3 repeats in inverted orientation compared to samples with blue tip. Informative PSV positions are shown as vertical lines with darker colour in each of the arrows. The orange dotted line indicates the start of the unique ‘spacer’ region. Any information that is not available is indicated by grey. **c.** Distribution of identified pseudogenes within the TSPY array. Numbers to the left indicate the total number of low divergence (≤2%) *TSPY* genes within the array, asterisks (*) indicate samples that due to the *IR3/IR3* inversions (see panel **b**) have the TSPY array in reverse orientation (reoriented for visualization), canonical protein-coding copies are shown in grey, additional pseudogenes (high divergence copies, >2%) located outside of the array are shown in black (note: a high divergence pseudogene located at the end of all arrays is not shown), and the coloured boxes are low divergence (≤2%) pseudogenes originating from six different events: three caused by nonsense mutations, two by a one nucleotide indel deletion, and one by a structural variation (in NA18989). The structural variation deletes the 5’ region of the *TSPY* gene copy (*∼*370 nucleotides of the proximal half of exon 1). Refer to panel **b** for sample IDs and phylogenetic relationships. **d.** Sequence identity heatmap of the *DYZ19* repeat array from NA19331 (using 1 kbp window size) highlighting the higher sequence similarity within central and distal regions. **e.** Genetic landscape of the chromosome Y centromeric region from HG03371 (E1b1a1a1a1c1a-CTS1313). This centromere harbours the ancestral 36-monomer higher-order repeats (HORs), from which the canonical 34-monomer HOR is derived (**Fig. S58**).

**Figure 4.**
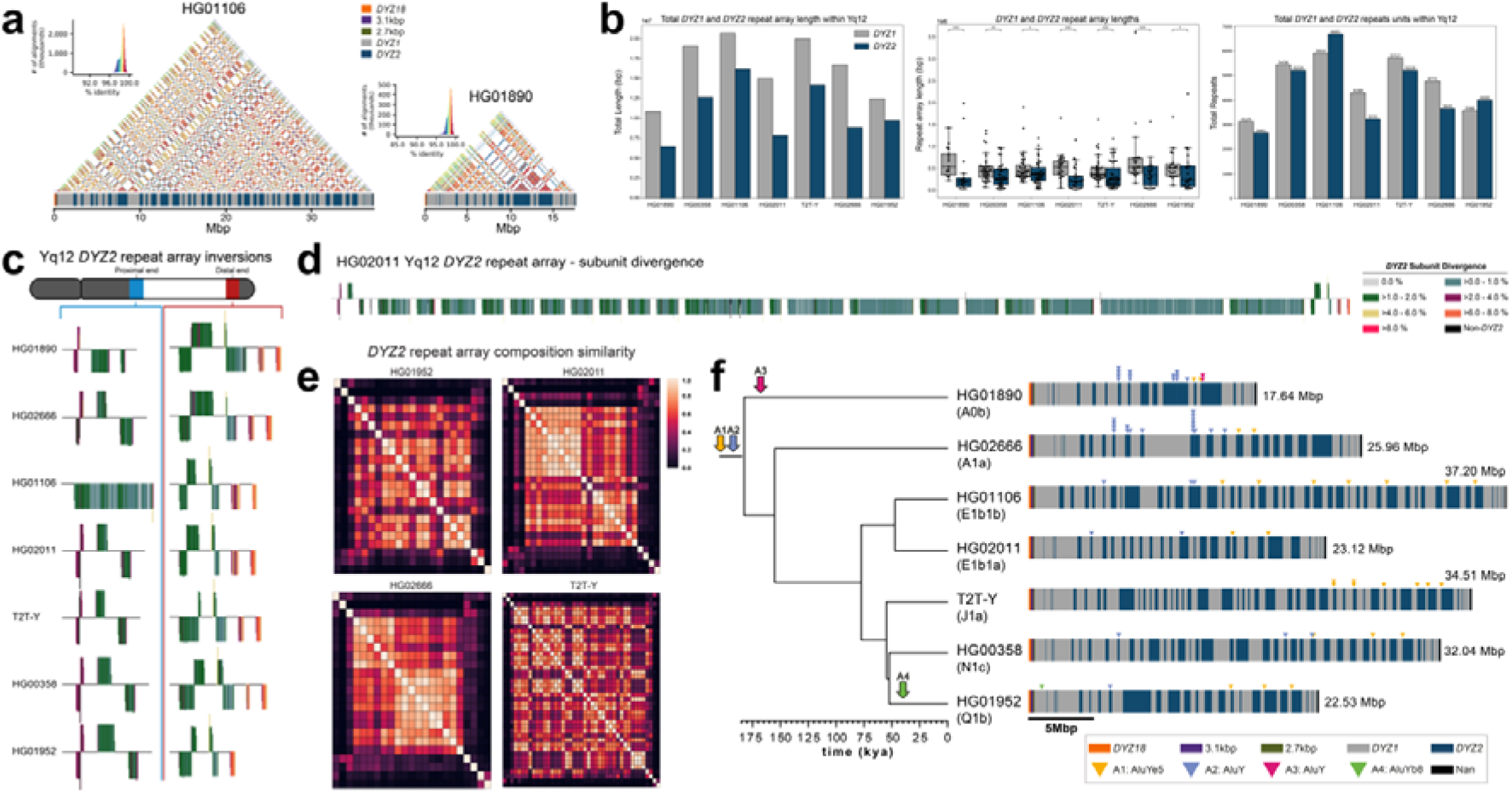
Yq12 heterochromatic region. **a.** Yq12 heterochromatic subregion sequence identity heatmap in 5 kbp windows for HG01106 and HG01890 with repeat array annotations. **b.** Bar plot of the total length of *DYZ1* and *DYZ2* repeat arrays for each sample (left), boxplots of individual array lengths (middle) and the total number of *DYZ1* and *DYZ2* repeat units (right) within contiguously assembled genomes. Black dots represent individual arrays and asterisks (*) denote a statistically significant difference between *DYZ1* and *DYZ2* array lengths (two-sided Mann-Whitney U test: p-value <□0.05, alpha=0.05, Methods). **c.** *DYZ2* repeat array inversions in the proximal and distal ends of the Yq12 region. *DYZ2* repeats are coloured based on their divergence estimate (see panel d) and visualized based on their orientation (sense - up, antisense - down). **d.** Detailed representation of *DYZ2* subunit divergence estimates for HG02011. Length of each line is a function of the subunit length. Orientation (sense - up, antisense - down). **e.** Heatmaps showing the inter-*DYZ2* repeat array subunit composition similarity within a sample. Similarity is calculated using the Bray-Curtis index (1 – Bray-Curtis Distance, 1.0 = the same composition). *DYZ2* repeat arrays are shown in physical order from proximal to distal (from top down, and from left to right). **f.** Mobile element insertions identified in the Yq12 subregion. We identified four putative *Alu* insertions across the seven gapless Yq12 assemblies. Their approximate location, as well as expansion and contraction dynamics of *Alu* insertion-containing *DYZ* repeat units, are shown (right). Following the insertion into *DYZ* repeat units, lineage-specific contractions and expansions occurred. Two *Alu* insertions (A1, and A2) occurred prior to the radiation of Y haplogroups (at least 180,000 years ago). Two additional *Alu* elements represent lineage-specific insertions. Based on these patterns, we can ascertain that arrays and/or repeat units with the same *Alu* insertion are related to each other. While the intra-repeat array expansions may be caused by replication slippage, NAHR may cause both intra- and inter-array expansion^42,43^, although gene conversion can not be excluded. The total length of the Yq12 region is indicated on the right.

The euchromatic regions showed comparatively little variation in size (**Fig. 2a; Fig. S7, Tables S14, S16)** with exception of the ampliconic subregion 2 that contains a copy-number variable TSPY repeat array, composed of 20.3 kbp repeat units. The TSPY array size varies by up to 467 kbp between individuals (**Figs. S19, S24; Tables S15-S18; Supplementary Results ‘Gene family architecture and evolution’; Methods**) and was consistently shorter among males within haplogroup QR (from 567 to 648 kbp, mean 603 kbp) compared to males in the other haplogroups (from 465 to 932 kbp, mean 701 kbp) (**Figs. S19, S24-S27**). The concordance of observed size variation with the phylogeny is well supported by relatively constant, phylogenetically-independent contrasts (PICs), across the phylogeny (**Figs. S25-S26; Methods**). Such phylogenetic consistency reinforces the high quality of our assemblies even across homogeneous tandem arrays, as more closely related Y chromosomes are expected to be more similar, and this consequently allows investigation of mutational dynamics across well-defined timeframes.

#### Distribution and frequency of genetic variants

We leveraged our assemblies to produce a set of variant calls for each Y chromosome, including structural variants (SVs), indels, and single-nucleotide variants (SNVs). In the MSY, we report on average 88 insertion and deletion SVs (≥50 bp), three large inversions (>1 kbp), 2,168 indels (<50 bp), and 3,228 SNVs per Y assembly (**Fig. S28; Table S19; Methods**) when compared to the GRCh38 Y reference. Variants were merged across all 43 samples to produce a nonredundant callset of 876 SVs (488 insertions, 378 deletions, 10 inversions), 23,459 indels (10,283 insertions, 13,176 deletions), and 53,744 SNVs (**Tables S20-S24; Supplementary Results ‘Orthogonal support to Y-chromosomal SVs and copy number variation’**). Based on SV insertions, we identified an average of 81 kbp (range of 46 to 155 kbp) of novel, non-reference sequences per Y chromosome. After excluding simple repeats and mobile element sequences, an average of 18 kbp (range of 0.6 to 47 kbp) of unique non-reference sequence per Y remained (**Table S25**).

Across the unique regions of the autosomes, we find 1.91 SVs, 165.66 indels, and 994.42 SNVs per Mbp per haplotype (**Table S26, Methods**). In the PAR1 region, on both the X and the Y chromosome, SV rates increased 1.98-fold to 3.79 SVs per Mbp (*p* = 3.38×10^-6^, Welch’s t-test) per haplotype, indels increased 1.56-fold to 259.14 per Mbp (*p* = 1.97×10^-4^, Welch’s t-test), and SNVs decreased slightly to 936.19 (*p* = 0.18, Welch’s t-test) across unique loci. While PAR1 has the same ploidy as the autosomes, it is much shorter (2.8 Mbp) and has a 10× increased recombination rate in males compared to females^21^, which may lead to the observed higher density of SVs and indels. A reduced level of variation observed in the proximal 500 kbp before the currently-established PAR1 boundary could be the result of a lower recombination rate closer to the sex-specific chromosomal regions, indicating a more distal location for the actual PAR1 boundary (**Figs. S29-S32**). As expected, the human chromosome X (excluding both PAR regions) exhibits lower genetic variation with 1.16 SVs (*p* = 1.54×10^-26^ Student’s t-test), 106.64 indels (*p* = 1.33×10^-46^, Student’s t-test), and 584.93 SNVs (*p* = 1.18×10^-83^, Student’s t-test) per Mbp of unique loci with most differences likely attributed to a lower effective population size for the X chromosome. The MSY has even less variation than seen for the X chromosome, with an average of 0.01 SVs, 2.11 indels, and 5.72 SNVs per Mbp (*p* < 1×10^-100^ for all, Welch’s t-test) of unique loci (**Table S26**).

We also identified 25 transposable element insertions in the 43 Y-chromosomal assemblies that are not present in the GRCh38 Y, including 18 *Alu* elements (4/18 within the Yq12) and seven LINE-1s (long interspersed element-1; no significant difference compared to the whole-genome distribution reported in^19^) (**Fig. 4f**; **Tables S27-S28**; **Methods; Supplementary Results ‘Yq12 heterochromatic subregion’**). Closer inspection across the three gaplessly assembled Y chromosomes, as well as the T2T Y chromosome, showed substantial differences in repeat composition between Y-chromosomal subregions (**Fig. S33; Tables S29-S30**). For example, the pseudoautosomal regions showed a clear increase in SINE (short interspersed element) content and reduction in LINE and LTR (long terminal repeat) content compared to the male-specific XTR, XDR, and ampliconic regions (**Figs. S33-S34**; **Table S31**).

#### Y-chromosomal inversions

Large inversions were identified using Strand-seq^22^ and manual inspection of assembly alignments, yielding as many as 14 inversions in the euchromatic regions and two inversions within the Yq12 across the studied males (**Figs. 3a**, **4c; Figs. S35-S36; Tables S32-S34**; **Methods; Supplementary Results ‘Y-chromosomal Inversions’**). Six of these matched the ten inversions identified above by variant calling (**Table S22**). The breakpoint intervals for 8/14 of the euchromatic inversions were refined to DNA regions as small as 500 bp (**Fig. 3b**; **Fig. S36-38; Table S35**; **Methods**). All these inversions are flanked by highly similar (up to 99.97%) and large (from 8.7 kbp to 1.45 Mbp) inverted SDs, and while determination of the molecular mechanism generating Y-chromosomal inversions remains challenging, most are likely a result of non-allelic homologous recombination (NAHR). Moreover, we found that most (12/14, 85%) euchromatic inversions are recurrent, with 2 to 13 toggling events in the Y phylogeny which translates to an inversion rate estimate ranging from 3.68 ⨉ 10^-5^ (95% C.I.: 3.25 - 4.17 ⨉ 10^-5^) to 2.39 ⨉ 10^-4^ (95% C.I.: 2.11 - 2.71 ⨉ 10^-4^) per locus per father-to-son Y transmission. The highest inversion recurrence is seen among the eight Y-chromosomal palindromes (called P1-P8, **Fig. 3a; Fig. S39; Table S32, Methods**). Taken together, we calculate a rate of one recurrent inversion per 603 (95% C.I.: 533 - 684) father-to-son Y transmissions. The per site per generation rate estimates for 12 Y-chromosomal recurrent inversions are significantly higher (>2-fold difference between median estimates, two-tailed Mann-Whitney-Wilcoxon test, n=44, p-value <0.0001) than the rates previously estimated for 32 autosomal and X-chromosomal recurrent inversions^23^.

There are two fixed inversions flanking the Yq12 subregion (**Fig. 4c; Fig. S40; Table S34; Supplementary Results ‘Y-chromosomal Inversions’**). The proximal inversion, observed in 10/11 individuals analyzed, ranged from 358.9 to 820.7 kbp in size (mean 649.0 kbp) (**Table S34**). The distal inversion, on the other hand, was observed in all 11 individuals and ranged from 259.5 to 641.4 kbp in size (mean 472.5 kbp). We found the breakpoints for these two inversions to be identical among all individuals. This suggests that the consistent presence of these two inversions at both ends of the Yq12 subregion may prevent unequal sister chromatid exchange from occurring, restricting expansion and contraction of the repeat units to the region flanked by these two inversions.

#### Evolution of palindromes and multi-copy gene families

To further reconstruct the evolution of Y-chromosomal palindromes, we investigated both the gene conversion patterns and evolutionary rates across the Y assemblies (**Figs. S41-S43**; **Tables S15, S36-S37**). The intra-arm gene conversion patterns across seven palindromes (P1, P3-P8, but excluding the P2 palindrome and DNA sequences that are shared between different palindromes; **Methods**) showed a significant bias towards G or C nucleotides (942 events to G or C vs. 701 to A or T nucleotides, p=2.75 x 10^-9^, Chi-square test), but no bias towards the ancestral state (357 events to derived vs. 374 to ancestral state; P=0.5295, Chi-square test; **Table S36**). Comparison of base substitution patterns for all eight palindromes and the eight XDR regions, across 13 Y chromosomes, indicated that different palindromes are evolving at different rates (**Methods**). The level of sequence variation (both in base substitutions and SVs) and estimated base substitution mutation rates were higher for palindromes P1, P2 and P3 which contain higher proportions of multicopy (i.e., >2 copies) segments compared to other palindromes (3.03 x 10^-8^ (95% C.I.: 2.80 - 3.27 x 10^-8^) vs. 2.12 x 10^-8^ (95% C.I.: 1.96 - 2.29 x 10^-8^) mutations per position per generation, respectively) (**Fig. 3a; Figs. S39, S41-S43; Table S37; Methods**). The increased variation of P1, P2 and P3 likely results from sequence exchange between multicopy regions.

The gene annotation of the Y-chromosomal assemblies showed no evidence of the loss of any MSY protein-coding genes in the 43 males analysed (**Tables S38-S42; Supplementary Results ‘Gene annotation’**). However, the investigation of three copy-number variable ampliconic gene families (*DAZ* (Deleted in Azoospermia)*, TSPY* (testis specific protein Y-linked 1), *RBMY1* (RNA-binding motif (RRM) gene on Y chromosome), **Supplementary Results ‘Gene family architecture and evolution’)** revealed substantial differences in their genetic diversity and evolution. While only two out of 43 samples (41 assemblies, T2T Y and GRCh38 Y) showed a difference in the *DAZ* copy number (two and six *DAZ* copies vs. four in all others), extensive variation was detected in the copy number of the 28 canonical exons (from 0 to 14 copies of a single exon) between samples **(Fig. S44; Tables S15, S43-S44, Methods)**. Consistent with previous reports, *RBMY1* genes were primarily located in four separate regions, while three samples had undergone larger rearrangements (**Figs. S45-S46; Supplementary Results** ‘**Gene family architecture and evolution**’)^24^. On average, eight *RBMY1* gene copies (from 5 to 11) were identified, with most of the variation caused by expansions or contractions in regions 1 and 2 (**Figs. S45-S46; Table S28**). A phylogenetic analysis of *RBMY1* genes revealed that the gene copies from regions 3 and 4 have likely given rise to *RBMY1* genes located in regions 1 and 2 (**Figs. S47-S48)**, additionally supported by the analysis of the chimpanzee (PanTro6) *RBMY1* sequences (**Supplementary Results** ‘**Gene family architecture and evolution**’).

The majority of the *TSPY* genes are located in a tandemly organized and highly copy-number variable TSPY array. While a single repeat unit containing the *TSPY2* gene is located upstream of the TSPY array in GRCh38, we inferred that the ancestral position of *TSPY2* is in between the TSPY repeat array and the Y centromere in reverse orientation (**Figs. S49-S50**; **Table S35**; **Supplementary Results sections ‘Y-chromosomal inversions’** and **‘Gene family evolution’**). Likely the result of two inversions or a complex rearrangement, the localization of TSPY2 upstream of the TSPY array is shared by all QR haplogroup (including the GRCh38 Y) individuals. On average, 33 *TSPY* gene copies (from 23 to 39, 46 in T2T Y, counts include low divergence (≤2%) *TSPY* gene copies from the TSPY repeat array and *TSPY2*) were identified per assembly (**Fig. 3c**, **Figs. S50-S51, Tables S15, S39**). Both network and phylogenetic analysis of *TSPY* gene sequences support identification of the ancestral gene copy (medium blue in **Figs. S50** and **S52**; **Methods**). Notably, six independently arisen pseudogenes were identified among the *TSPY* genes located within the 41 TSPY arrays analysed (39 contiguous assemblies, the T2T Y and the GRCh38 Y), with 31/41 samples carrying at least one pseudogene (**Fig. 3c**). The phylogenetic distribution suggests periodic purging of pseudogenes from the array, possibly through the removal of deleterious mutations by gene conversion. Evidence of NAHR and gene conversions was found both between the tandemly repeated *TSPY* gene copies and the *RBMY1* genes (**Figs. S45, S50)**.

#### Epigenetic variation

The ONT sequencing data also provide a means to explore the base-level epigenetic landscape of the Y chromosomes (**Fig. S53**). Here, we focused on DNA methylation at CpG sites, hereafter referred to as DNAme. In 41 samples (EBV-transformed lymphoblastoid cell lines) that passed QC (**Methods**), we first tested the association of chromosome Y assembly length on global DNAme levels as has previously been shown in Drosophila^25^. We detected a significant relationship between the chromosome Y assembly length on global DNAme levels, both genome wide and for the Y chromosome (linear model p=0.0477 and p=0.0469 (n=41); **Fig. S54; Supplementary Results ‘Functional analysis’**). We found 2,861 DNAme segments that vary across these Y chromosomes (**Fig. S55; Table S45**). Of note, 21% of the variation in DNAme levels is associated with haplogroups (Permanova p=0.003, n=41), while the same is true for only 4.8% of the expression levels (Permanova p=0.005, n=210, leveraging the Geuvadis RNA-seq expression data^26^; **Methods**). This association is particularly significant for five genes (*BCORP1* (**Fig. S56**), *LINC00280*, *LOC100996911*, *PRKY*, *UTY*), where both DNAme and gene-expression effects are observed (**Tables S45, S56**). Lastly, we find 194 Y-chromosomal genetic variants, including a 171 base-pair insertion and one inversion, that impact DNAme levels on chromosome Y (**Table S46**; **Supplementary Results ‘Functional analysis’**). This suggests that some of the genetic background, either on the Y chromosome or elsewhere in the genome, may impact the functional outcome (the epigenetic and transcriptional profiles) of specific genes on the Y chromosome.

### Genetic variation and evolution of the Y-chromosomal heterochromatic regions

#### Variation in the size and structure of centromeric/pericentromeric repeat arrays

In general, the chromosome Y centromeres are composed of 171 bp *DYZ3* α-satellite repeat units^3^, organized into a higher-order repeat (HOR) array, flanked on either side by short stretches of monomeric α-satellite. The α-satellite HOR arrays across gaplessly assembled Y centromeres ranged in size from 264 kbp to 1.165 Mbp (mean 671 kbp), with smaller arrays found in haplogroup R1b samples compared to other lineages (mean 341 kbp vs. 787 kbp, respectively; **Figs. S19, S24-S26, S57; Tables S15-16**; **Methods**)^27,28^. We determined that the *DYZ3* α-satellite HOR array is mostly composed of a 34-monomer repeating unit that is the most prevalent HOR type found in the 21 analysed samples **Fig. 3e, Fig. S58; Methods**). However, we identified two other HORs that were present at high frequency among the analysed Y chromosomes: a 35-monomer HOR found in 14/21 samples and a 36-monomer HOR found in 11/21 samples (**Methods**). While the 35-monomer HOR is present across different Y lineages in the Y phylogeny, the 36-monomer HOR has been lost in phylogenetically closely related Y chromosomes representing the QR haplogroups (**Fig. S57)**. Analysis of the sequence composition of these HORs revealed that the 36-monomer HOR likely represents the ancestral state of the canonical 35-mer and 34-mer HOR after deletion of the 22nd α-satellite monomer in the resulting HORs, respectively (**Fig. S58; Methods**).

The overall organization of the *DYZ3* α-satellite HOR array is similar to that found on other human chromosomes, with near-identical α-satellite HORs in the core of the centromere that become increasingly divergent towards the periphery^29–32^. There is a directionality of the divergent monomers at the periphery of the Y centromeres such that a larger block of diverged monomers is consistently found at the p-arm side of the centromere compared to the block of diverged monomers juxtaposed to the q-arm.

Adjacent to the *DYZ3* α-satellite HOR array on the q-arm is an *HSat3* repeat array which ranges in size from 372 to 488 kbp (mean 378 kbp), followed by a *DYZ17* repeat array, which ranges in size from 858 kbp to 1.740 Mbp (mean 1.085 Mbp). Comparison of the sizes of these three repeat arrays reveals no significant correlation among their sizes (**Fig. 3e; Figs. S59-S61; Table S15-S16**).

The *DYZ19* repeat array is located on the long arm, flanked by XDRs (**Fig. 1a**) and composed of 125 bp repeat units (fragment of an LTR) in head-to-tail fashion. This subregion was completely assembled across all 43 Y chromosomes and exhibits the highest variation with a 6.7-fold difference in size (from 63.5 to 428 kbp). The HG02492 individual (haplogroup J2a) with the smallest-sized *DYZ19* repeat array has an approximately 200 kbp deletion in this subregion (**Table S16**). In 43/44 Y chromosomes (including T2T Y), we find evidence of at least two rounds of mutation/expansion (**Fig. 3d**, green and red coloured blocks, respectively, **Figs. S21-S23**), leading to directional homogenization of the central and distal parts of the region in all Y chromosomes. Finally, we observed a recent ∼80 kbp duplication event shared by the 11 phylogenetically related haplogroup QR samples (**Figs. S21-S23**), which must have occurred approximately 36,000 years ago (**Figs. 1b, S1**), resulting in a substantially larger overall *DYZ19* subregion in these Y chromosomes.

Between the Yq11 and the Yq12 subregions lies the *DYZ18* subregion which comprises three distinct repeat arrays: a *DYZ18* repeat array and novel 3.1 kbp and 2.7 kbp repeat arrays (**Figs. S62-S69**). The 3.1 kbp repeat array is composed of degenerate copies of the *DYZ18* repeat unit, exhibiting 95.8% sequence identity (using SNVs only) across the length of the repeat unit. The 2.7 kbp repeat array seems to have originated from both the *DYZ18* (23% of the 2.7 kbp repeat unit shows 86.3% sequence identity to *DYZ18*) and *DYZ1* (77% of the 2.7 kbp repeat unit shows 97% sequence identity to *DYZ1*) repeat units (**Fig. S62**). All three repeat arrays (*DYZ18*, 3.1 kbp and 2.7 kbp) show a similar pattern and level of methylation compared to the *DYZ1* repeat arrays (**Fig. S70**), in that we observe constitutive hypermethylation.

#### Composition of the Yq12 heterochromatic subregion

The Yq12 subregion is the most challenging portion of the Y chromosome to assemble contiguously due to its highly repetitive nature and size. In this study, we completely assembled the Yq12 subregion for six individuals and compared it to the Yq12 subregion of the T2T Y chromosome (**Figs. 1a**, **4a,f; Tables S14-S16; Supplementary Results ‘Yq12 heterochromatic subregion’**). This subregion is composed of alternating arrays of repeat units: *DYZ1* and *DYZ2*^3,5, 33–36^. The *DYZ1* repeat unit is approximately 3.5 kbp and consists mainly of simple repeats and pentameric satellite sequences, and has been recently referred to as HSat3A6^4^. The *DYZ2* repeat (which has been recently referred to as HSat1B^31^) is approximately 2.4 kbp and consists mainly of a tandemly repeated AT-rich simple repeat fused to a 5’ truncated *Alu* element followed by an HSATI satellite sequence (**Fig. S62**).

The *DYZ1* repeat units are tandemly arranged into larger *DYZ1* repeat arrays, as are the *DYZ2* repeat units, and the *DYZ1* and *DYZ2* repeat arrays alternate with one another (**Fig. 4**). The total number of *DYZ1* and *DYZ2* arrays (range from 34 to 86, mean 61) were significantly positively correlated (Spearman correlation=0.90, p-value=0.0056, n=7, alpha=0.05) with the total length of the analysed Yq12 region (**Fig. S71**), whereas the length of the individual *DYZ1* and *DYZ2* repeat arrays were found to be widely variable (**Fig. 4b; Fig. S72**). The *DYZ1* arrays were significantly longer (range from 50,420 to 3,599,754 bp, mean 535,314 bp) than the *DYZ2* arrays (range from 11,215 to 2,202,896 bp, mean 354,027 bp, two-tailed Mann-Whitney U test (n=7) p-value < 0.05) (**Fig. 4b**); however, the total number of each repeat unit was nearly equal within each Y chromosome (*DYZ1* to *DYZ2* ratio ranges from 0.88 to 1.33, mean 1.09) (**Fig. 4b; Table S47**). From ONT data, we observed a consistent hypermethylation of the *DYZ2* repeat arrays compared to the *DYZ1* repeat arrays, the sequence composition of the two repeats is markedly different in terms of CG content (24% *DYZ2* versus 38% *DYZ1*) and number of CpG dinucleotides (1 CpG/150 bp *DYZ2* versus 1 CpG/35 bp *DYZ1*) potentially explaining the marked DNA methylation differences (**Fig. S53**).

Sequence analysis of the repeat units in the Yq12 suggests that the *DYZ1* and *DYZ2* repeat arrays and the entire Yq12 subregion may have evolved in a similar manner, and similarly to the centromeric region (see above). Specifically, repeat units near the middle of a given array showed a higher level of sequence similarity to each other than to the repeat units at the distal regions of the repeat array (**Fig. 4d; Fig. S73**). This suggests that expansion and contraction tends to occur in the middle of the repeat arrays, homogenizing these units yet allowing divergent repeat units to accumulate towards the periphery. Similarly, when looking at the entire Yq12 subregion, we observed that repeat arrays located in the middle of the Yq12 subregion tend to be more similar in sequence to each other than to repeat arrays at the periphery (**Fig. 4e; Figs. S73-S74**). This observation is supported by results from the *DYZ2* repeat divergence analysis, the inter-*DYZ2* array profile comparison, and the construction of a *DYZ2* phylogeny (**Fig. S75**; **Methods**).

## Discussion

The mammalian Y chromosome has been notoriously difficult to assemble owing to its extraordinarily high repeat content. Here, we present the Y-chromosomal assemblies of 43 males from the 1000 Genomes Project dataset and a comprehensive analysis of their genetic and epigenetic variation and composition. While both the GRCh38 Y and the T2T Y assemblies represent relatively recently emerged (TMRCA 54.5 kya [95% HPD interval: 47.6 - 62.4 kya], **Fig. S1**) European Y lineages, half of our Y chromosomes carry African Y lineages, including two of the deepest-rooted human Y lineages (A0b and A1a, TMRCA 183 kya [95% HPD interval: 160-209 kya]), which we gaplessly assembled allowing us to investigate how the Y chromosome has changed over 180,000 years of human evolution.

For the first time, we were able to comprehensively and precisely examine the extent of genetic variation down to the nucleotide level across multiple human Y chromosomes. The male-specific portion of the Y chromosome (MSY) can be roughly divided into two portions: the euchromatic and the heterochromatic regions. The single-copy protein-coding MSY genes, present in the GRCh38 Y reference sequence, are conserved in all 43 Y assemblies with few SNVs. The low SNV diversity in Y is concordant with previous studies and consistent with models of natural demographic processes such as extreme male-specific bottlenecks in recent human history and purifying selection removing deleterious mutations and linked variation^11,13,37^. The multi-copy protein-coding MSY genes are often copy number variable. We found that 5/8 multi-copy gene families showed variation in terms of copy number, with the highest variation observed in the *TSPY* gene family (23 to 39 copies, 46 in the T2T Y, **Fig. 3c**; **Table S39**). Investigation of three copy-number variable gene families (*TSPY*, *RBMY1* and *DAZ*) revealed different modes of evolution, likely resulting from differences in structural composition of the genomic regions. For example, the majority of the *TSPY* genes are located within a tandemly repeated array, undergoing frequent expansions and contractions, where we also find evidence of lineage-specific acquisition and purging of pseudogenes.

The euchromatic region harbours additional structural variation across the 43 individuals. Most notably, we identified 14 inversions that together affect half of the Y-chromosomal euchromatin, with only the most closely related pair of African Ys (from NA19317 and NA19347) showing the exact same inversion composition. Of these 14 inversions, 12 showed recurrent toggling in recent human history, including five novel recurrent inversions that were not previously reported^23^. We narrowed down the breakpoints for all of the inversions and have refined the breakpoints down to a 500 bp region for 8 of 14 inversions. The determination of the molecular mechanism causing the inversions remains challenging; however, the increased recurrent inversion rate on the Y chromosome compared to the rest of the human genome may be in part due to DNA double-strand breaks being repaired by intra-chromatid recombination^23,38^. The enrichment in highly similar (inverted) SDs^2,3^ prone to NAHR, coupled with reduced selection to maintain gene order, may explain the high prevalence of recurrent inversions on the Y. The majority of the recurrent inversions (8/14) occur between highly similar SDs termed palindromes P1-P8 (**Fig. 3a**). Three of the palindromes appear to be evolving at faster rates compared both to the other five palindromes and the unique XDR regions of the Y, likely due to sequence exchange between multi-copy (i.e., >2 copies) SDs.

In the PAR1 region, we find evidence of enrichment of indels and SVs compared to autosomes, and the rest of the X and the Y chromosomes, potentially resulting from a higher recombination rate in this region during male meiosis^21^. Interestingly, there is a reduction of genetic variation in the proximal 500 kbp of PAR1, indicating a reduced recombination rate here and suggesting that the actual PAR1 boundary probably lies distal to the currently established boundary^3^.

There are four heterochromatic subregions in the human Y chromosome: the (peri-)centromeric region, *DYZ18*, *DYZ19* and Yq12. Heterochromatin is usually defined by the preponderance of highly repetitive sequences and the constitutive dense packaging of the chromatin within^39^. When we examined the DNA sequence and the methylation patterns for these four heterochromatic subregions, the high repetitive sequence content and the high level of methylation (**Figs. S53, S70**) observed is consistent with the definition of heterochromatin. Furthermore, resolving the complete structural variation in the heterochromatic regions of the human Y chromosome provides novel molecular archaeological evidence for evolutionary mechanisms. For example, we show how the higher order structure at the centromeric region of the Y chromosome evolved from an ancestral 36-mer HOR to a 34-mer HOR, which predominates in the centromeres of human males^40^. Moreover, the degeneration of these repeat units of the (peri-)centromeric region of the Y chromosome has a directional bias towards the p-arm side. The presence of an *Alu* element right at the q-arm boundary, but not on the p-arm side, raises the possibility that following two *Alu* insertions, over 180,000 years ago, led to a subsequent *Alu*-*Alu* recombination that deleted the region in between and removed the diverged centromeric sequence block^41^. In the Yq12 subregion, we find evidence for localized expansions and contractions of the *DYZ1* and *DYZ2* repeat units, though the preservation of nearly 1:1 ratio among all males studied indicates functional or evolutionary constraints.

In this study, we fully sequenced and analysed 43 diverse Y chromosomes and identified the full extent of variation of this chromosome across more than 180,000 years of human evolution, offering a major advance to our understanding of how non-recombining regions of the genome evolve and persist. For the first time, sequence-level resolution across multiple human Y chromosomes has revealed new DNA sequences and new elements of conservation, and provided molecular data that give us important insights into genomic stability and chromosomal integrity. It also offers the possibility to investigate the molecular mechanisms and evolution of repetitive sequences across a well-defined timeframe without the encumbrances of meiotic recombination. Ultimately, the ability to effectively assemble the complete human Y chromosome has been a long-awaited yet crucial milestone towards understanding the full extent of human genetic variation and also provides the starting point to associate Y-chromosomal sequences to specific human traits and more thoroughly study human evolution.

## Methods

### 1. Sample selection

Samples were selected from the 1000 Genomes Project Diversity Panel^44^ and at least one representative was selected from each of 26 populations (**Table S1**). A total of 13/28 samples were included from the Human Genome Structural Variation Consortium (HGSVC) Phase 2 dataset, which was published previously^19^. In addition, for 15/28 samples data was newly generated as part of the HGSVC efforts (see the section ‘Data production’ for details’). We also included 15 samples from the Human Pangenome Reference Consortium (HPRC) (**Table S1**). Notably, there is an African Y lineage (A00) older than the lineages in our dataset (TMRCA 254 kya; 95% CI 192-307 kya^13,45^) that we could not include due to sample availability issues.

### 2. Data production

#### a. PacBio HiFi sequence production

##### University of Washington

Sample HG00731 data have been previously described^19^. Additional samples HG02554 and HG02953 were prepared for sequencing in the same way but with the following modifications: isolated DNA was sheared using the Megaruptor 3 instrument (Diagenode) twice using settings 31 and 32 to achieve a peak size of ∼15-20 kbp. The sheared material was subjected to SMRTbell library preparation using the Express Template Prep Kit v2 and SMRTbell Cleanup Kit v2 (PacBio). After checking for size and quantity, the libraries were size-selected on the Pippin HT instrument (Sage Science) using the protocol “0.75% Agarose, 15-20 kbp High Pass” and a cutoff of 14-15 kbp. Size-selected libraries were checked via fluorometric quantitation (Qubit) and pulse-field sizing (FEMTO Pulse). All cells were sequenced on a Sequel II instrument (PacBio) using 30-hour movie times using version 2.0 sequencing chemistry and 2-hour pre-extension. HiFi/CCS analysis was performed using SMRT Link v10.1 using an estimated read-quality value of 0.99.

##### The Jackson Laboratory

High-molecular-weight (HMW) DNA was extracted from 30M frozen pelleted cells using the Gentra Puregene extraction kit (Qiagen). Purified gDNA was assessed using fluorometric (Qubit, Thermo Fisher) assays for quantity and FEMTO Pulse (Agilent) for quality. For HiFi sequencing, samples exhibiting a mode size above 50 kbp were considered good candidates. Libraries were prepared using SMRTBell Express Template Prep Kit 2.0 (Pacbio). Briefly, 12 μl of DNA was first sheared using gTUBEs (Covaris) to target 15-18 kbp fragments. Two 5 µg of sheared DNA were used for each prep. DNA was treated to remove single strand overhangs, followed by DNA damage repair and end repair/ A-tailing. The DNA was then ligated V3 adapter and purified using Ampure beads. The adapter ligated library was treated with Enzyme mix 2.0 for Nuclease treatment to remove damaged or non-intact SMRTbell templates, followed by size selection using Pippin HT generating a library that has a size >10 kbp. The size selected and purified >10 kbp fraction of libraries were used for sequencing on Sequel II (Pacbio).

#### b. ONT-UL sequence production

##### University of Washington

High-molecular-weight (HMW) DNA was extracted from 2 aliquots of 30 M frozen pelleted cells using phenol-chloroform approach as described in^46^. Libraries were prepared using Ultra long DNA Sequencing Kit (SQK-ULK001, ONT) according to the manufacturer’s recommendation. Briefly, DNA from ∼10M cells was incubated with 6 μl of fragmentation mix (FRA) at room temperature (RT) for 5 min and 75°C for 5 min. This was followed by an addition of 5 μl of adaptor (RAP-F) to the reaction mix and incubated for 30 min at RT. The libraries were cleaned up using Nanobind disks (Circulomics) and Long Fragment Buffer (LFB) (SQK-ULK001, ONT) and eluted in Elution Buffer (EB). Libraries were sequenced on the flow cell R9.4.1 (FLO-PRO002, ONT) on a PromethION (ONT) for 96 hrs. A library was split into 3 loads, with each load going 24 hrs followed by a nuclease wash (EXP-WSH004, ONT) and subsequent reload.

##### The Jackson Laboratory

High-molecular-weight (HMW) DNA was extracted from 60 M frozen pelleted cells using phenol-chloroform approach as previously described^47^. Libraries were prepared using Ultra long DNA Sequencing Kit (SQK-ULK001, ONT) according to the manufacturer’s recommendation. Briefly, 50ug of DNA was incubated with 6 μl of FRA at RT for 5 min and 75°C for 5 min. This was followed by an addition of 5 μl of adaptor (RAP-F) to the reaction mix and incubated for 30 min at RT. The libraries were cleaned up using Nanodisks (Circulomics) and eluted in EB. Libraries were sequenced on the flow cell R9.4.1 (FLO-PRO002, ONT) on a PromethION (ONT) for 96 hrs. A library was generally split into 3 loads with each loaded at an interval of about 24 hrs or when pore activity dropped to 20%. A nuclease wash was performed using Flow Cell Wash Kit (EXP-WSH004) between each subsequent load.

#### c. Bionano optical genome maps production

Optical mapping data were generated at Bionano Genomics, San Diego, USA. Lymphoblastoid cell lines were obtained from Coriell Cell Repositories and grown in RPMI 1640 media with 15% FBS, supplemented with L-glutamine and penicillin/streptomycin, at 37°C and 5% CO_2_. Ultra-high-molecular-weight DNA was extracted according to the Bionano Prep Cell Culture DNA Isolation Protocol(Document number 30026, revision F) using a Bionano SP Blood & Cell DNA Isolation Kit (Part #80030). In short, 1.5 M cells were centrifuged and resuspended in a solution containing detergents, proteinase K, and RNase A. DNA was bound to a silica disk, washed, eluted, and homogenized via 1hr end-over-end rotation at 15 rpm, followed by an overnight rest at RT.

Isolated DNA was fluorescently tagged at motif CTTAAG by the enzyme DLE-1 and counter-stained using a Bionano Prep™ DNA Labeling Kit – DLS (catalog # 8005) according to the Bionano Prep Direct Label and Stain (DLS) Protocol(Document number 30206, revision G). Data collection was performed using Saphyr 2nd generation instruments (Part #60325) and Instrument Control Software (ICS) version 4.9.19316.1.

#### d. Strand-seq data generation and data processing

Strand-seq data were generated at EMBL and the protocol is as follows. EBV-transformed lymphoblastoid cell lines from the 1000 Genomes Project (Coriell Institute; **Table S1**) were cultured in BrdU (100 uM final concentration; Sigma, B9285) for 18 or 24 hrs, and single isolated nuclei (0.1% NP-40 substitute lysis buffer^48^ were sorted into 96-well plates using the BD FACSMelody and BD Fusion cell sorter. In each sorted plate, 94 single cells plus one 100-cell positive control and one 0-cell negative control were deposited. Strand-specific single-cell DNA sequencing libraries were generated using the previously described Strand-seq protocol^22,48^ and automated on the Beckman Coulter Biomek FX P liquid handling robotic system^49^. Following 15 rounds of PCR amplification, 288 individually barcoded libraries (amounting to three 96-well plates) were pooled for sequencing on the Illumina NextSeq500 platform (MID-mode, 75 bp paired-end protocol). The demultiplexed FASTQ files were aligned to the GRCh38 reference assembly (GCA_000001405.15) using BWA aligner (version 0.7.15-0.7.17) for standard library selection. Aligned reads were sorted by genomic position using SAMtools (version 1.10) and duplicate reads were marked using sambamba (version 1.0). Low-quality libraries were excluded from future analyses if they showed low read counts (<50 reads per Mbp), uneven coverage, or an excess of ‘background reads’ (reads mapped in opposing orientation for chromosomes expected to inherit only Crick or Watson strands) yielding noisy single-cell data, as previously described^48^. Aligned BAM files were used for inversion discovery as described in^23^.

#### e. Hi-C data production

Lymphoblastoid cell lines were obtained from Coriell Cell Repositories and cultured in RPMI 1640 supplemented with 15% FBS. Cells were maintained at 37°C in an atmosphere containing 5% CO_2_. Hi-C libraries using 1.5 M human cells as input were generated with Proximo Hi-C kits v4.0 (Phase Genomics, Seattle, WA) following the manufacturer’s protocol with the following modification: in brief, cells were crosslinked, quenched, lysed sequentially with Lysis Buffers 1 and 2, and liberated chromatin immobilized on magnetic recovery beads. A 4-enzyme cocktail composed of DpnII (GATC), DdeI (CTNAG), HinfI (GANTC), and MseI (TTAA) was used during the fragmentation step to improve coverage and aid haplotype phasing. Following fragmentation and fill-in with biotinylated nucleotides, fragmented chromatin was proximity ligated for 4 hrs at 25°C. Crosslinks were then reversed, DNA purified and biotinylated junctions recovered using magnetic streptavidin beads. Bead-bound proximity ligated fragments were then used to generate a dual-unique indexed library compatible with Illumina sequencing chemistry. The Hi-C libraries were evaluated using fluorescent-based assays, including qPCR with the Universal KAPA Library Quantification Kit and Tapestation (Agilent). Sequencing of the libraries was performed at New York Genome Center (NYGC) on an Illumina Novaseq 6000 instrument using 2×150 bp cycles.

#### f. RNAseq data production

Total RNA of cell pellets were isolated using QIAGEN RNeasy Mini Kit according to the manufacturer’s instructions. Briefly, each cell pellet (10 M cells) was homogenized and lysed in Buffer RLT Plus, supplemented with 1% β-mercaptoethanol. The lysate-containing RNA was purified using an RNeasy spin column, followed by an in-column DNase I treatment by incubating for 10 min at RT, and then washed. Finally, total RNA was eluted in 50 uL RNase-free water. RNA-seq libraries were prepared with 300 ng total RNA using KAPA RNA Hyperprep with RiboErase (Roche) according to the manufacturer’s instructions. First, ribosomal RNA was depleted using RiboErase. Purified RNA was then fragmented at 85°C for 6 min, targeting fragments ranging 250-300 bp. Fragmented RNA was reverse transcribed with an incubation of 25°C for 10 min, 42°C for 15 min, and an inactivation step at 70°C for 15 min. This was followed by a second strand synthesis and A-tailing at 16°C for 30 min, 62°C for 10 min. The double-stranded cDNA A-tailed fragments were ligated with Illumina unique dual index adapters. Adapter-ligated cDNA fragments were then purified by washing with AMPure XP beads (Beckman). This was followed by 10 cycles of PCR amplification. The final library was cleaned up using AMPure XP beads. Quantification of libraries was performed using real-time qPCR (Thermo Fisher). Sequencing was performed on an Illumina NovaSeq platform generating paired end reads of 100 bp at The Jackson Laboratory for Genomic Medicine.

#### g. Iso-seq data production

Iso-seq data were generated at The Jackson Laboratory. Total RNA was extracted from 10 M human cell pellets. 300 ng total RNA were used to prepare Iso-seq libraries according to Iso-seq Express Template Preparation (Pacbio). First, full-length cDNA was generated using NEBNext Single Cell/ Low Input cDNA synthesis and Amplification Module in combination with Iso-seq Express Oligo Kit. Amplified cDNA was purified using ProNex beads. The cDNA yield of 160–320Lng then underwent SMRTbell library preparation including a DNA damage repair, end repair, and A-tailing and finally ligated with Overhang Barcoded Adapters. Libraries were sequenced on Pacbio Sequel II. Iso-seq reads were processed with default parameters using the PacBio Iso-seq3 pipeline.

### 3. Construction and dating of Y phylogeny

The genotypes were jointly called from the 1000 Genomes Project Illumina high-coverage data from^50^ using the ∼10.4 Mbp of chromosome Y sequence previously defined as accessible to short-read sequencing^51^. BCFtools (v1.9)^52,53^ was used with minimum base quality and mapping quality 20, defining ploidy as 1, followed by filtering out SNVs within 5 bp of an indel call (SnpGap) and removal of indels. Additionally, we filtered for a minimum read depth of 3. If multiple alleles were supported by reads, then the fraction of reads supporting the called allele should be ≥0.85; otherwise, the genotype was converted to missing data. Sites with ≥6% of missing calls, i.e., missing in more than 3 out of 44 samples, were removed using VCFtools (v0.1.16)^54^. After filtering, a total of 10,406,108 sites remained, including 12,880 variant sites. Since Illumina short-read data was not available from two samples, HG02486 and HG03471, data from their fathers (HG02484 and HG03469, respectively) was used for Y phylogeny construction and dating.

The Y haplogroups of each sample were predicted as previously described^14^ and correspond to the International Society of Genetic Genealogy nomenclature (ISOGG, https://isogg.org, v15.73, accessed in August 2021). We used the coalescence-based method implemented in BEAST (v1.10.4^55^ to estimate the ages of internal nodes in the Y phylogeny. A starting maximum likelihood phylogenetic tree for BEAST was constructed with RAxML (v8.2.10^56^) with the GTRGAMMA substitution model. Markov chain Monte Carlo samples were based on 200 million iterations, logging every 1000 iterations. The first 10% of iterations were discarded as burn-in. A constant-sized coalescent tree prior, the GTR substitution model, accounting for site heterogeneity (gamma) and a strict clock with a substitution rate of 0.76 × 10^−9^ (95% confidence interval: 0.67 × 10^−9^– 0.86 × 10^−9^) single-nucleotide mutations per bp per year was used^57^. A prior with a normal distribution based on the 95% confidence interval of the substitution rate was applied. A summary tree was produced using TreeAnnotator (v1.10.4) and visualized using the FigTree software (v1.4.4).

The closely related pair of African E1b1a1a1a-CTS8030 lineage Y chromosomes carried by NA19317 and NA19347 differ by 3 SNVs across the 10,406,108 bp region, with the TMRCA estimated to 200 ya (95% HPD interval: 0 - 500 ya).

A separate phylogeny (see **Fig. 4f**) was reconstructed using seven samples (HG01890, HG02666, HG01106, HG02011, T2T Y from NA24385/HG002, HG00358 and HG01952) with contiguously assembled Yq12 region following identical approach to that described above, with a single difference that sites with any missing genotypes were filtered out. The final callset used for phylogeny construction and split time estimates using Beast contained a total of 10,382,177 sites, including 5,918 variant sites.

### 4. *De novo* Assembly Generation

#### a. Reference assemblies

We used the GRCh38 (GCA_000001405.15) and the CHM13 (GCA_009914755.3) plus the T2T Y assembly from GenBank (CP086569.2) released in April 2022. We note that we did not use the unlocalised GRCh38 contig “chrY_KI270740v1_random” (37,240 bp, composed of 289 *DYZ19* primary repeat units) in any of the analyses presented in this study.

#### b. Constructing *de novo* assemblies

All 28 HGSVC and 15 HPRC samples were processed with the same Snakemake^58^ workflow (see “Code Availability” statement in main text) to first produce a *de novo* whole-genome assembly from which selected sequences were extracted in downstream steps of the workflow. The *de novo* whole-genome assembly was produced using Verkko v1.0^15^ with default parameters, combining all available PacBio HiFi and ONT data per sample to create a whole-genome assembly:

`verkko -d work_dir/ --hifi {hifi_reads} --nano {ont_reads}`

We note here that we had to manually modify the assembly FASTA file produced by Verkko for the sample NA19705 for the following reason: at the time of assembly production, the Verkko assembly for the sample NA19705 was affected by a minor bug in Verkko v1.0 resulting in an empty output sequence for contig “0000598”. The Verkko development team suggested removing the affected record, i.e., the FASTA header plus the subsequent blank line, because the underlying bug is unlikely to affect the overall quality of the assembly. We followed that advice, and continued the analysis with the modified assembly FASTA file. Our discussion with the Verkko development team is publicly documented in the Verkko Github issue #66. The assembly FASTA file was adapted as follows:

`egrep -v “(^$|unassigned\-0000598)” assembly.original.fasta > assembly.fasta`

For the samples with at least 50X HiFi input coverage (termed high-coverage samples, **Tables S1-S2**), we generated alternative assemblies using hifiasm v0.16.1-r375^59^ for quality control purposes. Hifiasm was executed with default parameters using only HiFi reads as input, thus producing partially phased output assemblies “hap1” and “hap2” (cf. hifiasm documentation): `hifiasm -o {out_prefix} -t {threads} {hifi_reads}`

The two hifiasm haplotype assemblies per sample are comparable to the Verkko assemblies in that they represent a diploid human genome without further identification of specific chromosomes, i.e., the assembled Y sequence contigs have to be identified in a subsequent process that we implemented as follows.

We employed a simple rule-based strategy to identify and extract assembled sequences for the two quasi-haploid chromosomes X and Y. The following rules were applied in the order stated here:

Rule 1: the assembled sequence has primary alignments only to the target sequence of interest, i.e., to either chrY or chrX. The sequence alignments were produced with minimap2 v2.24^60^:

`minimap2 -t {threads} -x asm20 -Y --secondary=yes -N 1 --cs -c -- paf-no-hit`

Rule 2: the assembled sequence has mixed primary alignments, i.e., not only to the target sequence of interest, but exhibits Y-specific sequence motif hits for any of the following motifs: *DYZ1*, *DYZ18* and the secondary repeat unit of *DYZ3* from^3^. The motif hits were identified with HMMER v3.3.2dev (commit hash #016cba0)^61^:

`nhmmer --cpu {threads} --dna -o {output_txt} --tblout {output_table} -E 1.60E-150 {query_motif} {assembly}`

Rule 3: the assembled sequence has mixed primary alignments, i.e., not only to the target sequence of interest, but exhibits more than 300 hits for the Y-unspecific repeat unit *DYZ2* (see Section ‘**Yq12 *DYZ2* Consensus and Divergence**’ for details on *DYZ2* repeat unit consensus generation). The threshold was determined by expert judgement after evaluating the number of motif hits on other reference chromosomes. The same HMMER call as for rule 2 was used with an E-value cutoff of 1.6e-15 and a score threshold of 1700.

Rule 4: the assembled sequence has no alignment to the chrY reference sequence, but exhibits Y-specific motif hits as for rule 2.

Rule 5: the assembled sequence has mixed primary alignments, but more than 90% of the assembled sequence (in bp) has a primary alignment to a single target sequence of interest; this rule was introduced to resolve ambiguous cases of primary alignments to both chrX and chrY.

After identification of all assembled chrY and chrX sequences, the respective records were extracted from the whole-genome assembly FASTA file and, if necessary, reverse-complemented to be in the same orientation as the T2T reference using custom code.

#### c. Assembly evaluation and validation

##### Error detection in *de novo* assemblies

Following established procedures^19,62^, we implemented three independent approaches to identify regions of putative misassemblies for all 43 samples. First, we used VerityMap (v2.1.1-alpha-dev #8d241f4)^18^ that generates and processes read-to-assembly alignments to flag regions in the assemblies that exhibit spurious signal, i.e., regions of putative assembly errors, but that may also indicate difficulties in the read alignment. Given the higher accuracy of HiFi reads, we executed VerityMap only with HiFi reads as input:

`python repos/VerityMap/veritymap/main.py --no-reuse –reads {hifi_reads} -t {threads} -d hifi -l SAMPLE-ID -o {out_dir} {assembly_FASTA}`

Second, we used DeepVariant (v1.3.0)^63^ and the PEPPER-Margin-DeepVariant pipeline (v0.8, DeepVariant v1.3.0^64^) to identify heterozygous (HET) SNVs using both HiFi and ONT reads aligned to the *de novo* assemblies. Given the quasi-haploid nature of the chromosome Y assemblies, we counted all HET SNVs remaining after quality filtering (bcftools v1.15 “filter” QUAL>=10) as putative assembly errors:

/opt/deepvariant/bin/run_deepvariant --model_type=“PACBIO” -- ref={assembly_FASTA} --num_shards={threads} --reads={HiFi-to-assembly_BAM} --sample_name=SAMPLE-ID --output_vcf={out_vcf} -- output_gvcf={out_gvcf} --intermediate_results_dir=$TMPDIR

run_pepper_margin_deepvariant call_variant --bam {ONT-to- assembly_BAM} --fasta {assembly_FASTA} --output_dir {out_dir} -- threads {threads} --ont_r9_guppy5_sup --sample_name SAMPLE-ID -- output_prefix {out_prefix} --skip_final_phased_bam --gvcf

Third, we used the tool NucFreq^20^ to identify positions in the HiFi read-to-assembly alignments where the second most common base is supported by at least 10% of the alignments. BAM files were filtered with samtools using the flag 2308 (drop secondary and supplementary alignments) following the information in the NucFreq readme. Additionally, we only processed assembled contigs larger than 500 kbp to limit the effect of spurious alignments in short contigs. NucFreq was then executed with default parameters:

`NucPlot.py --obed OUTPUT.bed --threads {threads} --bed ASSM-CONTIGS.bed HIFI.INPUT.bam OUTPUT.png`

We note here that NucFreq could not successfully process the alignments for sample HG00512 due to an error in the graphics output. We thus omitted this sample from the following processing steps. Again following the information in the NucFreq readme, we then created flagged regions if more than five positions were flagged in a 500 bp window, and subsequently merged overlapping windows (**Table S12**).

As a final processing step, we merged the VerityMap- and NucFreq-flagged regions (subsuming HET SNVs called by either DeepVariant or PEPPER) by tripling each region’s size (flanking region upstream and downstream) and then merging all overlapping regions with bedtools:

`bedtools merge -c 4 -o collapse -i CONCAT-ALL-REGIONS.bed > OUT.MERGED.bed`

The resulting clusters and all regions separately were post-processed with custom code to derive error estimates for the assemblies (see “Code Availability” statement in main text and “Data availability” for access to BED files listing all flagged regions/positions and merged clusters; **Table S10**).

Since the PAR1 subregion was contiguously assembled from only 10 samples (**Table S14, S16)**, all highlighted regions as putative assembly errors by VerityMap were visually evaluated in the HiFi and ONT read alignments to the assembly using the Integrative Genomics Viewer (IGV v2.14.1)^65^ (**Table S11**).

Assembly QV estimates were produced with yak v0.1 (github.com/lh3/yak) following the examples in its documentation (see readme in referenced repository). The QV estimation process requires an independent sequence data source to derive a (sample-specific) reference k-mer set to compare the k-mer content of the assembly. In our case, we used available short read data to create said reference k-mer set, which necessitated excluding the samples HG02486 and HG03471 because no short reads were available. For the chromosome Y-only QV estimation, we restricted the short reads to those with primary alignments to our Y assemblies or to the T2T Y, which we added during the alignment step to capture reads that would align to Y sequences missing from our assemblies.

##### Assembly gap detection

We used the recently introduced tool Rukki (packaged with Verkko v1.2)^15^ to derive estimates of potential gaps in our assemblies. After having identified chrY and chrX contigs as described above, we used this information to prepare annotation tables for Rukki to identify chrX/chrY paths in the assembly graph:

`rukki trio -g {assembly_graph} -m {XY_contig_table} -p {out_paths} - -final-assign {out_node_assignment} --try-fill-bubbles --min-gap- size 1 --default-gap-size 1000`

The resulting set of paths including gap estimates was summarized using custom code (**Table S6**). See “Data availability” section for access to NucFreq plots generated for all samples.

##### Assembly evaluation using Bionano Genomics optical mapping data

To evaluate the accuracy of Verkko assemblies, all samples (n=43) were first *de novo* assembled using the raw optical mapping molecule files (bnx), followed by alignment of assembled contigs to the T2T whole genome reference genome assembly (CHM13 + T2T Y) using Bionano Solve (v3.5.1) pipelineCL.py.

`python2.7 Solve3.5.1_01142020/Pipeline/1.0/pipelineCL.py -T 64 -U -j`

`64 -jp 64 -N 6 -f 0.25 -i 5 -w -c 3 \`

`-y \`

`-b ${ bnx} \`

`-l ${output_dir} \`

`-t Solve3.5.1_01142020/RefAligner/1.0/ \`

`-a`

`Solve3.5.1_01142020/RefAligner/1.0/optArguments_haplotype_DLE1_saphy r_human.xml \`

`-r ${ref}`

To improve the accuracy of optical mapping Y chromosomal assemblies, unaligned molecules, molecules that align to T2T chromosome Y and molecules that were used for assembling contigs but did not align to any chromosomes were extracted from the optical mapping *de novo* assembly results. These molecules were used for the following three approaches: 1) local *de novo* assembly using Verkko assemblies as the reference using pipelineCL.py, as described above; 2) alignment of the molecules to Verkko assemblies using refAligner (Bionano Solve (v3.5.1)); and 3) hybrid scaffolding using optical mapping *de novo* assembly consensus maps (cmaps) and Verkko assemblies by hybridScaffold.pl.

`perl Solve3.5.1_01142020/HybridScaffold/12162019/hybridScaffold.pl \`

`-n ${fastafile} \`

`-b ${bionano_cmap} \`

`-c`

`Solve3.5.1_01142020/HybridScaffold/12162019/hybridScaffold_DLE1_conf ig.xml \`

`-r Solve3.5.1_01142020/RefAligner/1.0/RefAligner \`

`-o ${output_dir} \`

`-f -B 2 -N 2 -x -y \`

`-m ${bionano_bnx} \`

`-p Solve3.5.1_01142020/Pipeline/12162019/ \`

`-q`

`Solve3.5.1_01142020/RefAligner/1.0/optArguments_nonhaplotype_DLE1_sa phyr_human.xml`

Inconsistencies between optical mapping data and Verkko assemblies were identified based on variant calls from approach 1 using “exp_refineFinal1_merged_filter_inversions.smap” output file. Variants were filtered out based on the following criteria: a) variant size smaller than 500 base pairs; b) variants labeled as “heterozygous”; c) translocations with a confidence score of ≤0.05 and inversions with a confidence score of ≤0.7 (as recommended on Bionano Solve Theory of Operation: Structural Variant Calling - Document Number: 30110); d) variants with a confidence score of <0.5. Variant reference start and end positions were then used to evaluate the presence of single molecules which span the entire variant using alignment results from approach 2. Alignments with a confidence score of <30.0 were filtered out. Hybrid scaffolding results, conflict sites provided in “conflicts_cut_status.txt” output file from approach 3 were used to evaluate if inconsistencies identified above based on optical mapping variant calls overlap with conflict sites (i.e., sites identified by hybrid scaffolding pipeline representing inconsistencies between sequencing and optical mapping data) (**Table S48**). Furthermore, we used molecule alignment results to identify coordinate ranges on each Verkko assembly which had no single DNA molecule coverage using the same alignment confidence score threshold of 30.0, as described above, dividing assemblies into 10 kbp bins and counting the number single molecules covering each 10 kbp window (**Table S49**).

#### d. *De novo* assembly annotation

##### Annotation of Y-chromosomal subregion

The 24 Y-chromosomal subregion coordinates (**Table S13**) relative to the GRCh38 reference sequence were obtained from^7^. Since Skov et al. produced their annotation on the basis of a coordinate liftover from GRCh37, we updated some coordinates to be compatible with the following publicly available resources: for the pseudoautosomal regions we used the coordinates from the UCSC Genome Browser for GRCh38.p13 as they slightly differed. Additionally, Y-chromosomal amplicon start and end coordinates were edited according to more recent annotations from^66^, and the locations of *DYZ19* and *DYZ18* repeat arrays were adjusted based on the identification of their locations using HMMER3 (v3.3.2)^67^ with the respective repeat unit consensus sequences from^3^.

The locations and orientations of Y-chromosomal subregions in the T2T Y were determined by mapping the subregion sequences from the GRCh38 Y to the T2T Y using minimap2 (v2.24, see above). The same approach was used to determine the subregion locations in each *de novo* assembly with subregion sequences from both GRCh38 and the T2T Y (**Table S13**). The locations of the *DYZ18* and *DYZ19* repeat arrays in each *de novo* assembly were further confirmed (and coordinates adjusted if necessary) by running HMMER3 (see above) with the respective repeat unit consensus sequences from^3^. Only tandemly organized matches with HMMER3 score thresholds higher than 1700 for *DYZ18* and 70 for *DYZ19*, respectively, were included and used to report the locations and sizes of these repeat arrays.

A Y-chromosomal subregion was considered as contiguous if it was assembled contiguously from the subclass on the left to the subclass on the right (note that the *DYZ18* subregion is completely deleted in HG02572), except for pseudoautosomal regions where they were defined as >95% length of the T2T Y pseudoautosomal regions and with no unplaced contigs. Note that due to the requirement of no unplaced contigs the assembly for HG02666 appears to have a break in PAR2 subregion, while it is contiguously assembled from the telomeric sequence of PAR1 to telomeric sequence in PAR2 without breaks (however, there is a ∼14 kbp unplaced PAR2 contig aligning best to a central region of PAR2). The assembly of HG01890 however has a break approximately 100 kbp before the end of PAR2. Assembly of PAR1 remains especially challenging due to its sequence composition and sequencing biases^8,9^, and among our samples was contiguously assembled for 10/43 samples, while PAR2 was contiguously assembled for 39/43 samples.

##### Annotation of centromeric and pericentromeric regions

To annotate the centromeric regions, we first ran RepeatMasker (v4.1.0) on 26 Y-chromosomal assemblies (22 samples with contiguously assembled pericentromeric regions, 3 samples with a single gap and no unplaced centromeric contigs, and the T2T Y) to identify the locations of α-satellite repeats using the following command:

`RepeatMasker -species human -dir {path_to_directory} –pa {num_of_threads} {path_to_fasta}`

Then, we subsetted each contig to the region containing α-satellite repeats and ran HumAS-HMMER (v3.3.2; https://github.com/fedorrik/HumAS-HMMER_for_AnVIL) to identify the location of α-satellite higher-order repeats (HORs), using the following command:

`Hmmer-run.sh {directory_with_fasta} AS-HORs-hmmer3.0- 170921.hmm {num_of_threads}`

We combined the outputs from RepeatMasker (v4.1.0) and HumAS-HMMER to generate a track that annotates the location of α-satellite HORs and monomeric or diverged α-satellite within each centromeric region.

To determine the size of the α-satellite HOR array (reported for 26 samples in **Table S16,** while size estimates reported in results section include 23 gapless assemblies - see **Table S15**), we used the α-satellite HOR annotations generated via HumAS-HMMER (v3.3.2; described above) to determine the location of *DYZ3* α-satellite HORs, focusing on only those HORs annotated as “live” (e.g., S4CYH1L). Live HORs are those that have a clear higher-order pattern and are highly (>90%) homogenous^68^. This analysis was conducted on 21 centromeres (including the T2T Y, **Fig. S57**), excluding 5/26 samples (NA19384, HG01457, HG01890, NA19317, NA19331), where, despite a contiguously assembled pericentromeric subregion, the assembly contained unplaced centromeric contig(s).

To annotate the human satellite III (*HSat3*) and *DYZ17* arrays within the pericentromere, we ran StringDecomposer (v1.0.0) on each assembly centromeric contig using the *HSat3* and *DYZ17* consensus sequences described in Altemose, 2022, Seminars in Cell and Developmental Biology^69^ and available at the following URL: https://github.com/altemose/HSatReview/blob/main/Output_Files/HSat123_consensus_sequences.fa We ran the following command:

`stringdecomposer/run_decomposer.py {path_to_contig_fasta} {path_to_consensus_sequence+fasta} -t {num_of_threads} –o {output_tsv}`

The *HSat3* array was determined as the region that had a sequence identity of 60% or greater, while the *DYZ17* array was determined as the region that had a sequence identity of 65% or greater.

### 5. Downstream analysis

#### a. Effect of input read depth on assembly contiguity

We explored a putative dependence between the characteristics of the input read sets, such as read length N50 or genomic coverage, and the resulting assembly contiguity by training multivariate regression models (“ElasticNet” from scikit-learn v1.1.1, see “Code Availability” statement in main text). The models were trained following standard procedures with 5-fold nested cross-validation (see scikit-learn documentation for “ElasticNetCV”). We note that we did not use the haplogroup information due to the unbalanced distribution of haplogroups in our dataset. We selected basic characteristics of both the HiFi and ONT-UL input read sets (read length N50, mean read length, genomic coverage and genomic coverage for ONT reads exceeding 100 kbp in length, i.e., the so- called ultralong fraction of ONT reads; **Table S51**) as model features and assembly contig NG50, assembly length or number of assembled contigs as target variable.

#### b. Locations of assembly gaps

The assembled Y-chromosomal contigs were mapped to the GRCh38 and the CHM13 plus T2T Y reference assemblies using minimap2 with the flags `-x asm20 -Y -p 0.95 -- secondary=yes -N 1 -a -L --MD --eqx`. The aligned Y-chromosomal sequences for each reference were partitioned to 1 kbp bins to investigate assembly gaps. Gap presence was inferred in bins where the average read depth was either lower or higher than 1. To investigate the potential factors associated with gap presence, we analysed these sequences to compare the GC content, SD content, and Y subregion. Read depth for each bin was calculated using mosdepth^70^ and the flags *`–n -x`*. GC content for each bin was calculated using BedTools `nuc` function^71^. SD locations for GRCh38 Y were obtained from the UCSC genome browser, and for the CHM13 plus T2T Y from^2^. Y-chromosomal subregion locations were determined as described in Methods section ‘*De novo* assembly annotation with Y-chromosomal subregions’. The bin read depth and GC content statistics were merged into matrices and visualized using *matplotlib* and *seaborn*^72,73^.

#### c. Comparison of assembled Y subregion sizes across samples

Sizes for each chromosome’s (peri-)centromeric regions were obtained as described in Methods section ‘Annotation of pericentromeric regions’. The size variation of (peri-)centromeric regions (*DYZ3* alpha-satellite array, *Hsat3*, *DYZ17* array, and total (peri-)centromeric region), and the *DYZ19*, *DYZ18* and *TSPY* repeat arrays were compared across samples using a heatmap, incorporating phylogenetic context. The sizes of the (peri-)centromeric regions (*DYZ3* alpha-satellite array, *Hsat3* and *DYZ17* array) were regressed against each other using the OLS function in *statsmodels*, and visualized using *matplotlib* and *seaborn*^72^.

#### d. Comparison and visualization of *de novo* assemblies

The similarities of three contiguously assembled Y chromosomes (HG00358, HG02666, HG01890), including comparison to both GRCh38 and the T2T Y, was assessed using blastn^74^ with sequence identity threshold of 80% (95% threshold was used for PAR1 subregion) (**Fig. 2b**) and excluding non-specific alignments (i.e., showing alignments between different Y subregions), followed by visualization with genoPlotR (0.8.11)^75^. Y subregions were uploaded as DNA segment files and alignment results were uploaded as comparison files following the file format recommended by the developers of the genoplotR package. Unplaced contigs were excluded, and all Y-chromosomal subregions, except for Yq12 heterochromatic region and PAR2, were included in queries.

blastn -query $file1 -subject $file2 -subject_besthit -outfmt ’7 qstart qend sstart send qseqid sseqid pident length mismatch gaps evalue bitscore sstrand qcovs qcovhsp qlen slen’ –out ${outputfile}.out

`plot_gene_map(dna_segs=dnaSegs, comparisons=comparisonFiles, xlims=xlims, legend = TRUE, gene_type = “headless_arrows”, dna_seg_scale=TRUE, scale=FALSE)`

For other samples, three-way comparisons were generated between the GRCh38 Y, Verkko *de novo* assembly and the T2T Y sequences, removing alignments with less than 80% sequence identity. The similarity of closely related NA19317 and NA19347 Y-chromosomal assemblies was assessed using the same approach.

#### e. Sequence identity heatmaps

Sequence identity within repeat arrays was investigated by running StainedGlass^76^. For the centromeric regions, StainedGlass was run with the following configuration: `window = 5785 and mm_f = 30000`. We adjusted the colour scale in the resulting plot using a custom R script that redefines the breaks in the histogram and its corresponding colours. This script is publicly available here: https://eichlerlab.gs.washington.edu/help/glogsdon/Shared_with_Pille/StainedGlass_adjustedScale.R. The command used to generate the new plots is: StainedGlass_adjustedScale.R –b {output_bed} -p {plot_prefix}. For the *DYZ19* repeat array, window = 1000 and mm_f = 10000 were used, 5 kbp of flanking sequence was included from both sides, followed by adjustment of colour scale using the custom R script (see above).

For the Yq12 subregion (including the *DYZ18* repeat array), `window = 5000 and mm_f = 10000` were used, and 10 kbp of flanking sequence was included. In addition to samples with contiguously assembled Yq12 subregion, the plots were generated for two samples (NA19705 and HG01928) with a single gap in Yq12 subregion (the two contigs containing Yqhet sequence were joined into a single contig with 100 Ns added to the joint location). HG01928 contains a single unplaced Yqhet contig (approximately 34 kbp in size) which was not included. For the Yq11/Yq12 transition region, 100 kbp proximal to the *DYZ18* repeat and 100 kbp of the first *DYZ1* repeat array was included in the StainedGlass runs, using `window = 2000 and mm_f = 10000`.

#### f. Dotplot generation

Dotplot visualizations were created using the NAHRwhals package version 0.9 which provides visualization utilities and a custom pipeline for pairwise sequence alignment based on minimap2 (v2.24). Briefly, NAHRwhals initiates pairwise alignments by splitting long sequences into chunks of 1 - 10 kbp which are then aligned to the target sequence separately, enhancing the capacity of minimap2 to correctly capture inverted or repetitive sequence alignments. Subsequently, alignment pairs are concatenated whenever the endpoint of one alignment falls in close proximity to the startpoint of another (base pair distance cutoff: 5% of the chunk length). Pairwise alignment dotplots are created with a pipeline based on the ggplot2 package, with optional .bed files accepted for specifying colorization or gene annotation. The NAHRwhals package and further documentation are available at https://github.com/WHops/nahrchainer, please see ‘Data access’ section on how to access the dotplots.

#### g. Inversion analyses

##### Inversion calling using Strand-seq data

The inversion calling from Strand-seq data, available for 30/43 samples and the T2T Y, using both the GRCh38 and the T2T Y sequences as references was performed as described previously^23^.

Note on the P5 palindrome spacer direction in the T2T Y assembly: the P5 spacer region is present in the same orientation in both GRCh38 (where the spacer orientation had been chosen randomly, see Supplementary Figure 11 from^3^ for more details) and the T2T Y sequence, while high-confidence calls from the Strand-seq data from individual HG002/NA24385 against both the GRCh38 and T2T Y report it to be in inverted orientation. It is therefore likely that the P5 spacer orientations are incorrect in both GRCh38 Y and the T2T Y and in the P5 inversion recurrence estimates we therefore considered HG002/NA24385 to carry the P5 inversion (as shown on **Fig. 3a**, inverted relative to GRCh38).

##### Inversion calling from the *de novo* assemblies

In order to determine the inversions from the *de novo* assemblies, we aligned the Y-chromosomal repeat units/SDs as published by^66^ to the *de novo* assemblies as described above (see Section: ‘Annotation with Y-chromosomal subregions’). Inverted alignment orientation of the unique sequences flanked by repeat units/SDs relative to the GRCh38 Y was considered as evidence of inversion. The presence of inversions was further confirmed by visual inspection of *de novo* assembly dotplots generated against both GRCh38 and T2T Y sequences (see Methods section: Dotplot generation), followed by merging with the Strand-seq calls (**Table S32**).

##### Inversion rate estimation

In order to estimate the inversion rate, we counted the minimum number of inversion events that would explain the observed genotype patterns in the Y phylogeny (**Fig. 3a**). A total of 12,880 SNVs called in the set of 44 males and Y chromosomal substitution rate from above (see Methods section ‘Construction and dating of Y phylogeny’) was used. A total of 126.4 years per SNV mutation was then calculated (0.76 x 10^-9^ x 10,406,108 bp)^-1^, and converted into generations assuming a 30-year generation time^77^. Thus each SNV corresponds to 4.21 generations, translating into a total branch length of 54,287 generations for the 44 samples. For a single inversion event in the phylogeny this yields a rate of 1.84 x 10^-5^ (95% CI: 1.62 x 10^-5^ to 2.08 x 10^-5^) mutations per father-to-son Y transmission. The confidence interval of the inversion rate was obtained using the confidence interval of the SNV rate.

##### Determination of inversion breakpoint ranges

We focused on the following eight recurrent inversions to narrow down the inversion breakpoint locations: IR3/IR3, IR5/IR5, and palindromes P8, P7, P6, P5, P4 and P3 (**Fig. 3a**), and leveraged the ‘phase’ information (i.e., proximal/distal) of paralogous sequence variants (PSVs) across the SDs mediating the inversions as follows. First, we extracted proximal and distal inverted repeat sequences flanking the identified inversions (spacer region) and aligned them using MAFFT (v7.487)^78,79^ with default parameters. From the alignment, we only selected informative sites (i.e., not identical across all repeats and samples), excluding singletons and removing sites within repetitive or poorly aligned regions as determined by Tandem Repeat Finder (v4.09.1)^80^ and Gblocks (v0.91b)^81^, respectively. We inferred the ancestral state of the inverted regions following the maximum parsimony principle as follows: we counted the number of inversion events that would explain the distribution of inversions in the Y phylogeny by assuming a) that the reference (i.e., same as GRCh38 Y) state was ancestral and b) that the inverted (i.e., inverted compared to GRCh38 Y) state was ancestral. The definition of ancestral state for each of the regions was defined as the lesser number of events to explain the tree (IR3: reference; IR5: reference; P8: inverted; P7: reference; P5: reference; P4: reference; P3: reference). As we observed a clear bias of inversion state in both African (Y lineages A, B and E) and non-African Y lineages for the P6 palindrome (the African Y lineages have more inverted states (17/21) and non-African Y lineages have more reference states (17/23)), we determined the ancestral state and inversion breakpoints for African and non-African Y lineages separately in the following analyses.

We then defined an ancestral group as any samples showing an ancestral direction in the spacer region, and selected sites that have no overlapping alleles between the proximal and distal alleles in the defined ancestral group, which were defined as the final set of informative PSVs. For IR3, we used the ancestral group as samples with Y-chromosomal structure 1 (i.e., with the single ∼20.3 kbp TSPY repeat located in the proximal IR3 repeat) and ancestral direction in the spacer region. According to the allele information from the PSVs, we determined the phase (proximal or distal) for each PSV across samples. Excluding non-phased PSVs (e.g., the same alleles were found in both proximal and distal sequences), any two adjacent PSVs with the same phase were connected as a segment while masking any single PSVs with a different phase from the flanking ones to only retain reliable contiguous segments. An inversion breakpoint was determined to be a range where phase switching occurred between two segments, and the coordinate was converted to the T2T Y coordinate based on the multiple sequence alignment and to the GRCh38 Y coordinate using the LiftOver tool at the UCSC Genome Browser web page (https://genome.ucsc.edu/cgi-bin/hgLiftOver). Samples with non-contiguous assembly of the repeat regions were excluded from each analysis of the corresponding repeat region.

#### h. Molecular evolution of Y-chromosomal palindromes

##### Alignments of palindromes and the XDRs

To construct alignments of P1, P2, P3 palindromes, P1 yellow and P3 teal SDs (**Fig. S39**), we first mapped XDR8, blue, teal, green, red, grey, yellow, and other2 sequences derived from the GRCh38 Y reference (using coordinates derived from^66^) to each assembly using minimap2 (v2.24, see above). We then reconstructed the order of non-overlapping SDs by selecting the longest from any overlapping mappings for each SD. The P1, P2, P3, P1 yellow, and P3 teal sequences were collected from each sample only if the directionality and origin of two palindromic arms could clearly be determined in the context of inter-palindromic inversion status. For example, sample HG02486 has the GRCh38 Y segments mappings in the order of: XDR8, blue, teal, teal, blue, green, red, red, green, yellow, blue, grey, red, red, green, yellow, blue, grey, and other2 on the same contig, which matches with the expected SD order given the inversion status of HG02486 (one *gr/rg* inversion). As we do not know the exact breakpoint locations for the *gr/rg* inversion in HG02486, the green and red SDs could be mixtures of P1 and P2 origin (since both contain green and red SDs and the *gr/rg* inversions breakpoints are likely to be located within these regions) due to the inversion. Therefore, in order to be conservative and to avoid SDs which are potential mixes from different origins, we here excluded the green and red segments from the analysis. Similar filtering was applied to all samples in order to reduce the possibility of including variation that has been caused by e.g., inter-palindromic inversions. As a result, we collected 18, 24, 28, 34, and 39 sequences for P1, P2, P3, P1 yellow, and P3 teal, respectively (**Table S15**), and aligned them using MAFFT (v7,487)^78,79^ with default parameters. For the other palindromes (P4, P5, P6, P7 and P8), we used the same alignments from the inversion analysis described above (see Methods section ‘Determination of inversion breakpoint ranges’ for details). The eight XDR sequences were collected by mapping the XDR sequences derived from GRCh38 Y reference to each assembly using minimap2, and aligning them using MAFFT (v7.487)^78,79^ with default parameters. The raw alignments were summarized by averaging frequencies of major allele (including a gap) across 100 bp windows (**Figs. S41** and **S42**).

For all the aligned sequences, we trimmed both ends of each alignment so that the GRCh38 Y coordinate system could be used. To minimize alignment errors and recurrent mutations, we masked sites residing in repetitive or poorly aligned regions as defined by Tandem repeat finder (v4.09.1)^80^ and Gblocks (v0.91b)^81^. Additionally, sites with a gap in any of the samples or with more than two alleles were masked from all samples. Lastly, we manually curated alignments by masking regions with any potential structural rearrangements in order to only consider point mutations in the following analyses. The final curated alignments contain unmasked regions ranging from 57.63 to 97.40% of raw alignment depending on the region.

##### Estimation of point mutation rates in palindromes

To estimate the point mutation rates of the Y-chromosomal palindromes, we first selected a set of 13 samples for which, following the stringent filtering and manual curation described above, alignments of all palindromic regions and XDRs were available. We then counted the number of SNVs in the XDR regions, which was used to calibrate the number of generations spanned by the 13 samples following the approach described in Skov et. al.^7^ (and described in more detail below), using the mutation rate estimates of the XDRs (3.14 x 10^-8^ per position per generation, PPPG) from Helgason et. al.^82^. Mutation events in the palindromes were determined considering the phylogenetic relationships of the 13 samples following the approach in Skov et. al.^7^. Lastly, the mutation rate (PPPG) for palindromes was estimated by dividing the number of mutation events by the estimated generations and 2 times the unmasked palindrome length for each palindrome (**Table S37**). For P1 and P3 palindromes, we analysed regions with and without multi-copy (i.e., >2 copies) SDs separately.

##### Detection of gene conversion events in palindromes

The gene conversion analysis was carried out for palindromes P1 yellow, P3 teal, P4, P5, P6, P7 and P8 with 34, 39, 36, 33, 43, 44 and 44 samples for the respective palindromes (including the T2T Y; **Tables S15, S36**). For each position in the alignment, we determined the genotypes for all internal nodes based on their child nodes and assigned gene conversion or mutation events for each node following the approach described in Skov et. al.^7^. Starting with all observed genotypes in the tree, we filled out genotypes of all ancestral nodes based on their child nodes. We then determined gene conversion or mutation events if the genotype of the parent was different from that of the child(ren). In case multiple scenarios could explain the phylogenetic tree and the observed genotypes at a particular position, the one with the lowest number of mutations was selected. Positions, for which the best scenario included more than one mutation, were excluded from this analysis. The bias towards the ancestral state or GC bases was statistically tested using the Chi-square test.

We would like to note that gene conversion events towards the ancestral state might be underestimated compared to the actual number of events as it is not possible to detect a gene conversion event towards the ancestral state in case it had occurred on the same branch where the mutation generating the paralogous sequence variant took place. In order to adjust for this bias in the detection of ‘to-ancestral’ and ‘to-derived’ gene conversion events, we changed the derived homozygous genotypes to ancestral homozygous genotypes in all gene conversion events detected in the initial gene conversion analysis. Using the modified genotypes, we then determined the gene conversion events using the same approach of the initial analysis, and recalculated the ancestral bias by discarding the gene conversion events that were not identified in the new analysis.

#### i. Variant calling

##### Variant calling using *de novo* assemblies

Variants were called from assembly contigs using PAV (v2.1.0)^19^ with default parameters using minimap2 (v2.17) contig alignments to GRCh38 (primary assembly only, ftp://ftp.1000genomes.ebi.ac.uk/vol1/ftp/data_collections/HGSVC2/technical/reference/20200513_hg38_NoALT/). Supporting variant calls were done against the same reference with PAV (v2.1.0) using LRA^83^ alignments (commit e20e67) with assemblies, PBSV (v2.8.0) (https://github.com/PacificBiosciences/pbsv) with PacBio HiFi reads, SVIM-asm (v1.0.2)^84^ with assemblies, Sniffles (v2.0.7)^85^ with PacBio HiFi and ONT, DeepVariant (v1.1.0)^16,63^ with PacBio HiFi, Clair3 (v0.1.12)^86^ with ONT, CuteSV (v2.0.1)^87^ with ONT, and LongShot (v0.4.5)^88^ with ONT. A validation approach based on the subseq command was used to search for raw-read support in PacBio HiFi and ONT^19^.

A merged callset was created from the PAV calls with minimap2 alignments across all samples with SV-Pop^19,89^ to create a single nonredundant callset. We used merging parameters “`nr::exact:ro(0.5):szro(0.5,200)`” for SV and indel insertions and deletions (Exact size & position, then 50% reciprocal overlap, then 50% overlap by size and within 200 bp), “`nr::exact:ro(0.2)`” for inversions (Exact size & position, then 20% reciprocal overlap), and “`nrsnv::exact`” for SNVs (exact position and REF/ALT match). The PAV minimap2 callset was intersected with each orthogonal support source using the same merging parameters. SVs were accepted into the final callset if they had support from two orthogonal sources with at least one being another caller (i.e., support from only subseq PacBio HiFi and subseq ONT was not allowed). Indels and SNVs were accepted with support from one orthogonal caller. Inversions were manually curated using dotplots and density plots generated by PAV. A chrX callset was constructed from chrX assemblies, and QC was applied using the same callers and parameters for the chrY callset, which was subsequently used for variant density estimations in PAR1.

An increase in SVs near contig ends can indicate errors, which we did not see evidence for with a minimum distance to a contig end of 6.9 kbp for SV insertions and 198 kbp for SV deletions. All SVs were anchored more than 1 kbp into contig ends except a single deletion in HG00673 (624 bp), and an average of 1.53 insertions and 0.77 deletions were anchored less than 10 kbp from contig ends.

To search for likely duplications within insertion calls, insertion sequences were re-mapped to the reference with minimap2 (v2.17) with parameters `“-x asm20 -H --secondary=no -r 2k -Y -a --eqx -L -t 4`”.

To characterize variant densities in PAR1, MSY, and chrX against autosomes, we split variants into four callsets: 1) MSY, 2) autosomes, 3) chrX outside PARs, and 4) PAR1 from both chrX and chrY (post-QC PAV calls). The MSY and PAR1 callsets were derived from this study, and the autosomes and chrX was derived from Ebert 2021^19^. For chrX, we included only female samples to avoid technical biases in the analysis. We excluded tandem repeats and SDs to avoid overwhelming our signal by higher mutability rates and potential technical biases within these regions. For variant call rates, we computed the variants per Mbp eliminating any uncallable and repetitive loci from the denominator, which includes assembly gaps, centromeres and difficult to align regions around them (low-confidence filter published with Ebert 2021), tandem repeats, SDs, and any loci that were not contiguously mappable for variant calling as flagged by PAV. For all statistics, the choice between Student’s and Welch’s t-test was made by an F-test with a p-value cutoff at 0.01.

##### Validation of large SVs using optical mapping data

Orthogonal support for merged PAV calls were evaluated by using optical mapping data (**Table S52**). Molecule support was evaluated using local *de novo* assembly maps which aligned to GRCh38 reference assembly. This evaluation included all 29 SVs 5 kbp or larger in size (including 15 insertions and 14 deletions; **Table S21**) Although variants <5 kbp could be resolved by optical mapping technique, there were loci without any fluorescent labels which could lead to misinterpretation of the results. Variant reference (GRCh38) start and end positions were used to evaluate the presence of single molecules which span the variant breakpoints using alignment results using Bionano Access (v1.7). Alignments with a confidence score of <30.0 were filtered out.

##### TSPY repeat array copy number analysis

To perform a detailed analysis of the TSPY repeat array, known to be highly variable in copy number^90^, the consensus sequence of the repeat unit was first constructed as follows. The repeat units were determined from the T2T Y sequence, the individual repeat unit sequences extracted and aligned using MAFFT (v7.487)^78,79^ with default parameters. A consensus sequence was generated using EMBOSS cons (v6.6.0.0) command line version with default parameters, followed by manual editing to replace sites defined as ‘N’s with the major allele across the repeat units. The constructed TSPY repeat unit consensus sequence was 20,284 bp.

The consensus sequence was used to identify TSPY repeat units from each *de novo* assembly using HMMER3 v3.3.2^67^, excluding five samples (HG03065, NA19239, HG01258, HG00096, HG03456) with non-contiguous assembly of this region. TSPY repeat units from each assembly were aligned using MAFFT as described above, followed by running HMMER functions “esl-alistat” and “esl-alipid” to obtain sequence identity statistics (**Table S18**).

Dotplots of the TSPY repeat array were generated using EMBOSS dotpath (v6.6.0.0) command line version with default parameters, with varying window sizes (2, 5 and 10 kbp). The TSPY repeat array locations as reported in **Table S17** were used, with 5 kbp of flanking sequence added to both sides.

##### Phylogenetic analysis of *TSPY*, *RBMY1,* and *DAZ* ampliconic Genes

We used Ensembl^91^ to retrieve the exon sequences for the *TSPY* (*TSPY1*), *RBMY1* (*RBMY1B*), and *DAZ* (*DAZ1*) ampliconic genes. The sequences of these exons were incorporated with the curated Dfam library^92^ into a custom RepeatMasker^93^ library. A local RepeatMasker installation was then utilized to screen all (43 assemblies, T2T Y and GRCh38) Y chromosome assemblies for exon sequence hits. For each assembly, we identified all low-divergence exon hits below 2% divergence. Individual *TSPY* and *RBMY1* gene copies were counted as protein-coding if all gene exons (six exons for *TSPY* and twelve exons for *RBMY1*) were present and classified as low divergence. Whereas for *DAZ*, there was a high variation in exon copy number across individual *DAZ* gene copies. Therefore, an individual *DAZ* gene copy was defined as simply all low divergence exons that are present in the same orientation within a *DAZ* gene cluster. After retrieving the individual exon sequences using SAMtools^52^, the exon sequences of each individual gene copy were ‘fused’ together using Python. Assemblies containing breaks in contigs within the TSPY array (excluding GRCh38) or *DAZ* gene clusters were dropped from all subsequent analyses. For each gene family, we generated a multiple sequence alignment (MSA, one alignment per ampliconic gene family) using MUSCLE and manually curated the alignment if needed. Multiple sequence alignment of *DAZ* sequences was deemed unfruitful due to the large variability in gene exon copy numbers. Subsequently, the *TSPY* and *RBMY* MSAs were given to IQ-Tree^94^ for phylogenetic analysis (see Phylogenetic Tree Analysis of *DYZ2* Consensus Sequence for details).

Additionally, we performed a network analysis for *TSPY* and *RBMY1* to identify clusters of sequences. First, we constructed a gene copy sequence distance matrix by computing the hamming distance between each sequence. Afterwards, we built an undirected weighted network, using NetworkX^95,96^, where each gene copy sequence is represented as a node and a weighted edge/link is placed between nodes that are the most similar to each other (i.e., lowest hamming distance). Following network construction, we utilized an asynchronous label propagation algorithm (as implemented in NetworkX)^97^ to identify gene sequence communities (i.e., clusters/subgraphs/subnetworks) within the network. Within each community, the node with the most connections (i.e., the ‘hub’ node) was identified and their sequence was considered the best representation of the community. If a network community was comprised of less than five nodes (*TSPY*), or three nodes (*RBMY1*), the sequence of each node within said community was compared to that of all ‘hub’ nodes and then reassigned to the community of the ‘hub’ node they most closely resembled (i.e., least hamming distance). Separate community size cut offs were utilized due to the large difference in total sequences (nodes) within each ampliconic gene network (*TSPY*: 1344 sequences, *RBMY1*: 353 sequences). Next, we created a community consensus sequence of each community. This was constructed from the MSA of the sequences of all gene copies within a community using MUSCLE and applying a majority rule approach to build the consensus sequence. The sequence of every node within a community was then compared to their community’s consensus sequence. The gene copy community assignments were projected onto the gene phylogenetic tree in order to better understand the evolutionary relationships between network communities.

#### j. Mobile element insertion analysis

##### Mobile element insertion (MEI) calling

We leveraged an enhanced version of PALMER (Pre-mAsking Long reads for Mobile Element inseRtion,^98^) to detect MEIs. Reference-aligned (to both GRCh38 Y and T2T Y) Y contigs from Verkko assembly were used as input. PALMER identified putative non-reference insertions (i.e., L1, SVA or Alu elements) using a pre-masking module based on a library of mobile element sequences. PALMER then identifies the hallmarks of retrotransposition events induced by target-primed reverse transcription, including target site duplication (TSD) motifs, transductions, and poly(A) tract sequences, and etc. Further manual inspection was carried out based on the information of large inversions, SVs, heterochromatic regions, and concordance with the Y phylogeny. Low confidence calls overlapping with large SVs or discordant with the Y phylogeny were excluded, and high confidence calls were annotated with further genomic content details.

In order to compare the ratios of non-reference mobile element insertions from the Y chromosome to the rest of the genome the following approach was used. The size of the GRCh38 Y reference of 57.2 Mbp was used, while the total GRCh38 reference sequence length is 3.2 Gbp. At the whole genome level, this results in a ratio for non-reference *Alu* elements of 0.459 per Mbp (1470/3.2 Gbp) and for non-reference LINE-1 of 0.066 per Mbp (210/3.2 Gbp)^19^. In chromosome Y, the ratio for non-reference *Alu* elements and LINE-1 is 0.315 per Mbp (18/57.2 Mbp) and 0.122 per Mbp (7/57.2 Mbp), respectively. The ratios within the MEI category were compared using the Chi-square test.

#### k. Gene annotation

##### Genome Annotation - liftoff

Genome annotations of chromosome Y assemblies were obtained by using T2T Y and GRCh38 Y gff annotation files using liftoff ^99^.

`liftoff -db $dbfile -o $outputfile.gff -u $outputfile.unmapped –dir $outputdir -p 8 -m $minimap2dir -sc 0.85 -copies $fastafile –cds $refassembly`

To evaluate which of the GRCh38 Y protein-coding genes were not detected in Verkko assemblies, we selected genes which were labeled as “protein_coding” from the GENCODEv41 annotation file (i.e., a total of 63 protein-coding genes) We compared protein coding genes’ open reading frames (ORFs) to the open reading frames obtained from Ensembl 109 to check whether any pseudogenes were miscategorized. First, exon sequence coordinates were collected from liftoff results. Then, transcripts which have the highest sequence identity were selected and used for evaluating ORFs. Concatenated exon fasta sequences were uploaded to AliView (v1.28)^100^. Exon sequences were aligned using “Realign everything” option and sequences were translated using a built-in tool.

#### l. Methylation analysis

Read-level CpG DNA-methylation (DNAme) likelihood ratios were estimated using nanopolish version 0.11.1. Nanopolish (https://github.com/jts/nanopolish) was run on the alignment to GRCh38 for all samples including the two Genome in a bottle samples^101^ (28 HGSVC, 15 HPRC and 2 GIAB (NA24385/HG002 and NA24149/HG003), totalling 45 samples before QC), for the three complete assemblies (HG00358, HG01890, HG02666) we additionally mapped the reads back to the assembled Y chromosomes and performed a separate nanopolish run. Based on the GRCh38 mappings we first performed sample quality control (QC). We find four samples, (NA19331, NA19347, HG03009 and HG03065), with genome wide methylation levels below 50%, which were QCed out. Using information on the multiple runs on some samples we observed a high degree of concordance between multiple runs from the same donor, average difference between the replicates over the segments of 0.01 [0-0.15] in methylation beta space.

After QC we leverage pycoMeth to *de novo* identify interesting methylation segments on chromosome Y. pycoMeth (version 2.2)^102^ Meth_Seg is a Bayesian changepoint-detection algorithm that determines regions with consistent methylation rate from the read-level methylation predictions. Over the 139 QCed flow cells of the 41 samples, we find 2,861 segments that behave consistently in terms of methylation variation in a sample. After segmentation we derived methylation rates per segment per sample by binarizing methylation calls thresholded at absolute log-likelihood ratio of 2.

To test for methylation effects of haplogroups we first leveraged the permanova test, implemented in the R package vegan^103,104^, to identify the impact of “aggregated” haplotype group on the DNAme levels over the segments. Because of the low sample numbers per haplotype group we aggregated haplogroups to meta groups based on genomic distance and sample size. We aggregated A,B and C to “ABC”, G and H to “GH”, N and O to “NO”, and Q and R to “QR”. The E haplogroup and J haplogroup were kept as individual units for our analyses. In the analysis we correct for sequencing center and global DNAme levels. Next to Y haplogroups we assessed the link between chromosome Y assembly length and global DNAme levels, either genome wide or on chromosome Y, using the permanova test and linear models. In the permanova test we added chromosome Y assembly length as an extra explanatory variable. The linear model was built up like the permanova test, correcting for the sequencing center and haplogroup. Next to the chromosome Y global analysis we also tested individual segments for differential meta-haplogroup DNAmeusing the Kruskal Wallis test. Regions with FDR<=0.2, as derived from the Benjamini-Hochberg procedure, are reported as DMRs. For follow up tests on the regions that are found to be significantly different from the Kruskal Wallis test we used a one versus all strategy leveraging a Mann–Whitney U test.

Next to assessing the effects of haplogroup and chromosome Y length we also tested for local methylation Quantitative Trait Loci (*cis*-meQTL) using limix-QTL^105,106^. Specifically, we tested the impact of the genetic variants called on GRCh38 (see Methods **“Variant calling using *de novo* assemblies”**), versus the DNAme levels in the 2,861 segments discovered by pycoMeth. For this we leveraged an LMM implemented in limixQTL, methylation levels were arcsin transformed and we leveraged population as a random effect term. Variants with a MAF >10% and a call rate >90%, leaving 11,226 variants to be tested. For each DNAme segment we tested variants within the segment or within 100,000 bases around it. Yielding a total of 245,131 tests. Using 1,000 permutations we determined the number of independent tests per gene and P values were corrected for this estimated number of tests using the Bonferroni procedure. To account for the number of tested segments we leveraged a Benjamini-Hochberg procedure over the top variants per segment to correct for this.

#### m. Expression analysis

Gene expression quantification for the HGSVC^19^ and the Geuvadis dataset^26^ was derived from the^19^. In short, RNA-seq QC was conducted using Trim Galore! (v0.6.5)^107^ and reads were mapped to GRCh38 using STAR (v2.7.5a)^108^, followed by gene expression quantification using FeatureCounts (v2)^109^. After quality control gene expression data is available for 210 Geuvaids males and 21 HGSVC males.

As with the DNAme analysis we leveraged the permanova test to quantify the overall impact of haplogroup on gene expression variation. Here we focused only on the Geuvadis samples initially and tested for the effect of the signal character haplotype groups, specifically “E”, “G”, “I”, “J”,”N”,”R” and “T”. Additionally we tested for single gene effects using the Kruskal Wallis test, and the Mann–Whitney U test. For *BCORP1* we leveraged the HGSVC expression data to assess the link between DNAme and expression variation.

#### n. Iso-Seq data analysis

Iso-Seq reads were aligned independently with minimap v2.24 (-ax splice:hq -f 1000) to each chrY Verkko assembly, as well as the T2T v2.0 reference including HG002 chrY, and GRCh38. Read alignments were compared between the HG002-T2T chrY reference and each *de novo* Verkko chrY assembly. Existing testis Iso-seq data from seven individuals was also analysed (SRX9033926 and SRX9033927).

#### o. Hi-C data analysis

We analysed 40/43 samples for which Hi-C data was available (Hi-C data is missing for HG00358, HG01890 and NA19705). For each sample, GRCh38 reference genome was used to map the raw reads and Juicer software tools (version 1.6)^110^ with the default aligner BWA mem (version: 0.7.17)^111^ was utilized to pre-process and map the reads. Read pairs with low mapping quality (MAPQ < 30) were filtered out and unmapped reads, such as abnormal split reads and duplicate reads, were also removed. Using these filtered read pairs, Juicer was then applied to create a Hi-C contact map for each sample. To leverage the collected chrY Hi-C data from these 40 samples with various resolutions, we combined the chrY Hi-C contact maps of these 40 samples using the *mega.sh* script^110^ given by Juicer to produce a “mega” map. Knight-Ruiz (KR) matrix balancing was applied to normalize Hi-C contact frequency matrices^112^.

We then calculated Insulation Score (IS)^113^, which was initially developed to find TAD boundaries on Hi-C data with a relatively low resolution, to call TAD boundaries at 10 kilobase resolution for the merged sample and each individual sample. For the merged sample, the FAN-C toolkit (version 0.9.23b4)^114^ with default parameters was applied to calculate IS and boundary score (BS) based on the KR normalized “mega” map at 10 kb resolution and 100 kb window size (utilizing the same setting as in the 4DN domain calling protocol)^115^. For each individual sample, the KR normalized contact matrix of each sample served as the input to the same procedure as in analysing the merged sample. The previous merged result was treated as a catalog of TAD boundaries in lymphoblastoid cell lines (LCLs) for chrY to finalize the location of TAD boundaries and TADs of each individual sample. More specifically, 25 kb flanking regions were added on both sides of the merged TAD boundary locations. Any sample boundary located within the merged boundary with the added flanking region was considered as the final TAD boundary. The final TAD regions were then derived from the two adjacent TAD boundaries excluding those regions where more than half the length of the regions have “NA” insulation score values.

The average and variance (maximum difference between any of the two samples) insulation scores of our 40 chrY samples were calculated to show the differences among these samples and were plotted aligned with methylation analysis and chrY assembly together. Due to the limited Hi-C sequencing depth and resolution, some of the chrY regions have the missing reads and those regions with “NA” insulation scores were shown as blank regions in the plot. Kruskal-Wallis (One-Way ANOVA) test (SciPy v1.7.3 scipy.stats.kruskal) was performed on the insulation scores (10 kb resolution) of each sample with the same 6 meta haplogroups classified in the methylation analysis to detect any associations between differentially insulated regions (DIR) and differentially methylated regions (DMR). Within each DMR, P values were adjusted and those insulated regions with FDR <= 0.20 were defined as the regions that are significantly differentially insulated and methylated.

#### p. Yq12 heterochromatin analyses

##### Yq12 partitioning

RepeatMasker (v4.1.0) was run using the default Dfam library to identify and classify repeat elements within the sequence of the Yq12 region^92^. The RepeatMasker output was parsed to determine the repeat organization and any recurring repeat patterns. A custom Python script that capitalized on the patterns of repetitive elements, as well as the sequence length between *Alu* elements was used to identify individual *DYZ2* repeats, as well as the start and end boundaries for each *DYZ1* and *DYZ2* array.

##### Yq12 *DYZ2* consensus and divergence

The two assemblies with the longest (T2T Y from HG002) and shortest (HG01890) Yq12 subregions were selected for *DYZ2* repeat consensus sequence building. Among all *DYZ2* repeats identified within the Yq12 subregion, most (sample collective mean: 46.8%) were exactly 2,413 bp in length. Therefore, five-hundred *DYZ2* repeats 2,413 bp in length were randomly selected from each assembly, and their sequences retrieved using Pysam (version 0.19.1)^116^, (https://github.com/pysam-developers/pysam). Next, a multiple sequence alignment (MSA) of these five-hundred sequences was performed using Muscle (v5.1)^117^. Based on the MSA, a *DYZ2* consensus sequence was constructed using a majority rule approach. Alignment of the two 2,413 bp consensus sequences, built from both assemblies, confirmed 100% sequence identity between the two consensus sequences. Further analysis of the *DYZ2* repeat regions revealed the absence of a seven nucleotide segment (‘ACATACG’) at the intersection of the *DYZ2* HSATI and the adjacent *DYZ2* AT-rich simple repeat sequence. To address this, ten nucleotides downstream of the HSAT I sequence of all *DYZ2* repeat units were retrieved, an MSA performed using Muscle (v5.1)^117^, and a consensus sequence constructed using a majority rule approach. The resulting consensus was then fused to the 2,413 bp consensus sequence creating a final 2,420 bp *DYZ2* consensus sequence. *DYZ2* arrays were then re-screened using HMMER (v3.3.2) and the 2,420 bp *DYZ2* consensus sequence.

In view of the AT-rich simple repeat portion of *DYZ2* being highly variable in length, only the *Alu* and HSATI portion of the *DYZ2* consensus sequence was used as part of a custom RepeatMasker library to determine the divergence of each *DYZ2* repeat sequence within the Yq12 subregion. Divergence was defined as the percentage of substitutions in the sequence matching region compared to the consensus. The *DYZ2* arrays were then visualized with a custom Turtle (https://docs.python.org/3.5/library/turtle.html#turtle.textinput) script written in Python. To compare the compositional similarity between *DYZ2* arrays within a genome, a *DYZ2* array (rows) by *DYZ2* repeat composition profile (columns; *DYZ2* repeat length + orientation + divergence) matrix was constructed. Next, the SciPy (v1.8.1) library was used to calculate the Bray-Curtis Distance/Dissimilarity (as implemented in scipy.spatial.distance.braycurtis) between *DYZ2* array composition profiles^118^. The complement of the Bray-Curtis dissimilarity was used in the visualization as typically a Bray-Curtis dissimilarity closer to zero implies that the two compositions are more similar (**Fig. 4e, S74**).

##### Yq12 *DYZ1* array analysis

Initially, RepeatMasker (v4.1.0) was used to annotate all repeats within *DYZ1* arrays. However, consecutive RepeatMasker runs resulted in variable annotations. These variable results were also observed using a custom RepeatMasker library approach with inclusion of the existing available *DYZ1* consensus sequence^3^. In light of these findings, *DYZ1* array sequences were extracted with Pysam, and then each sequence underwent a virtual restriction digestion with HaeIII using the Sequence Manipulation Suite^119^. HaeIII, which has a ‘ggcc’ restriction cut site, was chosen based on previous research of the *DYZ1* repeat in monozygotic twins^120^. The resulting restriction fragment sequences were oriented based on the sequence orientation of satellite sequences within them detected by RepeatMasker (base Dfam library). A new *DYZ1* consensus sequence was constructed by retrieving the sequence of digestion fragments 3,569 bp in length (as fragments this length were in the greatest abundance in 6 out of 7 analysed genomes), performing a MSA using Muscle (v5.1), and then applying a majority rule approach to construct the consensus sequence.

To classify the composition of all restriction fragments a k-mer profile analysis was performed. First, the relative abundance of k-mers within fragments as well as consensus sequences (*DYZ18*, 3.1 kbp, 2.7 kbp, *DYZ1*) were computed. A k-mer of length 5 was chosen as *DYZ1* is likely ancestrally derived from a pentanucleotide^4,121^. Next, the Bray-Curtis dissimilarity between k-mer abundance profiles of each fragment and consensus sequence was computed, and fragments were classified based on their similarity to the consensus sequence k-mer profile (using a 75% similarity minimum) **(Fig. S63).** Afterwards, the sequence fragments with the same classification adjacent to one another were concatenated, and the fully assembled sequence was provided to HMMER (v3.3.2) to detect repeats and partition fragment sequences into individual repeat units^67^. The HMMER output was filtered by E-value (only E-value of zero was kept). Once individual repeat units (*DYZ18*, 3.1 kbp, 2.7 kbp, and *DYZ1*) were characterized (**Fig. S64**), the Bray-Curtis dissimilarity of their sequence k-mer profile versus the consensus sequence was computed and then visualized with the custom Turtle script written in Python **(Fig. S65).** A two-sided Mann-Whitney U test (SciPy v1.7.3 scipy.stats.mannwhitneyu^118^) was utilized to test for differences in length between *DYZ1* and *DYZ2* arrays for each sample with a completely assembled Yq12 region (n=7) (T2T Y HG002:MWU=541.0, pvalue=0.000786; HG02011:MWU=169.0, pvalue=0.000167; HG01106:MWU=617.0, pvalue=0.038162; HG01952:MWU=172.0, pvalue=0.042480; HG01890:MWU=51.0, pvalue=0.000867; HG02666:MWU=144.0, pvalue=0.007497; HG00358:MWU=497.0, pvalue=0.008068;) (**Fig. 4b**). A two-sided Spearman rank-order correlation coefficient (SciPy v1.7.3 scipy.stats.spearmanr^118^) was calculated using all samples with a completely assembled Yq12 (n=7) to measure the relationship between the total length of the analysed Yq12 region and the total *DYZ1* plus *DYZ2* arrays within this region (correlation=0.90093, p-value=0.005620) (**Fig. S71**). All statistical tests performed were considered significant using an alpha=0.05.

##### Phylogenetic Tree Analysis of *DYZ2* Consensus Sequences

We retrieved *DYZ2* repeat sequence, which was previously identified on all other human acrocentric chromosomes (13, 14, 15, 21, and 22)^42^, for our phylogenetic analyses from CHM13 using BLAST^74^. More specifically, we queried the T2T-CHM13v1.1 reference genome^31^ using our *DYZ2* consensus sequence and retrieved all matches with an e-value of zero and greater than 500 nucleotides in length using SAMtools^52^. Next, we performed a multiple sequence alignment (MSA), for each chromosome separately, using MUSCLE^117^. We then generated a chromosome specific consensus sequence using a majority rule approach. To reflect sequence variation within the Yq12 heterochromatin region, we also constructed two Y chromosome *DYZ2* consensus sequences. One Yq12 *DYZ2* consensus sequence was built from *DYZ2* repeat sequences originating from arrays outside of the Yq12 inversions (i.e., end blocks). The second consensus sequence was constructed from *DYZ2* sequences located within arrays internal to the Yq12 inversions (i.e., middle blocks). Next, we performed an MSA of all *DYZ2* consensus sequences using MUSCLE. We elected to use only the HSATI and *Alu* portions of the *DYZ2* consensus sequences as the AT-rich simple repeat region was highly variable across consensus sequences. Next, a phylogenetic tree was inferred using maximum likelihood from the MSA with IQ-Tree (a GTR + Gamma model was used)^94^. IQ-Tree generated 1000 bootstrap (UFboot^122^) replicates from which an unrooted consensus tree was generated. The phylogenetic tree was then rooted at the midpoint and visualized using FigTree [http://tree.bio.ed.ac.uk/software/figtree/].

##### Yq12 Mobile Element Insertion Analysis

RepeatMasker output was screened for the presence of additional transposable elements, in particular mobile element insertions (MEIs). Putative MEIs (i.e., elements with a divergence <4%) plus 100 bp of flanking were retrieved from the respective assemblies. Following an MSA using Muscle, the ancestral sequence of the MEI was determined and utilized for all downstream analyses, (This step was necessary as some of the MEI duplicated multiple times and harboured substitutions). The divergence, and subfamily affiliation, were determined based on the MEI with the lowest divergence from the respective consensus sequence. All MEIs were screened for the presence of characteristics of target-primed reverse transcription (TPRT) hallmarks (i.e., presence of an A-tail, target site duplications, and endonuclease cleavage site)^123^.

##### Application of T2T-CHM13-derived repeat annotation pipeline on 43 assembled chrYs

Repeat discovery and annotation were performed on HG002 chrY^124^ following the pipeline laid out in Hoyt et al. 2022^125^. Subsequently, this repeat annotation compilation pipeline was applied to the 43 assembled chrYs presented herein for annotation with RepeatMasker4.1.2-p1of all known repeats using the Dfam3.3 library^92^ as well as CHM13 and HG002 chrY derived repeat models (noted in ^125^ and ^124^). Repeat annotations were summarized for all 44 assembled chrYs at the repeat class level using RepeatMasker’s script, buildSummary.pl^92^ with a corresponding genome file denoting the size of the assembly (base pairs) and reported in **Table S30**.

##### Regional repeat assessment of four fully assembled chrYs (including HG002)

Four chrYs were completely assembled from PAR1 to PAR2 (see **Table S9**) providing an opportunity to compare repeat variation within and between Y-chromosomal subregions. Therefore, Y-chromosomal subregions were extracted from the RepeatMasker compilation output (containing known and new repeat models) and summarized with buildSummary.pl^92^ with a corresponding genome file denoting the length of the region to be summarized. Repeat classes were summarized per region based on base pair composition, rather than counts, and similar regions were combined (e.g., PAR1 + PAR2 = PARs) and presented in **Fig. S33** and reported in **Table S29**.

##### BLAST estimates of *DAZ* and *RBMY1* composite repeat copy number

BLAST custom databases were generated from all chrY assemblies and used to detect instances of the *DAZ* and *RBMY1* composite repeat units per assembly. The consensus sequences for these two composite repeat units were derived from HG002 chrY, and are reported in **Table S43** and in^124^. The RBMY1 composite repeat unit contains the whole gene, while that of *DAZ* lies within the gene. Due to the fact that composite repeats are composed of three or more repeating sequences (i.e., TEs, satellites, composite subunits, simple/low complexity repeats) as defined in^125^, of which are scattered throughout the genome, we required at least an 85% length match to detect predominantly full-length copies while still allowing for variation in the ends. While this requirement for length matching prevents the detection of individual repeats within the composite from being counted as a composite, it does have the limitation of not detecting a full-length copy, as polymorphic TE insertions may interfere. Copy number estimate results for all 44 chrYs are reported in **Table S43-S44.**

### 6. Statistical analysis and plotting

All statistical analyses in this study were performed using R (http://CRAN.R-project.org/) and Python (http://www.python.org). The respective test details such as program or library version, sample size, resulting statistics and p-values are stated in the running text. Figures were generated using R and Python’s Matplotlib (https://matplotlib.org), seaborn^72^ and the “Turtle” graphics framework (https://docs.python.org/3/library/turtle.html).

### 7. Data Availability

All data generated are available via the HGSVC data portal at https://ftp.1000genomes.ebi.ac.uk/vol1/ftp/data_collections/HGSVC2/working/ and https://ftp.1000genomes.ebi.ac.uk/vol1/ftp/data_collections/HGSVC3/working/. HPRC year 1 data files, PacBio HiFi, ONT long-read sequencing and Bionano Genomics optical mapping and data files were downloaded from the following url: https://humanpangenome.org/year-1-sequencing-data-release/. Large supplementary data files such as the assembled genomes are available at ftp://ftp.1000genomes.ebi.ac.uk/vol1/ftp/data_collections/HGSVC3/working/20230412_sigY_assemb ly_data .

### 8. Code Availability

Project code implemented to produce the assemblies and the basic QC/evaluation statistics is available at github.com/marschall-lab/project-male-assembly. All scripts written and used in the study of the Yq12 subregion are available at https://github.com/Markloftus/Yq12.

## Supporting information

Supplementary Results and Figures

## Author contributions

PacBio production sequencing: Q.Z., K.M.M., A.P.L., J.K.; ONT production: Q.Z., K.H.; Strand-seq production: P.Hasenfeld., J.O.K.; ONT re-basecalling and methylation calling: P.A.A., W.T.H.; Genome assembly: P.E., F.Y., T.M.; Assembly analysis and evaluation: P.E., P.H., F.Y., W.H., F.T.; Assembly-based variant calling: P.E., P.A.A., P.H., C.R.B.; Variant QC, merging, and annotation: P.A.A., P.H.; Short-read calling, phylogeny construction and dating: P.H.; Analysis of Bionano Genomics optical maps: F.Y.; Strand-seq inversion detection and genotyping: D.P.; MEI discovery and integration: W.Z., M.L., M.K.K.; Inversion analysis: P.H., D.P., K.K., M.L., M.K.K.; Gene conversion and evolutionary rate: K.K., P.H., M.K.K.; Gene families: M.L., M.K.K.; Analyses on Y subregions: P.E., P.H., M.L., F.Y., G.A.L., P.A.A., W.H., K.K., F.T., M.K.K., E.E.E., C.L.; RNA-seq analysis: M.J.B.; Methylation and meQTL analysis: M.J.B.; HiC analysis: C.Li., X.S.; Repeat annotation: S.J.H., R.J.O.; Iso-Seq analysis: P.D., E.E.E.; Gene annotations F.Y., P.D.; Supplementary materials: P.H., P.E., M.L., F.Y., P.A.A., G.A.L., M.J.B., W.Z., W.H., K.K., C.Li, S.J.H., P.D., F.T., J.Y.K., Q.Z., K.M.M., P.Hasenfeld, X.S., M.K.K.; Display items: P.H., P.E., M.L., F.Y., G.A.L., W.H., K.K., F.T., M.K.K.; Manuscript writing: P.H., P.E., M.L., P.A.A., G.A.L., M.J.B., W.Z., M.K.K., C.L. with contributions from all other authors. All authors contributed to the final interpretation of data. HGSVC Co-chairs: C.L., J.O.K., E.E.E., T.M.

## Acknowledgements

We thank Arang Rhie and Adam Phillippy for coordination and discussions; Dr. Yali Xue for discussions and advice throughout the project; Jonathan Wood and the Genome Reference Informatics Team at the Wellcome Sanger Institute for suggestions and feedback on assembly evaluation; Laurits Skov for advice and sharing his scripts for gene conversion detection; the Human Pangenome Reference Consortium (HPRC, https://humanpangenome.org) for making their data publicly available; the Centre for Information and Media Technology at Heinrich Heine University Düsseldorf and the Scientific Services at the Jackson Laboratory including the Genome Technologies Service for their expert assistance with the work described herein and Research IT for providing computational infrastructure and support and the Phillippy Lab (NIH/NHGRI) for their comprehensive Verkko support. We are grateful to the people who generously contributed samples as part of the 1000 Genomes Project.

## Consortia

The members of the Human Genome Structural Variation Consortium (HGSVC) are Haley J. Abel, Hufsah Ashraf, Peter A. Audano, Anna O. Basile, Christine Beck, Marc Jan Bonder, Harrison Brand, Marta Byrska-Bishop, Mark J.P. Chaisson, Yu Chen, Ken Chen, Zechen Chong, Nelson T. Chuang, Wayne E. Clarke, André Corvelo, Scott E. Devine, Peter Ebert, Jana Ebler, Evan E. Eichler, Uday S. Evani, Susan Fairley, Paul Flicek, Sky Gao, Mark B. Gerstein, Maryam Ghareghani, Ira M. Hall, Pille Hallast, William T. Harvey, Patrick Hasenfeld, Alex R. Hastie, Wolfram Höps, PingHsun Hsieh, Sarah Hunt, Miriam K. Konkel, Jan O. Korbel, Sushant Kumar, Charles Lee, Alexandra P. Lewis, Chong Li, Bin Li, Yang I. Li, Jiadong Lin, Mark Loftus, Tsung-Yu Lu, Rebecca Serra Mari, Tobias Marschall, Ryan E. Mills, Zepeng Mu, Katherine M. Munson, David Porubsky, Benjamin Raeder, Tobias Rausch, Allison A. Regier, Jingwen Ren, Bernardo Rodriguez-Martin, Ashley D. Sanders, Martin Santamarina, Xinghua Shi, Chen Song, Oliver Stegle, Michael E. Talkowski, Luke J. Tallon, Jose M.C. Tubio, Aaron M. Wenger, Xiaofei Yang, Kai Ye, Feyza Yilmaz, Xuefang Zhao, Weichen Zhou, Qihui Zhu, and Michael C. Zody.

