## Supplementary Results and Figures for "Assembly of 43 diverse human Y chromosomes reveals extensive complexity and variation"

Table of contents


Supplementary Results 2
- *De novo* assembly evaluation 2
- Effect of input read characteristics on assembly contiguity 4
- Orthogonal support to Y-chromosomal SVs and copy number variation 5
- Genetic variation of PAR1 region 6
- Gene annotation 7
- Gene family architecture and evolution 8
- Y-chromosomal inversions 12
- Yq12 heterochromatic subregion 16
  - A Yq12 overview 16
  - Yq12 *DYZ1* and *DYZ2* repeat analyses 17
  - Yq12 mobile element insertions (MEIs) 19
- Functional analysis 20

Supplementary Figures 23-112

References 113

### Supplementary Results

#### *De novo* assembly evaluation

We evaluated a series of contig alignments, in which we combined different samples and assemblies as target and query for the alignment process. We selected all contig alignments for the respective assemblies to the T2T Y reference as a baseline, and ran the following experiments (**Figure S76**, from left to right): pairing the two closely related African samples NA19317 and NA19347 (with the TMRCA estimated to be only 200 years ago (ya) [95% highest posterior density [HPD] interval: 0 - 500 ya] and therefore considered as quasi-replicates); considering the four pairs of high- and lower-coverage assemblies; aligning the Verkko hybrid assemblies to HiFi-only assemblies built with hifiasm v0.16.1-r375[1], and generating self-alignments. In all these scenarios, the collected statistics support the view that the Verkko assemblies have been robustly assembled and contain sample-specific sequences. For example, the fraction of the query sequences aligned with the maximal mapping quality (MAPQ) of 60 is highest for the self-alignments (**Figure S76**, middle row), followed by the quasi-replicate African pair and the alignments to the HiFi-only hifiasm assemblies. The alignments of the high- and lower-coverage pairs (**Figure S76**, middle row, blue boxes) show a drop relative to the aforementioned combinations, which is consistent with the higher error rate for the lower-coverage assemblies (**Figure S6**). Next, we checked the overall (dis-)similarity of the Y assemblies with respect to their k-mer content (**Methods**). Relative to the GRCh38 Y assembly, all our assemblies plus the T2T Y show a coherent behavior, sharing a substantial fraction of their constituent k-mers (**Figure S77**). Notably, the four pairs of high and lower coverage assemblies do not exhibit any inconsistencies, suggesting that, despite elevated error rate and increased fragmentation at lower coverages, the assemblies still represent a sample-specific Y chromosome.

We investigated the locations of assembly gaps via aligning all identified Y contigs to the GRCh38 and CHM13 plus T2T Y reference sequences and assessing the Y-chromosomal alignments via alignment coverage (**Figs. S78-82, Methods**). Contigs aligning well to the reference sequence were expected to show coverage of 1, while assembly gaps and poor alignments due to misalignment, misassembly or structural differences between the assembly and reference sequence should show no or >1 coverage. The results highlight the Yq12 heterochromatin as the most poorly aligning subregion (**Figure S80**), consistent with the presence of the highest number of Y assembly breaks across samples (**Tables S14, S16**) and an overall high variation in the composition of this region across samples (**Figure 4f**). In the (peri-)centromeric region majority of the poor alignments were localized to the highly repetitive centromeric *DYZ3* α-satellite array (**Figure S82**). In euchromatic regions the PAR1 and ampliconic subregion 7 were the most challenging to contiguously assemble. The majority of the poorly aligning regions in ampliconic region 7 overlap with the P1 palindrome (**Figure S79**) composed of ~1.45 Mbp inverted segmental duplications 99.97% sequence identity[2], while in the PAR1 the poorly aligning regions are broadly distributed (**Figure S81**).

We complemented the alignment-based detection of assembly gaps with the recently introduced tool Rukki (as part of the Verkko v1.2 release package). Leveraging orthogonal information such as parental k-mers, Rukki can identify phasing paths in the complete assembly graph, thereby resolving more complex structures ("bubbles") into linear assemblies. We mimicked the necessary input data for Rukki using the identified chromosome X and Y contigs (**Methods**), and obtained an average gap size estimate of 48,985 bp (homopolymer-compressed) for a total of 136 paths through the assembly graph, leaving 1264 contigs disconnected. The average gapless length of the paths amounted to 98.85% of their total length. We note here that, by resolving bubbles in the graph, Rukki could also connect several contigs without introducing gaps, which implies that the actual fragmentation of our assemblies is slightly lower than what we had initially assumed (**Table S6**).

We also compared in more detail the assemblies of the closely related pair of African Y chromosomes from NA19317 and NA19347, assembled to a similar level of contiguity (NA19317 contiguously assembled from PAR1 to Yq12 in a single contig, while NA19347 has an additional break at the (peri-)centromeric region). In agreement with the TMRCA estimate, the Y assemblies show high similarity in structure and sequence (**Figure S4, Table S8**). Across 23.96 Mbp (PAR1, (peri-)centromeric, Yq12 and PAR2 regions were excluded as either not contiguously assembled or recombine with the X chromosome), only 233 nucleotide substitutions and 583 indels summing to a total of 976 base pairs were identified, translating to a sequence identity of 99.9959% (**Table S8**). A total of 286/583 indels represent expansions and contractions at polynucleotide tracts and short tandem repeats (STRs). This was further supported by structural variants called by PAV (**Table S21**). A total of 23 SVs were called where these two samples differ from each other, all localized either in tandem repeat regions or repeat arrays (i.e., centromeric region, *DYZ19*, *DYZ18* and Yq12) with higher mutation rates than the more unique regions of the genome.

In addition, we used the Bionano optical genome mapping (OGM) data to evaluate the quality of the Verkko *de novo* Y-chromosomal assemblies. For 18/43 samples, no inconsistencies were identified between the Verkko and the OGM assemblies (**Table S48**). For the remaining 25/43 samples, 94 inconsistencies were identified, however, analysis of single optical mapping DNA molecules confirmed that 82% (77/94) were correctly resolved in Verkko assemblies. For the remaining 17/94 inconsistent sites (from 10 samples), the accuracy could not be evaluated due to the lack of single optical mapping DNA molecule data spanning these sites. Taken together, the orthogonal validation using optical mapping data identified no inconsistencies in the Verkko assemblies.

We performed an additional assembly evaluation step by aligning single optical mapping molecules to the Verkko assemblies (**Methods**). This approach identified a total of 2,351 10-kbp windows in 43 samples which were not covered by optical mapping molecules (**Table S49**). Most of these windows (1,798/2,351; in 43/43 samples) overlapped with PAR1, PAR2, (peri-)centromeric and Yq12 heterochromatic subregions. Additionally, 300/2,351 windows (in 26/43 samples) overlapped with ampliconic regions (more specifically

AMPL1, AMPL2, AMPL5, AMPL6 and AMPL7), all of which are challenging to contiguously assemble due to their sequence composition. Only a small proportion of windows (253/2,351 windows) from 3/43 samples overlapped with X-transposed and X-degenerate regions (XTR1, XDR1 and XDR3; **Table S49**), highlighting regions that would require further investigation. It is important to keep in mind that while optical mapping data offers an independent orthogonal validation for the generated assemblies, it loses resolution at heterochromatic regions (due to the lack of restriction enzyme cutting sites) and can struggle to correctly characterize highly repetitive and complex genomic regions.

Last, we used several tools to assess the base-level correctness of our assemblies on the basis of consistently occurring rare k-mers (VerityMap[3]) and of heterozygous positions in the sequence (DeepVariant[4], PEPPER[5], NucFreq[6]; **Methods**). Since NucFreq's way of identifying potentially erroneous regions is inherently dependent on precise read-to-assembly alignments, which is challenging in large parts of the Y chromosomes such as the (peri-)centromere and the Yq12/HET, we first checked how many regions flagged by NucFreq were located in those sequence contexts. NucFreq called a median of 37 regions (median size 69,574 bp) per sample (mean 63, size 166,366 bp), of which ~81.9% (median; mean ~69.4%) overlapped with said region types (**Table S12**). Additionally, as part of a more detailed analysis of PAR1 in 10 samples (**Table S11**), we manually curated both VerityMap- and NucFreq-flagged regions and frequently observed false positives in areas of HiFi coverage dropout, likely caused by elevated levels of GA/TC dinucleotides in the sequence [7–9]. We thus implemented a clustering strategy to identify parts of the assemblies where both VerityMap and NucFreq flagged nearby regions (**Methods**). This procedure resulted in a median of ~99.987% (mean ~99.69%) of the assembled sequence not being part of a merged VerityMap/NucFreq cluster. The more lenient setting counting all regions as indicators of potential assembly errors resulted in estimates of ~99.814% (median) and ~99.533% (mean) of the assembled sequence not being flagged for spurious signal.

#### Effect of input read characteristics on assembly contiguity

We explored the potential effect of the varying input read set characteristics, such as genomic coverage or read length N50, on the outcome of the hybrid assembly process. First, we randomly selected four of the high-coverage samples (HG02666, HG01457, NA19384, NA18989; HiFi coverage at least 50×, in the following denoted with the prefix "HC" for high coverage where needed to disambiguate) and re-assembled those with about half of the available HiFi reads, i.e., using around 30× coverage, which is comparable to most of the HiFi datasets used in this study (**Tables S1-S2**). The lower-coverage assemblies show higher fragmentation as indicated by a considerably larger number of assembled contigs and a smaller contig NG50 statistic (**Tables S4-S5**). This observation is compatible with the assumption that higher input read coverage has a positive effect on assembly quality in terms of contiguity. However, given that not all Y chromosomes of the high-coverage samples could be assembled contiguously from telomere to telomere (**Tables S5, S9**), it is evident that this factor alone is not sufficient as an explanatory variable. Moreover, for all four lower-coverage assemblies, the total assembled length of the Y sequence is increased by two to six megabases compared to their high-coverage counterparts, which suggests that the total assembly length may be of limited value when comparing Y assemblies created with substantially different HiFi input coverage (**Table S5**). We deepened our analysis by training multivariate regression models (**Methods**) to investigate the relationship between the input read set and quality-related assembly statistics of interest such as the contig NG50. For this analysis, we augmented our dataset with the four lower-coverage assemblies described above. The results of the regression analysis confirmed that HiFi input coverage and mean ONT-UL read length are relevant factors to achieve higher contig NG50 values (**Tables S50-S51**), yet cannot be sufficient as explained above. Given the small size of our dataset from a statistical point of view, e.g., including only two samples from haplogroup A (HG02666, HG01890), and these two Y chromosomes could be assembled in a single contig from telomere to telomere, it is challenging to derive a robust statement about the factors governing overall assembly quality.

It is possible that additional confounding factors originate from differences, e.g., in cell line culture such as varying numbers of passages. These details are commonly not known for the biological material at hand and can thus also not be assessed in the standard library QC.

#### Orthogonal support to Y-chromosomal SVs and copy number variation

In order to evaluate and validate the Y-chromosomal SVs and copy number variation (CNVs) identified from the *de novo* assemblies we used several independent approaches as described below.

We evaluated assembly-derived structural variants called with PAV (using the GRCh38 Y reference sequence, **Table S20-S21, Methods**) by using optical mapping data as an orthogonal support (**Methods**). The 29 evaluated variants included all 15 insertions and 14 deletions 5 kbp or larger in size. Across 21/29 SVs the genotype concordance between optical mapping data and PAV calls was 91% (87/96 calls, **Table S52**). The remaining 8/29 SVs were located in the (peri-)centromeric and proximal PAR2 regions where optical mapping does not have sufficient resolution.

The non-recombining nature of the male-specific regions of the Y chromosome offers the advantage that phylogenetically more closely related Y chromosomes are expected to be more similar. Therefore, the concordance of the genetic variants with the Y-chromosomal phylogeny offers strong support to the accuracy of both Y assemblies and the reported variants. Phylogenetically closely related samples show high levels of similarity in terms of the overall Y assembly size and sequence similarity (**Figures 2a, 3a, S4, S17, S19; Tables S8, S16**). For example, the assemblies of samples closely related to GRCh38 are structurally highly similar to the GRCh38 as one would expect (**Figures 3a, S14, S16**). Similar patterns are seen for complex regions, such as the centromere, the TSPY and *DYZ19* repeat arrays and the Yq11/Yq12 transition region (including the *DYZ18*, 3.1-kbp, 2.7-kbp repeat arrays) as phylogenetically closely related samples show high levels of similarity both in terms of repeat array sizes and structure (**Figure 3c**, **Figures S21-S27, S57, S59-60, S67-S69;**

**Tables S16-S18**). Inversion calls follow a similar pattern - despite high inversion recurrence rate (see details in main text and **Tables S32-S33**), closely related samples are similar in terms of their inversion calls.

The identified inversions are also well supported by previous reports (see Supplementary Results 'Y-chromosomal inversion' below for additional details). Additionally, the availability of Strand-seq data for 30/43 samples and the T2T Y, offered orthogonal support to the reported inversion calls. Across 10 inversions and confident Strand-Seq calls relative either to GRCh38 or the T2T Y as a reference, all 293/293 genotype calls were concordant with the assembly-based inversion calls (**Figure 3a, Table S32-S33**).

In addition, we compared our findings to previously published results either for the same samples, or phylogenetically closely related samples. For example, the *RBMY1* copy number estimates for 14 samples overlapping with[10] (that used read-depth information from low-coverage Illumina data) are highly concordant with 13/14 samples showing either exactly the same or plus 1 total *RBMY1* copy number estimates (**Table S53**). Across all 1,218 studied 1000 Genomes Project samples, the *RBMY1* gene copy number ranged from 3 to 13, with 35.5% of samples carrying 8 copies[10]. Our copy number estimates ranged from 5 to 11 copies, with 43.2% of samples carrying 8 copies, matching well with previous estimates. Additionally, samples with closely related Y lineages carried similar *RBMY1* copy numbers - for example we report 10 copies for HG00358 (N1a-Z1940) and 10 copies is known to be the most prevalent copy number in this haplogroup based on both droplet digital PCR and using Illumina read depth)[10].

Two of the assembled samples (HG00358, haplogroup N1c-Z1940 and NA18989, haplogroup C1a-CTS6678) carry rearrangements in the *AZFc*/ampliconic 7 subregion, known to be present in those Y haplogroups - ~1.8 Mbp *b2/b3* deletion and a likely *gr/rg* duplication, respectively **(Figures 2a-b; Figures S10, S18 and S35a**)[11–13]

#### Genetic variation of PAR1 region

We investigated in more detail the genetic variation of the PAR1 subregion in 10 samples where this subregion was contiguously assembled and in the T2T Y (**Tables S14-S16**). In order to evaluate the PAR1 assembly quality, we first visually checked all regions highlighted by VerityMap (**Table S11**) in HiFi and ONT read alignments to the *de novo* assembly. The examination of the 5 regions flagged by VerityMap as potential assembly errors resulted in the conclusion that the 3 false positives were likely caused by known sequencing biases in PacBio HiFi data (GA/TC-rich regions).

The 10 assembled and the T2T Y PAR1 regions ranged in size from 2,398,871 to 2,460,891 bp (mean 2,430,552 bp), with the largest difference of approximately 62 kbp in size (**Table S16)**. Approximately than half of the size variation across samples is accounted for by recently described satellite region (e.g., pasiphae and kalyke) localized close to the telomere, showing up to 55-fold differences in size across samples (**Figure S31**; **Table S55**)[9,14].

Our results match well with previous analysis reporting higher genetic diversity at the PAR1 region compared to the male-specific regions of the Y chromosome. The mean pairwise sequence identity in the 10 contiguously assembled males was 90.68% (range 75.83% to 97.58%, median 93.14%) and 99.88% (range 99.60% to 99.99%, median 99.93%) in PAR1 and the combined XDR regions (**Table S54)**, respectively. We also examined the variability of the PAR1 region by constructing graphs from the 10 *de novo* assemblies and the T2T reference (**Methods**) and plotted the total length of the sequence variants in large (>1 kbp) bubbles of the graph (**Figures S31-S32**). The first ~2 Mbp of the PAR1 region showed higher variability compared to the following ~4 Mbp of the chromosome Y sequence. This observation was consistent with chromosome X where Y sequence variants were only detectable in the first ~2 Mbp of the PAR1 region. Higher level of sequence similarity was also evident from the pairwise comparison of sequence identity both between the chrY PAR1 regions (**Figure S29**) and between the CHM13 chrX, T2T Y and the chrY and chrX PAR1 regions of the 10 samples where PAR1 was contiguously assembled (**Figure S30**). The chrY PAR1 exhibited a very distinct distribution and composition of transposable elements compared to the male-specific Y chromosome (**Figures S33-S44**; **Tables S29-S31**). While the PAR1 region wass distinguished by the enrichment in SINE elements, the male-speific region shows enrichment in LINE elements.

#### Gene annotation

To annotate genomes of *de novo* assemblies of 43 male samples, we used liftoff and GENCODEv41 GRCh38 annotations, and T2T-CHM13v2.0_RefSeq_Liftoff_v4 chrY annotations (**Table S41**). Annotation of 43 Y chromosomes presented a number of genes ranging from 580 (HG00358) to 758 (HG02666). The number of protein-coding genes ranged from 82 (HG00358 and HG03732) to 114 (HG02666), and the number of pseudogenes from 339 (HG00358) to 457 (NA18989) (**Table S41**). Majority of differences between GRCh38 Y and T2T Y annotations were due to previously unassembled regions, gaps, and ampliconic gene copy numbers (**Tables S38-S39, S42**). The single-copy protein-coding genes were present in all samples, except for 14 genes in PAR1, 1 gene in XDR1 and 1 gene in PAR2 in a total of 14 individuals, overlapping with poorly assembled regions in those individuals (**Tables S38-S42**). In addition, there are 8 multi-copy protein-coding gene families located in the ampliconic regions, 5 of which showed variation in copy number across the analysed samples. 3/8 protein-coding gene families (*VCY*, *PRY* and *HSFY*) showed a constant copy number (2 copies) across all samples. Two of the assembled samples (HG00358, haplogroup N1c-Z1940 and NA18989, haplogroup C1a-CTS6678) carry known rearrangements in the *AZFc*/ampliconic 7 subregion - ~1.8 Mbp *b2/b3* deletion and a likely *gr/rg* duplication, respectively[11–13] and show variation in copy number in genes (*DAZ*, *BPY2* and *CDY1*) affected by these rearrangements. The *BPY2* gene copy number ranges from 1 to 5 (41/44 samples show a constant copy number of 3), *CDY* from 3 to 5 copies (39/44 samples show a constant copy number of 4) and *DAZ* from 2 to 6 copies (42/44 samples show a constant copy number of 4). Note that the accuracy of gene annotation and copy number determination might be impacted by fragmented assembly in case of a few samples

(**Table S38**). The highest variation in copy number across the 43 *de novo* samples was observed for *RBMY* gene (from 5 to 11 copies, 6 copies in T2T Y) and *TSPY* (from 24 to 40, 47 copies in T2T Y; **Tables S38-S39**).

#### Gene family architecture and evolution

The human Y chromosome contains two protein-coding gene families known to be highly variable in copy number: the *TSPY* and the *RBMY1*.

The *TSPY* is one of the largest and most homogeneous tandemly organized protein-coding genes in the human genome[15]. A single *TSPY* gene is located within an approximately 20.3 kbp repeat unit, which are tandemly organized in head-to-tail fashion in a repeat array on the Yp arm. In addition, a single separate repeat unit (containing the *TSPY2* gene) in GRCh38 Y is located upstream of the repeat array[2]vary in copy number from 11 to 72 copies between individuals, with a peak of frequencies in the range of 21-35 copies. The *TSPY* gene copy number has been positively associated with sperm count and sperm concentration[16].

Here, we have analysed the TSPY repeat units, their composition and variation in more detail. First of all, analysis of the TSPY repeat unit locations revealed that in the majority of the samples (33/44, including the T2T Y) the single separate TSPY repeat unit is located in between the major repeat array and the centromere. In the rest, phylogenetically closely related haplogroup QR samples including the GRCh38 Y, the single copy resides upstream of the repeat array, likely a result of two inversions shared by these samples (**Figure S49** and see **Supplementary Results 'Y-chromosomal inversion'** below for additional details).

The size of the TSPY repeat array varied in size from 465 to 932 kbp (mean 677 kbp, median 688 kbp) and the respective repeat unit copy number in the repeat array ranged from 23 to 46 copies (mean 33, median 34 copies) across the T2T Y and 39 males where this region was contiguously assembled (**Figure 3c, Figure S51; Tables S11-S18, S38**).

The TSPY repeat array units showed similar patterns of sequence identity within samples. The single separate repeat unit containing the *TSPY2* gene was always the most diverse, followed by the two proximal copies in the repeat array (Figure S24). However, the overall sequence similarity between repeat units was high, with an average of 58% (range from 48.2% to 71.6%, median 58.3%) of all pairwise repeat unit comparisons within samples showing ≥99.5% sequence identity, while 76.1-87.4% (mean 82.8%, median 83.4%) showed ≥99% sequence identity (Figure S24, Table S18). Assuming that very high sequence identity between repeat units indicates recent expansion events within the array, we saw on average 4 repeat units (range from 0 to 17) per sample showing ≥99.9% sequence identity in pairwise comparisons. While smaller expansion events involving a single separate or two neighboring repeat units were seen in 39/40 samples, 13 samples also showed evidence of larger recent duplication events involving ≥3 consecutive repeat units with ≥99.9% sequence identity. A total of six samples (HG01928, HG01952, NA18534, NA19317, NA19347 and NA19705) carried 3 consecutive units (approx. 60.9 kbp duplication) with ≥99.9% sequence identity, four samples (HG00731, HG02011, HG02554 and HG02717) 4 consecutive units (approx. 81.1 kbp duplication) and one sample

(HG01890) 6 consecutive repeat units indicating a duplication event approximately 121.8 kbp in size. In addition, two samples have likely undergone 2 duplication events: HG02666 of 3 and 4 units (approx. 60.9 kbp and approx. 81 kbp duplications, respectively), T2T Y 4 and 5 consecutive units (approx. 81 kbp and 101.4 kbp, respectively) (Figures S24, S27).

We performed a closer investigation of three Y chromosome copy number variable ampliconic genes: *TPSY*, *RBMY1*, and *DAZ*. It should be noted that a subset of samples contained assembly breaks directly within the *TSPY* array (NA19239, HG03065, HG01258, HG00096), *DAZ* gene clusters (HG03065, HG00673), and *RBMY1* copies (HG03065). As such, those samples were excluded from further analyses of those genes. In our analyses, we utilized an exon-centric approach to identify all potential low divergence protein-coding copies of these genes (see **Methods**). As expected, the genomic architecture of the three genes was vastly different. While *RBMY1* copies are generally arranged into four separate groups/regions (median group distance: 250,495 bp, mean group distance: 614,057 bp), *DAZ* copies are organized within clusters, each containing two *DAZ* copies per cluster. *TSPY*, except for *TSPY2*, which is present as a single copy, exists as an array of tandemly repeated gene copies of variable copy number length (see above). We identified copy number variation across haplogroups for all three genes. On average, there were 8 *RBMY1* (max: 11, min: 5), 33 *TSPY* (max: 46, min: 23, counts include 1 copy of *TSPY2*), and 4 *DAZ* (max: 6, min: 2) gene copies per assembly (**Table S40**). Interestingly, our analyses revealed that nearly all *DAZ* genes (172 total copies) contained extensive exon copy number variation both across and within the 43 assemblies (41 assemblies, the T2T Y and GRCh38) (**Figure S44**). Just 7 of the 28 *DAZ* exons (exons: 1, 17, 18, 25, 26, 27, and 28) were considered '*stable*' as their copy number was found to vary little (< 3 gene copies across all samples had exon copy number variation) across *DAZ* gene copies, and only 3 of these exons (exons: 1, 25, and 26) were found in only one copy. In contrast, all other exons varied in their copy number with the most variation detected toward the distal end of the *DAZ* gene in exons 20-24 with exon 24 displaying the largest variation ranging from 0 – 14 (mean: 3.24, std: 3.12) copies. Interestingly, we noticed that copies of *DAZ* containing 4 duplications of exons 2 – 6 (25%; 43 of 172 *DAZ* copies) had the least variation in all other exons, and this *DAZ* copy was retained in all 43 assemblies. Due to this considerable sequence variation across *DAZ* copies, a multiple sequence alignment was uninformative for the highly variable regions, even with manual curation, resulting in low confidence of phylogenetic analyses. For this reason we elected not to pursue further analyses of the *DAZ* gene. Unlike *DAZ*, the other two analysed ampliconic genes did not exhibit exon copy-number variation. However, over a quarter (28.72%; 386 of 1344) of *TSPY* copies were found to contain a previously described 18 nucleotide duplication within the first exon[17].

Next, we constructed a maximum likelihood tree of all *TSPY* gene copies (nucleotide sequences; 1344 copies in total) from 41 (39 assemblies, T2T Y and GRCh38) samples with continuous assemblies in the TSPY region (**Figure S52**). A network-based clustering analysis discerned 32 distinct *TSPY* clusters (min cluster size: 5 copies, max cluster size: 272 copies) (see **Methods**). The network cluster assignments were considered satisfactory as most (90.6%) *TSPY* copies were pristine to the consensus sequence of their assigned *TSPY* cluster, and only 1.71% (23 of 1344) of all *TSPY* copies contained more than one SNV. Due to the high similarity across *TSPY* sequences, the majority of *TSPY* clusters were the result of only a few (mainly 1-2) unique SNVs (i.e., diagnostic nucleotides). The phylogenetic tree suggests that two of the largest *TSPY* clusters (network cluster 1: medium blue, network cluster 2: yellow) diverged early during Y chromosome radiation (**Figure S52**). We believe *TSPY* copies assigned to network cluster 1 are likely representative of the ancestral *TSPY* sequence for two reasons. The first reason being that a manual inspection of the *TSPY* multiple sequence alignment showed that *TSPY* copies assigned to network cluster 1 do not carry the diagnostic nucleotides of the other *TSPY* network clusters. Therefore, *TSPY* copies from network cluster 1 seem to be a '*baseline*' sequence. The second reason emanates from the results of a secondary network analysis of cluster consensus sequences (see **Methods** and the visualized network on **Figure S50**) which revealed that more *TSPY* clusters are similar to network cluster 1 than any other cluster. Interestingly, the phylogeny, as well as the secondary network analysis, indicates that *TSPY2* is more similar to the *TSPY* copies in network cluster 2 than to those in cluster 1 (**Figure S52**).

The composition of *TSPY* arrays within Y lineages were overall highly similar (**Figure S50**). Though, the two deepest-rooted assemblies (e.g., HG01890 and HG02666) appeared quite distinct (i.e., *TSPY* copies contained more SNVs and less cluster representation) compared to the other Y African lineage arrays suggesting possible concerted evolution of these arrays. The *TSPY* arrays from the two closest related samples (NA19317 and NA19347) were identical (Figure S3). Additionally, while all *TSPY* arrays terminated with a pseudogene (not visualized), most arrays harboured one additional pseudogene (mean: 0.87, max: 2, min: 0) (**Figure 3c**). We identified six separate truncating events across all samples, three of which are nonsense mutations, two are caused by a one nucleotide deletion, and one structural variation deleted the 5' region of *TSPY (*~370 nucleotides of the proximal half of exon 1) (**Figure 3c**). All nonsense mutations, but one, will likely result in nonsense-mediated decay. The nonsense mutation in HG03009 is in the last exon and terminates the N-terminus three amino acids prematurely.

The sequence pattern within *TSPY* arrays suggests frequent occurrence of rearrangements such as non-allelic homologous recombination and gene conversion. Most recombination/gene conversion events identified are retained within lineages and constitute entire clusters (**Figure S50**, Network Clusters: 4, 7, 8, 11, 12, 13, 16, 17, 18, 21), although some of these events are sample specific. For instance, within the T2T assembly (HG002) we find evidence for a gene conversion event within the third *TSPY* copy of the array. This *TSPY* copy contains two cluster diagnostic nucleotides ~400 bases apart within the first exon (T-74 and G-478).

We performed a similar set of analyses for *RBMY1* that largely confirmed the presence of four separate regions containing *RBMY1* genes with intact open reading frames (region 1 most proximal, region 4 most distal) across the 44 (42 assemblies, T2T Y and GRCh38) analysed assemblies[10] (**Figures S45**). Though, one sample (NA19239) exhibited an additional 5th *RBMY1* containing region (located between regions 2 and 3), which most likely was the result of a duplication event of two *RBMY1* copies within the distal end of region 2 (**Figures S45**). Additionally, two assemblies (HG02572 and HG03579) contained only 3 *RBMY* regions due to deletion of region 4 and region 3, respectively. Inversions of *RBMY* regions were also found within two assemblies. The first assembly (NA18989) contains (at least) one inversion involving regions 1 and 2. The second assembly (HG02666) carries an inversion of regions 1 and 2 followed by a second inversion of all *RBMY* regions (resulting in a region order of: 4, 3, 1, and 2)(Figure 3a, see Supplementary Results 'Y-chromosomal inversion' below for additional details).

A network-based cluster assignment of *RBMY1* resulted in 11 homogeneous clusters (min cluster size: 3, max cluster size: 101)(**Figure S45**). The vast majority (91.9%) of *RBMY1* copies showed no SNVs, and only 3.11% (11 of 353) of all *RBMY1* copies had more than one SNV, compared to the consensus sequence of their assigned *RBMY1* network cluster. Across assemblies nearly all *RBMY1* copy number variation, as well as lineage-specific events, occurred within regions 1 and 2 (**Figure S45**). In contrast, regions 3 and 4 exhibit little copy number variation harbouring one *RBMY1* copy each (RBMY1F and RBMY1J, respectively; excluding HG03579 and HG02572 which contained the previously described region deletions). To further elucidate the phylogenetic relationships between the regions and *RBMY1* copies, we performed a maximum likelihood analysis (**Methods**). Our phylogeny suggests a radiation of *RBMY1* copies located within regions 1 and 2 from those in regions 3 and 4 (**Figure S47**). Considering these results, in conjunction with the maintenance of the 4 regions across assemblies, we pursued an additional set of analyses (phylogenetic and network-based clustering) for *RBMY1* at the protein sequence level (**Figures S46, S48**). The *RBMY1* protein sequence network-based cluster analysis resulted in fewer unique clusters (9 network clusters, min cluster size: 2, max cluster size: 98) suggesting less variation at the protein sequence level compared to the nucleotide level. RBMY1F (region 3) and RBMY1J (region 4) are identical at the nucleotide and protein sequence level across most haplogroups except the R lineage, which harbours a synonymous SNV **(Figure S45)**. Furthermore, we found that all *RBMY1* copies are functionally intact apart from a single *RBMY1* copy in HG01457. This *RBMY1* pseudogene is located in region 3 and has a 7nt deletion in exon 10 resulting in a premature stop codon and likely nonsense-mediated decay. RBMY1B (Nucleotide Network cluster: 6, Protein Network Cluster: 6) appears unique to the African lineages (**Figures S45, S47**). Intriguingly, the most upstream *RBMY1* copy in GRCh38 is identified as *RBMY1B* and harbours the same amino acid composition (**Figure S46**). Assuming that inclusion of a different haplogroup is not the cause, it may represent a recurrent substitution, or, more likely, be the result of gene conversion or non-allelic homologous recombination.

To elucidate the origin of RBMY1B, we performed comparative analyses involving all *RBMY1* copies at the nucleotide and amino acid level. RBMY1B which harbours one amino acid characteristic for RBMY1D (H126Q) and one from RBMY1F/J (C471Y), either arose through non-allelic homologous recombination between RBMY1D and RBMY1F/J or, alternatively, C471Y represents the ancestral allele followed be a missense mutation resulting in H126Q. The abundance of 471C across *RBMY1* copies in the most ancestral lineage (HG01890; all RBMY1A copies harbour this substitution) favours 471C representing the ancestral state of RBMY1B. To determine the *most* ancestral RBMY1 sequence, we retrieved *RBMY1* exon sequences from the chimpanzee (PanTro6) reference genome. Between the chimpanzee and human *RBMY* sequences we found extensive species-specific variation in the *RBMY1* exonic sequence. However, RBMY1F/J (regions 3 and 4) appears to be ancestral to the other *RBMY1* genes/regions as all chimpanzee *RBMY1* copies share the diagnostic amino acid composition (S-116, H-296) present in RBMY1F/J.

As both the phylogenetic and network-based clustering analyses (nucleotide and protein) suggested the divergence of *RBMY1* copies located in regions 1 and 2 from those in regions 3 and 4, we elected to perform an additional functional analysis of all *RBMY1* copies using InterProScan [18]. InterProScan revealed that all *RBMY1* copies contained the same protein functional domains. However, *RBMY1F/J* copies (regions 3 and 4) contain a missense mutation that codes for an altered amino acid residue, which happens to be one of the ancestral diagnostic residues (S116), within the RNA recognition motif (RRM) domain suggesting similar functional capabilities, but possibly distinct RNA targets. Both RRM domains have been maintained throughout modern human Y-haplogroup radiation. This combined with evidence for multiple, some undiscovered roles, such as the expression in spermatogonia, *RBMY1* protein presence in different regions of sperm (midpiece region and tail)[19], and observation of both oncogenic/anti-oncogenic functions in hepatocellular cancer[20,21], may suggest that not all *RBMY1* copies may not be functionally equivalent.

#### Y-chromosomal inversions

Inversions have remained one of the most challenging structural variation types to reliably genotype, especially when flanked by large highly similar segmental duplications. The Y-chromosomal *de novo* assemblies resolved to base pair level enabled us to confidently identify a total of 16 inversions (14 in the euchromatic regions and 2 in the Yq12 heterochromatic region) from the 44 individuals (43 assembled here and the T2T Y) analysed here, to narrow down the breakpoint locations for 10/16 inversions and improve the inversion rate estimates due to higher phylogenetic resolution compared to previous reports[22,23].

The 14 euchromatic inversions were identified from the *de novo* Verkko assemblies and independently called using Strand-seq data mapped to both GRCh38 Y and the T2T Y reference sequences, available for 31/44 samples (see Methods section **'Inversion analyses'**) (**Tables S32-S33**). In addition, 6/14 of the euchromatic inversions overlapped with inversions called using PAV (**Tables S20, S22**; see Methods section **'Variant calling using *de novo* assemblies'**). All 14 inversions are flanked by inverted repeats (**Figure 3a**), showing up to 99.99% sequence similarity between the repeats and up to 1.45 Mbp in size (the P1 palindrome)[2]. The sizes of inversions range from approximately 30 kbp (the P7 palindrome) to 5.94 Mbp (the IR5/IR5 inversion in HG02666, see more details below) (**Table S32, S35**). Combining the maximum sizes of euchromatic regions affected by inversions sums to a total of approximately 12.18 Mbp or 54.6% of GRCh38 MSY euchromatic composition. 12/14 euchromatic inversions are recurrent, toggling in the Y phylogeny from two (the *blue2/blue3* or *b2/b3* inversion, **Figure 3a, Figure S39**) to 13 times (P3 palindrome composed of 283 kbp inverted repeats which are separated by a 170 kbp spacer region[2]). The two inversions identified in single individuals (the *blue1/blue4* or *b1/b4* in HG01890 and *IR5/IR5* in HG02666, see more details below) are the largest, approximately 4.2 Mbp and 5.94 Mbp in size. Overall, across all 44 samples included in the current study, only the most closely related pair of African Y chromosomes (carried by NA19317 and NA19347) show identical composition in terms of inversions (**Figure 3a; Table S32**), highlighting the high structural variability of the human Y chromosome.

Taking advantage of the sequence resolution offered by the Verkko assemblies, we succeeded in determining the likely breakpoint ranges for 8 euchromatic inversions down to 500-bp region (**Figure 3b; Figures S36-S38; Table S35;** Methods section **'Determination of inversion breakpoint ranges'**), allowing us to determine the inversion sizes more accurately. According to the GRCh38 Y coordinates, the average sizes of breakpoint ranges for palindromes P3-P8 are 33,381 bp (ranging from 1,115 bp in P3 to 181,342 bp in P4) and 33,203 bp (ranging from 1,117 bp in P3 to 181,342 bp in P4) for proximal and distal copies of inverted repeats or palindrome arms. The location of breakpoint ranges tend to be located closer to the spacer region, suggesting that the distance between the breakpoints in the proximal and distal arms impacts the triggering of an inversion event (**Figure S37**). The inversion sizes (for palindromes P3-P8 and IR3) range from 29,426 bp (palindrome P7) to 3,679,407 bp (IR3) with an average of 714,036 bp, when estimating it based on the start coordinate on the proximal repeat/arm and the end coordinate on distal repeat/arm of the breakpoint ranges. The inversion size, as well as the size of the breakpoint range, are positively correlated with the size of palindrome, except for the IR3 repeats where the unique spacer region (~3.5Mb) is substantially larger than that of any Y palindrome (Spearman's correlation coefficient between breakpoint range and proximal/distal arm size: 0.8857 (p-value 0.0333), and between inversion size and proximal/distal arm size: 1.00 (p-value 0.0028) based on GRCh38 coordinates).

Large inversions are mostly responsible for the fact that all three of our contiguously assembled Y chromosomes are structurally distinct from each other across multi-Mbp euchromatic regions (**Figure 2b-c; Figures S8-S10, S18, S35**), and from both GRCh38 and the T2T Y sequences, which also differ from each other due to a known >1.9 Mbp polymorphic *gr/rg* inversion carried by the T2T Y[11,24]. The structural composition of the *AZFc* region in the deepest-rooting Y chromosome (HG01890 A0b-L1038) can be explained by two inversions (between the *blue 1* and *blue 4* amplicons, and another between the *blue 2* and *blue 3* amplicons, **Figure S39**), up to 4.1 and 1.2 Mbp in size (considering the start and end coordinates of the respective blue amplicons in the GRCh38 Y), respectively, or three inversions (additionally requires the *gr/rg* inversion) when compared to the T2T Y (**Figures 2b-c; Figures S8, S18 and S35a**).

The second deepest-rooting Y chromosome from HG02666 (A1a-M31) carries a P5/P1 inversion and additionally a smaller inversion between *blue 2* and *blue 3* amplicons (**Figures 2b-c; Figures S9, S18**). We were able to pinpoint the inversion breakpoints of the P5/P1 inversion into 504-bp intervals (**Figure S38**) within ERV1 repeat elements in the IR5 repeats located in inverted orientations in the distal arm of P5 palindrome and in the proximal arm of the P1 palindrome. The resulting inversion is 5.941 Mbp in size relative to the Verkko assembly for HG02666, or 6.001 Mbp relative to GRCh38 Y and likely caused by non-allelic homologous recombination (NAHR). Recombination between palindromes P5 and P1 (both P5/proximal-P1 and P5/distal-P1 deletions, known as *AZFb* deletions) are known to cause massive deletions and spermatogenic failure, with most breakpoints identified within a hotspot region within 30 kbp from the center of the P5 palindrome [25]. Interestingly, the inversion breakpoints identified here do not overlap with the deletion hotspots as they are located ~81.7 kb from the center of the P5 palindrome. Closer inspection of the sequences of the *blue 2* and *blue 3* repeats from HG01890 and HG02666 indicates that these are independent inversions and were therefore counted as independent events in inversion rate calculations.

The Y assembly for HG00358 (N1c-Z1940) contains a known ~1.8 Mbp *b2/b3* deletion fixed in haplogroup N samples (**Figures 2a-b; Figures S10, S18 and S35a**)[11].

We detected the *gr/rg* inversion, one of the major structural differences between the GRCh38 Y and the T2T Y sequences, in seven samples (**Figure 3a; Table S32**), including the two other haplogroup J samples (HG02492 J2a-M47 and HG01259 J1-M267) which are most closely related to the T2T Y. The presence of *gr/rg* inversions is also supported by Bionano optical mapping data. Our results on *gr/rg* phylogenetic distribution fit well with previous reports both in terms of the presence of this inversion in haplogroups B2b-M112, E1b1b1b1a-M81, and its absence in other Y lineages overlapping between the two studies, although matching the results exactly is not possible due to lower resolution of typed phylogenetically informative markers by Repping and colleagues[11]. This most likely also explains the absence of the *gr/rg* inversion in their haplogroup J samples, while indicating that the inversion is not shared by haplogroup J samples as our phylogeny might suggest, but instead occurred independently in J1-M267 and its sublineages (carried by HG01259 and the T2T Y) and J2a-M47 (carried by HG02492). However, since we were not able to determine the inversion breakpoints for the *gr/rg* inversion, we took the conservative approach and counted a total of 5 independent inversions in the phylogeny (instead of 6 in case the inversions in J1 and J2a were independent). Overall, the concordance with previous studies supports structurally correct assembly of this complex region in our dataset.

The largest recurrent inversion among our samples is found on the p-arm, mediated by the inverted IR3 repeats, each approximately 290-300 kbp in size. The IR3 inversions are known to be polymorphic and reported to be approximately 3.3-3.8 Mbp in size[22,23]. Interestingly, we discovered that most (33/44) Y chromosomes, including the T2T Y, show a distinct composition of IR3 repeats compared to the GRCh38 Y sequence (**Figure S49**). In GRCh38 Y, the distal IR3 repeat contains a single copy of the ~20.3 kbp *TSPY* repeat (including the *TSPY2* gene; see Method section **'TSPY repeat copy number analysis'**) in direct orientation, while in the majority of samples the single TSPY repeat is located in the proximal *IR3* repeat in inverted orientation (**Figure 3b; Figure S49**). Analysis of the *IR3* repeat sequences revealed that the phylogenetically closely related Y haplogroup QR samples (including GRCh38 Y, mostly haplogroup R1b) have likely undergone two inversions - a ~3.67 - 3.68 Mbp (relative to GRCh38 Y sequence) inversion changing the location and orientation of the single TSPY repeat from the distal to proximal repeat, while another, ~3.24 - 3.28 Mbp inversion reverted the region located between the *IR3* repeats (**Figure S49; Table S35**). In addition to these two events shared by all QR lineage Y chromosomes, the *IR3/IR3* inversion was identified in four samples which now carry the genomic region in between the *IR3* repeats in inverted orientation compared to other samples **(Figure 3a**), totalling to six inversion events across all analysed samples. The inversion breakpoint ranges were narrowed down to regions of 6.7 to 40.1 kbp in size (**Figure 3b; Table S35**). In two samples (NA19239 and HG03492) the inversion breakpoints were located closer to the unique spacer region, leading to inversions of ~3.2 Mbp in size. Interestingly, the inversion breakpoint region in HG03492 overlaps with the second inversion region shared by all QR samples. In HG03732 and NA19331 the inversions were larger, ~3.4 Mbp in size, and inversion breakpoints were located closer to the center of IR3 repeats.

Additionally, we highlight an inverted duplication which affects roughly two thirds of the 161 kbp unique sequence in the P3 palindrome, spawns a second copy of the *TTTY5* gene and effectively elongates the segmental duplications in this region (**Figure S35b**). A detailed sequence view reveals a high sequence similarity between the duplication and its template, and its placement in Y phylogeny supports emergence of this variant in the common ancestor of haplogroup E1a2 carried by NA19239, HG03248 and HG02572 (**Figure 1a; Figures S1, S35b**).

In addition to the inversions in the euchromatic regions of the Y chromosome, we also identified inversions at the proximal and distal ends of the Yq12 heterochromatic region, one at each end (**Figure 4c**). The inversion breakpoint analyses at the nucleotide level revealed distinct breakpoints, further supporting the presence of these two inversion events (**Figure S40; Table S34**). Alternatively, a complex rearrangement with multiple breakpoints, resulting in orientation changes of the *DYZ1* and *DYZ2* repeat units within the distal and proximal ends of the Yq12 region, could have occurred.

As some variation was noticed within the proximal inversion region across the 11 analysed samples, breakpoint analysis was performed for each assembly separately. For 9/11 examined assemblies, the 5' breakpoint of the proximal inversion was identified within a *DYZ2* repeat unit at the 3' end of the *Alu* sequence immediately upstream of a second 'orphaned' *Alu* A-tail (Adenosine-rich sequence) segment (**Figure S40a**). The 3' breakpoint of the proximal inversion resides within the intersection of an AT-rich simple repeat region of a *DYZ2* subunit and a *DYZ1* subunit (**Figure S40a**). For all of the assemblies analysed, the 5' breakpoint of the distal inversion is situated at the boundary of an AT-rich simple repeat and the 5' end of an *Alu* sequence ('head' of the *Alu*) within a *DYZ2* repeat unit (**Figure S40b**). Finally, the 3' breakpoint of the distal inversion lies between an AT-rich simple repeat and the remaining portion of the *Alu* sequence head right before the HSATI satellite (**Figure S40b**).

Across the eleven analysed samples, three distinct patterns within the proximal inversion region were observed. While the majority of assemblies shared the breakpoints described above, two assemblies – HG01106, and HG01890 – showed a deviating pattern. In HG01106 the entire proximal inversion region seems deleted and additional studies are required to determine if this is shared by other closely related Y chromosomes, or is sample-specific (rearrangements having occurred in the lymphoblastoid cell line can not be excluded). To determine the ancestral state of the inversion region, the HG01890 Y assembly was further investigated. This was deemed particularly important, as HG01890 represents the deepest rooting Y chromosome lineage in the current dataset. Comparison of HG01890 with the other Y assemblies revealed the likely presence of deletions encompassing both the 5' and 3' breakpoints of the proximal inversion.

#### Yq12 heterochromatic subregion

##### A Yq12 overview

Our comparison of the Yq12 subregion of T2T Y and GRCh38 Y revealed that the distal section, situated closest to the PAR2 subregion, is structurally distinct from the rest of Yq12 and fully assembled in GRCh38 Y reference sequence. As no evidence of structural variation was found within this region, we focused on the previously incompletely assembled proximal sections of this region, including the *DYZ18* repeat array (**Figure 1a**; **Tables S13, S15-S16**) in our subsequent analyses of the seven samples (HG01890, HG02666, HG00358, HG01106, HG01952, HG02011 plus the T2T Y) with contiguously assembled Yq12 heterochromatic regions. The largest completely assembled Yq12 subregion is the 7th largest Yq12 subregion observed among the 44 samples analysed (**Fig. S17b**). Therefore, the assembly outcome is likely determined not only by the size of the region.

First, we assessed the previously mostly unassembled Yq12 region for its repetitive sequence composition. Within each of the analysed genomes, we observed an alternating pattern of two distinct segments (**Methods**). One segment consists mainly of a tandemly repeated AT-rich simple repeat fused to a 5' truncated *Alu* element, followed by an HSATI satellite. Comparison with the Yq12 literature revealed that this arrangement represents a previously described ~2.4 kbp tripartite repeat element, *DYZ2*[26,27]. The subunit composition in the second segment was less well defined. We noticed that these sequences mainly contain simple repeats and pentameric satellite sequences, with over 95% (33,677 of 35,370) of all satellites identified as HSATII. Further analyses revealed an association of this sequence with a ~3.5 kbp repeat called *DYZ1*[2,28–31]. The *DYZ1* repeat unit showed more variation in size (range from 1,165 to 3,608 bp, with 95% of all *DYZ1* repeat units longer than 3,000 bp with a mean length of 3,543 bp) compared to the *DYZ2* repeat units (range from 1,275 to 3,719 bp, with 93.7% of all *DYZ2* repeats 2,420 bp in size) (**Methods**). Consequently, our analyses support that the alternating repeat segments be identified as *DYZ1* and *DYZ2* arrays. Interestingly, the total number of arrays within assemblies is positively correlated to the length of the analysed Yq12 region (two-sided Spearman: 0.90; p-value=0.0056, **Figure S71, Methods**).

Next, we extended the *DYZ1* and *DYZ2* array analyses to the two assemblies (HG01928 and NA19705) with a single gap within the Yq12. Additionally, we included the assemblies of the two most closely related individuals (NA19317 and NA19347) with an estimated divergence time of ~200 years despite the presence of multiple contigs to gain a better understanding of the evolution of this region. For the two assemblies with multiple contigs, we focused our analyses on the arrays that are continuously assembled and reside at the proximal and distal ends of the Yq12 region. As expected, we identified copy number variation both with regard to the number of *DYZ1* and *DYZ2* arrays and *DYZ1* and *DYZ2* repeat units within the arrays in all four assemblies **(Figure S72b)**. However, the number of *DYZ1* and *DYZ2* repeat arrays within the assembled regions was identical within the two most closely related genomes **(Figures S72b, S83)**. Furthermore, the *DYZ2* repeat unit copy numbers within 14/20 *DYZ2* arrays between NA19317 and NA19347 were identical **(Figure S83)**. Comparison of these 14 *DYZ2* arrays with identical repeat unit copy number (encompassing a total of 2,231,881 nucleotides) revealed only five single nucleotide variants (SNVs) – none of which represented CpG mutations – and one indel within a homopolymeric adenosine tract. Of the remaining six *DYZ2* arrays, four were located in the proximal or distal ends of the Yq12 region and showed only minor variation in the *DYZ2* repeat unit copy number (+/- 1 *DYZ2* repeat units). The last two arrays were not included in the analyses because of their immediate adjacency to an incomplete assembly region.

##### Yq12 *DYZ1* and *DYZ2* repeat analyses

We examined inter-individual variation with regard to subunit composition of Yq12 *DYZ2* arrays in greater detail. Across the seven assemblies with fully assembled Yq12 region, the total *DYZ2* repeat units within the Yq12 region ranged from a minimum of 2,661 *DYZ2* subunits (HG01890) to a maximum of 6,681 *DYZ2* subunits (HG01106), with a mean of 4,380 units. *DYZ2* repeat units ranged in size from a minimum of 1,275 bp to a maximum of 3,719 bp, though 98.6% (30,242 out of 30,656 of all *DYZ2* repeat units across complete assemblies) were between 2,000-2,999 bp in length, with a median length of 2,420 bp (93.7% of all *DYZ2* repeats were 2,420 bp). Sequence composition analysis suggests that this variation in sequence length is primarily caused either by expansion or contraction within the AT-rich simple repeat segment of these elements (sample collective mean: 1,415 bp, standard deviation (SD): 383 bp). The single origin *DYZ2 Alu* sequence had a consistent length (sample collective mean: 290 bp, SD: 2 bp) and was primarily identified as *Alu*Y, though at roughly 20% divergence, the sequence is too diverged to confidently exclude *Alu*S origin. The HSATI satellite portion of the *DYZ2* subunit varied somewhat in size (sample collective mean: 566 bp, SD: 16 bp).

Our comparison to the *DYZ2* consensus sequence revealed that *DYZ2* repeat units located within arrays and positioned closer to the center of the Yq12 region were, on average, less diverged (i.e., potentially younger) **(Figure 4d; Figure S73)**. In contrast, more divergent *DYZ2* repeats were enriched toward the proximal and distal boundaries of the Yq12 region, with the putative oldest elements detected within the arrays situated between the distal inversion and the 3' end of the *DYZ* repeat arrays. Interestingly, this divergence pattern also seemed to be partially reflected within the individual *DYZ2* arrays where the divergence of *DYZ2* repeats situated closer to the center was generally lower compared to those near the ends. To investigate ongoing mutation dynamics, we also performed the *DYZ2* divergence analysis for the two most closely related genomes (NA19317 and NA19347). As expected, based on the previous *DYZ2* array comparisons, high similarity was uncovered between both genomes, and a similar divergence pattern as observed within the other genomes **(Figure S73b)**.

To further investigate the evolution of the Yq12 heterochromatin, we performed a phylogenetic analysis of the *DYZ2* repeat (**Figure S75).** As the *DYZ2* repeat arrays outside the inversion (the most centromeric and telomeric arrays) showed evidence for a different sequence composition, we built a *DYZ2* consensus sequence of this region from all seven fully assembled Yq12 samples (including T2T Y, **Table S15**) using a majority rule approach. In addition, we constructed a consensus sequence utilizing the central *DYZ2* copies (between the two fixed inversions). As Babcock et al. 2007[32] identified that *DYZ2* sequence is present on all other acrocentric chromosomes (chr 13, 14, 15, 21 and 22), we also constructed a *DYZ2* consensus sequence for each acrocentric chromosome and reconstructed a phylogeny of *DYZ2* using a maximum likelihood approach. Our analysis supports that the internal elements are younger than the copies toward both ends of the Yq12 heterochromatin region. Furthermore, we can infer that the most distal and proximal copies are more similar to the *DYZ2* copies on the other acrocentric chromosomes.

Next, we constructed a *DYZ2* repeat composition profile for each *DYZ2* array within a genome. Our inter-*DYZ2* array profile comparison (see **Methods**), performed for each genome separately, revealed a trend towards *DYZ2* arrays closely situated to one another having higher repeat composition similarity **(Figures 4e; Figure S74)**. Curiously, these *DYZ2* array composition similarity heatmaps (**Figure S74**) also exhibit what appear to be signals of past waves of amplifications/duplications of *DYZ2* arrays located between the peripheral Yq12 inversions.

Next, we investigated the Yq12 *DYZ1* repeat units in greater detail. Due to the low sequence complexity of the pentameric HSATII satellite and the simple repeat, we were unable to utilize the same approaches as those performed for the *DYZ2* arrays. Furthermore, an analysis using the previously published *DYZ1* consensus sequence[2] as a query sequence revealed an overall high divergence (~25%), further confounding downstream analyses. Based on these findings, two different approaches were pursued: (1) a virtual restriction digestion of the *DYZ1* array sequences with HaeIII that cuts DNA at ggcc sites[33], and (2) a targeted HMMER analysis[34]. The HaeIII restriction enzyme was selected based on previous molecular biology experiments of the *DYZ1* repeats in the Yq12 subregion, where the enzyme was shown to cut the repeat unit once, primarily resulting in fragments with 3,564 bp in length[33]. While our virtual digestion of the putative Yq12 *DYZ1* array regions of all complete assemblies showed a similar enrichment for 3,564 bp size fragments, we also observed considerable sequence length variation (Min: <25 bp, Max: >200 kbp) **(Figure S84)**. Visualization of the distribution of fragment lengths within *DYZ1* arrays revealed a highly similar pattern across the seven complete Yq12 assemblies **(Figure S84)**.

To explore the repeat composition of restriction fragments, we performed a k-mer profile similarity analysis. Considering that the first *DYZ1* array is adjoining the Yq11 *DYZ18*, 3.1-kbp, and 2.7-kbp repeat transition region, each digestion fragment was classified as being a unit, or a composition, of either *DYZ18*, 3.1-kbp repeat, 2.7-kbp repeat, or *DYZ1*. Compellingly, the findings of the *DYZ18* and transition region analysis within the Yq11 were supported and reiterated by this analysis **(Figures S63, S66)**. The k-mer profile dissimilarity analysis indicated that the 3.1-kbp repeat showed higher similarity to the *DYZ18* repeat (91%), and the 2.7-kbp repeat to *DYZ1* (85%), suggesting that the Yq11/Yq12 transition zone repeats (3.1-kbp and 2.7-kbp) are possibly derived from *DYZ18* and *DYZ1* **(Figure S66)**. Lastly, the virtual digest and HMMER analyses were combined where after digestion fragment classification, a targeted HMMER analysis was performed to partition restriction fragments into their individual repeat subunits **(Figure S63)**.

While previous studies reported a ratio of *DYZ1* to *DYZ2* repeat units as 2 to 1[27,35,36], we observed a nearly equal repeat unit ratio (collective sample mean *DYZ1*:*DYZ2* ratio: 1.09) within the Yq12 **(Figure 4b; Table S47)**. These findings align with our observation of a nearly 60:40 ratio of total nucleotides accounted for by *DYZ1* and *DYZ2* across all analysed assemblies. Finally, the dissimilarity of *DYZ1* repeats versus the constructed *DYZ1* consensus sequence was computed and visualized (see **Methods**). This analysis mirrored findings of the *DYZ2* repeat divergence analysis, with *DYZ1* subunits located near the center of *DYZ1* arrays tending to be less dissimilar (i.e., less diverged) than those found near the boundaries of arrays **(Figure S65)**.

#### Yq12 mobile element insertions (MEIs)

The Yq12 region was screened for the presence of mobile element insertions (MEIs) generated by the target-primed reverse transcription mechanism in both the *DYZ1* and *DYZ2* arrays. Four putative *Alu* insertions were identified across the seven samples with full Yq12 assemblies (**Figure 4f**). While three of the insertions resided within the *DYZ2* repeat unit, the fourth insertion was located within the *DYZ1* repeat unit. Based on the divergence (3% or less), all four putative insertions appeared considerably younger than the *Alu* sequence of the composite *DYZ2* repeat unit. Furthermore, all *Alu* elements harboured hallmarks of classical MEIs such as target site duplications, termination in an adenosine-rich tail, and endonuclease cleavage site **(Table S28)**. Two of the insertions were identified as *Alu*Y and one each as *Alu*Ye5, and *Alu*Yb8. Both *Alu*Y insertions occurred within the AT-rich simple repeat region of the *DYZ2* repeat; though at different locations and not within the same repeat unit. The *Alu*Yb8 element inserted into a *DYZ1* repeat; while the *Alu*Ye5 element inserted immediately upstream of the 5' *Alu* sequence of one *DYZ2* repeat and in 'sense orientation' relative to *DYZ2*.

*Alu* elements are unique in that the ancestral state (i.e., absence of the MEI) is known and the precise removal of a MEI is exceedingly rare[37]. Based on this, the approximate age of the insertions, and presence in all Y chromosome lineages, it can be inferred that the two *Alu*Y insertions have occurred early in human Y chromosome evolution prior to the rise of the now known Y chromosome lineages. Only the T2T Y assembly lacked evidence for one of the two *Alu*Y insertions. Based on its phylogenetic placement, this likely results from a deletion or gene conversion of repeat units harbouring the insertion **(Figure 4f)**. The *Alu*Ye5 insertion is unique to HG01890, and the *Alu*Yb8 element to HG01952. Further analysis revealed that the *Alu*Yb8 element is shared with HG01928 (assembly of the Yq12 subregion is not contiguous), supporting insertion in a common ancestor of HG01952 and HG01928 **(Figure 4f; Table S28)**.

While there is little evidence for post-insertion expansion of the *Alu*Yb8 element in the *DYZ1* repeat, the MEIs within a *DYZ2* repeat show varying degree of expansion with considerable inter-individual variation (**Figure 4f**). For example, one *Alu*Y insertion was identified in six out of seven assemblies with a copy number range from one (in HG01106) to seventeen (in HG02666). This further highlights the enormous inter-individual variation of the human Yq12 region. Furthermore, from the MEI patterns it can be inferred that the insertions occurred into different repeat arrays and that the expansion/duplication occurred independently for each MEI. Interestingly, each MEI insertion and their extensions occupy distinct areas within the Yq12 region with no overlap between the different MEIs **(Figure 4f)**.

These findings, in conjunction with the overall *DYZ1* and *DYZ2* array expansion/contraction dynamics, point toward random unequal crossing over between sister chromatids for the subsequent expansions of the *Alu* elements as well as the duplication or deletion of *DYZ1* and *DYZ2* arrays[38]. Unequal crossing over would also explain the expansion and contraction of repeats within these arrays without changing the repeat pattern[38], though gene conversion and replication slippage as contributing factors cannot be ruled out. The lower interindividual variation with regard to array number, array size, and *DYZ1/2* repeat units of the inversion regions and arrays distal to the inversions at the proximal and distal ends of the characterized repeat region is in agreement with the known recombination and crossing-over suppression of inversions[39]. Furthermore, a reduction in unequal crossing over near/within the Yq12 inversions could protect against deleterious effects outside the heterochromatin region such as gene-containing regions of the Y chromosome.

#### Functional analysis

DNA methylation calls on the ONT reads were derived from Nanopolish [40], after methylation calling and QC (**Methods**) we used pycoMeth[41] to *de novo* segment the methylation profiles of the 41 QC passing samples (**Figure S55**). This resulted in the identification of 2,861 independent segments (**Table S45**). We first assessed the link between DNAme levels and chromosome Y assembly length, identifying that there is a borderline significant negative link between DNAme levels and assembly length both genome wide **(Figure S54a)** and on chromosome Y specifically **(Figure S54b)**, P=0.0469 within chromosome Y and P=0.0477 autosome-wide. Though this effect is only borderline significant it might suggests that also in humans there is a link between repressive chromatin modification and chromosome Y length as previously reported found in Drosophila before [42]. Next to assess the global impact of the different haplogroups on the segmentation we used a permanova test. Specifically, we grouped haplogroups into 6 meta groups based on sample size and genetic distance, haplogroup A, B and C (“ABC” 4 samples), G and H (“GH” 2 samples), N and O (“NO” 6 samples), and Q and R ("QR" 11 samples), E (19 samples), J (4 samples - including NA24149, the father of HG002/NA24385), **Methods**). These grouped haplogroups explain 21% of the global variation in DNAme levels profiles (Permanova, *P* 0.0029). Here we also included chromosome Y assembly length, and find that on a segment level it explains 3% of the variation in DNAme levels on the chromosome Y segments (*P* 0.068). This indicates that the observed link between chromosome Y assembly length and DNAme levels is also influenced by haplogroup, and needs further validation to follow this up amongst others to validate this in a non cell-line setting.

Next to these global analyses we found on a segment level that 340 segments are differentially methylated (DM) (FDR 20%, **Table S45**, (**Methods**)) depending on haplogroup. Interestingly 218 (64%) of the segments have decreased DNAme levels in the QR haplogroups. The 340 DM segments are enriched to overlap regulatory information (Fisher exact *P < 2.2e-16,* odds ratio: 6.72*),* but depleted in overlap to genes (Fisher exact *P 2.088e-05,* odds ratio: 0.52*,* methods, **Table S45**).

Next to the effects of haplogroups on DNAme we tested for local DNA methylation quantitative trait loci (meQTLs). We leveraged the limixQTL pipeline to test for effects of genetic variation with 100,000 bases around the DNAme segment as identified using pycoMeth. We controlled for population structure by controlling for population as a random effect, and leveraged permutations to determine significance of effects (supplementary methods). We identified 10 segments with significant meQTLs (FDR 20%) and found a total of 194 meQTL effects. The majority of the effects are linked to SNVs (109), with 1 variant being an INV, and 1 effect being from a 171 base-pair insertion (**Table S46**).

Given that expression data is available only on a subset of the HGSVC and HPRC samples (21/44) we focussed on the 210 males from the Geuvadis project[43] to assess the effects of haplogroups on gene expression level. We find 64 of the 205 genes on chromosome Y expressed in the Geuvadis LCL gene expression data (**Table S56**). As with DNAme we first tested for global expression variation, here we leveraged the first character of the haplogroup as grouping ("E":44, "G":4, "I":23, "J":18,"N":22,"R":96, "T":3 (group:nSamples)), and find that Y haplogroup explains 4.8% of the variation in gene expression (Permanova, *P 0.005*), and in total 22 genes are significantly differentially expressed (FDR 10%). Even though the samples and Y haplotype distribution is different between the DNAme samples and the Geuvadis data we find 5 genes (*BCFORP1*, *LINC00280*, *LOC100996911*, *PRKY*, *UTY*) that have both DNAme effects as well as gene expression effects. Specifically, *BCORP1* is interesting as the effect directions on average match between the Geuvadis and HGSVC expression datasets and the expression effect is negatively correlated (r -0.3; p:0.1) between the overlapping HGSVC samples (**Figure S56**).

To demonstrate the utility of these highly contiguous Y assemblies in representing the genic diversity of other individuals, we analysed full-length cDNA sequences (PacBio Iso-Seq) of testis samples from seven anonymous donors (**Methods**). Of 30 Y-chromosomal genes expressed with at least five cDNA reads, 23 had improved transcript alignments compared to the T2T Y reference sequence, which provided only equal or inferior alignments (**Figure S85; Table S57**). Most notably, *DAZ2* transcripts had alignments improved by 15.5% on average, due to the variable internal repeat structure. Across all genes, a full 19% of the improved alignments came from the Y assembly of a single sample, HG01596. We also generated Iso-seq data on eight matched samples corresponding to *de novo* Y assemblies (**Figure S86; Table S58**). Aligning to a matched *de novo* Y assembly instead of the T2T Y reference improved between 14-51% of cDNA alignments.

Hi-C data has been widely utilized to characterize the 3D structure of the genome and identify chromatin structures, such as topologically associated domains (TADs) that play central roles in gene regulation. Previous research has primarily focused on Hi-C data analysis in autosomes, while here we investigate the variation of chromatin structures in diverse Y chromosomes. Using Hi-C data available from 40 samples, we identified TADs and TAD boundaries for Y chromosomes of these individuals by evaluating their insulation scores, which indicate the variations of the contact density of every Hi-C bin compared to adjacent bins (**Figures S87-S88**; **Methods**)[44]. Regions with high insulation scores are more likely to be found inside TADs and regions among TADs intend to have low insulation scores. In total, 112 TAD boundaries at 10 kbp resolution were detected in our merged callset of 40 samples (**Table S59**). We illustrated the average and variance (maximum difference between any of the two samples) of insulation scores of each sample to indicate the changes of chromatin structures together with the corresponding methylation profiles and chrY assembly (**Figure S55b**). For the 340 DMRs which are detected in the aforementioned methylation analysis, we performed Kruskal-Wallis H tests (FDR 20%) with the same 6 meta haplogroups on the insulation scores (10 kbp resolution) in each DMR to detect regions that are differentially methylated as well as differentially insulated. Among the 26 DMRs that intersected with 21 differentially insulated regions (DIRs), we found one of such region (DMR: chrY-7289920-7290751, DIR: chrY-7290001-7300000) that harbours the PRKY gene which is both differentially DNA methylated and differentially expressed (**Table S60**).

### Supplementary Figures

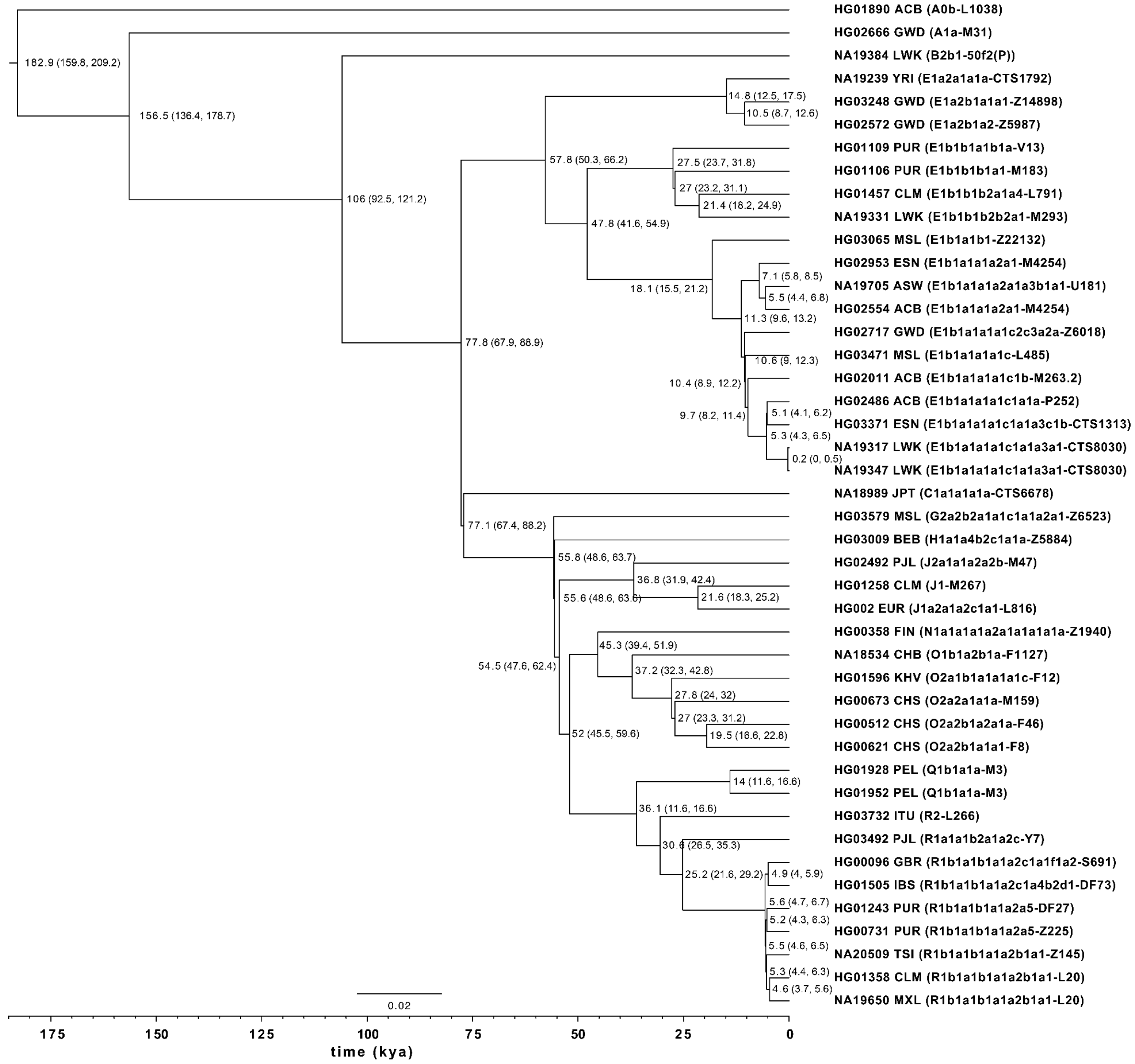


**Figure S1.** Phylogenetic relationships of the analysed Y chromosomes. Split times as estimated according to the BEAST analysis are shown with 95% HPD interval in brackets (kya - thousand years ago). Sample ID is followed by population designation, full Y haplogroup label according to ISOGG v15.73 and terminal marker ID. Population abbreviations: ACB - African Caribbean in Barbados; ASW - African Ancestry in SW USA; BEB - Bengali in Bangladesh; CHB - Han Chinese in Beijing, China; CHS - Han Chinese South; CLM - Colombian in Medellín, Colombia; ESN - Esan in Nigeria; FIN - Finnish in Finland; GBR - British From England and Scotland; GWD - Gambian in Western Division – Mandinka; IBS - Iberian Populations in Spain; ITU - Indian Telugu in the U.K.; JPT - Japanese in Tokyo, Japan; KHV - Kinh in Ho Chi Minh City, Vietnam; LWK - Luhya in Webuye, Kenya; MSL - Mende in Sierra Leone; MXL - Mexican Ancestry in Los Angeles CA USA; PEL - Peruvian in Lima Peru; PJL - Punjabi in Lahore, Pakistan; PUR - Puerto Rican in Puerto Rico; TSI - Toscani in Italia; YRI - Yoruba in Ibadan, Nigeria.

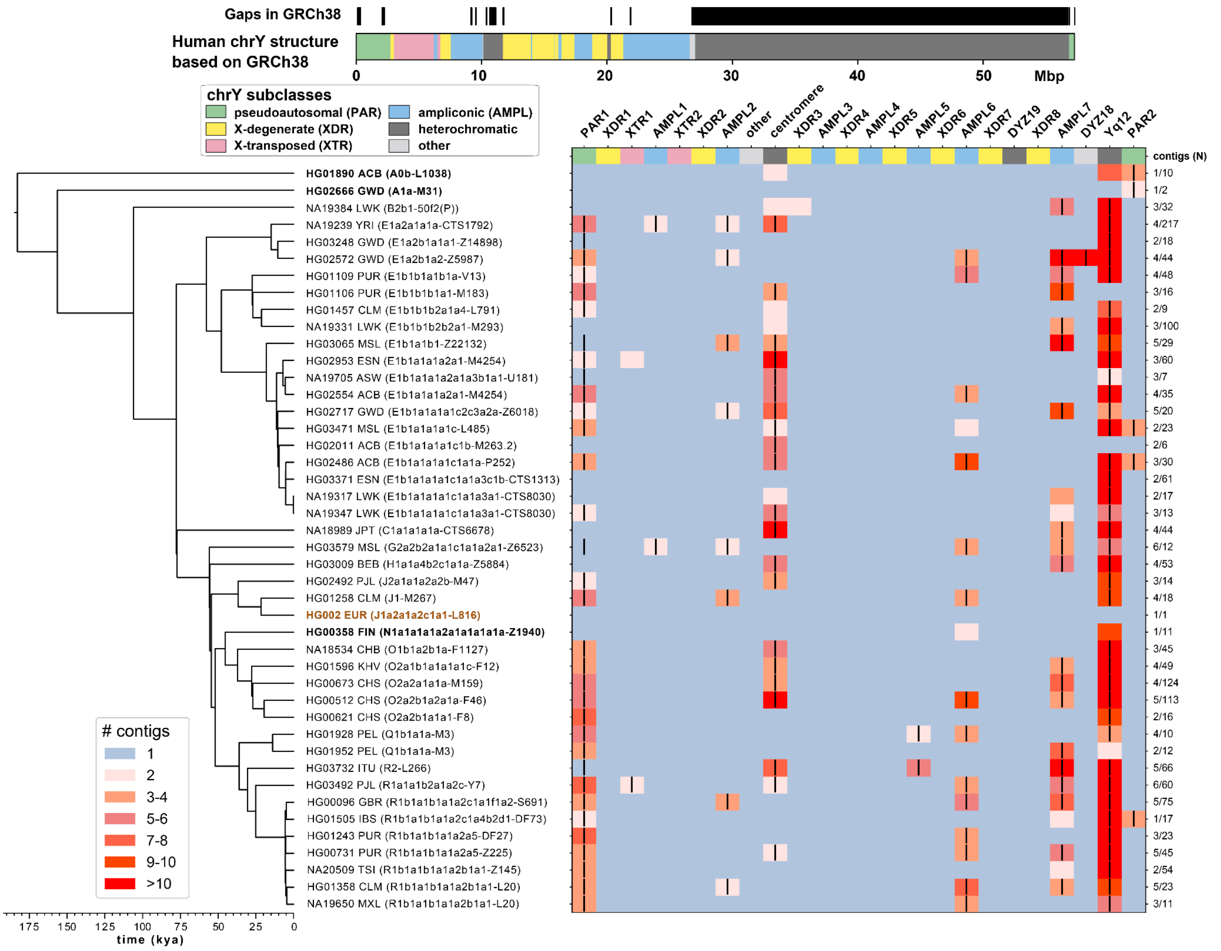


**Figure S2.** Phylogenetic relationships of the analysed Y chromosomes and assembly completeness. Phylogenetic relationships of the analysed Y chromosomes with branch lengths drawn proportional to the estimated times between successive splits according to BEAST analysis. Summary of Y assembly completeness with the number of contigs containing sequence from specific sequence class indicated with different colours (on the right - number of Y contigs needed to achieve the plotted assembly contiguity/total number of assembled Y contigs for each sample). Sample IDs include the population abbreviation, and the full Y lineage and terminal marker in brackets. See **Figure S1** for population abbreviations.

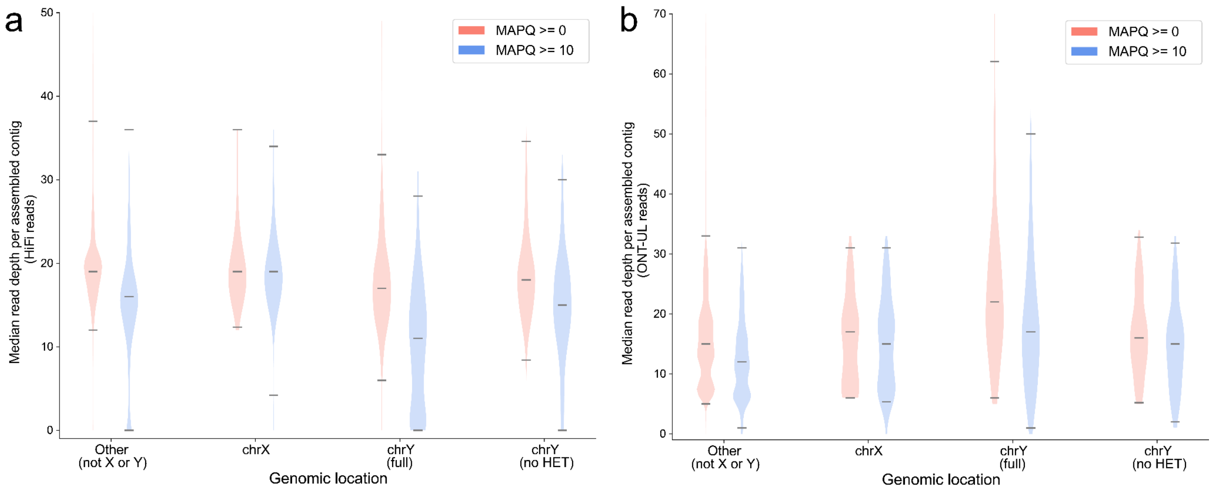


**Figure S3.** Median read depth across assembled contigs at two different MAPQ thresholds. **a.** PacBio Hifi reads, and **b.** ONT-UL reads. Genomic regions are depicted as follows from left to right: autosomes, chromosome X, chromosome Y and chromosome Y excluding heterochromatic regions (i.e., peri/centromeric region, *DYZ19* and Yq12 heterochromatic regions).

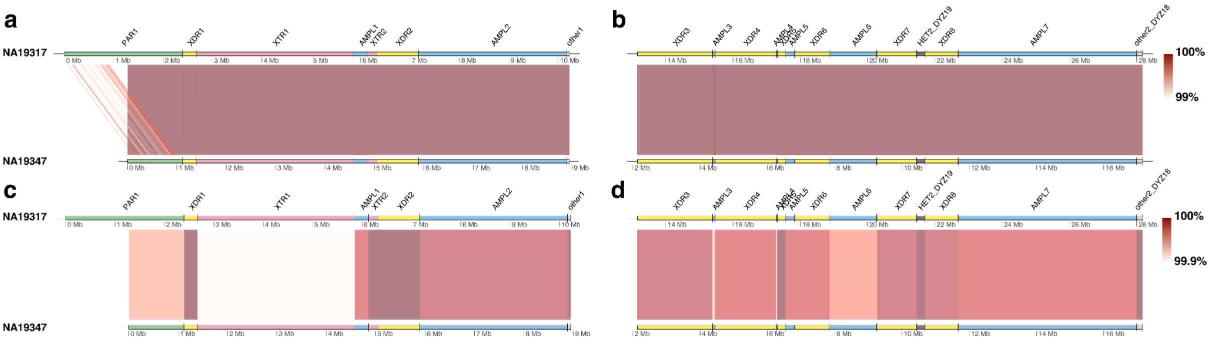


**Figure S4.** Comparison of the Y assemblies from closely related African Y chromosomes (NA19317 vs NA19347). Comparison of contiguously assembled regions spanning: **a, c.** from PAR1 until the end of other1, and **b, d.** from XDR3 to the end of *DYZ18* using two sequence identity thresholds (upper ≥99%, lower ≥99.9%). Pairwise sequence alignments of 21/24 contiguously assembled Y-chromosomal subregions showed sequence identity ranging from 99.982% to 100% (**Table S8**), with 100% sequence identify in three subregions (other1, *DYZ19* and *DYZ18*). Eight subregions (XDR1, XTR2, XDR2, XDR3, AMPL3, AMPL4, XDR6, and XDR8) have no substitutions and the number of indels range from 2 to 26. XTR1 subregion shows the lowest sequence identity (99.982%) with 185 mismatches and 389 indels/gaps in the alignment.

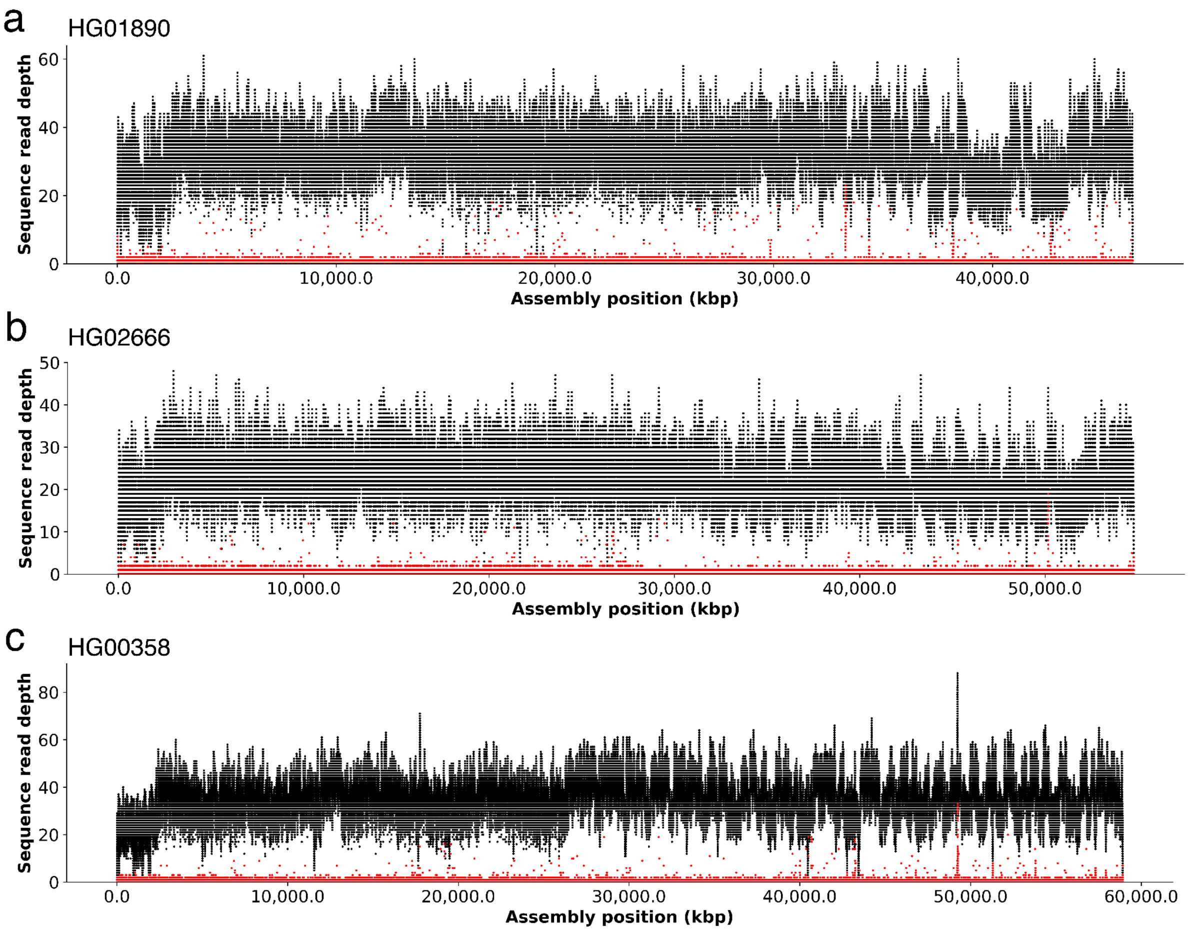


**Figure S5.** NucFreq HiFi read depth and nucleotide frequency plots for three samples. Black bars show HiFi read depth and red scatter depicts the abundance of the second most common base in the read-to-assembly alignment.

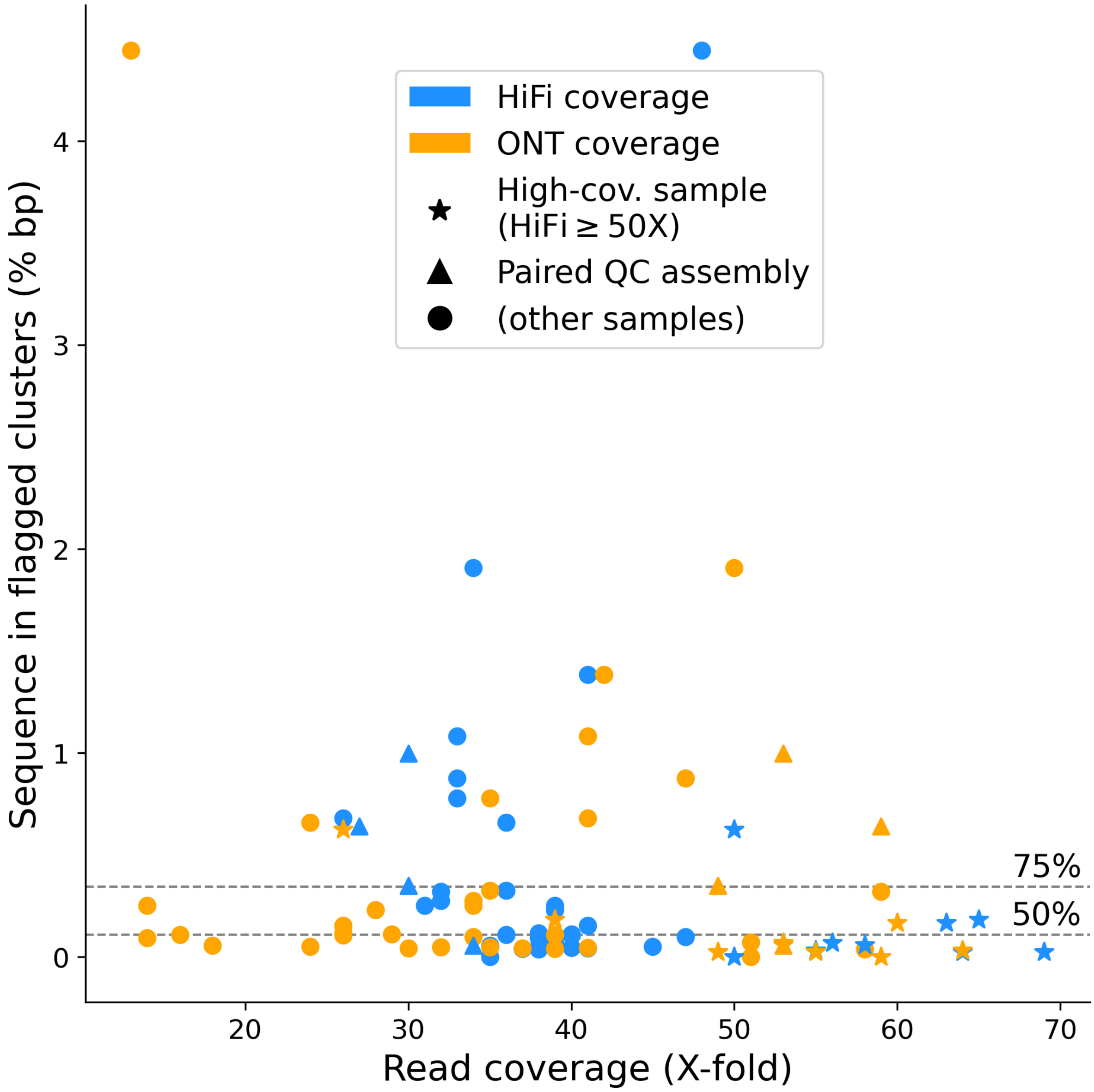


**Figure S6.** Scatter plot of input read coverage for both ONT (orange) and HiFi (blue) per sample (X-fold coverage relative to a ~3.1 Gbp genome size, x-axis) and putative assembly errors (% clustered flagged sequence, y-axis). "Star" markers highlight high-coverage samples. "Triangle" markers indicate assemblies created for QC purposes using approximately half of the HiFi coverage of the respective high-coverage sample. Dashed horizontal lines indicate the second and third quartile of samples.

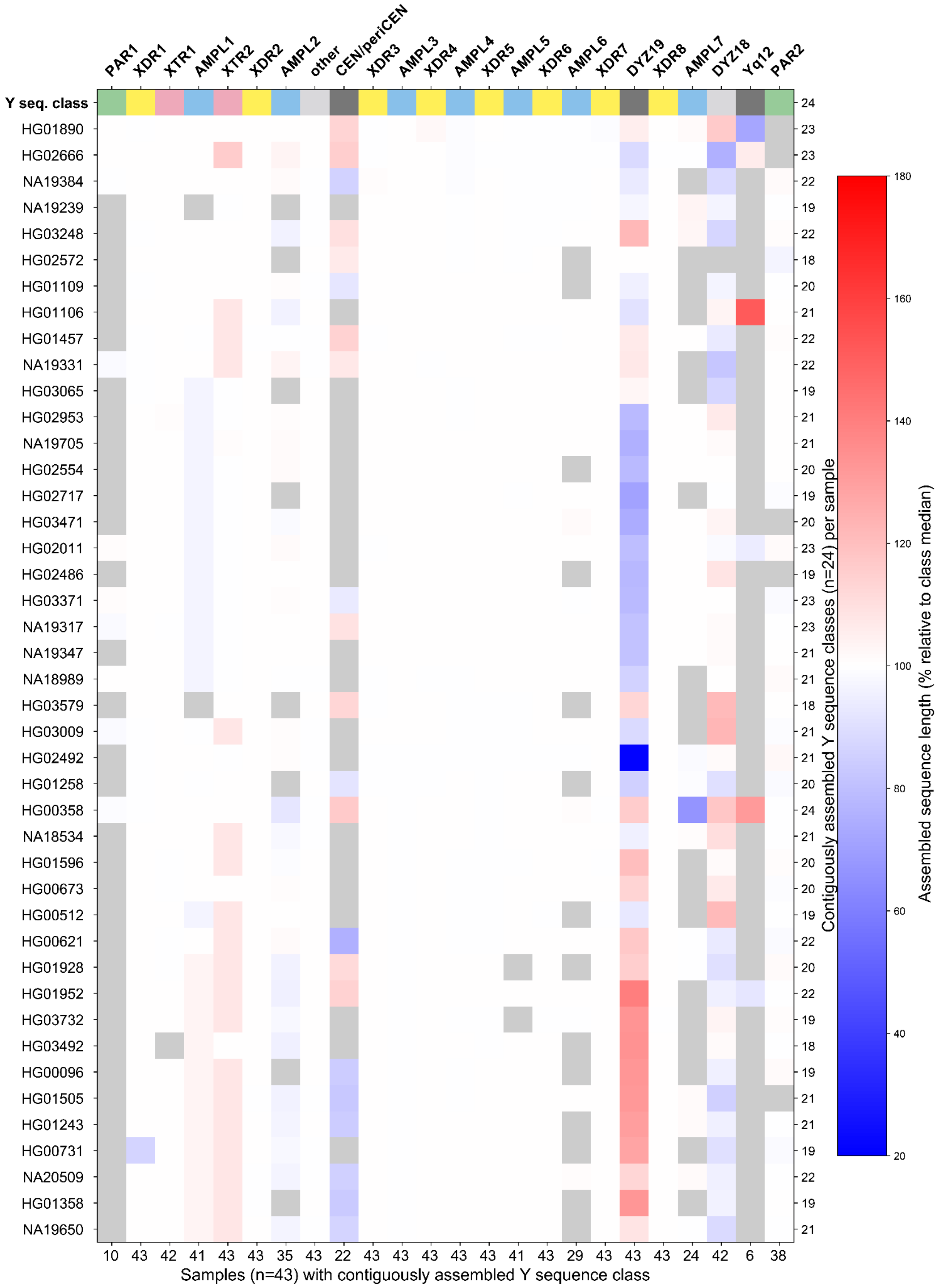


**Figure S7.** Size variation of contiguously assembled Y-chromosomal subregions shown as a heatmap relative to the median size of contiguously assembled subregions (shown as 100%). Boxes in grey indicate regions not contiguously assembled (Methods). Numbers on the bottom indicate contiguously assembled samples for each subregion out of a total of 43 samples, and numbers on the right indicate the contiguously assembled Y subregions out of 24 regions for each sample.

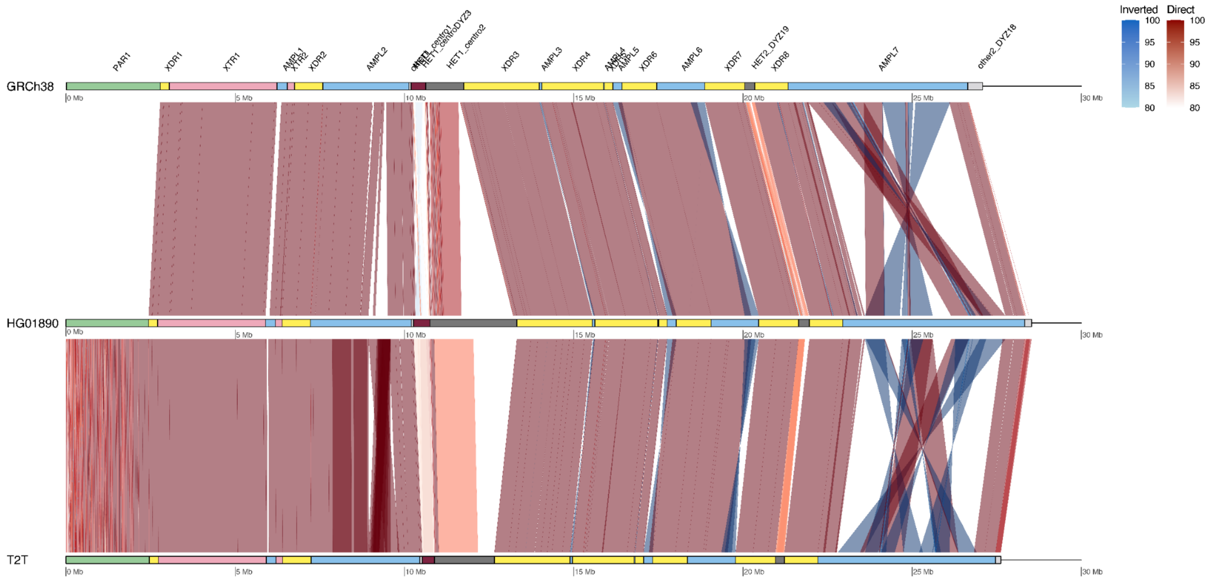


**Figure S8.** Comparison between GRCh38, HG01890 and T2T Y.

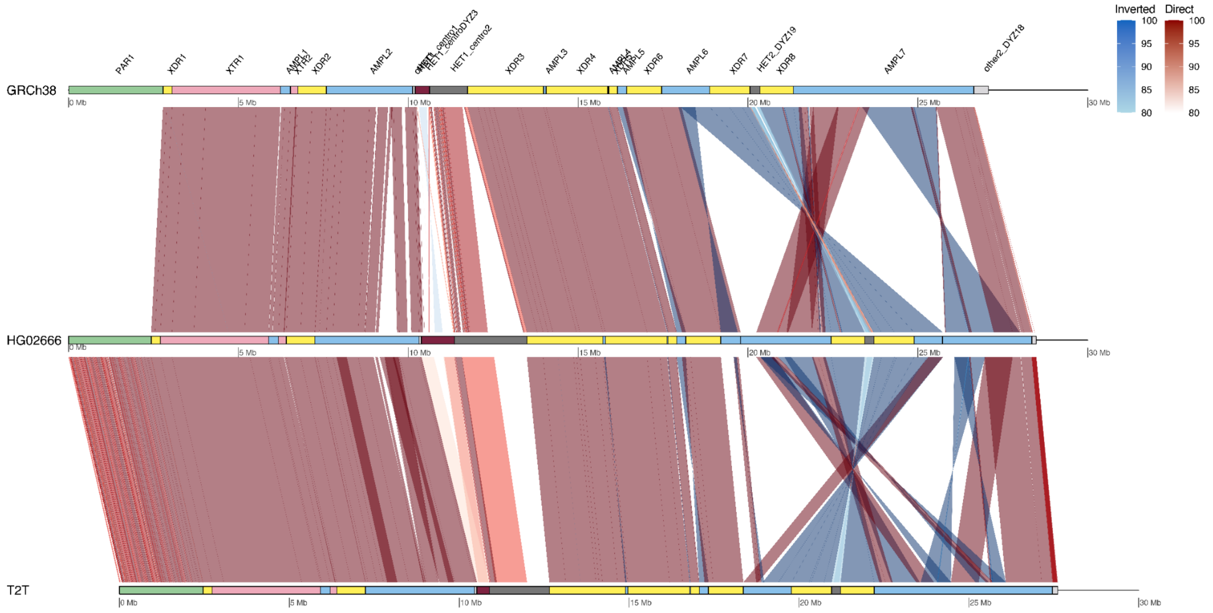


**Figure S9.** Comparison between GRCh38, HG02666 and T2T-T.

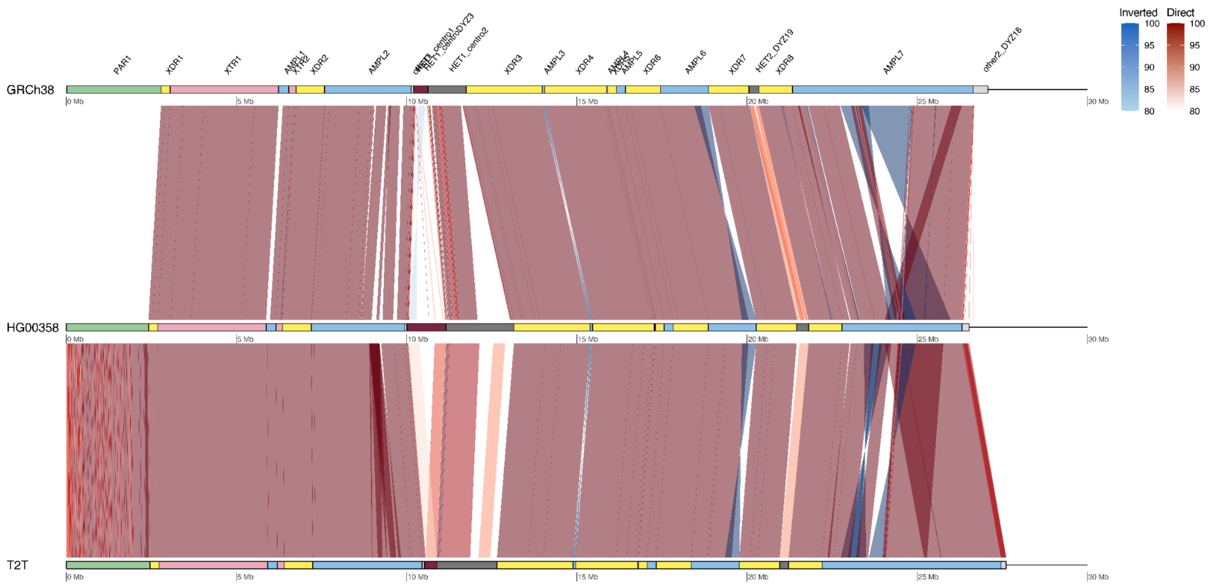


**Figure S10.** Comparison between GRCh38, HG00358 and T2T Y.

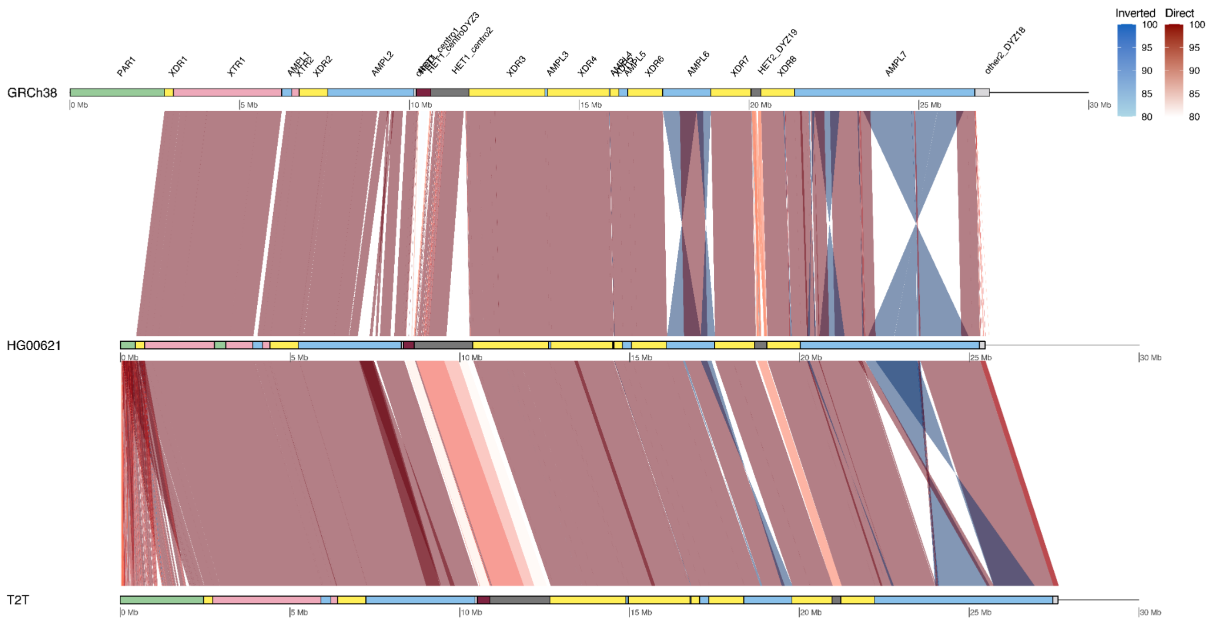


**Figure S11.** Comparison between GRCh38, HG00621 and T2T Y.

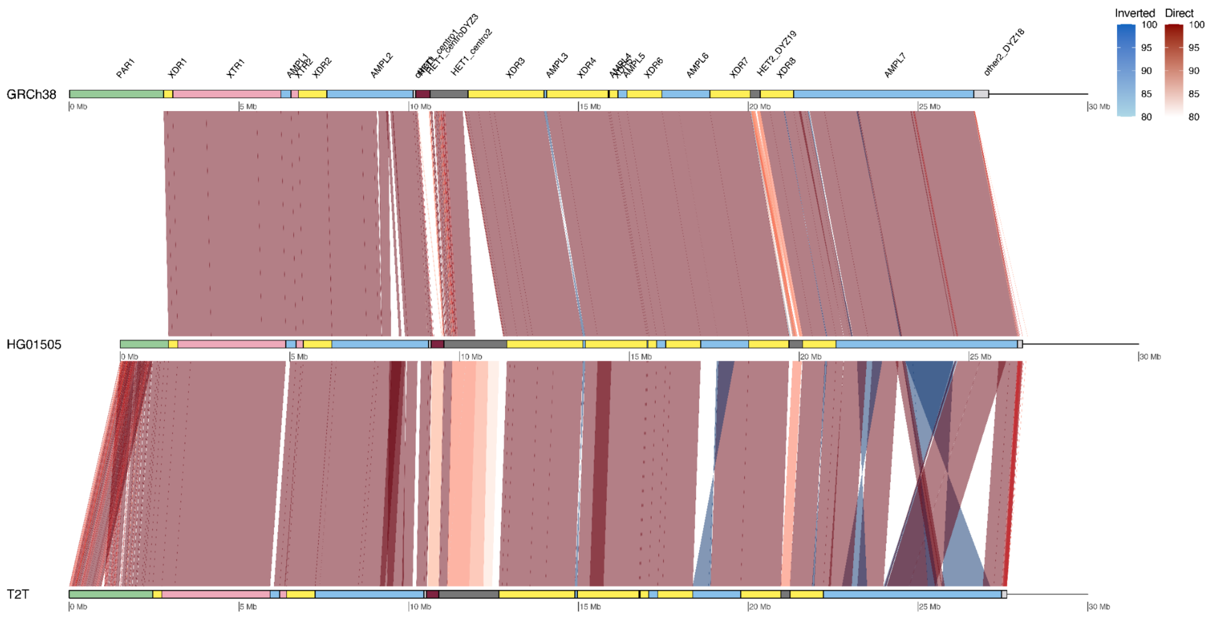


**Figure S12.** Comparison between GRCh38, HG01505 and T2T Y. Note - GRCh38 and HG01505 are phylogenetically closely related, both representing haplogroup R1b. Highly similar assembly and lack of large difference between GRCh38 and HG01505 supports accuracy of our de novo assemblies.

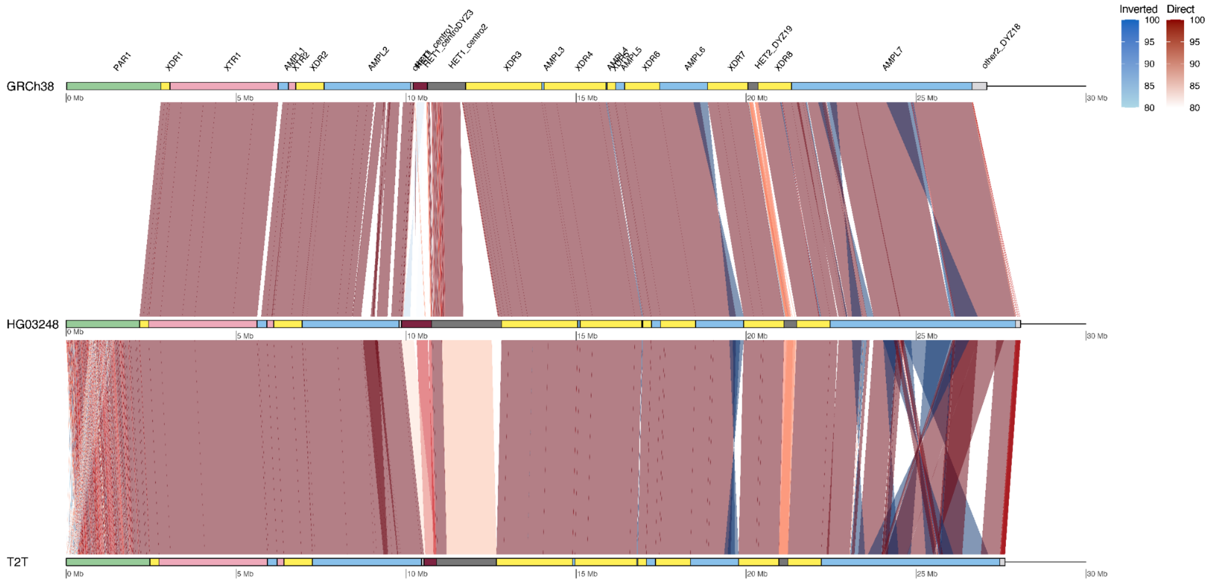


**Figure S13.** Comparison between GRCh38, HG03248 and T2T Y.

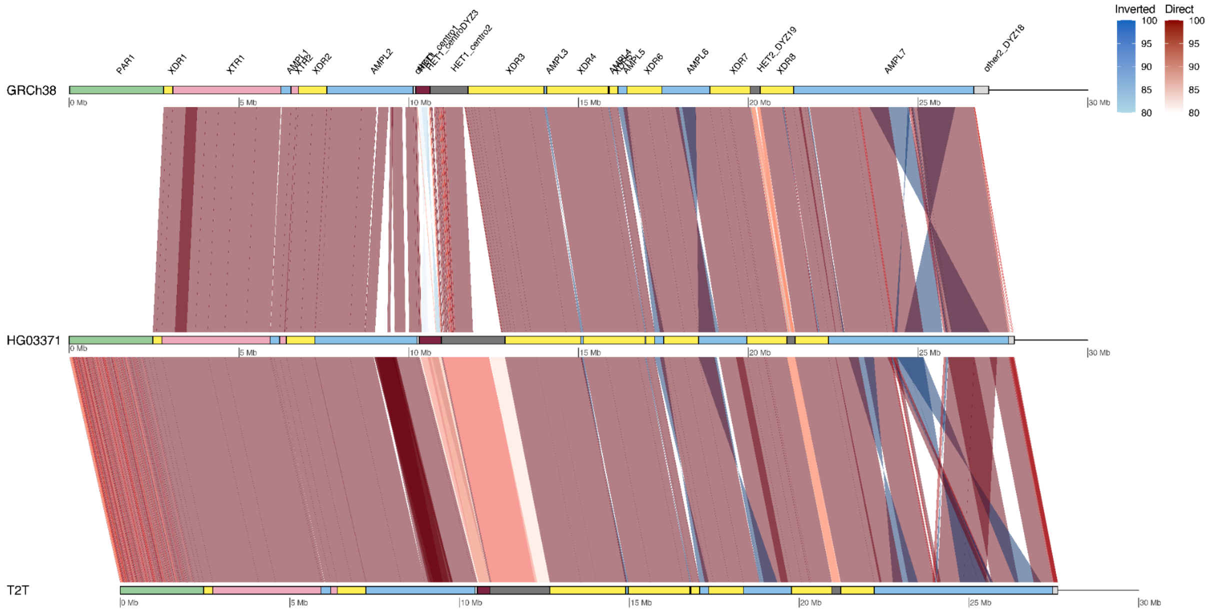


**Figure S14.** Comparison between GRCh38, HG03371 and T2T Y.

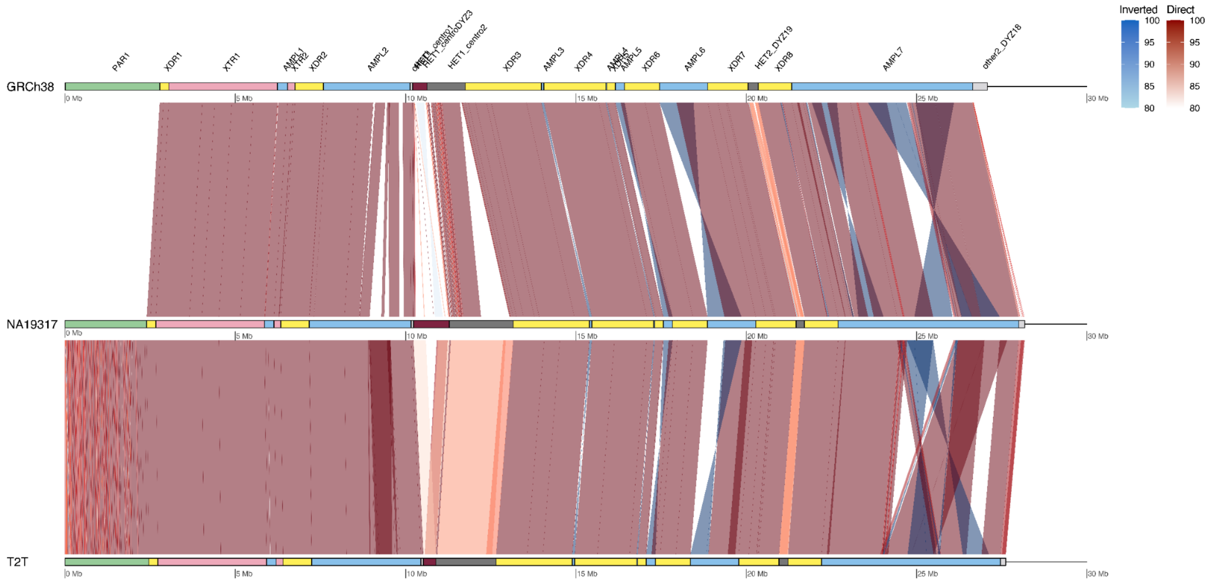


**Figure S15.** Comparison between GRCh38, NA19317 and T2T Y.

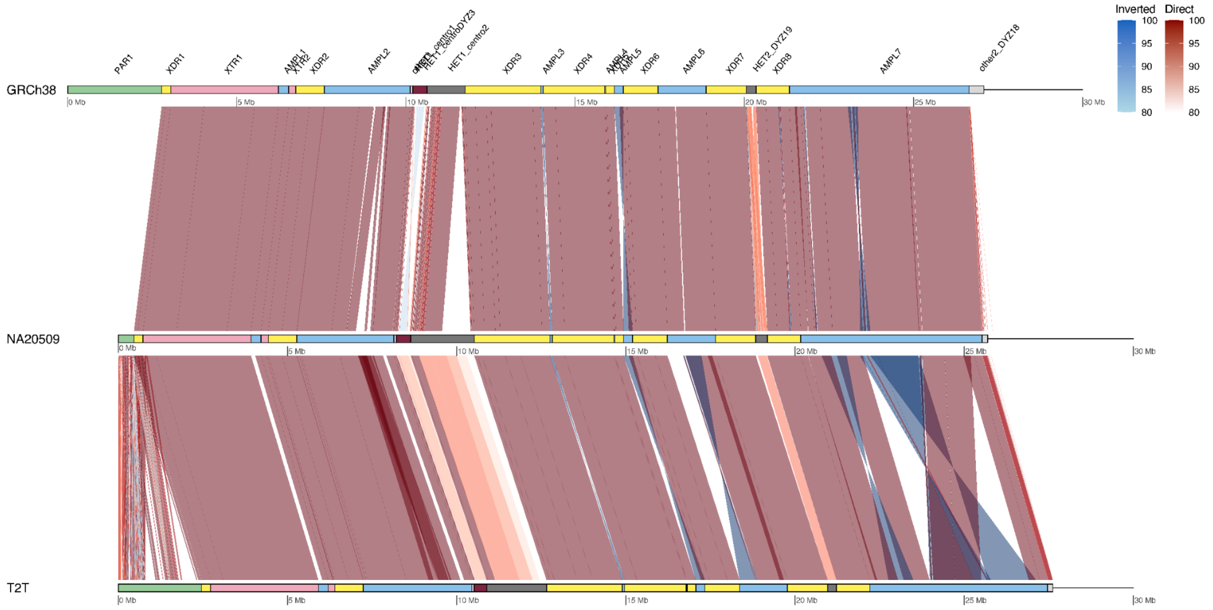


**Figure S16.** Comparison between GRCh38, NA20509 and T2T Y. Note - GRCh38 and NA20509 are phylogenetically closely related, both representing haplogroup R1b. Highly similar assembly and lack of large difference between GRCh38 and NA20509 supports accuracy of our de novo assemblies.

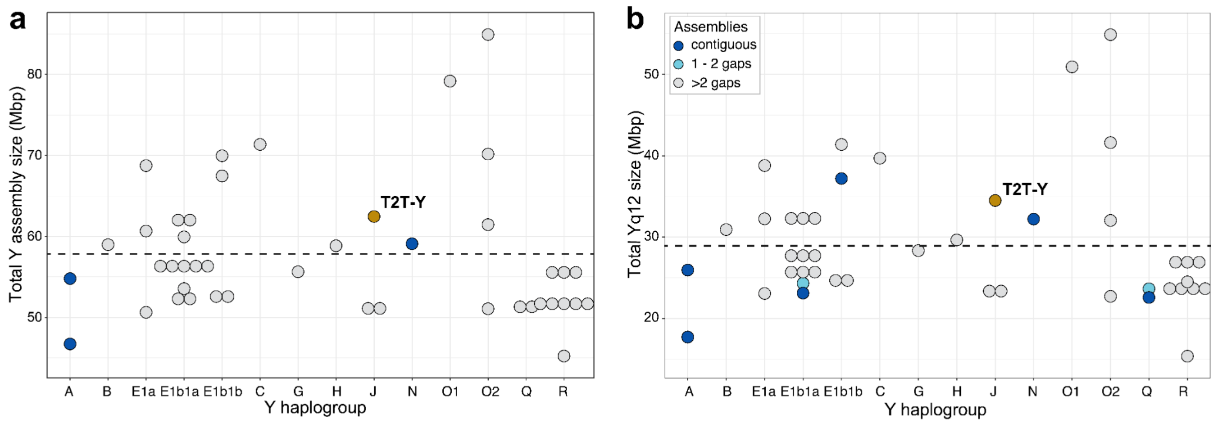


**Figure S17**. Y assembly sizes across Y haplogroups. **a**. The total combined Y assembly size. **b**. The total combined Yq12 subregion size. Samples with contiguous assembly, with 1-2 or more gaps and the T2T Y are indicated with different colours. Black dashed line indicated the mean (57.6 Mbp for total Y assembly and 29.0 Mbp for the Yq12 subregion).

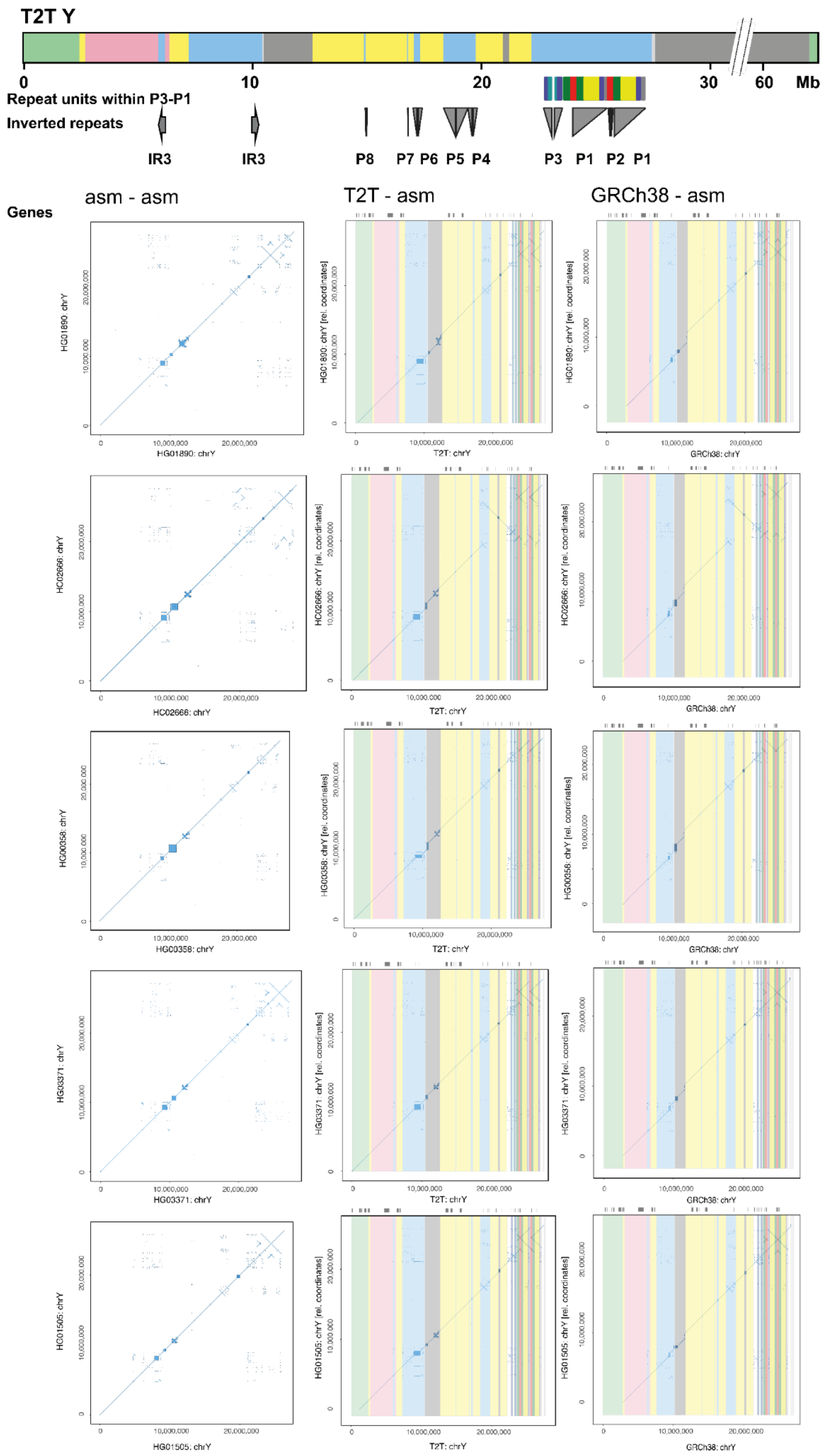


**Figure S18**. Dotplots of five samples contiguously assembled across the euchromatic regions (from PAR1 until Yq12 heterochromatic region) with self dotplot on the left, compared to T2T Y in center and to GRCh38 on the right, annotated with sequence classes and SD repeat units in ampliconic 7 region.

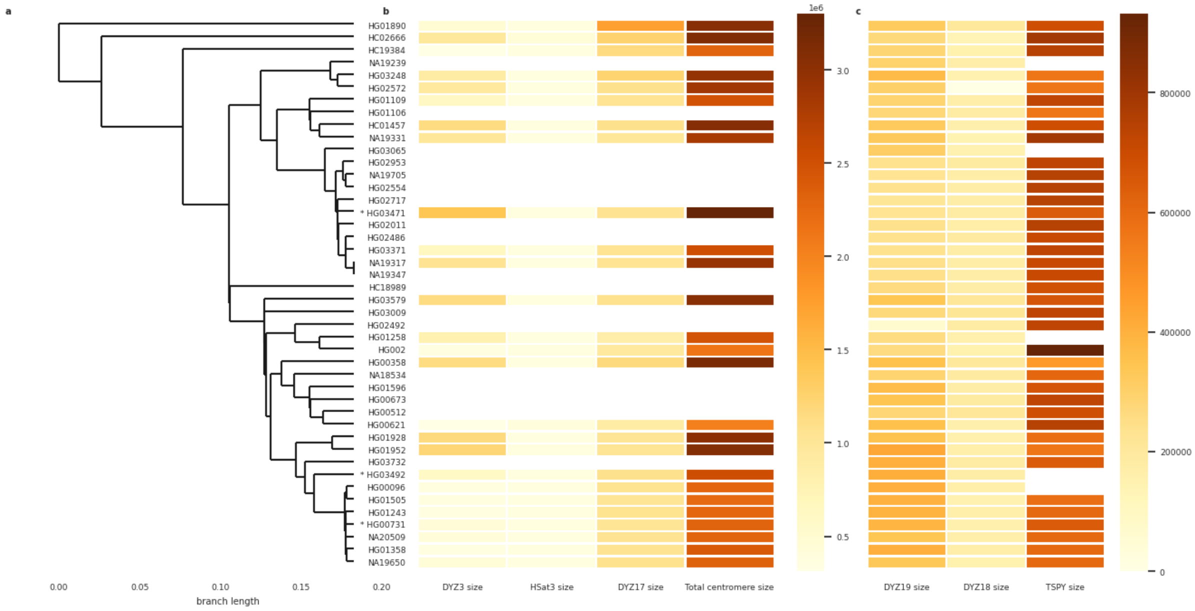


**Figure S19.** Size variation of the (peri-)centromeric region and repeat arrays (*DYZ3* alpha-satellite array, *Hsat3*, *DYZ17* array, and total (peri-)centromeric region) on the left and the *DYZ19*, *DYZ18*, and the TSPY copy-number variable repeat arrays on the right, with sizes shown as a heatmap. **a.** Phylogenetic clustering of the samples, as described in **Fig. S1**. **b.** Size variation heatmap for each pericentromeric region, and the total centromere size in millions of base pairs. White fill indicates that the size information of the region is not available due to non-contiguous assembly of the region. Asterisk to the left of the sample name indicates samples (HG00731, HG03471, and HG03492) with one assembly gap in the (peri-)centromeric region. **c.** Size variation heatmap for *DYZ19*, *DYZ18* and TSPY repeat arrays. The sizes of the (peri-)centromeric regions (*DYZ3* alpha-satellite array, *Hsat3*, and *DYZ17* array) were regressed against each other, but none achieved significant correlations.

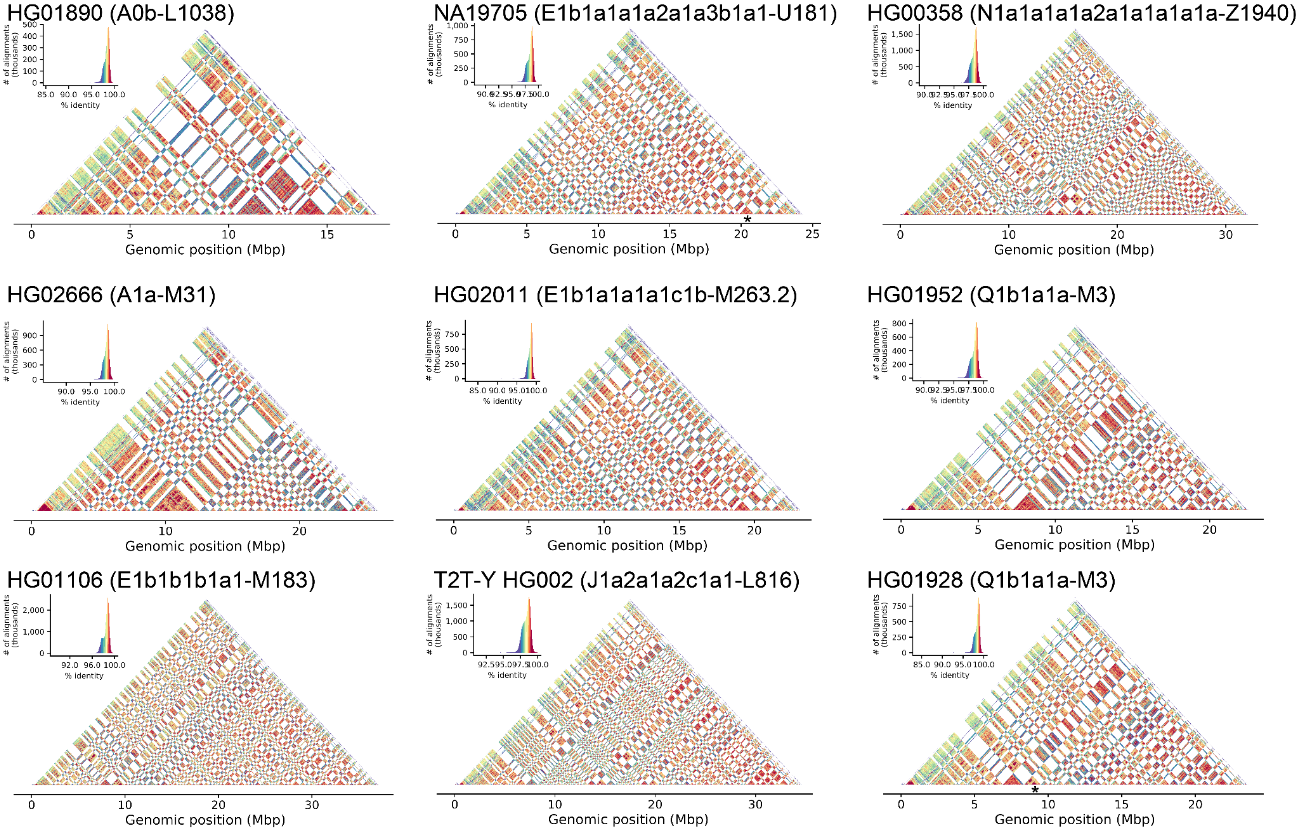


**Figure S20.** Sequence identity heatmaps of the Yq12 subregion for six contiguously assembled samples (HG01890, HG02666, HG01106, HG02011, HG00358 and HG01952), two samples (NA19705 and HG01928) with a single gap in the Yq12 subregion (gap location marked with asterisk) and the T2T Y from HG002 using 5kb window size.

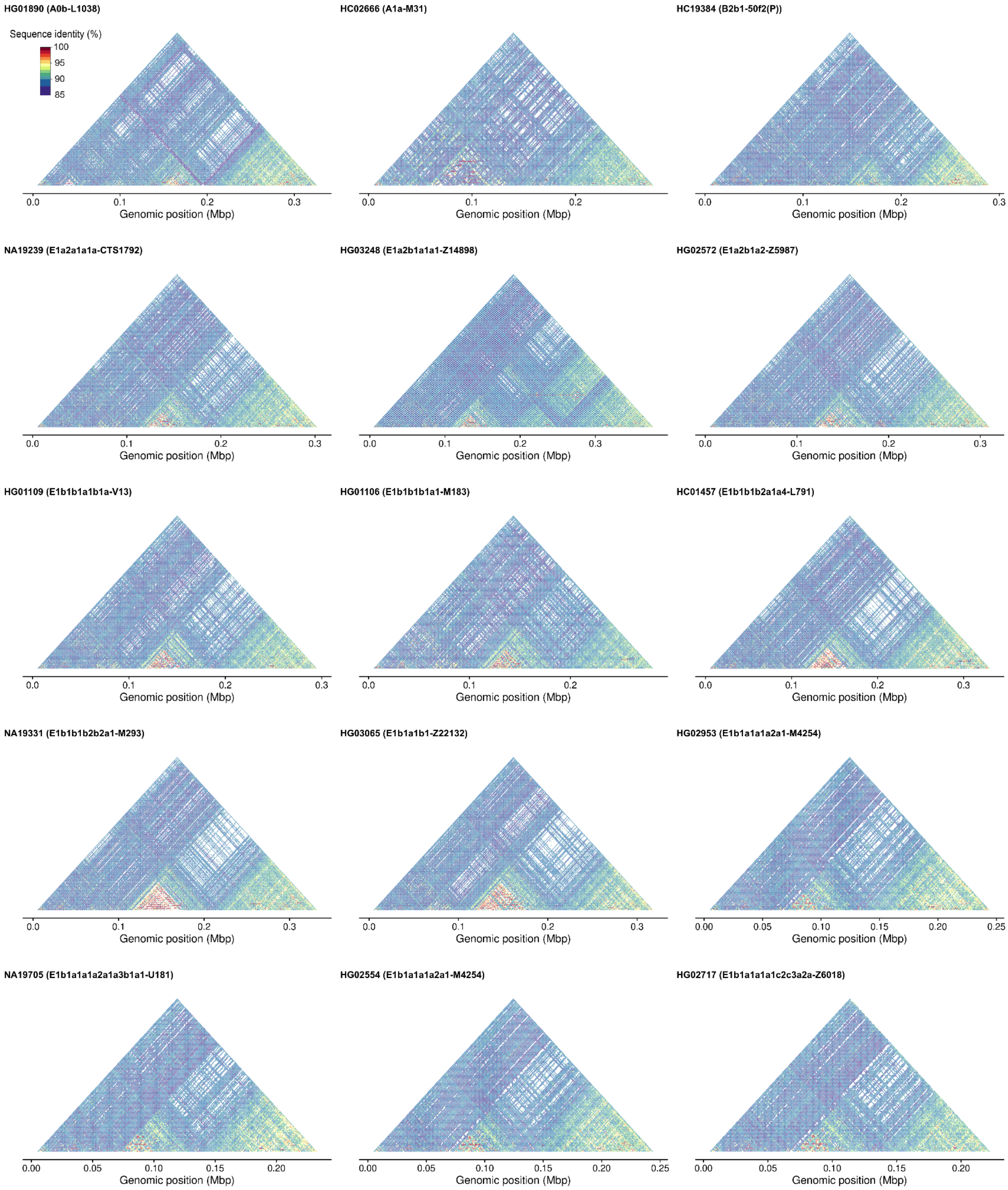


**Figure S21.** Sequence identity heatmaps of the DYZ19 subregion across samples, including the T2T Y, in phylogenetic order. 5000 bp of flanking sequence was added to the DYZ19 genomic interval and 1 kbp window size was used when running StainedGlass.

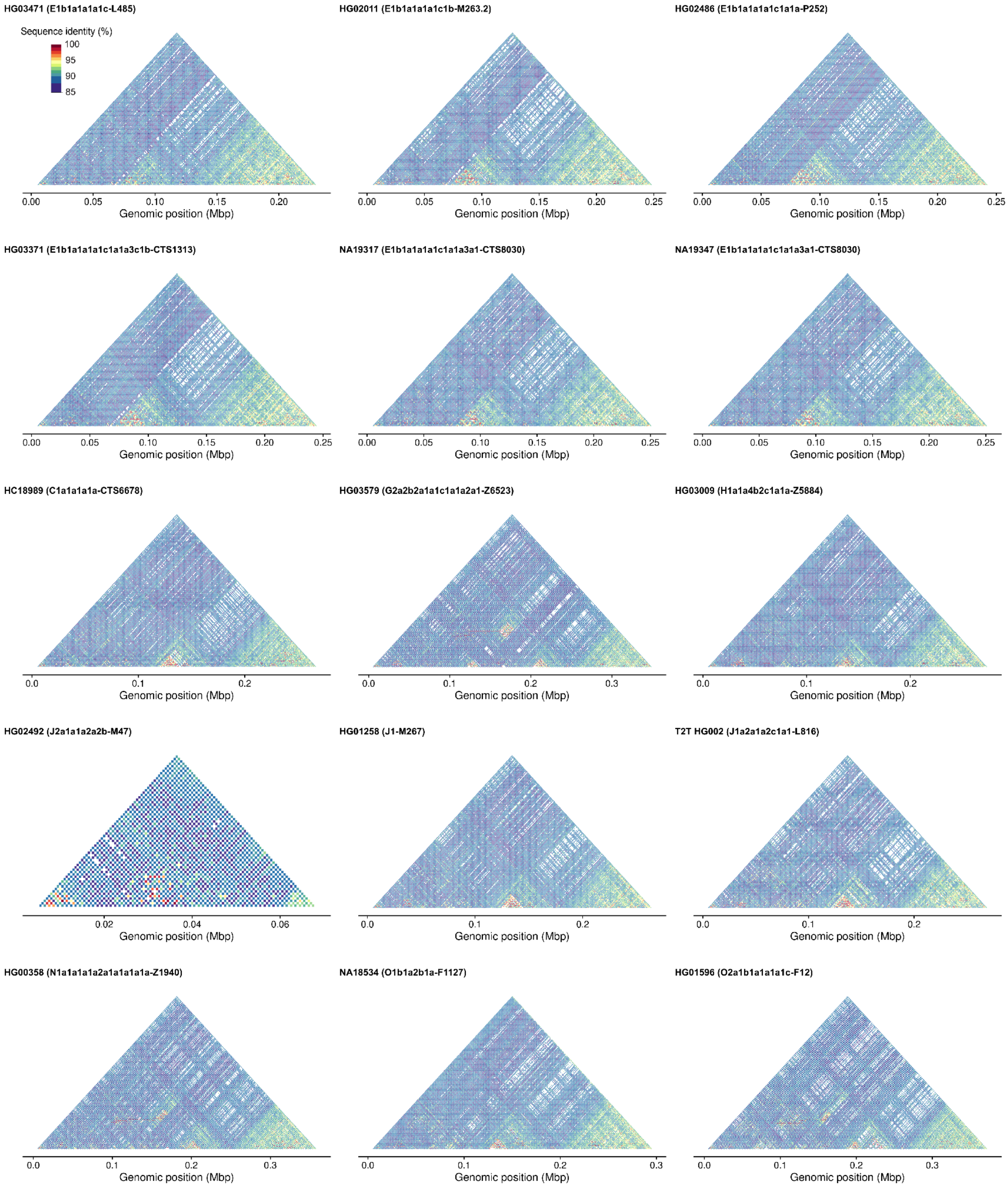


**Figure S22.** Sequence identity heatmaps of the DYZ19 subregion across samples, including the T2T Y, in phylogenetic order. 5000 bp of flanking sequence was added to the DYZ19 genomic interval and 1 kbp window size was used when running StainedGlass.

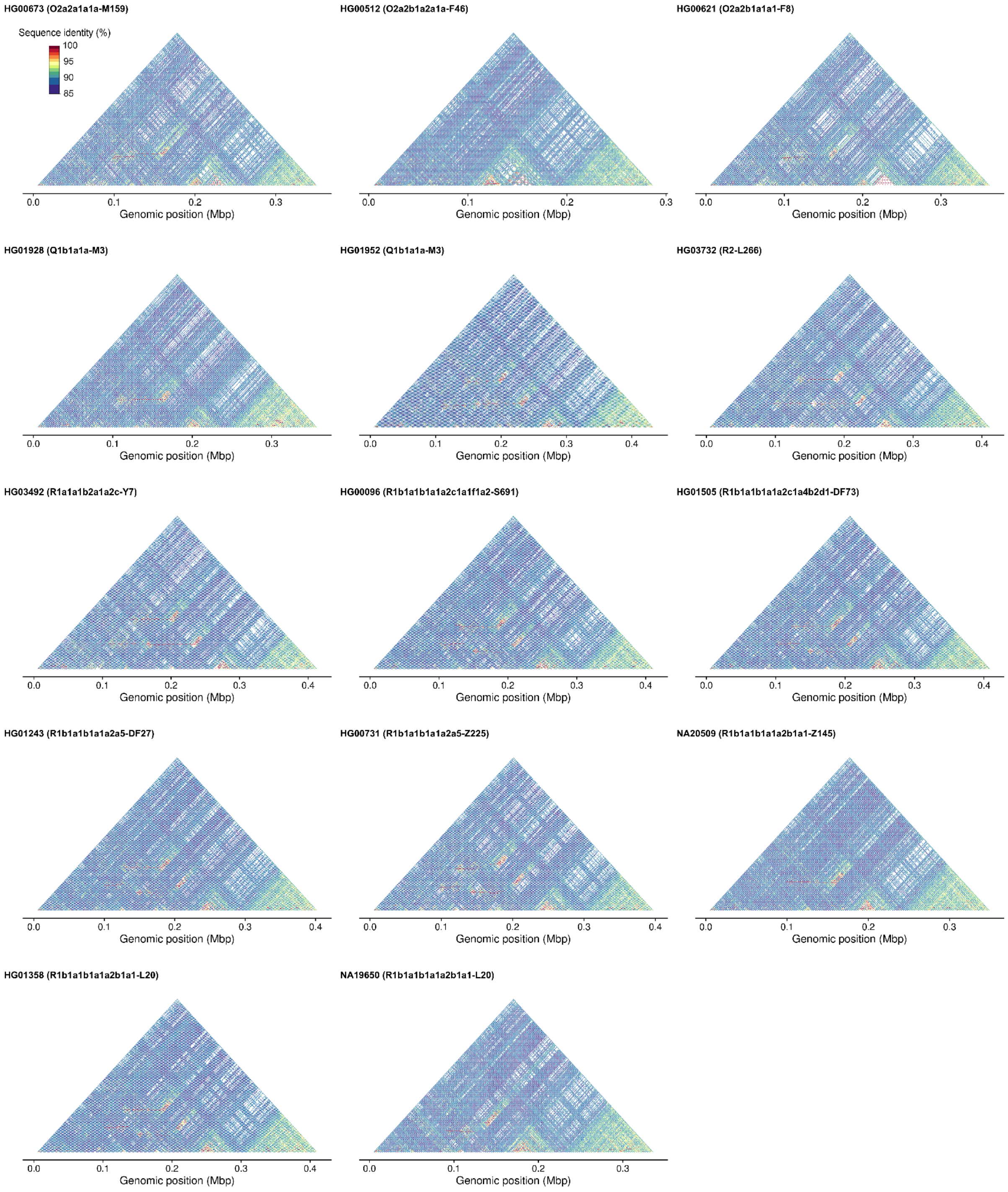


**Figure S23.** Sequence identity heatmaps of the *DYZ19* subregion across samples, including the T2T Y, in phylogenetic order. 5000 bp of flanking sequence was added to the *DYZ19* genomic interval and 1 kbp window size was used when running StainedGlass.

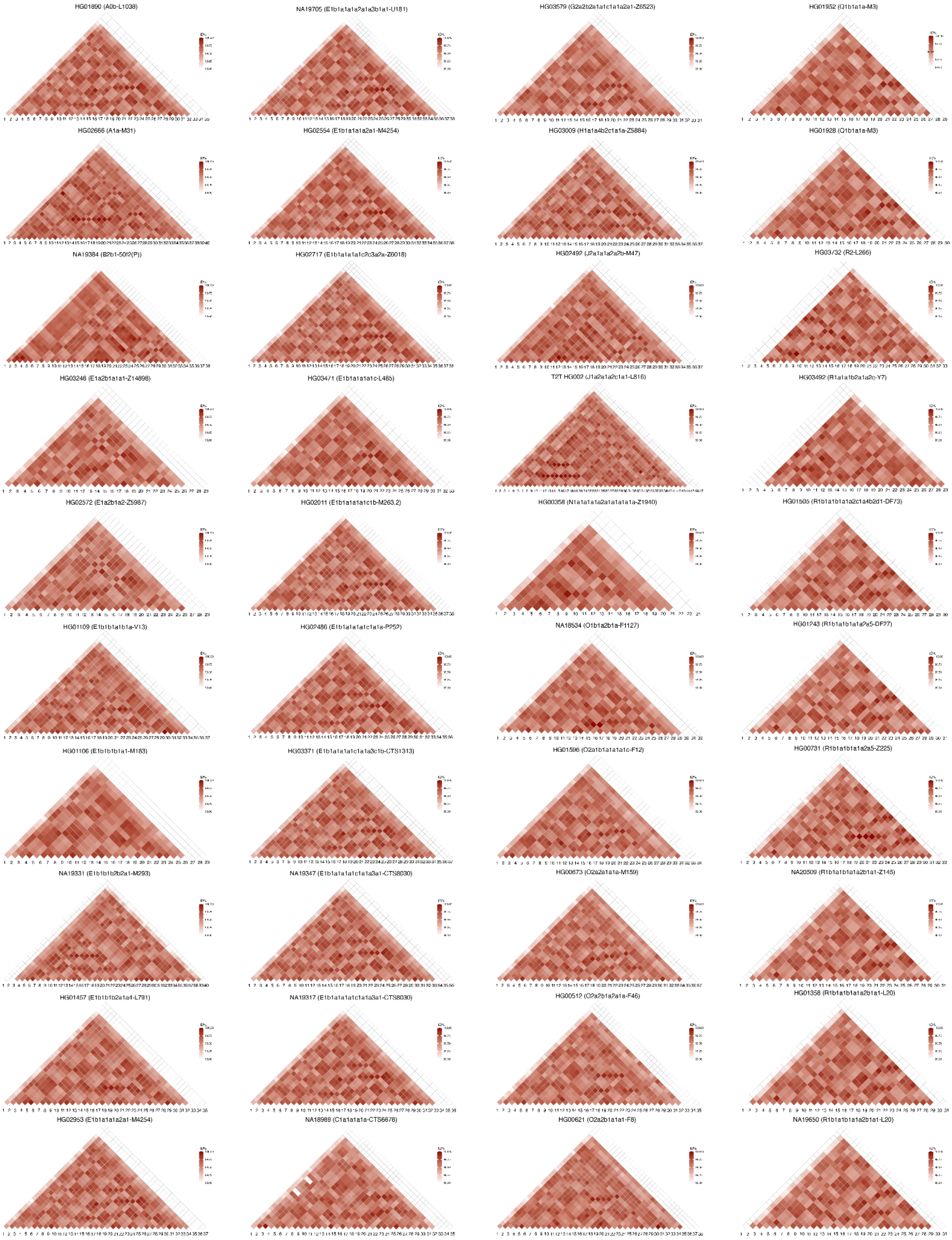

**Figure S24.** Sequence identity heatmaps of ~20.3-kbp long TSPY repeat units for 40 males, including the T2T Y, in phylogenetic order (from top to down from the deepest-rooting sample). Red shades from lighter to darker indicate sequence identity from 99-100%, respectively, while white fill indicates sequence identity below 99%. Due to the IR3/IR3 inversion, three samples (NA19331, HG03732 and HG03492) are displayed in different orientations compared to other samples. In majority of samples the single separate TSPY repeat unit containing the *TSPY2* gene copy is located in between the repeat array and the Y centromere (see **Figure S49** for details) and is therefore the last copy in the heatmap, while in haplogroup QR samples the single copy is located upstream of the repeat array and is therefore the first copy on the heatmap.

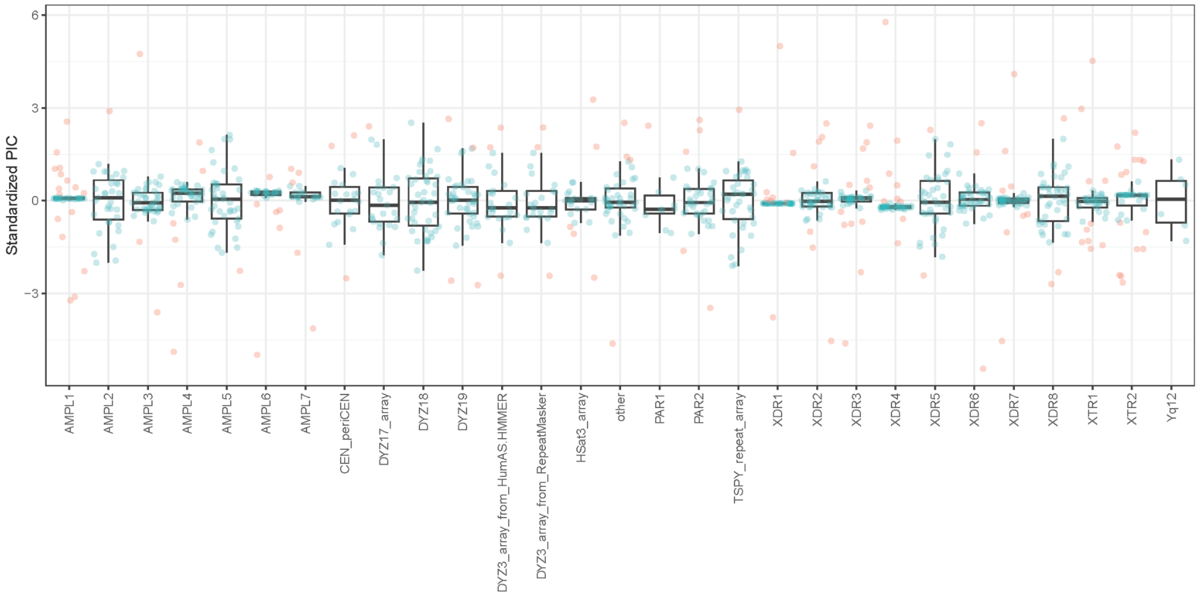


**Figure S25.** Phylogenetically Independent Contrasts (PIC) calculated from contiguously assembled Y-chromosomal sub-region sizes. PICs from each sub-region were standardized for visualization. The green and red dot indicates PICs falling within +/-1.5*IQR (Interquartile range) and PICs falling outside +/-1.5*IQR (outlier), respectively. 86.56% of PICs falls within +/-1.5*IQR on average.

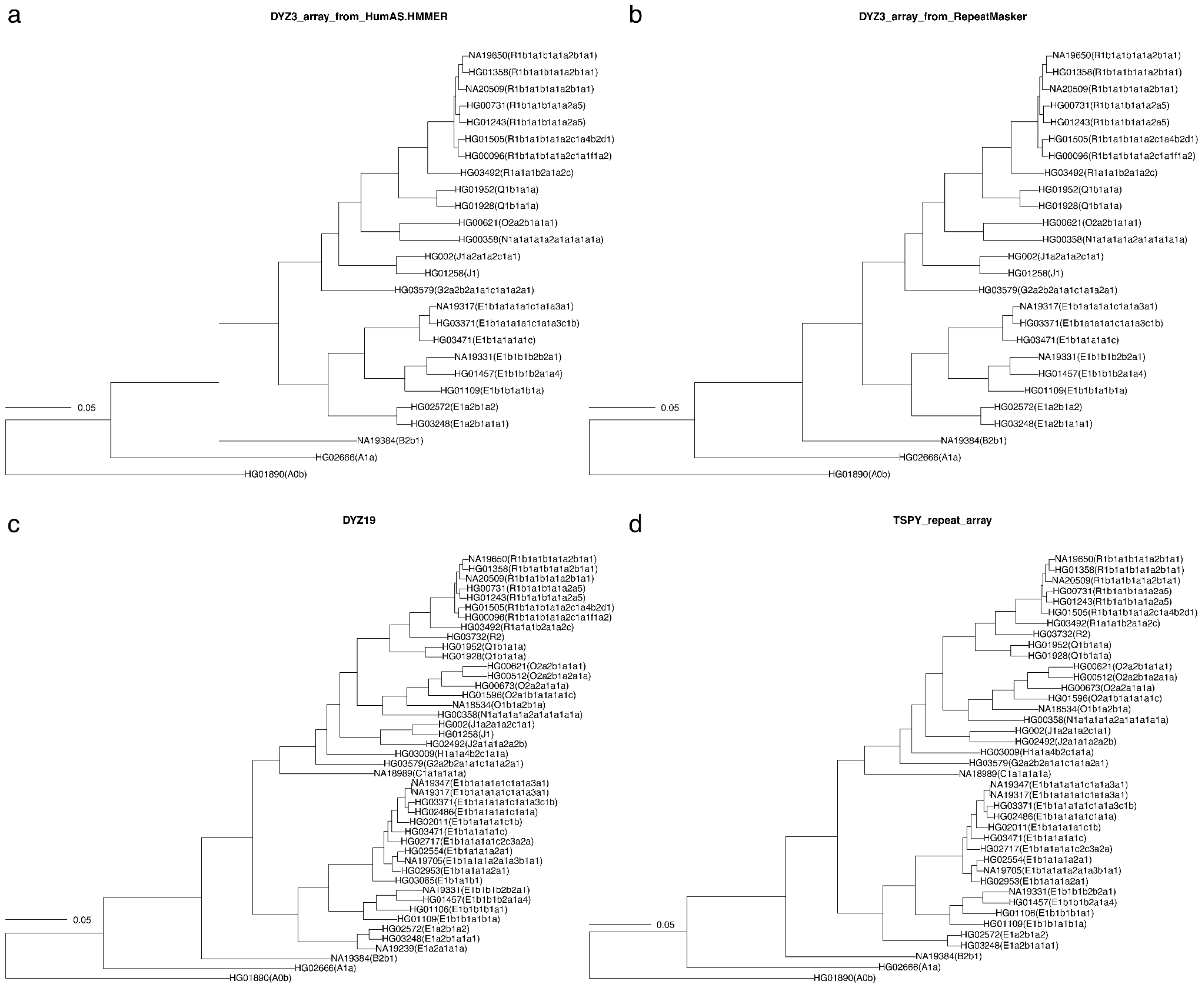


**Figure S26.** Rescaled trees with rescaled branch lengths according to Phylogenetically Independent Contrasts (PIC) calculated using the assembly sizes for **a.** the *DYZ3* array as defined by HumAS-HMMER, **b.** the *DYZ3* array as defined by RepeatMasker, **c.** the *DYZ19* and **d.** the TSPY repeat array.

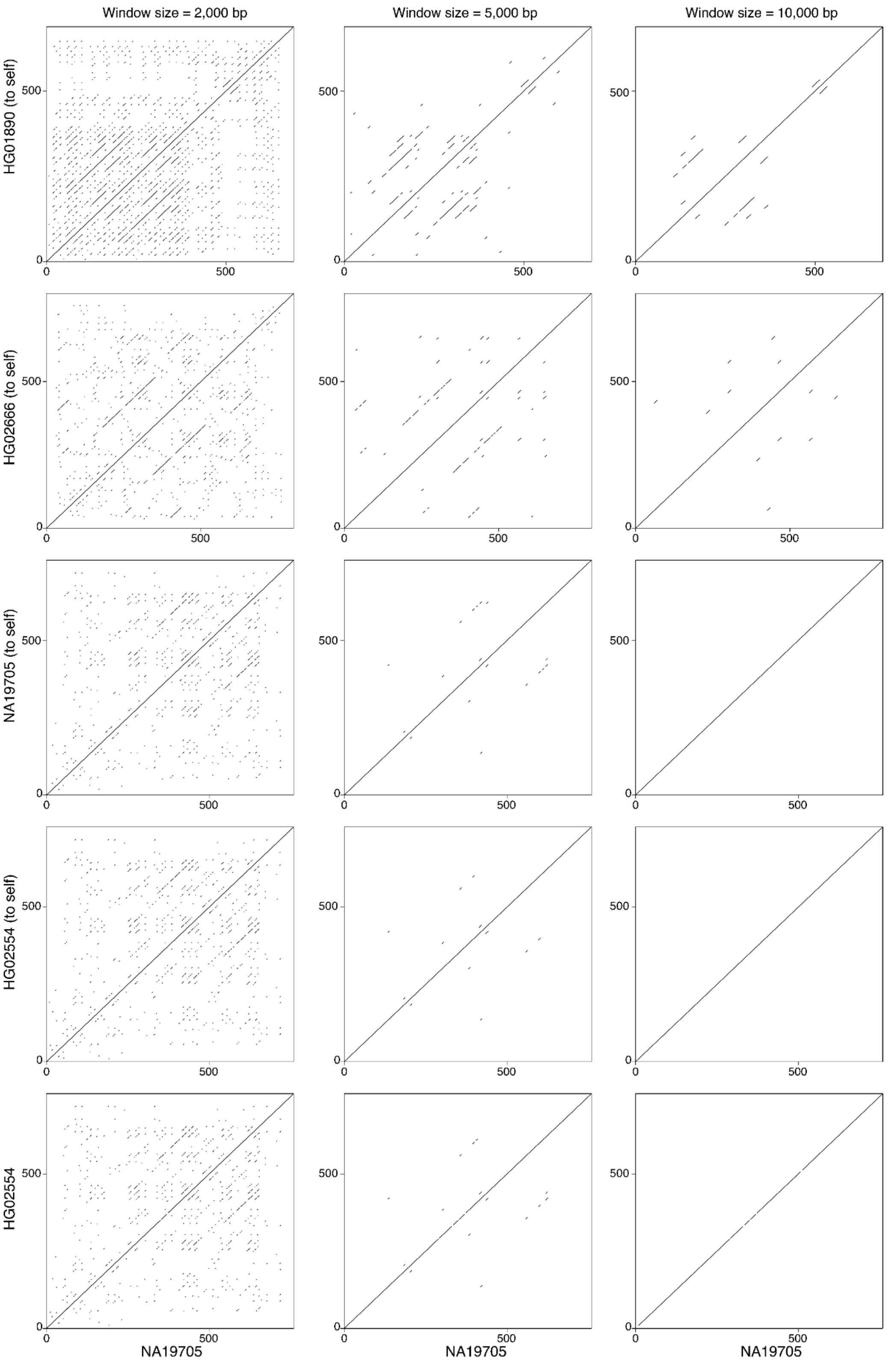


**Figure S27.** Dotplots of the TSPY repeat array with 5 kbp of flanking regions showing identical sequence blocks 2, 5, and 10 kbp in size. From the top - four samples with the comparison to self - HG01890 (A0b-L1038), HG02666 (A1a-M31), NA19705 (E1b1a1a1a2a1a3b1a1-U181) and HG02554 (E1b1a1a1a2a1-M4254). On the bottom - comparison of NA19705 and HG02554. The axes show region size in kbp. Note - NA19705 and HG02554 are phylogenetically closely related with the TMRCA estimated to be 5.5 kya [95% HPD interval: 4.4 - 6.8 kya] (**Figure S1**) with very similar TSPY repeat array sizes (749,363 vs 749,376 bp, respectively) and showing highly similar sequence identity patterns.

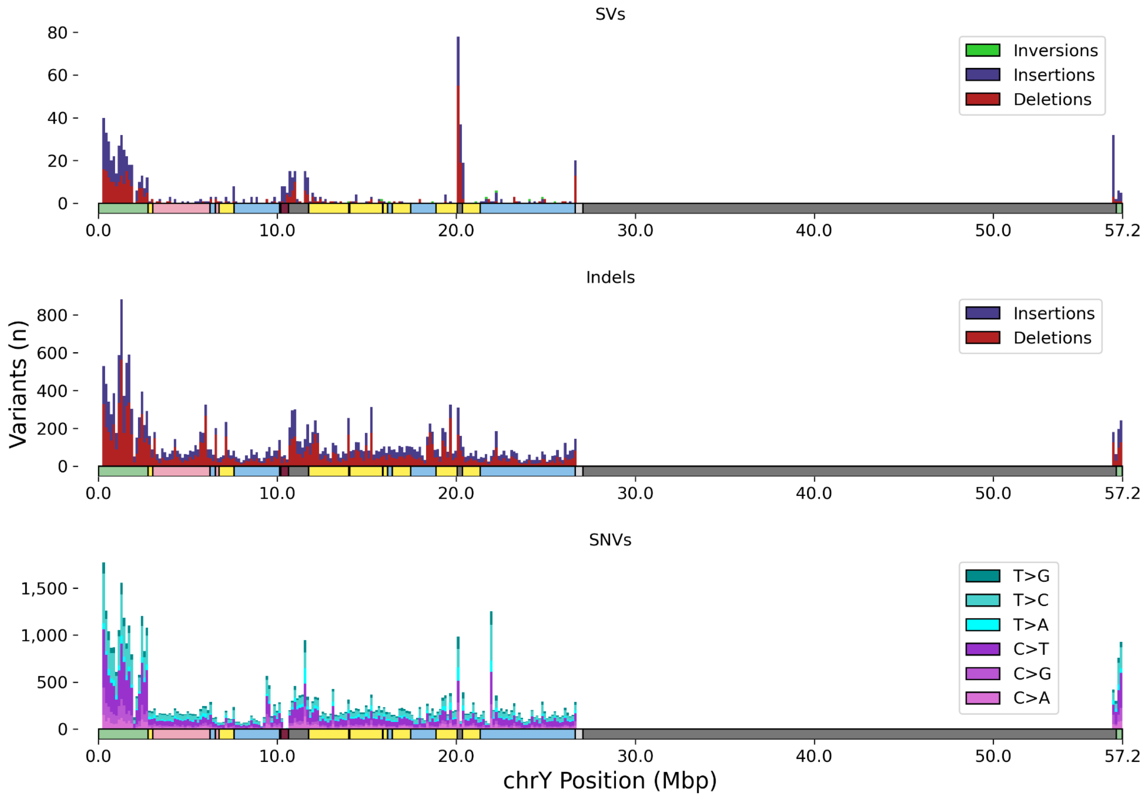


**Figure S28.** Distribution of variant sizes for SVs (≥ 50 bp, top), Indels (< 50 bp, middle), and SNV (bottom) across the Y chromosome (colour by region) as identified using PAV. High peaks in heterochromatin are apparent for SVs, but not SNVs and indels.

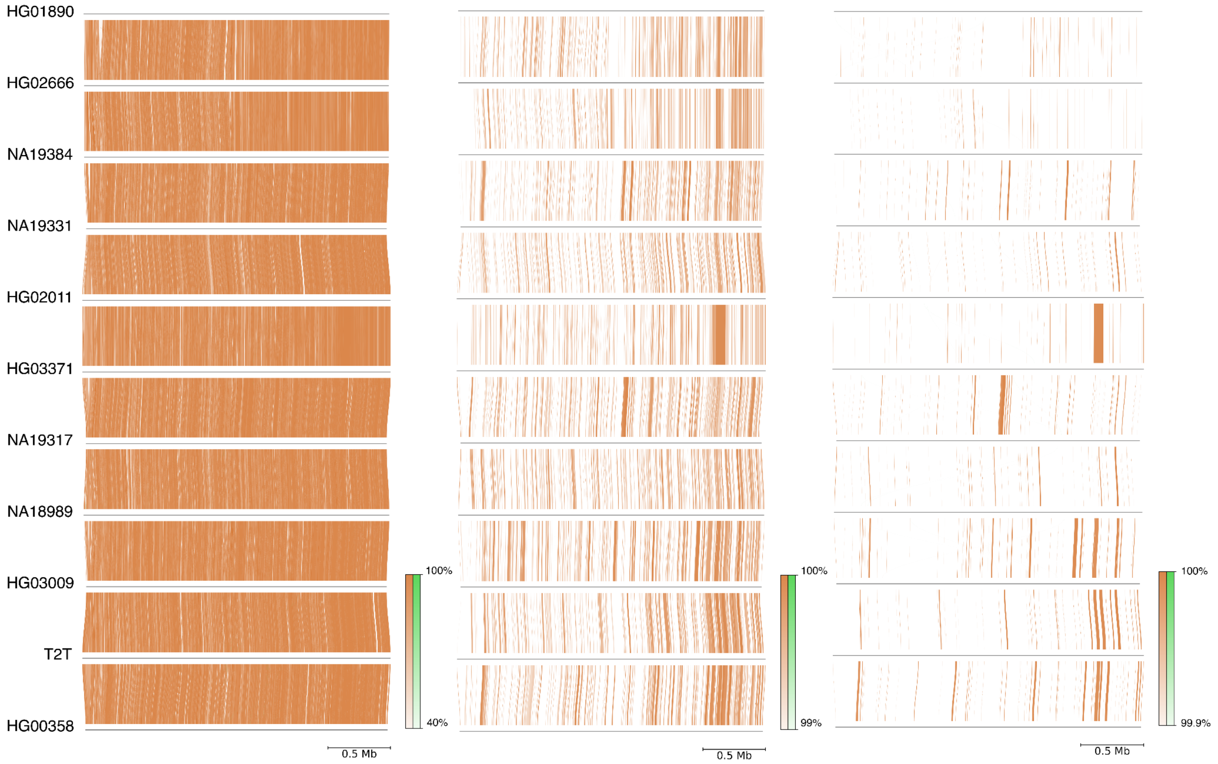


865
866 **Figure S29.** Comparison of sequence identity between the 10 contiguously assembled and the T2T Y PAR1 regions, using
867 different sequence identity thresholds.

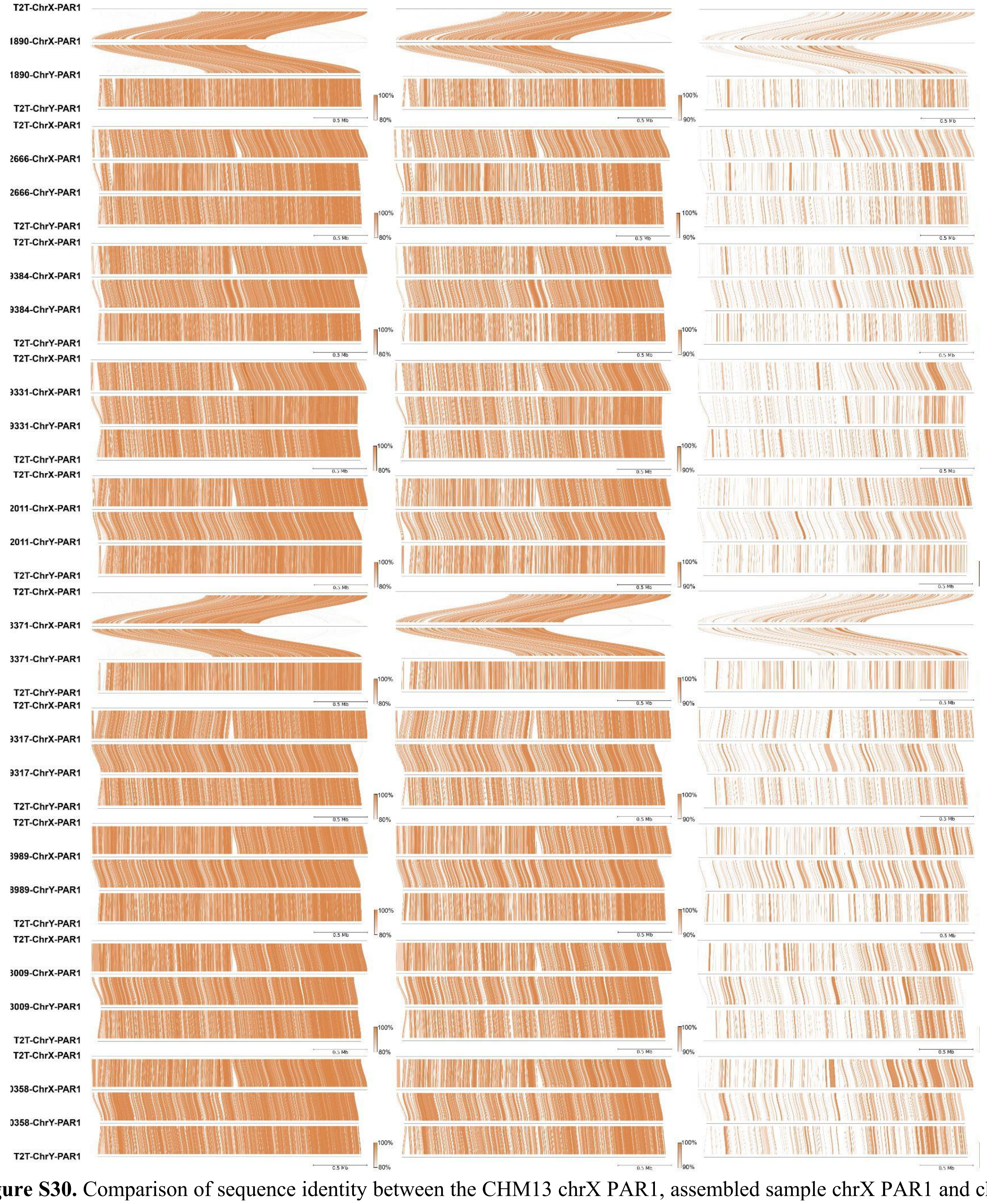


**Figure S30.** Comparison of sequence identity between the CHM13 chrX PAR1, assembled sample chrX PAR1 and chrY PAR1 and the T2T Y PAR1, using different sequence identity thresholds. Note that in two samples (HG01890 and HG03371) approximately 1 Mbp of the telomeric end of chrX PAR1 appears to be missing, either due to a true terminal deletion or an assembly artifact.

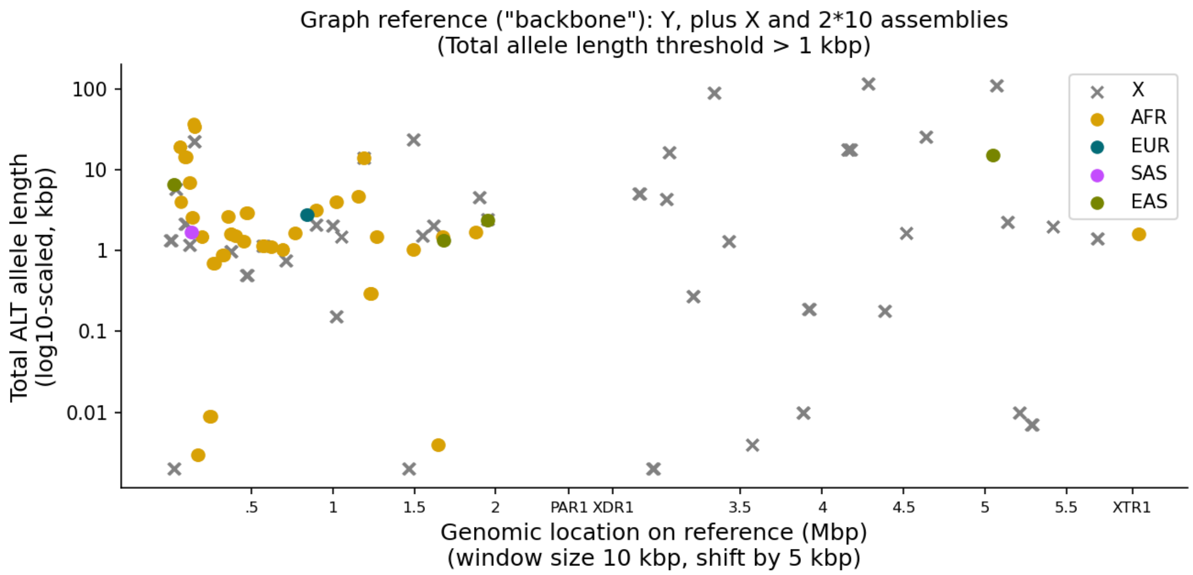


**Figure S31.** PAR1 variation - Y minigraph. Sequence variation in the first 6 Mbp (x-axis) of chromosome Y represented as "bubbles" in a graph constructed using T2T Y as reference (graph backbone), plus T2T X and all X and Y *de novo* assemblies of the 10 samples with a contiguously assembled PAR1 region. Chromosome Y sequence variation is depicted as the total length of all ALT alleles per bubble stratified by continental group (y-axis, coloured circles). Chromosome X sequence variants were aggregated over all X assemblies in the graph (grey X marker). Bubbles with a total allele length (REF plus ALT) of less than 1 kbp are not shown.

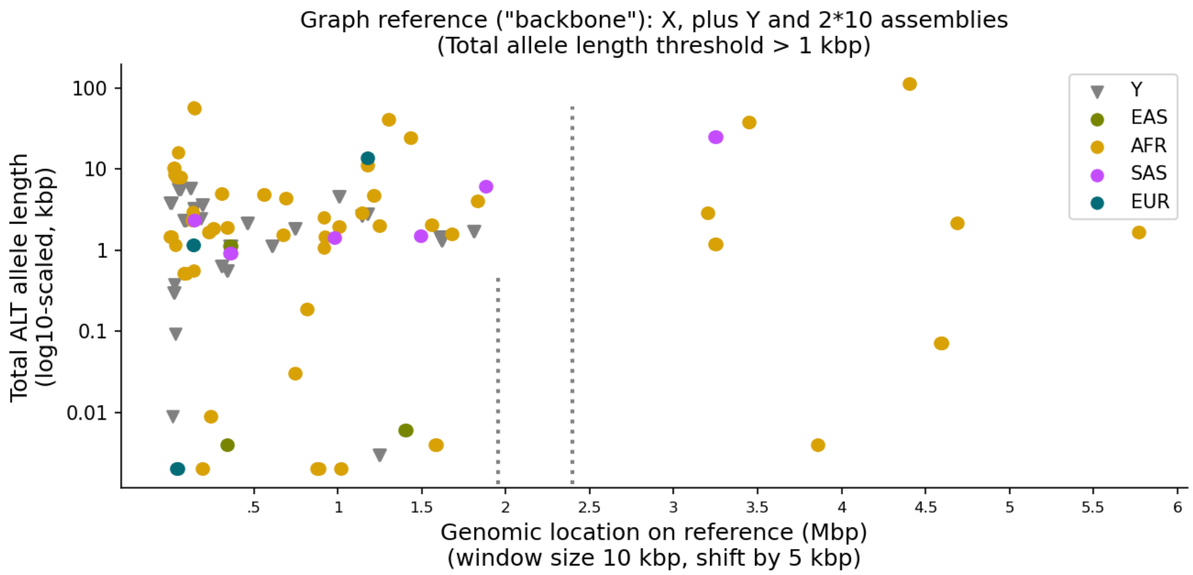


**Figure S32.** PAR1 variation - minigraph X. Sequence variation in the first 6 Mbp (x-axis) of chromosome X represented as "bubbles" in a graph constructed using T2T X as reference (graph backbone), plus T2T Y and all X and Y *de novo* assemblies of the 10 samples with a contiguously assembled PAR1 region. Chromosome X sequence variation is depicted as the total length of all ALT alleles per bubble stratified by continental group (y-axis, coloured circles). Chromosome Y sequence variants were aggregated over all Y assemblies in the graph (grey triangle marker). Bubbles with a total allele length (REF plus ALT) of less than 1 kbp are not shown. Vertical lines mark the last position of a variation bubble before the PAR1 boundary in chromosome Y (left line), and the extrapolated PAR1 boundary in chromosome X (right line) based on the distance between the last variation bubble and the PAR1 boundary in chromosome Y (~504 kbp).

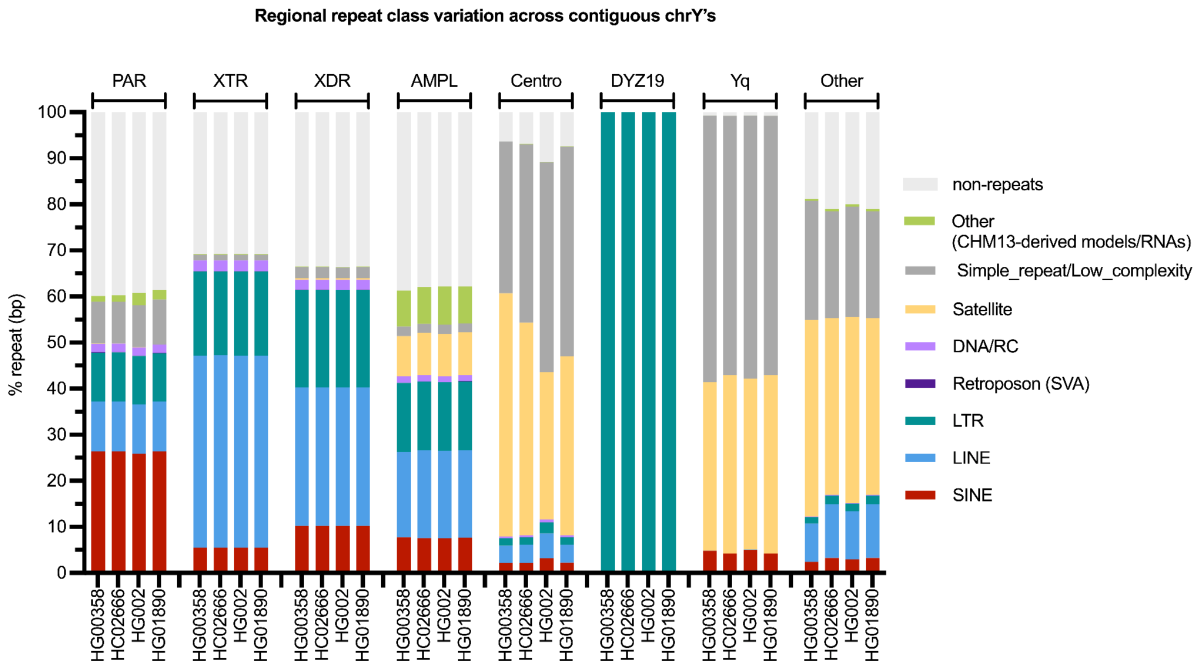


**Figure S33.** Regional repeat class variation across the three contiguously assembled and the T2T Y chromosome sequences. Please note HG01890 was included despite containing a single gap in the PAR2 region.

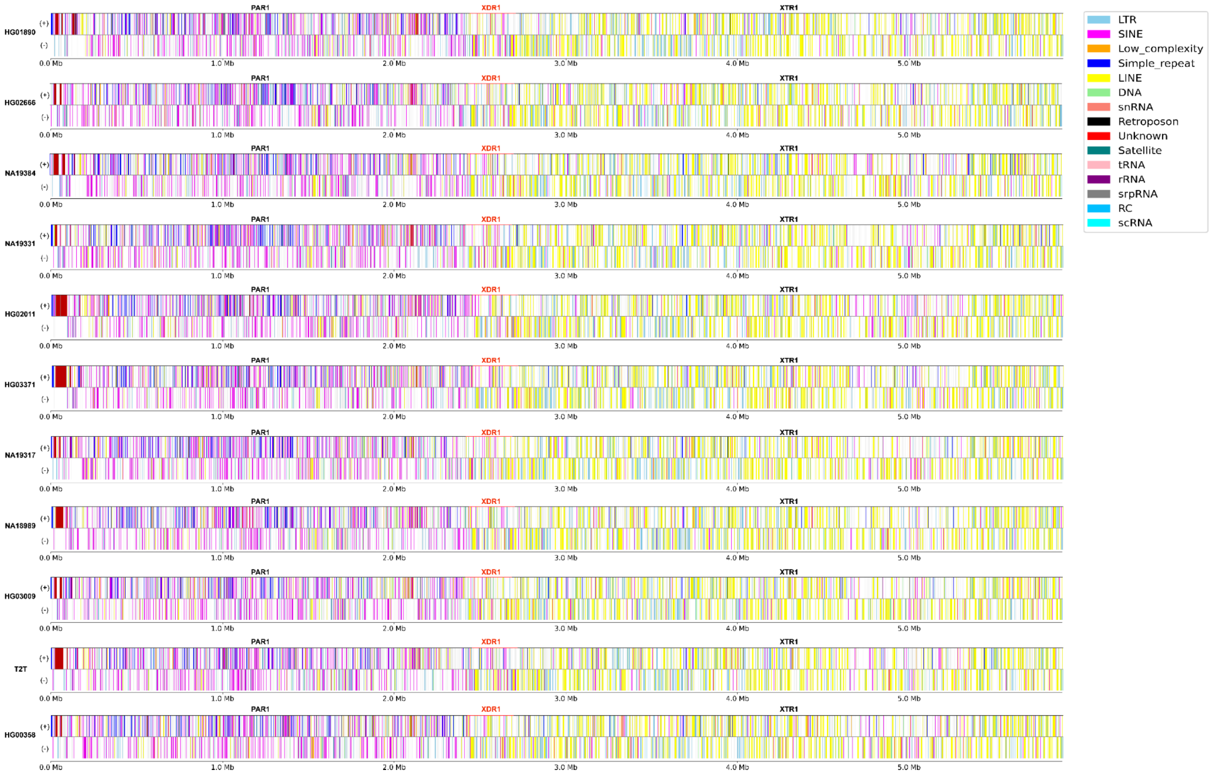


**Figure S34.** Repeat element distribution across 10 contiguously assembled and the T2T Y-chromosomal PAR1 regions. Repeat elements on sense (+) and antisense (-) strand are shown, coloured according to repeat class. Extensive differences in size can be seen between samples, especially in the satellite arrays located close to the telomere (in dark red). The locations of PAR1, XDR1 and XTR1 subregions in each individual are shown in black, red and black, respectively. Please note that the maroon colour of the "Unknown" elements close to the telomere is caused by significant clustering of those elements.

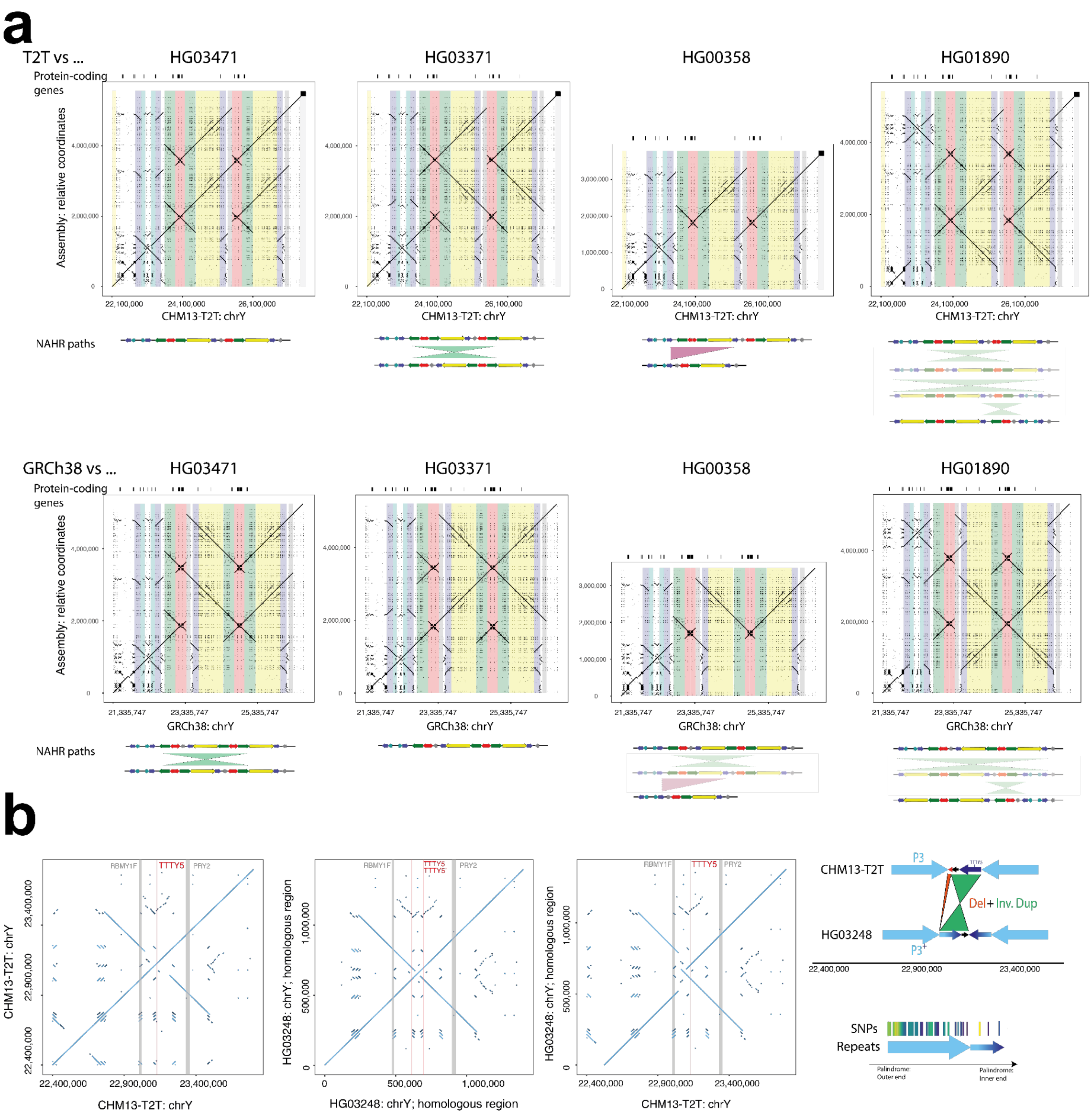


**Figure S35.** Examples of structural variation identified in the de novo assembled Y chromosomes. **a.** Inversions identified in the AZFc/ampliconic 7 subregion. Top - comparison between the T2T Y and the de novo assemblies, bottom - GRCh38 Y and the de novo assemblies. Potential NAHR path is shown below the dotplot. **b.** Inverted duplication affecting roughly two thirds of the 161 kbp unique 'spacer' sequence in the P3 palindrome, spawning a second copy of the TTTY5 gene and elongating the LCRs in this region. A detailed sequence view reveals a high sequence similarity between the duplication and its template and its placement in Y phylogeny supports emergence of this variant in the common ancestor of haplogroup E1a2 carried by NA19239, HG03248 and HG02572 (**Fig. 3a**).

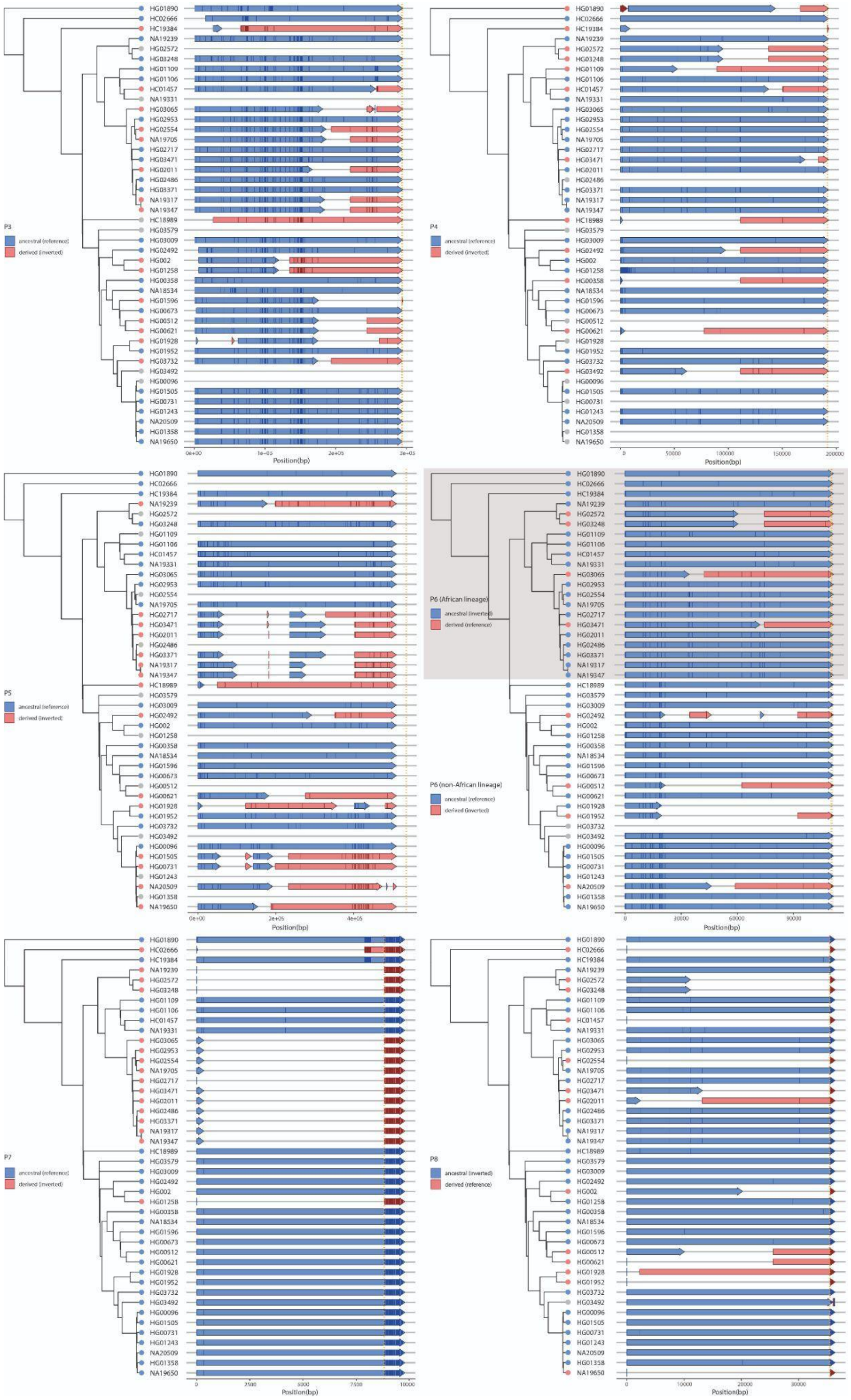


**Figure S36.** Breakpoint locations identified for 6 euchromatic inversions in palindromes P3, P4, P5, P6, P7 and P8. The red tip colours (derived state) in the phylogenetic tree indicate samples which have undergone an inversion and therefore carry the 'spacer' region in inverted orientation compared to samples with blue tip (ancestral state). Informative PSV positions are shown as vertical lines with darker colour in each of the arrows. The orange dotted line indicates the start of the unique 'spacer' region. Any information that is not available is indicated by grey. In P6, breakpoint locations were determined separately for African Y lineages (haplogroups A, B and E, grey shaded area) and non-African Y lineages, using two different sets of ancestral and derived states.

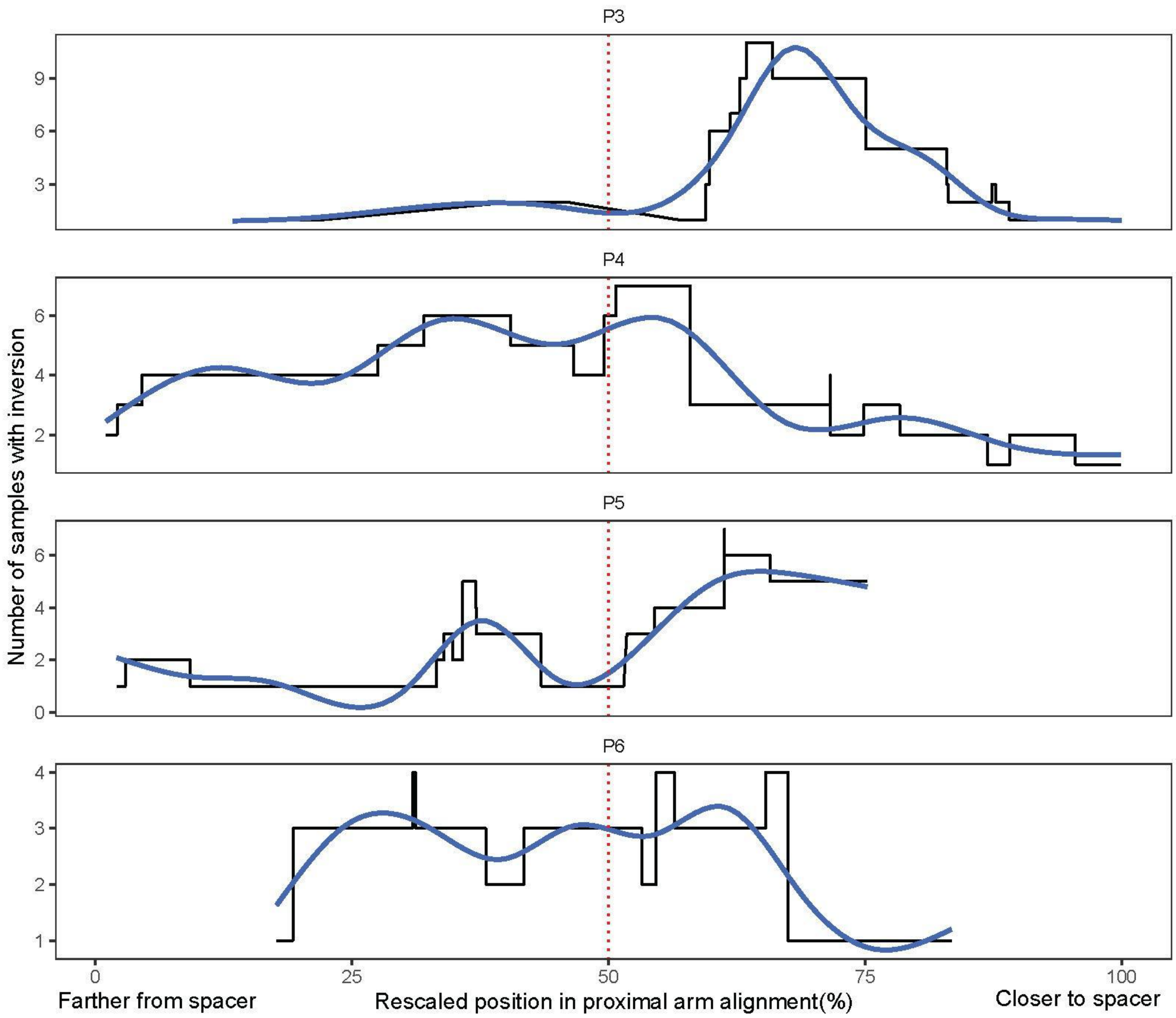


**Figure S37.** Rescaled breakpoint locations identified for 4 euchromatic inversions in palindromes P3, P4, P5, and P6. The start and end positions of each breakpoint range were rescaled to have the same start (0%) and end position (100%) across 4 palindromes. The y-axis indicates the number of samples that have inversion breakpoints at the corresponding position in the x-axis. The trend line indicated in blue is displayed by a smoothing function implemented in ggplot2 (geom_smooth, method =”gam”). P7 and P8 were excluded due to the small number of informative PSVs and therefore, wide breakpoint ranges.

| Distance between neighbouring PSVs | 125 | 125 | 37 | 76 | 86 | 84 | 76 | 233 | 193 | 22 | 351 | 105 | 27 | 294 | 2 | 1 | 1 | 1 | 1 | 8 | 35 | 8 | |
|---|---|---|---|---|---|---|---|---|---|---|---|---|---|---|---|---|---|---|---|---|---|---|---|
| IR5-1_HG02666 | G | A | G | G | A | A | G | C | G | - | T | A | T | C | C | - | - | - | - | - | T | T | T |
| IR5-2_HG02666 | T | G | A | A | G | G | A | T | A | C | T | G | G | C | C | - | - | - | - | - | T | T | T |
| IR5-3_HG02666 | G | A | G | A | A | A | G | C | G | - | T | G | G | A | T | T | T | T | A | T | C | C | C |
| IR5-4_HG02666 | T | G | A | A | G | G | A | T | A | C | T | G | G | A | T | T | T | T | A | T | C | C | C |
| IR5-1_NA19384 | G | A | G | G | A | A | G | C | G | - | T | A | T | C | C | - | - | - | - | - | T | T | T |
| IR5-2_NA19384 | G | A | G | G | A | A | G | C | G | - | T | A | T | C | C | - | - | - | - | - | T | T | T |
| IR5-3_NA19384 | T | G | A | A | G | G | A | T | A | C | C | G | G | A | T | T | T | T | A | T | C | C | C |
| IR5-4_NA19384 | T | G | A | A | G | G | A | T | A | C | T | G | G | A | T | T | T | T | A | T | C | C | C |
| IR5-1_HG01890 | G | A | G | G | A | A | G | C | G | - | T | A | T | C | C | - | - | - | - | - | T | T | T |
| IR5-2_HG01890 | G | A | G | G | A | A | G | C | G | - | T | A | T | C | C | - | - | - | - | - | T | T | T |
| IR5-3_HG01890 | T | G | A | A | G | G | A | T | A | C | T | G | G | C | T | T | T | T | A | T | C | C | C |
| IR5-4_HG01890 | T | G | A | A | G | G | A | T | A | - | T | G | G | A | T | T | T | T | A | T | C | C | C |

**Figure S38.** Inversion breakpoint identification for the IR5/IR5 inversion in HG02666. The alignment shows all 4 IR5 repeats from three samples (HG02666 - inverted, NA19384 and HG01890 - reference orientation), with only informative PSV positions and genotypes shown (i.e., sites identical between the IR5 repeats and across individuals have been removed for visualization purposes). In NA19384 and HG01890 the IR5 repeats located within the P5 palindrome (IR5-1 and IR5-2) show a distinct PSV pattern from the IR5 copies located within the P1 palindrome. HG02666 which carries an inversion, the change of this pattern indicates the location of the inversion breakpoints and is highlighted by a black box. Inversion breakpoints relative to GRCh38 are: chrY:18,036,429-18,036,932 and chrY:24,036,893-24,037,396 for proximal and distal breakpoints, respectively.

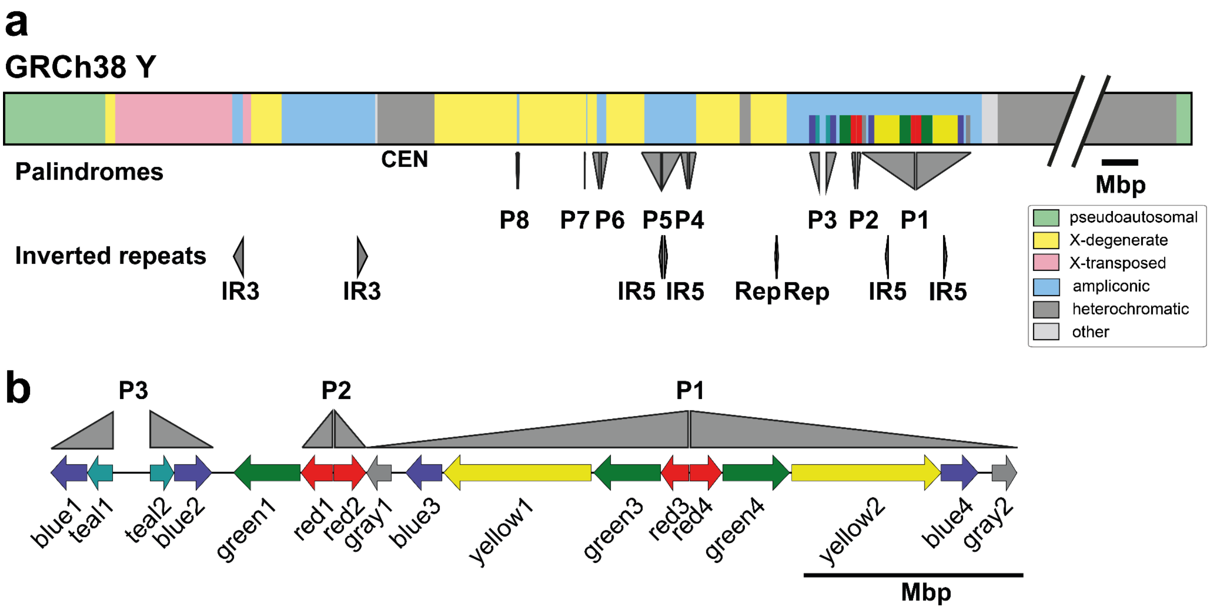


**Figure S39.** Schematic representation of inverted repeats involved in inversions. **a.** The GRCh38 Y reference structure with annotations of segmental duplications in *AZFc*/ampliconic subregion 7, with palindromes (P8-P1) and inverted repeats (IR) shown below. The repeat coordinates relative to GRCh38 Y reference sequence were obtained from Teitz et al [13]. **b.** Annotation of segmental duplications in *AZFc*/ampliconic subregion 7 following the naming originally proposed by Kuroda-Kawaguchi et al [45].

**a**

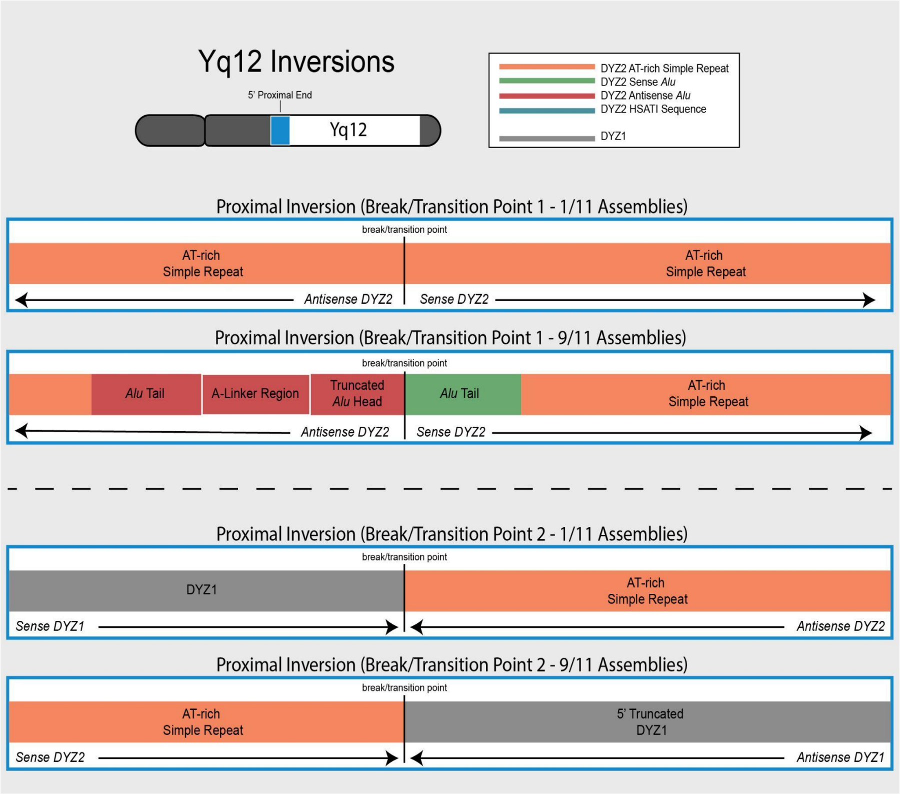


**b**

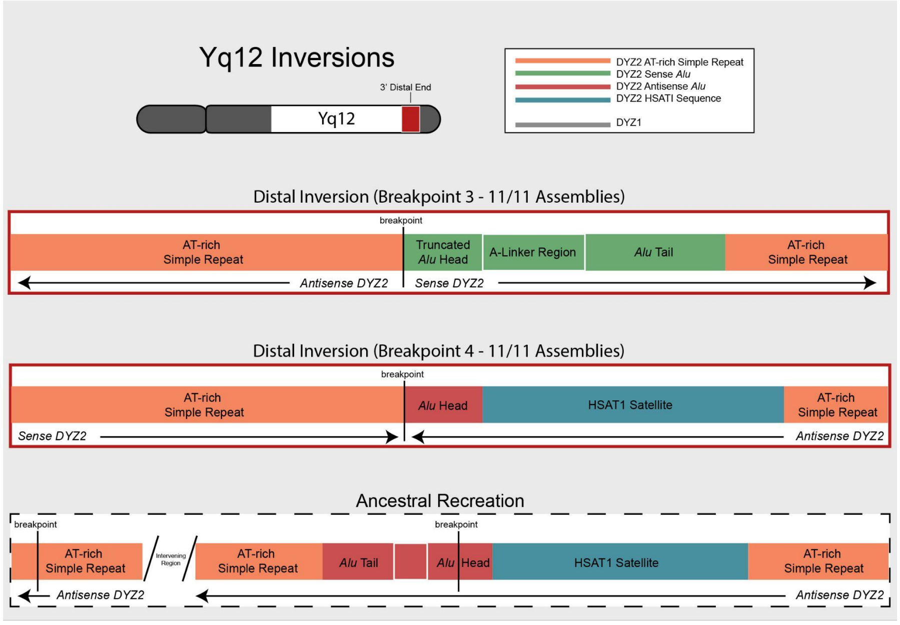


**Figure S40.** Inversions identified in the proximal and distal regions of the Yq12 subregion. **a.** Shows the proximal inversion break/transition regions. The top two rows show the inversion found in HG01890 and the bottom two rows the nine other genomes. The proximal inversion region is deleted in HG01106. The inversion break/transition points are described as such since the coordinates for where region changes orientation is located (see **Table S34**), but the exact breakpoint has not been elucidated. **b.** Shows the distal inversion breakpoints (top row) shared in all genomes, as well as an ancestral recreation of the region before the inversion.

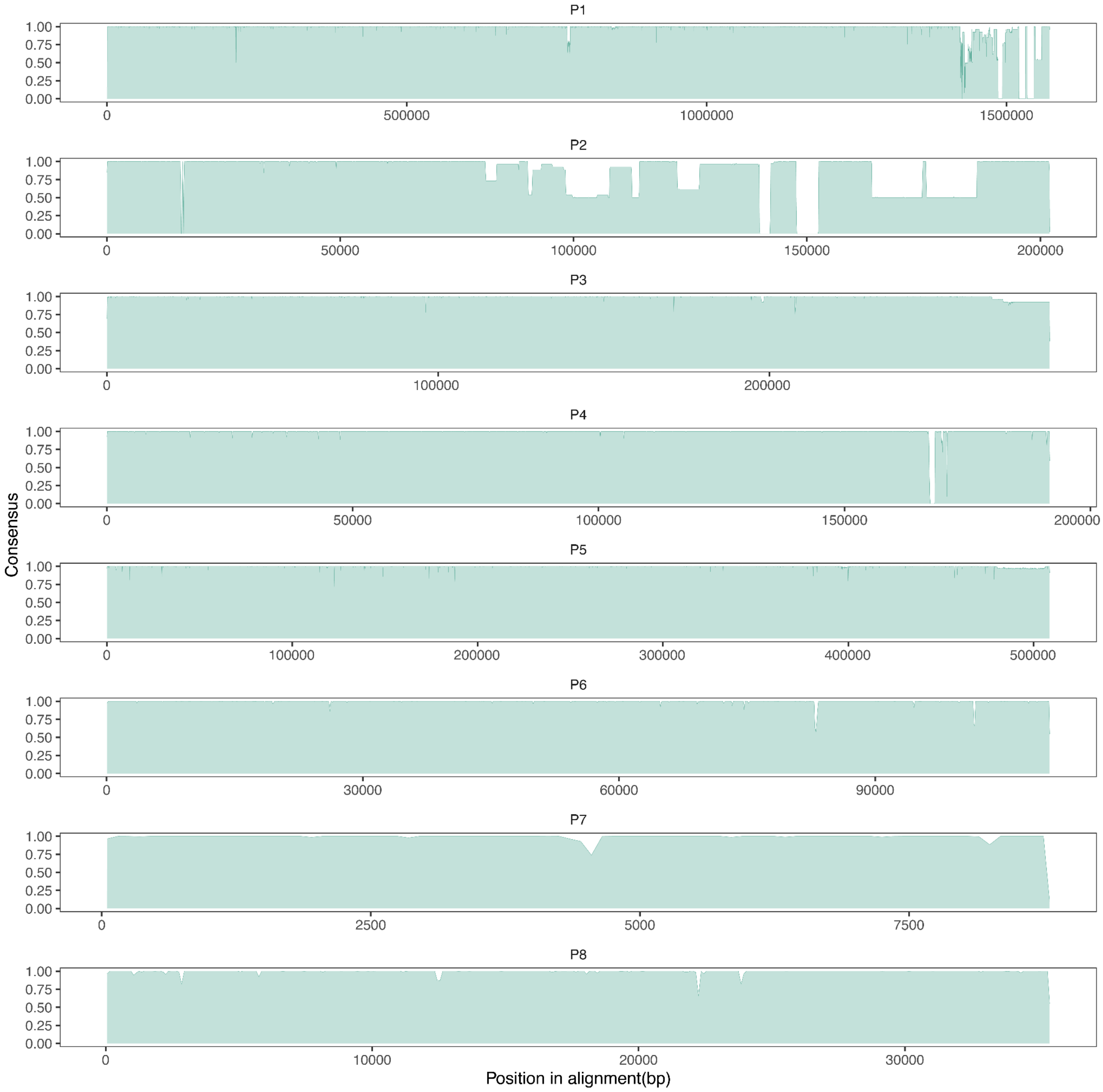


**Figure S41.** Alignment summary of 8 Y-chromosomal palindromes. The "consensus" on y-axis indicates a frequency of major allele (including a gap) calculated at each position in the alignment and averaged across 100 bp windows. The alignments from the same set of 13 samples are shown as used for **Figure S42**.

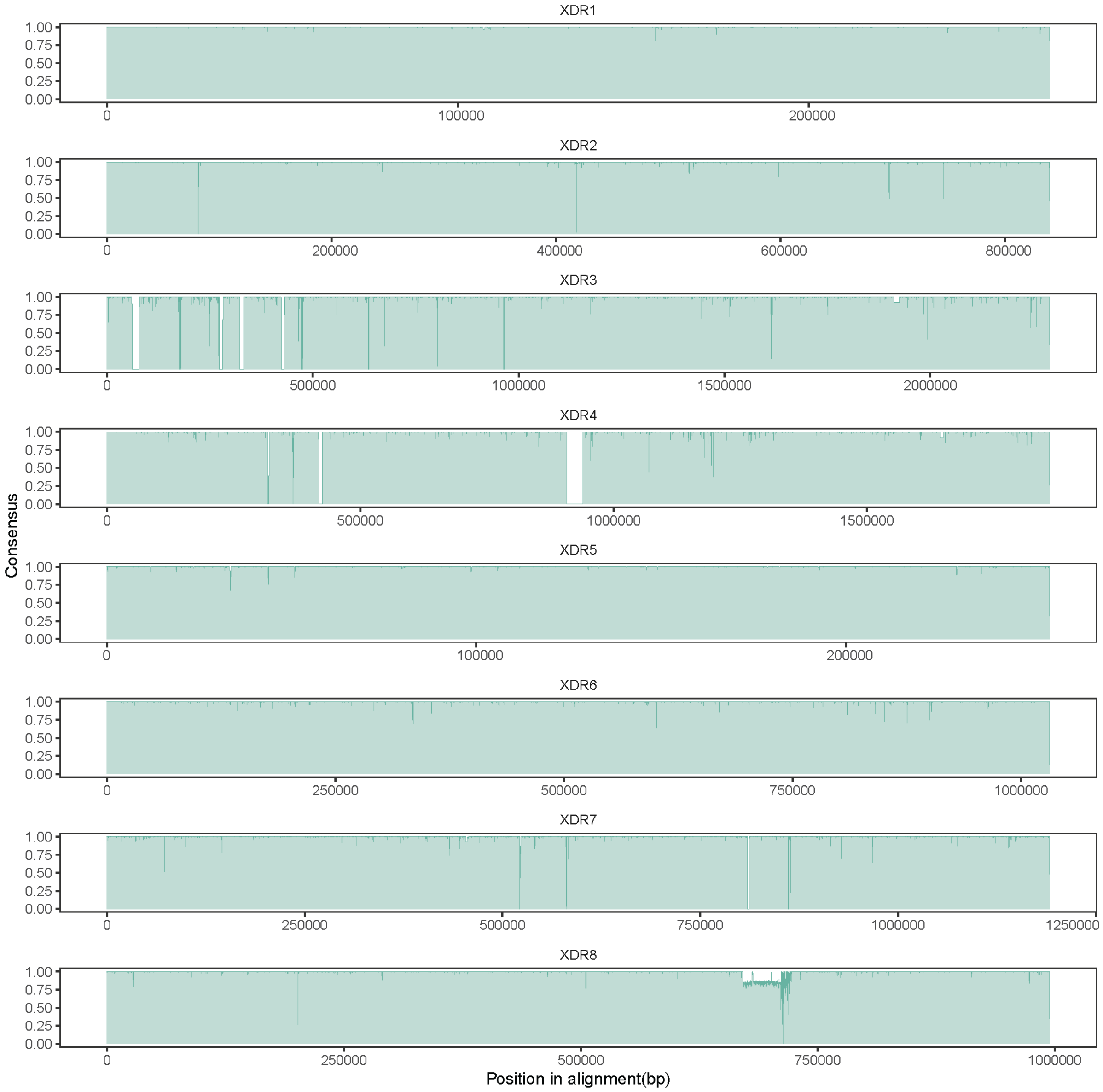


**Figure S42.** Alignment summary of XDR regions. The "consensus" on y-axis indicates a frequency of major allele (including a gap) calculated at each position in the alignment and averaged across 100 bp windows. The alignments from the same set of 13 samples are shown as used for **Figure S41**.

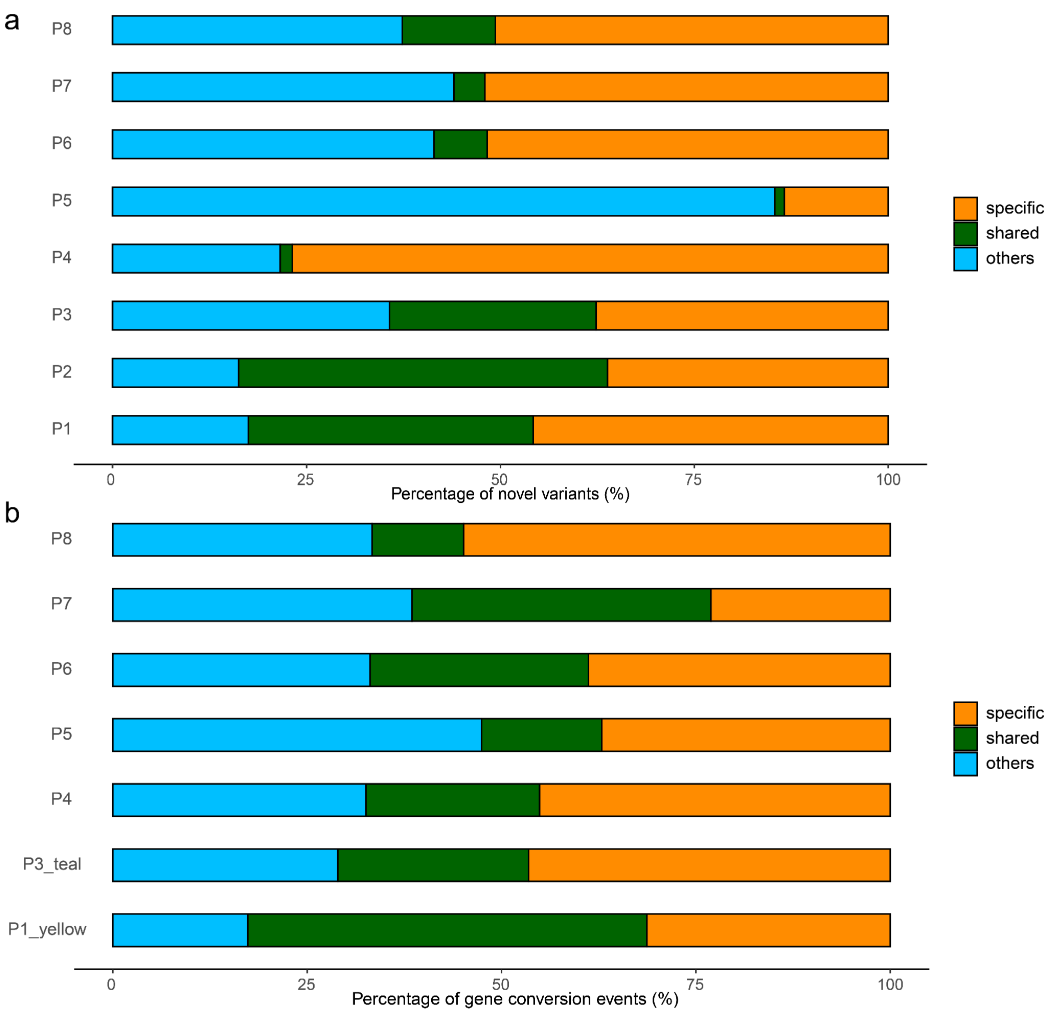


**Figure S43.** New variations and gene conversion events identified in Y haplogroups A, B and E versus others. **a.** Percentage of novel variants specific to (denoted by "specific") and shared by (denoted by "shared") A, B, and E haplogroups. The remaining variants are classified as "others". **b.** Percentage of gene conversion events specific to (denoted by "specific") and shared by (denoted by "shared") A, B, and E haplogroups. The remaining variants are classified as "others".

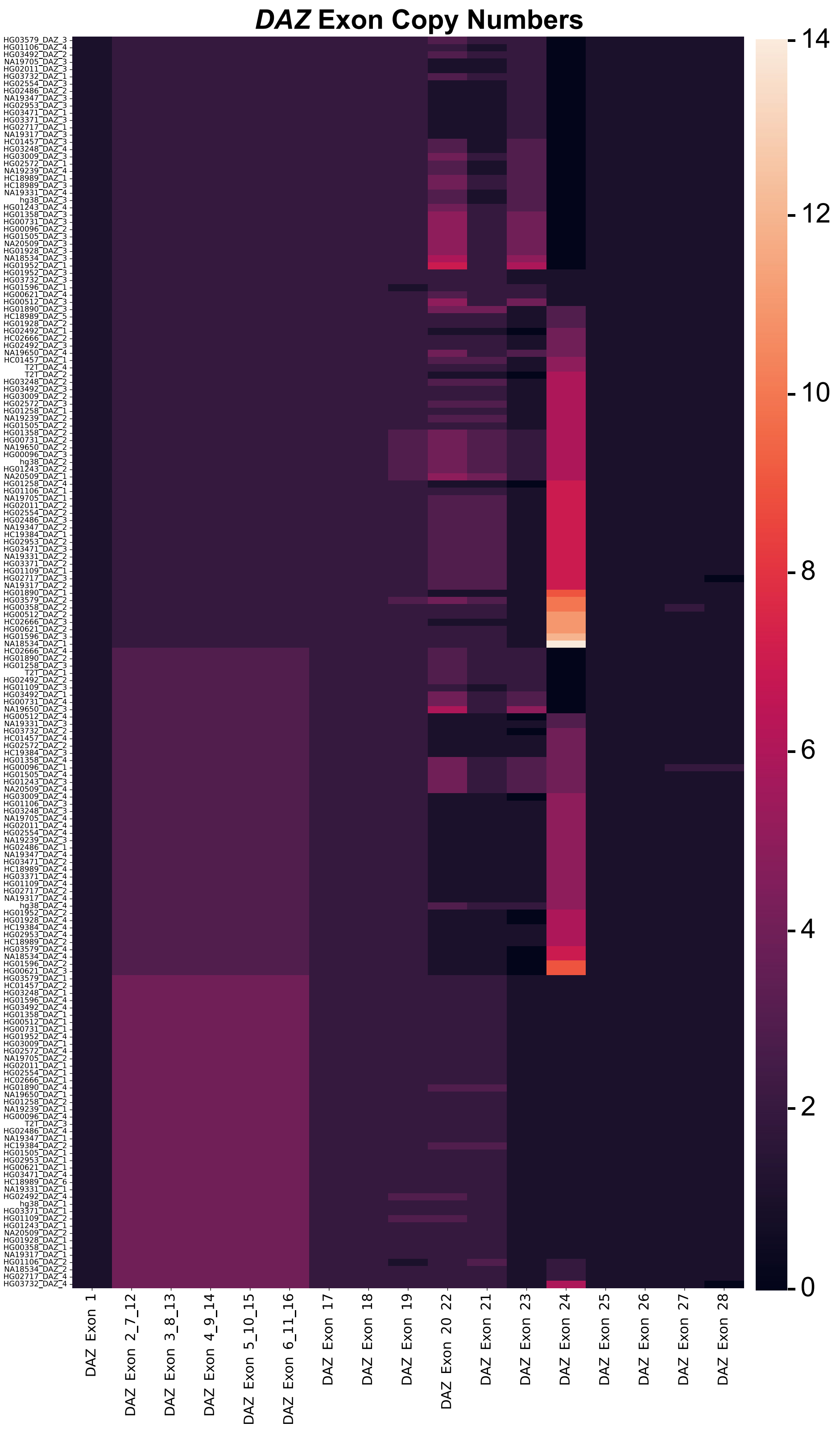


**Figure S44.** Heatmap showing the detected number of exons within individual DAZ genes across the 43 (41 + T2T Y and GRCh38) analysed assemblies. Exons with the exact same nucleotide sequence were grouped with one another (e.g., Exons 2, 7, 12). Results show a large variation in exon compositions of DAZ genes. Exon 24 of DAZ has the largest variation in copy number ranging from zero to fourteen copies. The large variation in exon copy numbers made it fruitless to align sequences and construct trees.

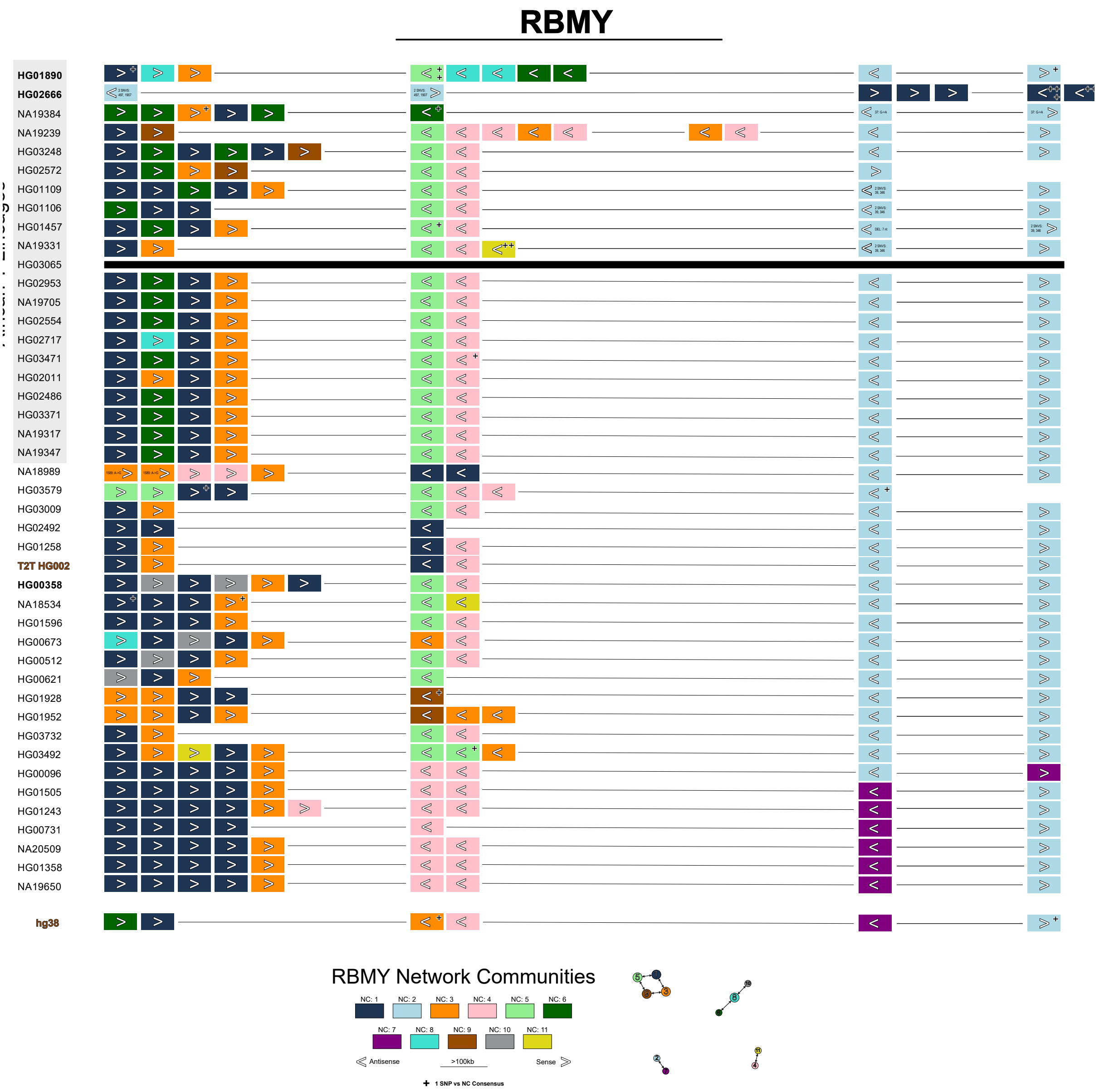


**Figure S45.** Network Analysis and Visualization of *RBMY1* gene copies. This figure shows the *RBMY1* gene copies and regions within each Y chromosome assembly that were analysed (42 + T2T + GRCh38). Individual *RBMY1* gene copies are shown as rectangles. The colour of an individual *RBMY1* copy is based on the network community (NC) assignment (see **Methods**). An arrow pointing to the right indicates the gene copy was in the sense orientation and an arrow pointing to the left indicates the gene copy was in antisense orientation. A black line between gene copies indicates that they are no less than 100 kbp away from one another and a black cross (+) indicates a single SNP compared to the NC consensus sequence. *RBMY1* gene copy SNV differences compared to the assigned NC consensus sequence are only annotated if more than one copy shares the same SNV. Additionally, a secondary directed network was created from the network community consensus sequences (shown near the legend). The size of a node is representative of the total *RBMY1* gene sequences that were assigned to the network community. An edge going from one node to another represents that the second node was the first's best match (i.e., most similar sequence; ties are allowed and shown as multiple edges on a node). The width of the edge represents how similar two nodes are where a thicker edge indicates a more similar sequence (i.e., less SNVs). Overall, the visualization shows (albeit simplified) the four primary *RBMY1* gene copy regions, although one sample (NA19239) has a partial duplication of gene region 2 and another (HG02666) suggests at least one inversion of the regions.

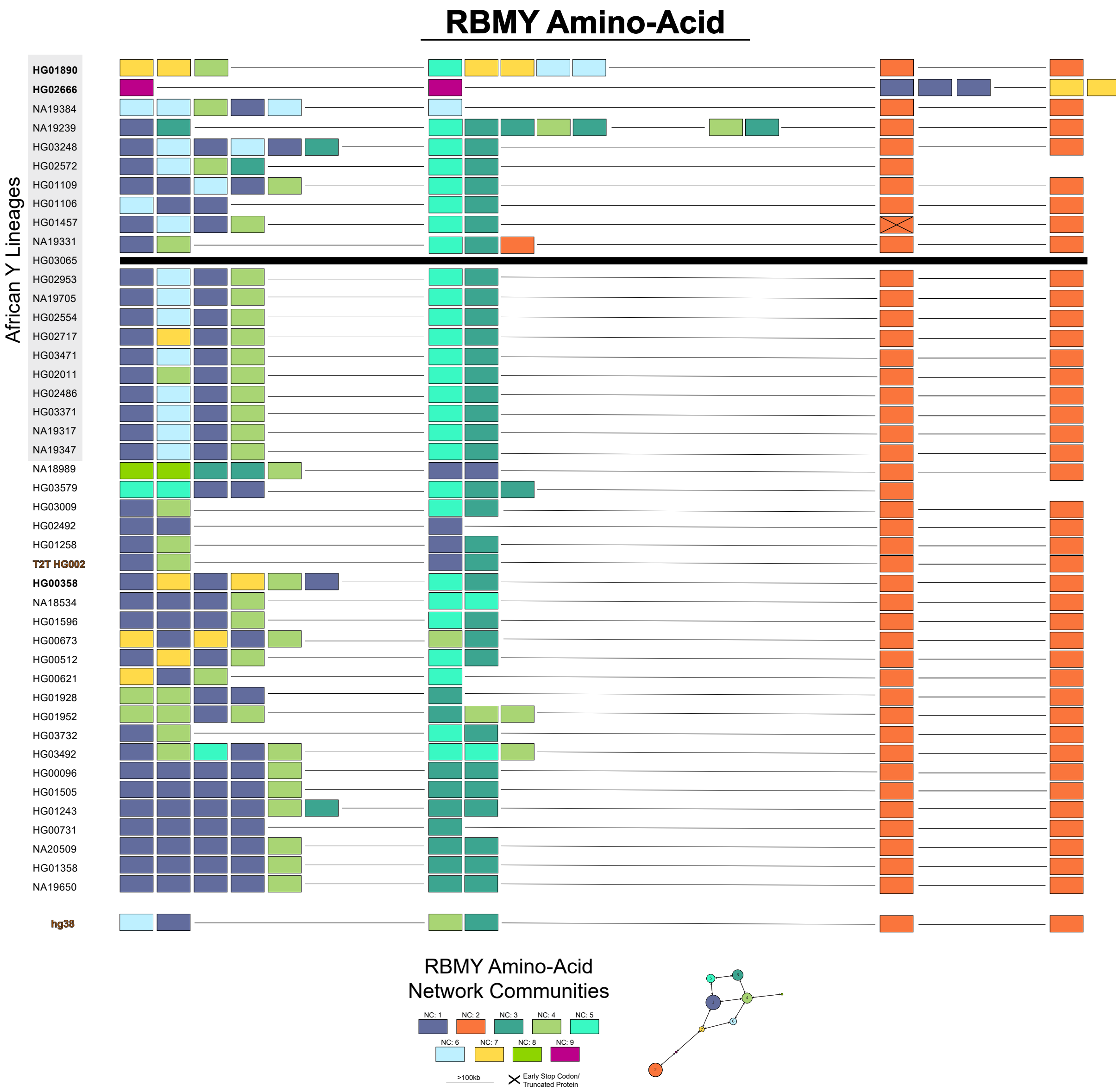


**Figure S46.** Network Analysis and Visualization of *RBMY1* Amino Acid Sequences. This figure shows the *RBMY1* gene copies and regions within each Y chromosome assembly that were analysed (42 + T2T + GRCh38). Individual *RBMY1* gene copies are shown as rectangles. The colour of an individual *RBMY* copy is based on the network community (NC) assignment (note: these colours are unique compared to the nucleotide network analyses, see **Methods**). A black line between gene copies indicates that they are no less than 100 kbp away from one another. Additionally, a secondary directed network was created from the translated network community consensus sequences (shown near the legend). The size of a node is representative of the total *RBMY1* gene copies that were assigned to the network community. An edge going from one node to another represents that the second node was the first's best match (i.e., most similar sequence; ties are allowed and shown as multiple edges on a node). The width of the edge represents how similar two nodes where a thicker edge indicates a more similar amino acid sequence. Overall, this visualization shows that some of the nucleotide network communities (previously shown **Figure S45**) contained sequence variants that were located in the untranslated regions resulting in less translated sequence variation. RBMY1F/J (orange rectangles; NC: 2) are maintained across all assemblies except for HG02666 which contains a recombinant version in regions 1 and 2 (purple rectangles; NC: 9). The most proximal *RBMY1* copy within region 1 of GRCh38 shows the same amino acid sequence as the African lineage specific RBMY1B. The amino acid sequence of one copy of RBMY1F/J within region 3 of HG01457 codes for a premature stop codon.

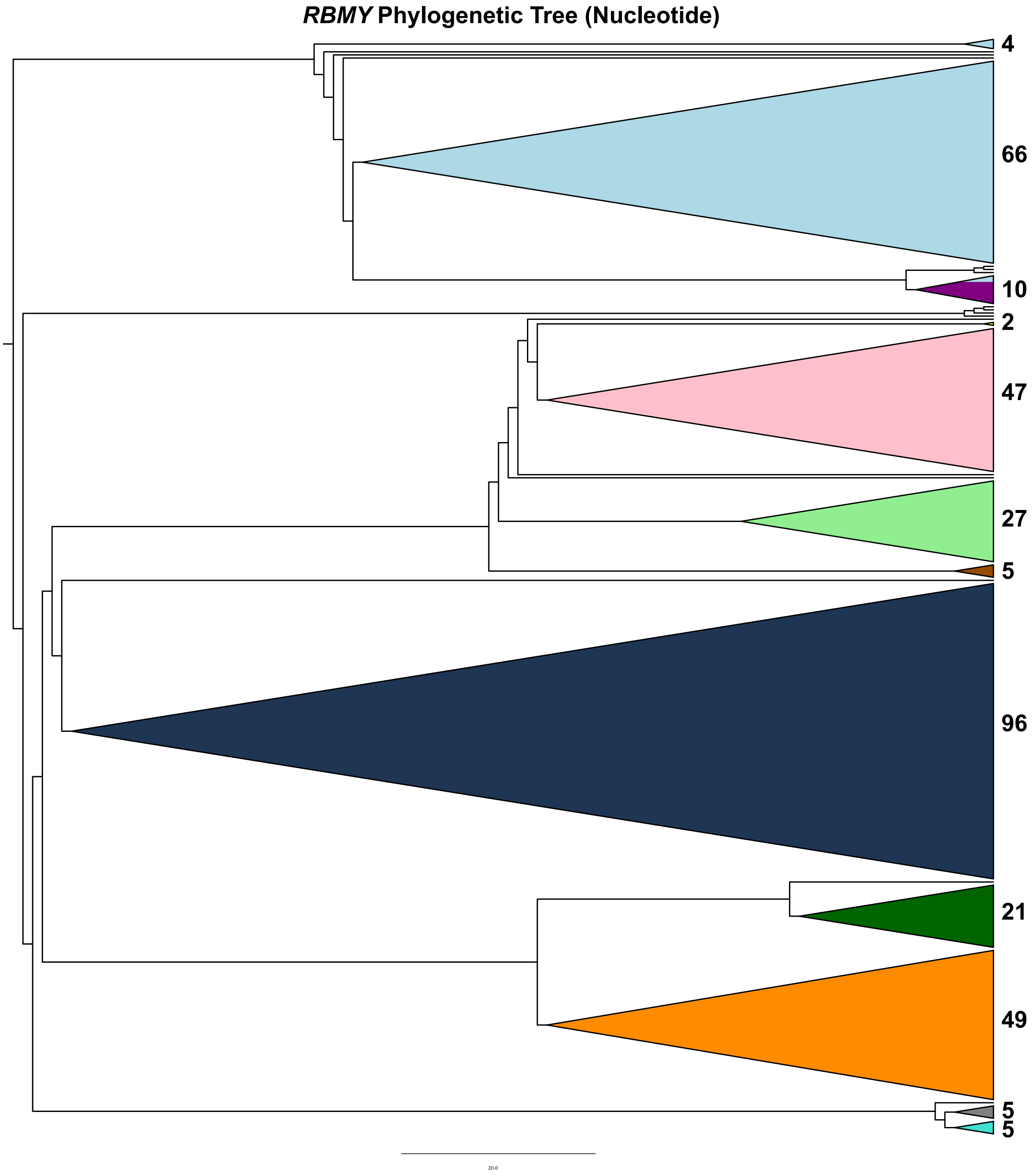


**Figure S47.** Phylogenetic tree of the *RMBY1* genes. Shown is the unrooted consensus tree of *RBMY1* genes constructed from 1000 bootstrap replicate trees (see **Methods**). Briefly, the exonic nucleotide sequences of all *RBMY1* genes were retrieved and 'fused' together to form individual gene copy exon sequences. The sequences were aligned using Muscle and a phylogenetic tree was reconstructed using a maximum likelihood approach. This tree is rooted at the midpoint, and the total count of *RBMY1* copies is shown on the right. *RBMY1* colours come from the network community assignments seen in **Figure S45**. On the Y chromosome *RBMY1* copies are primarily located within four regions. Results show *RBMY1* copies located in regions 1 and 2 (primarily dark blue, orange, dark/light green, and pink) clearly branch off from those located downstream in regions 3 and 4 (primarily light blue and purple copies).

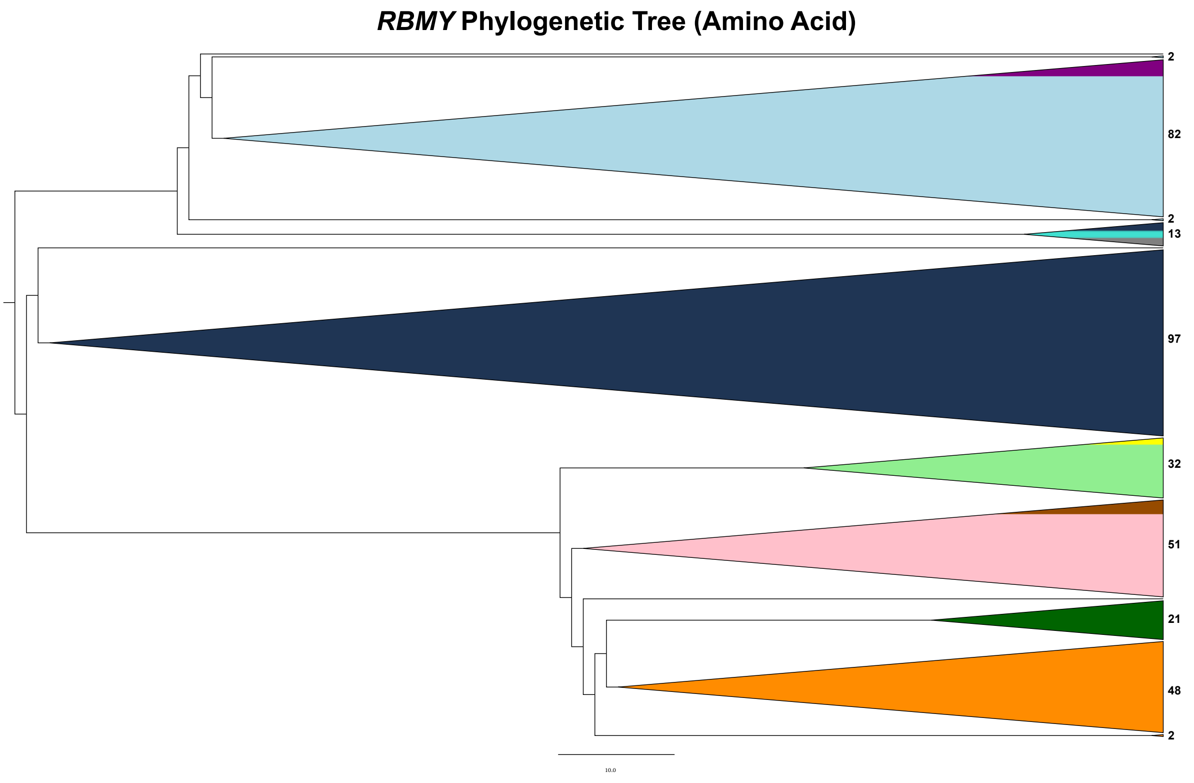


**Figure S48.** *RBMY1* Amino Acid Phylogenetic Tree Analysis. Shown is the unrooted consensus tree of translated *RBMY1* sequences (amino acids) constructed from 1000 bootstrap replicate trees (see **Methods**). Briefly, the exon sequences of all *RBMY1* genes were retrieved and 'fused' together to form individual gene copy exon sequences which were subsequently translated (using the same reading frame to produce the canonical protein sequence found in Ensembl). The translated sequences were aligned using Muscle and a phylogenetic tree was reconstructed using a maximum likelihood approach. This tree is rooted at the midpoint. *RBMY* copies are coloured based on the network clusters from the previous nucleotide analyses (see **Figure S45**). Counts of total *RBMY1* gene copies are shown on the right. The tree shows the amino acid sequence of RBMY1F/J (light blue/purple) diverges from the other *RBMY1* gene copies early in the tree. Functional analysis (see Results), suggests that all translated *RBMY1* sequences contain the same functional domains, but RBMY1F/J contain three amino acid substitutions one of which exists within the RNA-binding motif (R116S).

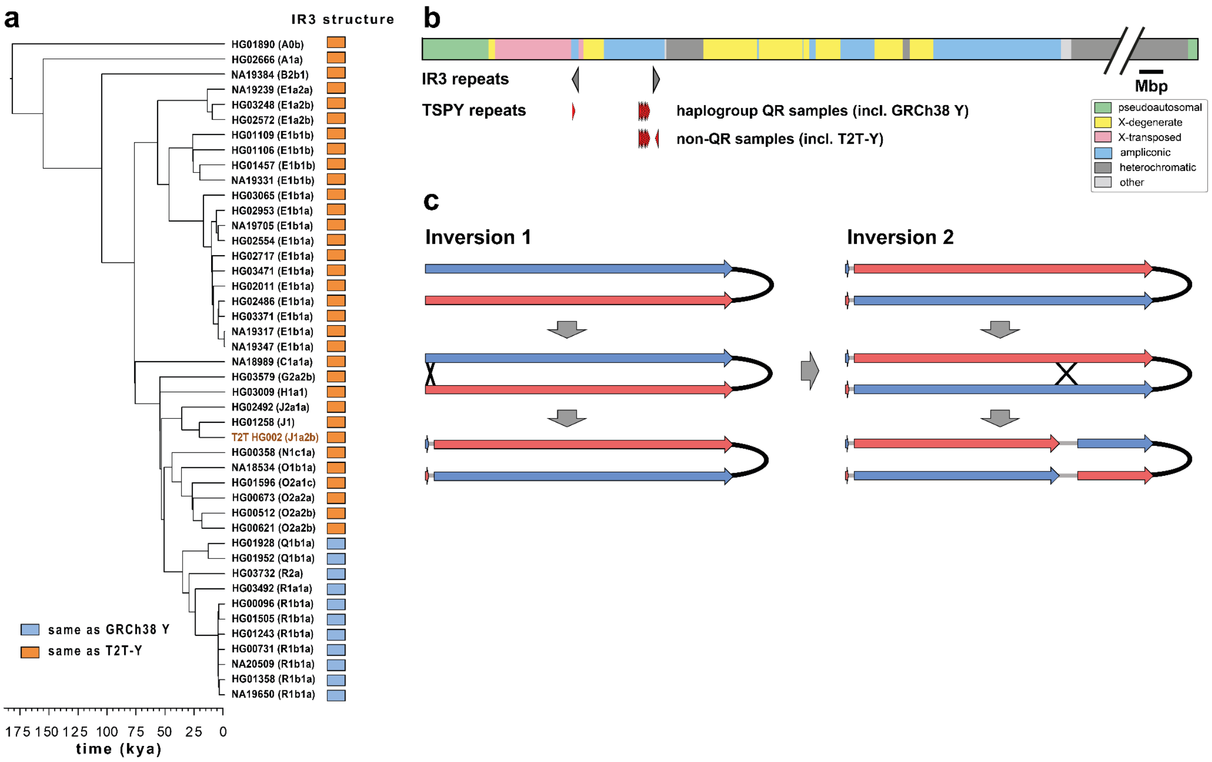


**Figure S49.** Phylogenetic distribution of different IR3 repeat compositions and the responsible IR3 inversion. **a.** Distribution of two different IR3 repeat compositions in the Y-chromosomal phylogeny. In orange - samples containing a single TSPY repeat in the proximal IR3 repeat in inverted orientation, in blue - samples containing a single TSPY repeat in the distal IR3 repeat in direct orientation. **b.** Schematic representation of IR3 composition and approximate locations of TSPY repeats relative to the Y chromosome structure. **c.** Identified inversions in phylogenetically related QR haplogroup samples - one changing the location and orientation of the single TSPY repeat from proximal to distal IR3 repeat, and another reversing the orientation of the region in between IR3 repeats. The inversions are indicated by black crosses. Blue and red arrows indicate distal and proximal IR3 repeats, respectively.

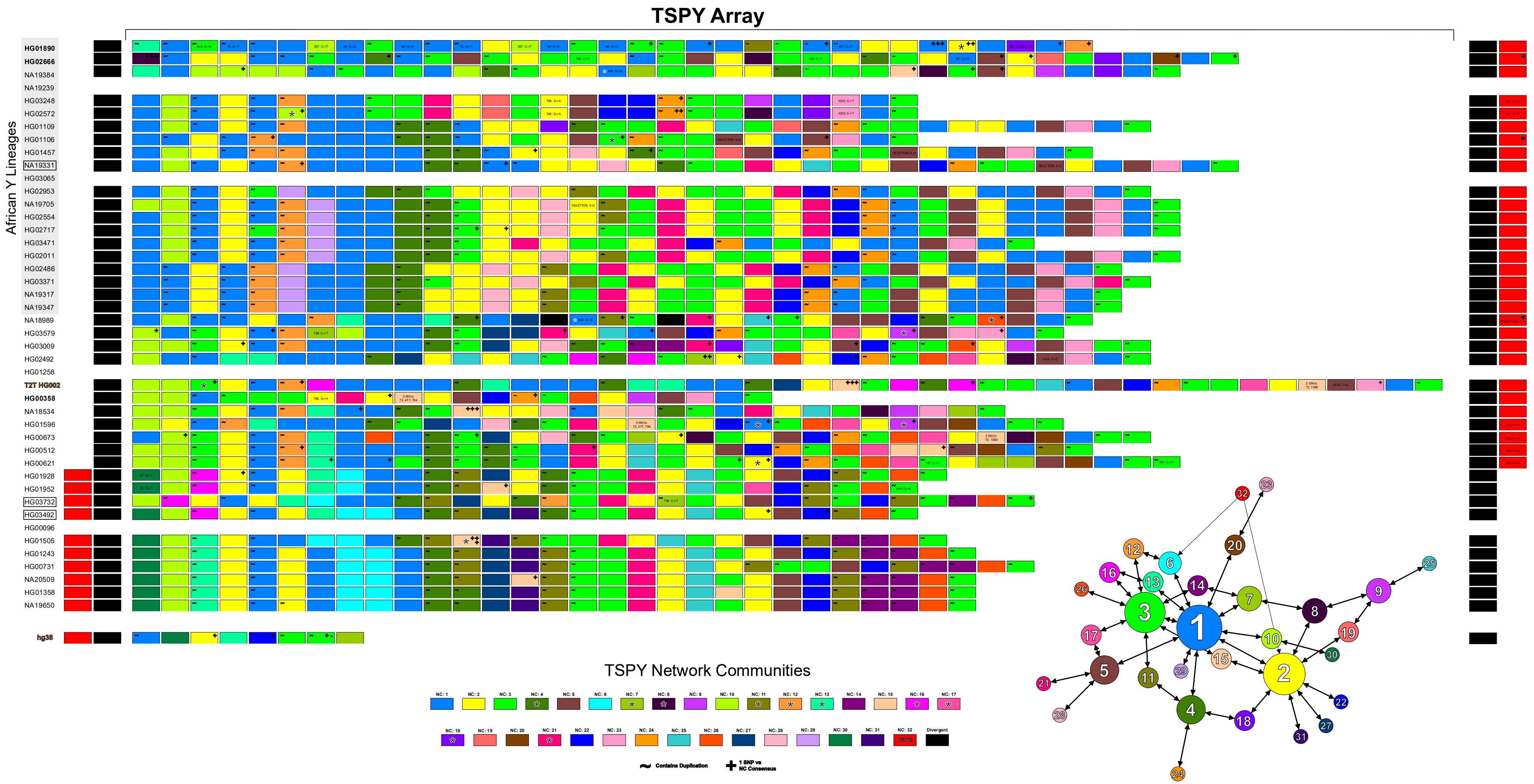

TSPY Array
African Y Lineages
HG01890
HG02666
NA19384
NA19239
HG03248
HG02572
HG01109
HG01106
HG01457
NA19331
HG03065
HG02953
NA19705
HG02554
HG02717
HG03471
HG02011
HG02486
HG03371
NA19317
NA19347
NA18989
HG03579
HG03009
HG02492
HG01258
T2T HG002
HG00358
NA18534
HG01596
HG00673
HG00512
HG00621
HG01928
HG01952
HG03732
HG03492
HG00096
HG01505
HG01243
HG00731
NA20509
HG01358
NA19650
hg38
TSPY Network Communities
Contains Duplication
1 SNP vs NC Consensus
Divergent

**Figure S50.** Network Analysis and Visualization of *TSPY* gene copies. The figure illustrates the TSPY array of each contiguous Y chromosome assembly in this region. Individual *TSPY* gene copies are shown as rectangles. The colour of each individual *TSPY* gene copy is based on the network community (NC) assignment (see **Methods**). A black rectangle around the sample name indicates that the TSPY array is in antisense orientation due to the sample carrying the *IR3/IR3* inversion. These arrays were re-oriented to better visualize the evolution of the array. Stars indicate possible gene conversion (GC) or recombination (R) events. If an entire community is likely the result of a GC/R event then a star was placed on the NC legend rectangle. Also shown is the *TSPY2* gene copy (red rectangle) that is located outside of the TSPY array. Additionally, a secondary directed network created from the consensus sequence of NCs is shown. The size of a node is representative of the total number of *TSPY* gene sequences that were assigned to the community. An edge going from one node to another represents that the two clusters are the most similar to each other (ties are allowed and shown as multiple edges leading from a node). The edge width represents the similarity between two nodes where a thicker edge indicates less SNVs. Overall, our results suggest that the medium blue NC (i.e., Network Community 1) is representative of the ancestral *TSPY* gene sequence. Expansions, contractions, gene conversions, recombinations, and deletions are widely seen throughout the TSPY array.

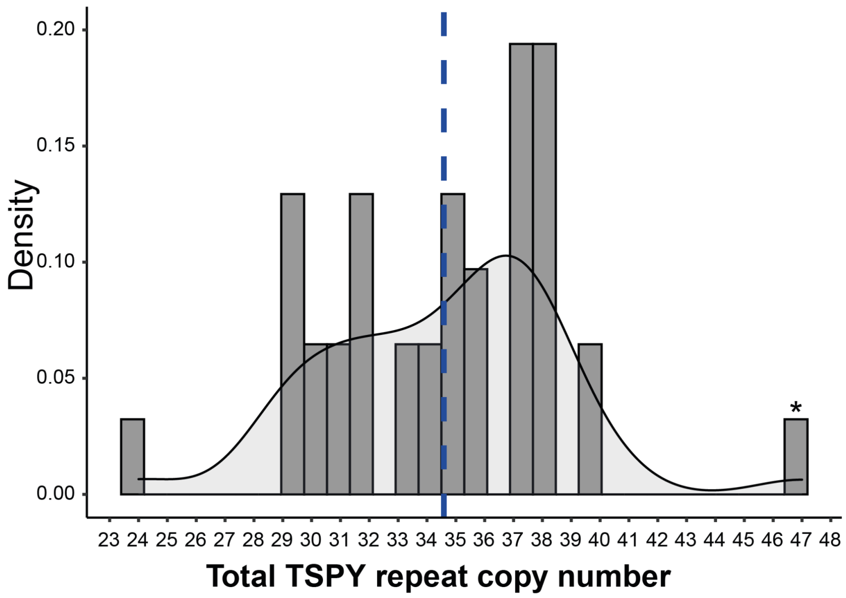


**Figure S51**. The total copy number distribution of the TSPY repeat units across 39 samples (T2T Y is marked with an asterisk).

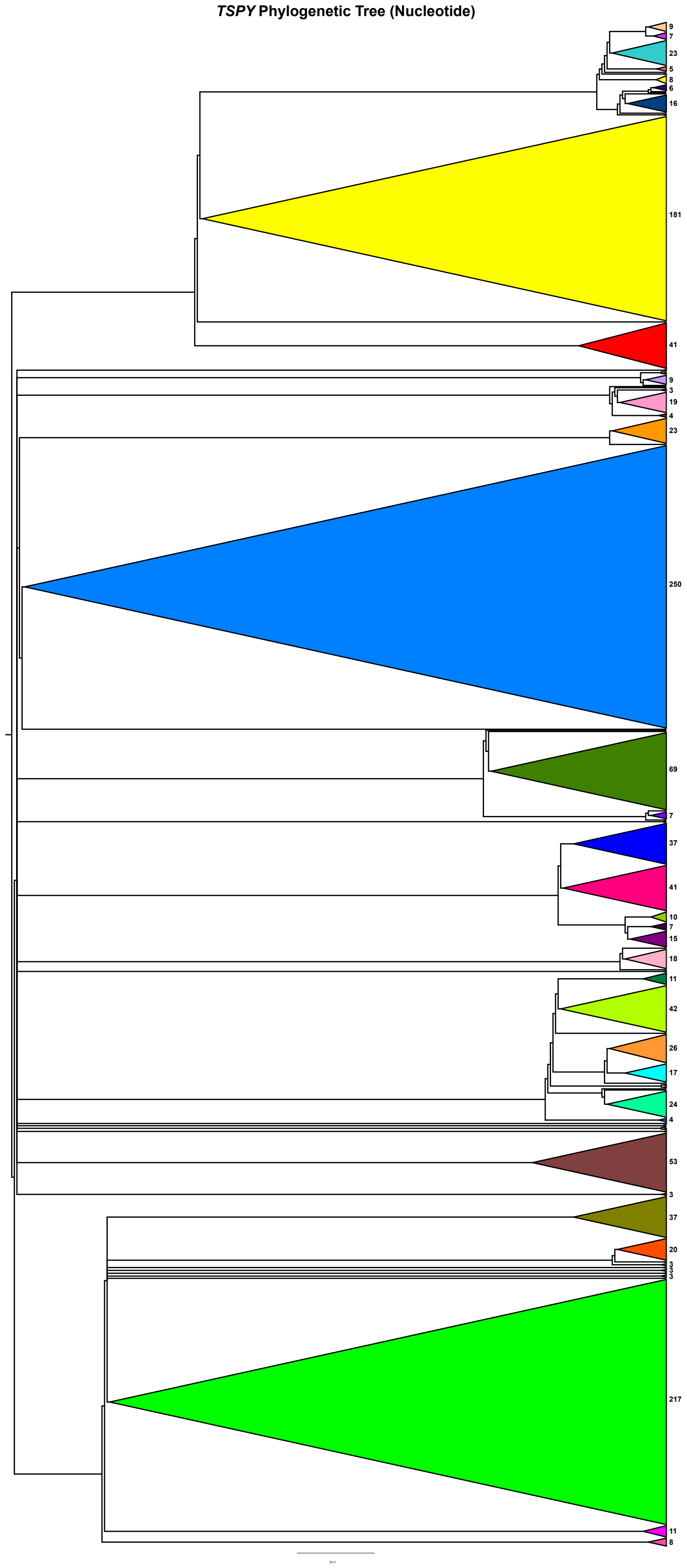


**Figure S52.** *TSPY* Phylogenetic Tree Analysis of nucleotide sequences. Shown is the consensus tree of *TSPY* genes constructed from 1000 bootstrap replicate trees and visualized using FigTree (see **Methods**). Briefly, the exonic nucleotide sequences of all *TSPY* genes were retrieved and 'fused' together to form individual gene copy exon sequences. The sequences were aligned using Muscle and a phylogenetic tree was reconstructed using a maximum likelihood approach. The consensus tree was rooted at the midpoint and the total *TSPY* copy number is shown on the right. TSPY colours come from the network community assignments seen in **Fig. S50.** The early split/rise of the medium blue cluster (n=250 *TSPY* copies, also known as network cluster 1 in **Fig. S50**) within the tree in conjunction with our manual comparison of *TSPY* sequences, as well as their presence across all lineages suggests that the medium blue *TSPY* copies are the most ancestral.

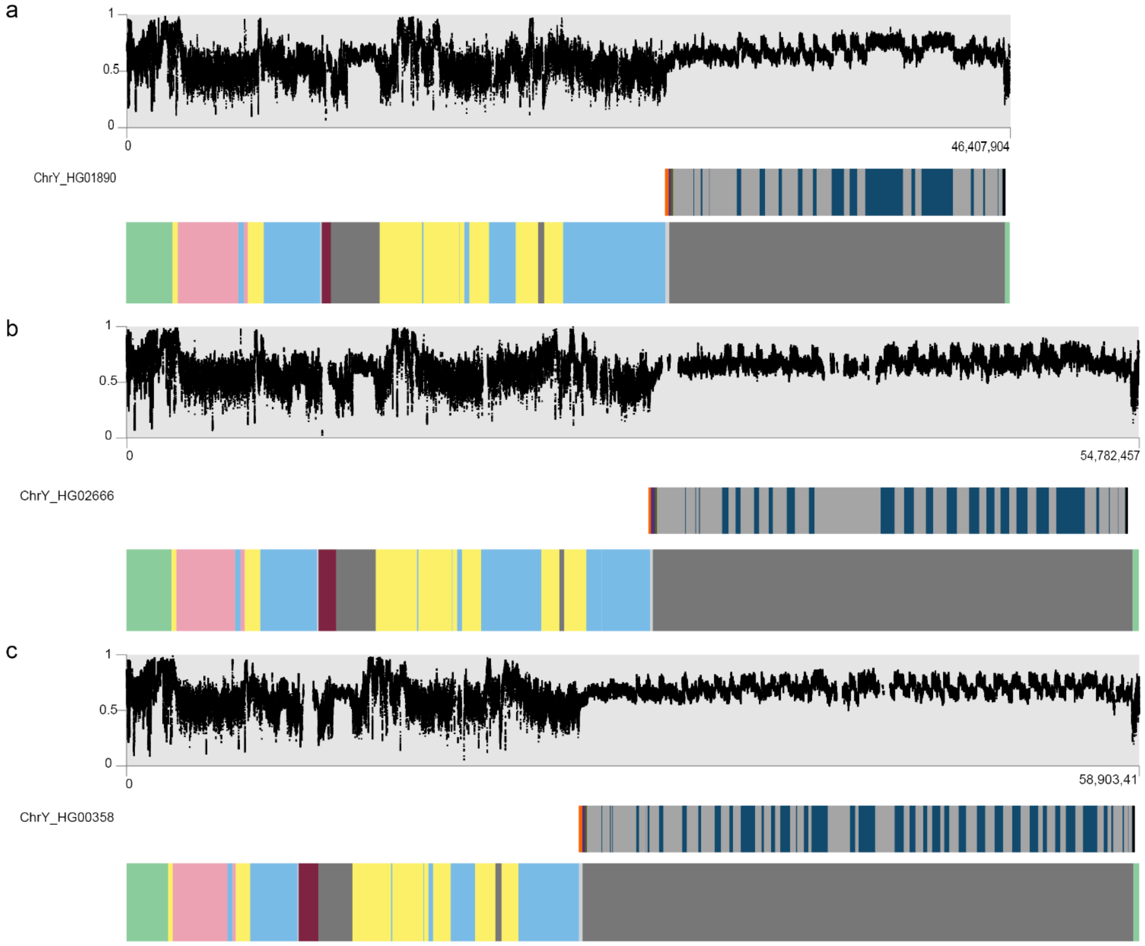


**Figure S53.** Methylation patterns as determined from the ONT data across the three contiguously assembled Y chromosomes, with repeat array locations (orange - *DYZ18*, purple - 2.7kb-repeat, green - 3.1kb-repeat, grey - *DYZ1*, blue - *DYZ2*) and Y-chromosomal subregion locations (see **Fig. 1a** for details) shown below as bar plots.

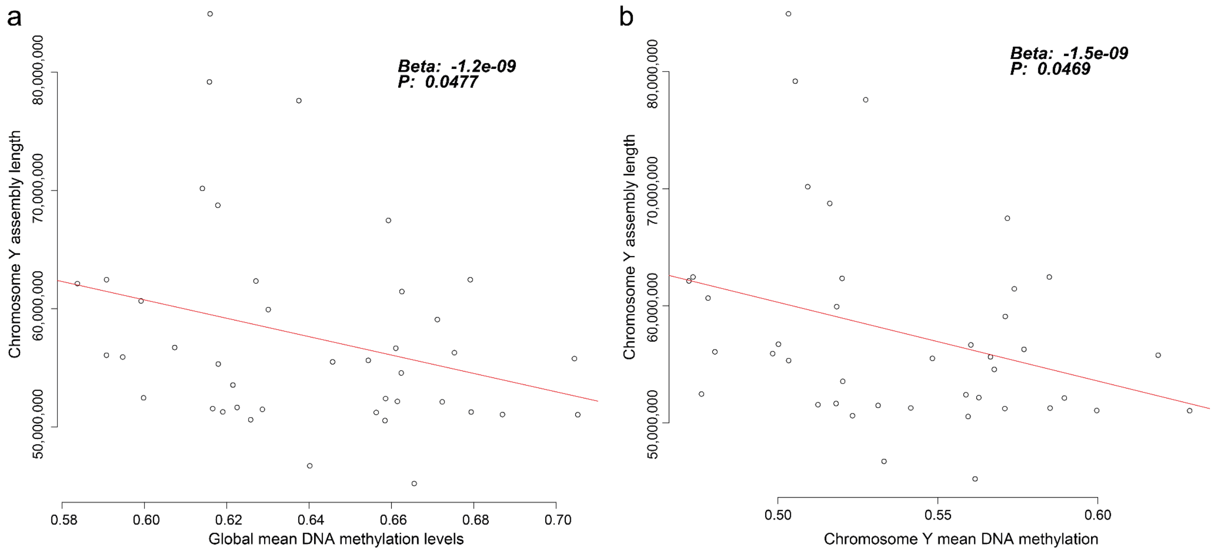


**Figure S54.** The relation between chromosome Y assembly length and DNAme levels. Shown are dot plots showing the relation between DNAme and chr Y assembly length, including the effect size (beta) and significance derived from a linear model (see Methods). **a.** DNAme levels on the autosomes versus chromosome Y assembly length. **b.** DNAme levels on chromosome Y versus chromosome Y assembly length.

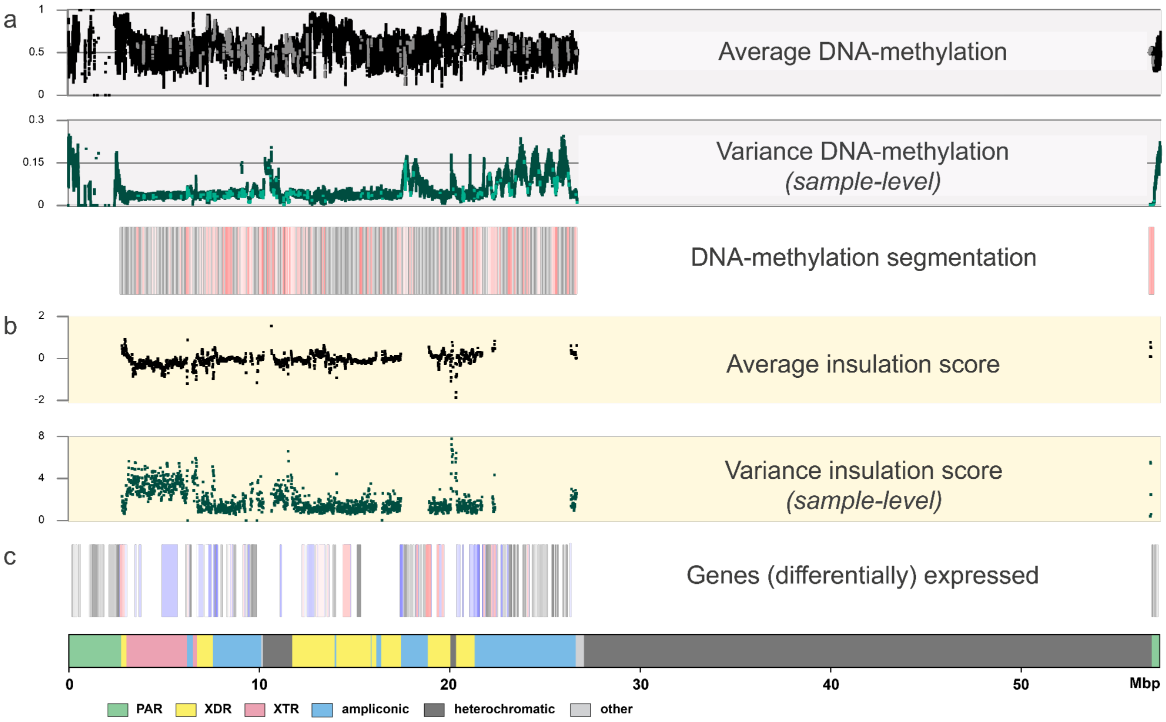


**Figure S55.** Functional analyses on the Y chromosome with DNA-methylation, RNA expression and HiC information as anchored to GRCh38. **a.** The top three panels show DNA-methylation levels and variation over the studied chromosomes. In black (top line) the average methylation is shown, in green (middle) the variation in DNA-methylation levels across the studied genomes. The bottom boxes represent the DNA methylation segmentation using PycoMeth-seg. In grey shades 2,861 methylation segments, and in red shades the 340 significantly differentially methylated segments (DMS). The CpG sites that fall in a DMS are coloured in a lighter shade in the top two panels. **b.** Two panels showing average insulation scores (top) and variance of insulations scores between any two samples (bottom) across 40 samples with Hi-C data with 10 kbp resolution. Regions with lower insulation scores are more insulated and more likely to be TAD boundaries, while regions with higher scores are most likely to stay inside TADs. **c.** The Geuvadis based gene-expression analysis, shown are the 205 genes on Y chromosome (grey shades), the 64 genes expressed in the Geuvadis LCLs (blue shades), of which 22 are differentially expressed (red shades).

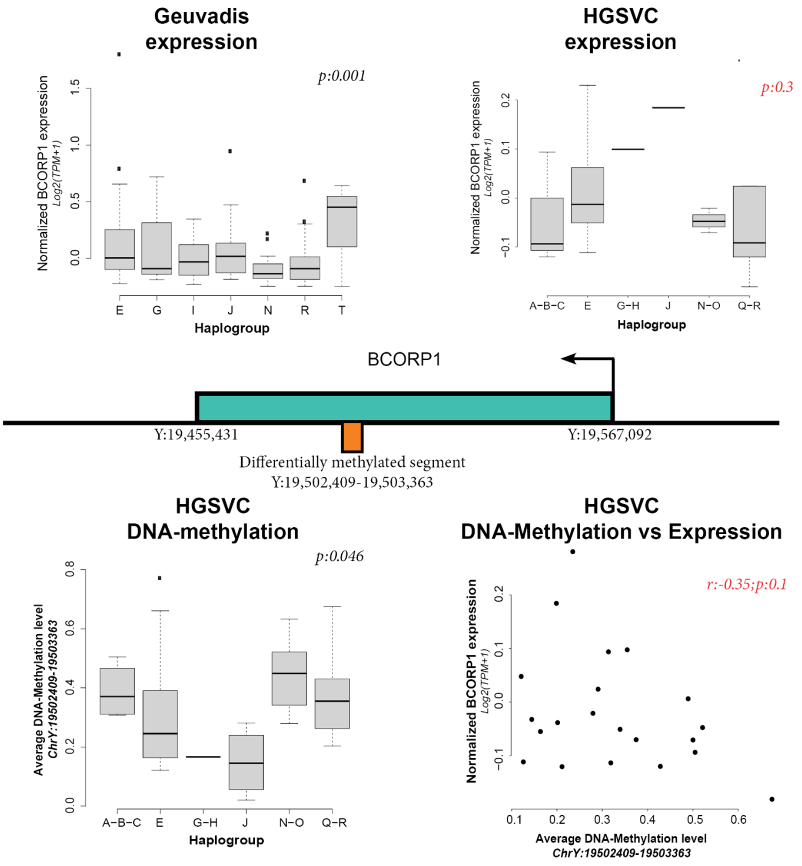


**Figure S56**. Schematic representation of the *BCORP1* gene and the effects of the haplogroup on gene expression and DNA-methylation (DNAme) levels. In the center an illustration of the *BCORP1* (in green), in the center of the gene a differentially DNAme segment is identified (in orange). The differential DNAme effect identified in the HGSVC samples is plotted in the bottom left boxplot. The *BCORP1* is found to be differentially expressed in the Geuvadis samples (top left boxplot). The expression effect is suggestive in the 21 HGSVC samples, expression of haplogroup E is on average higher than haplogroups G,H,J,N,O,Q,R. The expression effect of the haplogroup is inversely related to the DNA-methylation effect in this segment (Pearson R -0.35), with a suggestive P value of 0.1 indicating the relation in this small sample set.

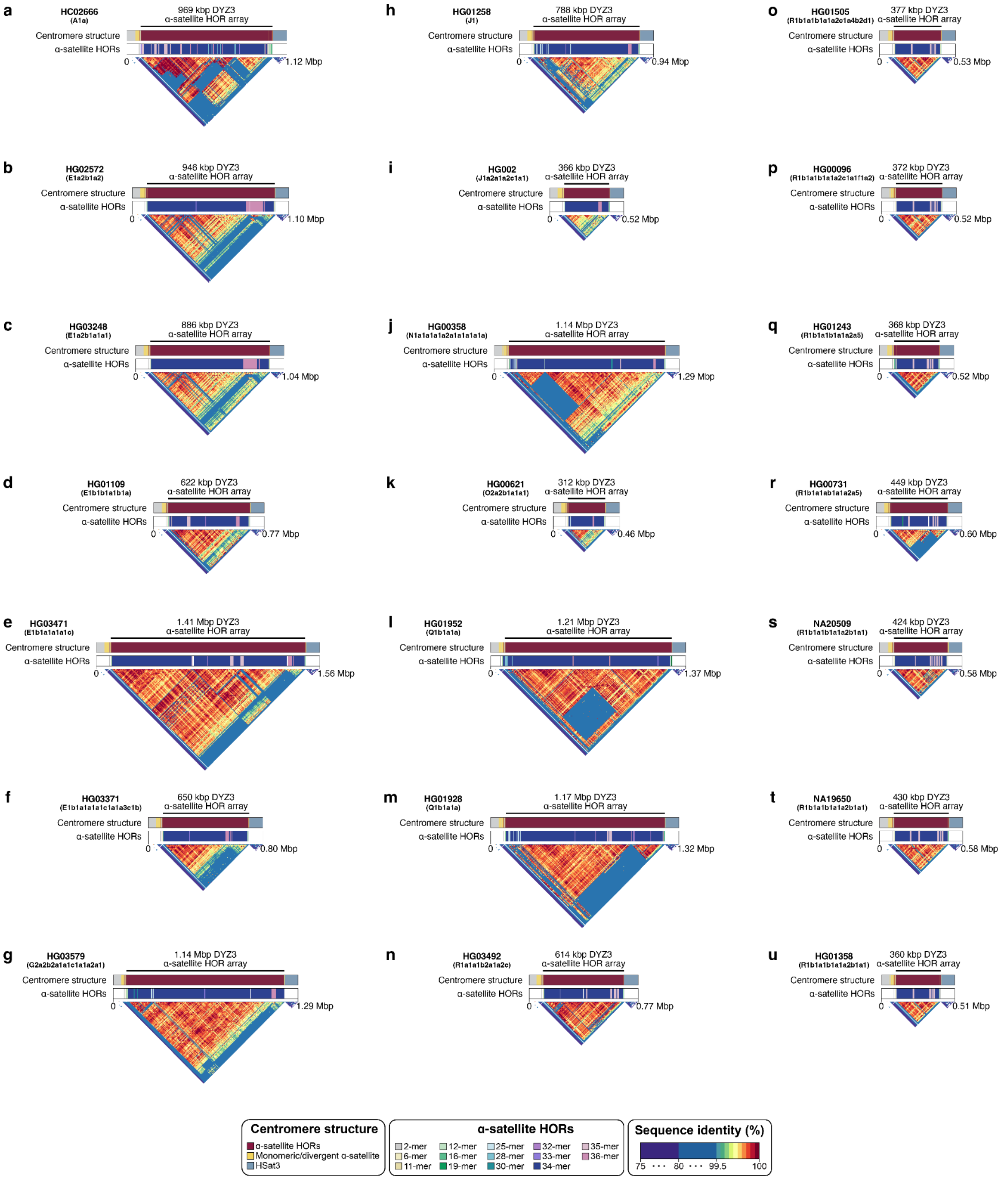


**Figure S57.** Organization of the chromosome Y centromeric region from 21 genomes representing all major superpopulations. The structure (top), α-satellite HOR organization (middle), and sequence identity heat map (bottom) for each centromere is shown and reveals the presence of novel HORs in over half of the centromeres. Note - the sizes of the *DYZ3* α-satellite array are shown on top as determined using RepeatMasker (**Methods**). The centromeres are ordered phylogenetically from the deepest-rooting sample, panel **a** to **u**.

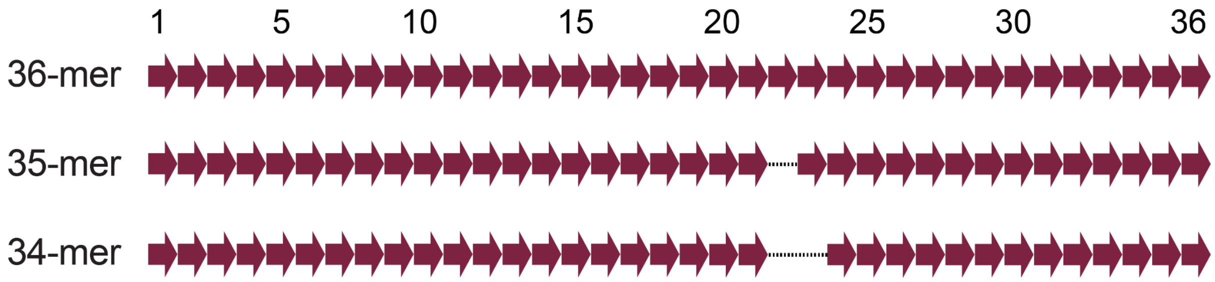

**Figure S58.** The 34-monomer α-satellite HOR was formed via two sequential steps in which a single α-satellite monomer residing at the 22nd position was deleted. The 34-monomer α-satellite HOR dominates all chromosome Y centromeres.

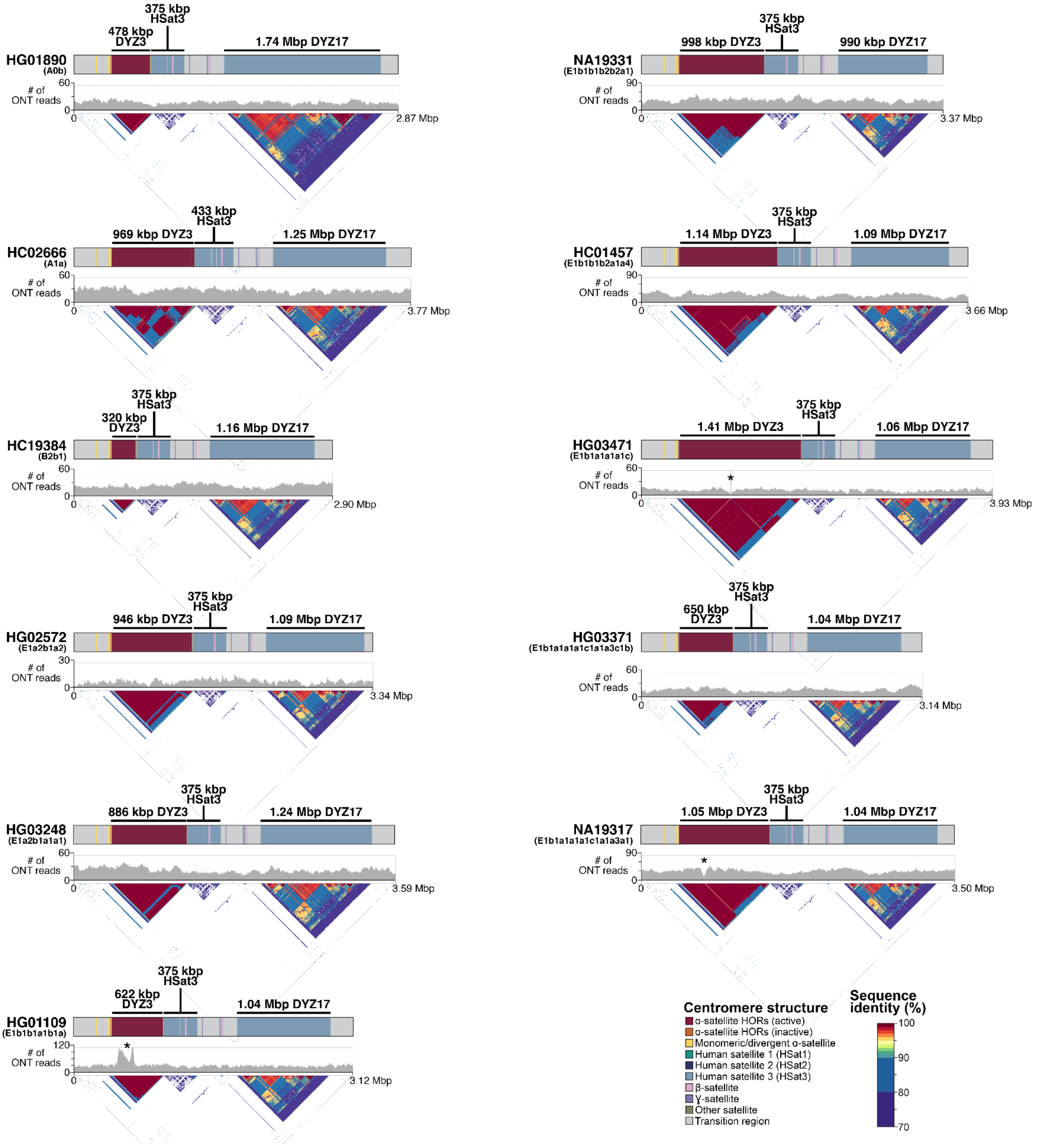


**Figure S59.** Genetic landscape of the Y-chromosomal pericentromeric region from samples carrying African Y lineages. The top panel shows locations and composition of the pericentromeric region with repeat array sizes shown for each Y chromosome (the *DYZ3* α-satellite array size as determined using RepeatMasker, **Methods**). The middle panel shows (UL-)ONT read depth and bottom sequence sequence identity head maps generated using StainedGlass pipeline (using 5-kb window size). Asterisks indicate two samples (HG01109 and NA19317) with a possible assembly collapse/error, and one sample (HG03471) with a single gap in the *DYZ3* array.

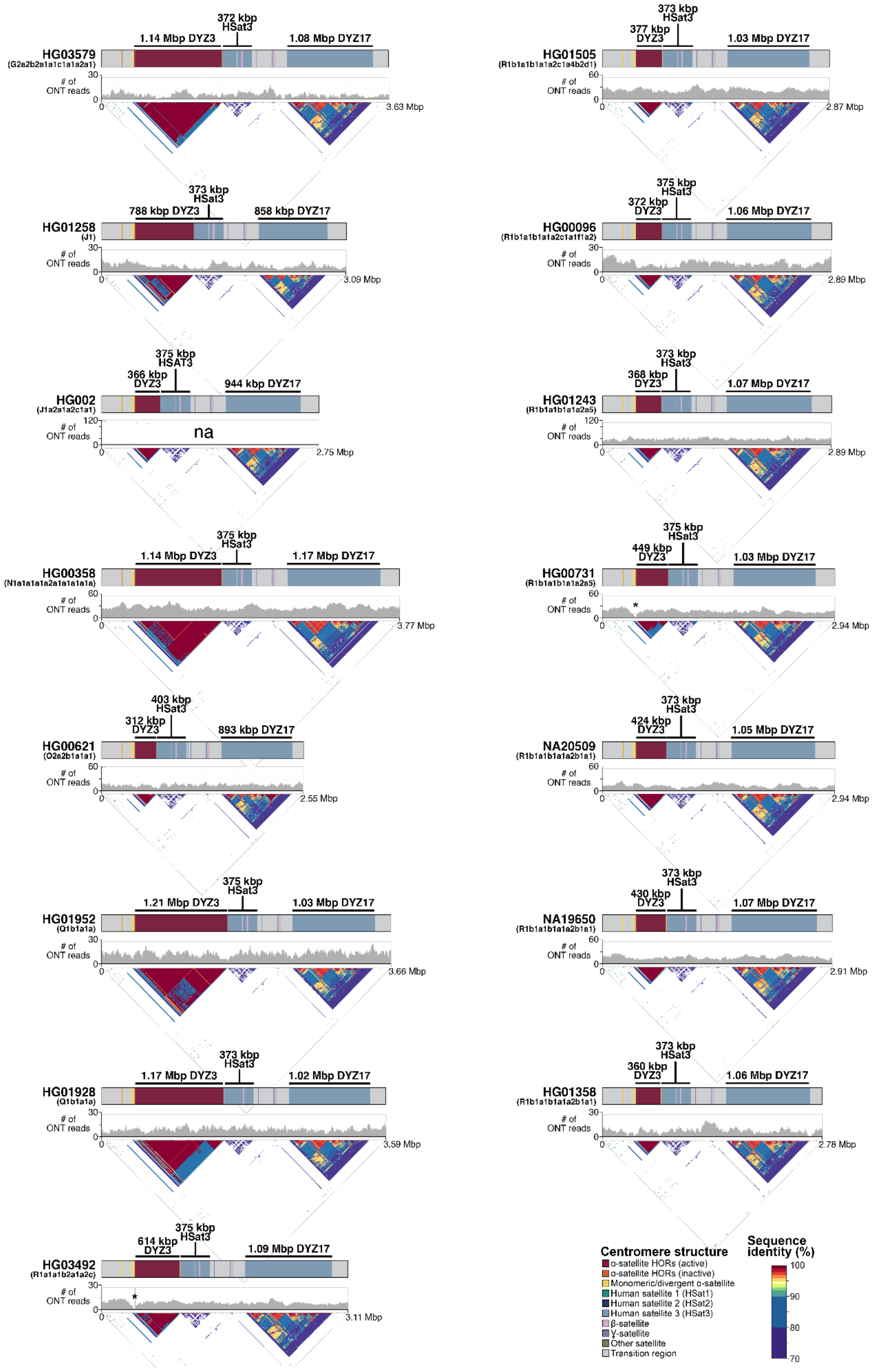


**Figure S60.** Genetic landscape of the Y-chromosomal pericentromeric region from samples carrying non-African Y lineages. The top panel shows locations and composition of the pericentromeric region with repeat array sizes shown for each Y chromosome (the *DYZ3* α-satellite array size as determined using RepeatMasker, **Methods**). The middle panel shows (UL-)ONT read depth and bottom sequence sequence identity head maps generated using StainedGlass pipeline (using 5-kb window size). Asterisks indicate two samples (HG03492 and HG00731) with a single gap in the *DYZ3* array. na - not available.

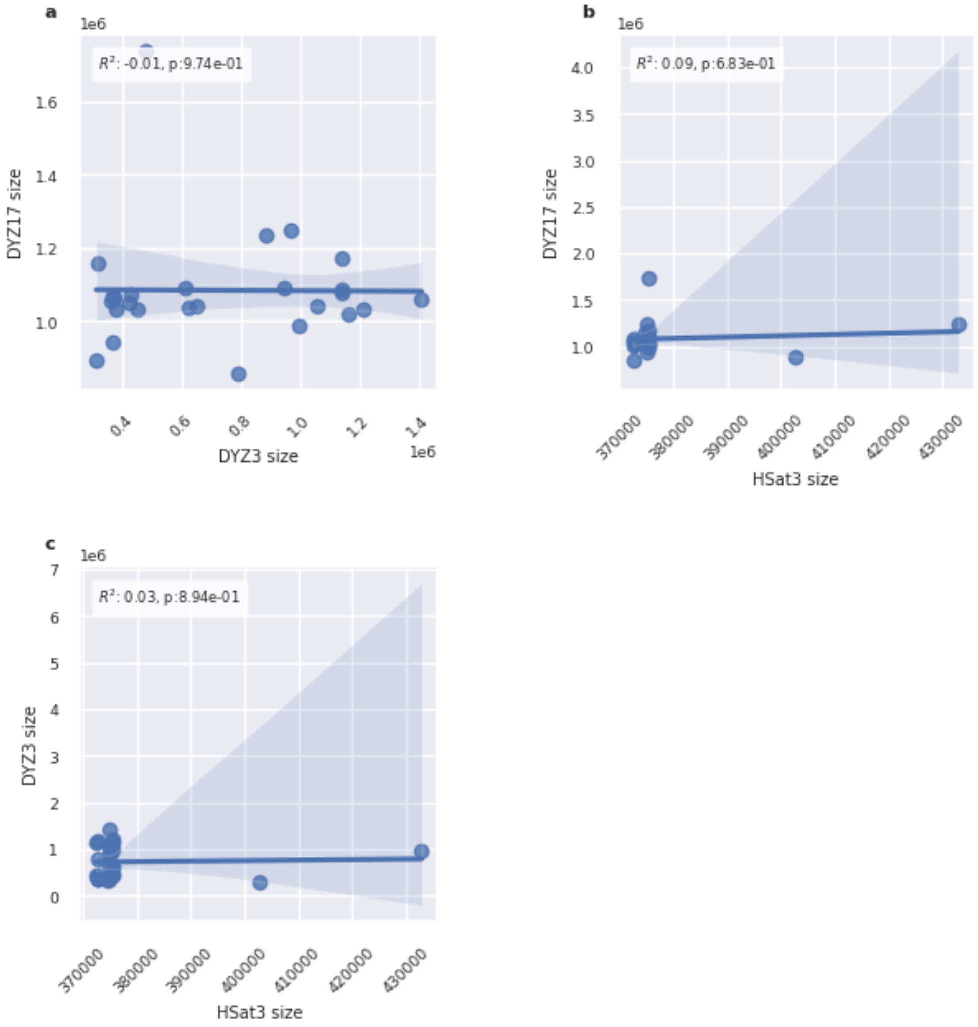


**Figure S61.** Regression plots between the sizes of (peri-)centromeric repeat arrays: *DYZ3* alpha-satellite array, Hsat3, and the *DYZ17* array. We report the correlation coefficient and the p-value on the upper-left corner box. No correlations attained a significant p-value.

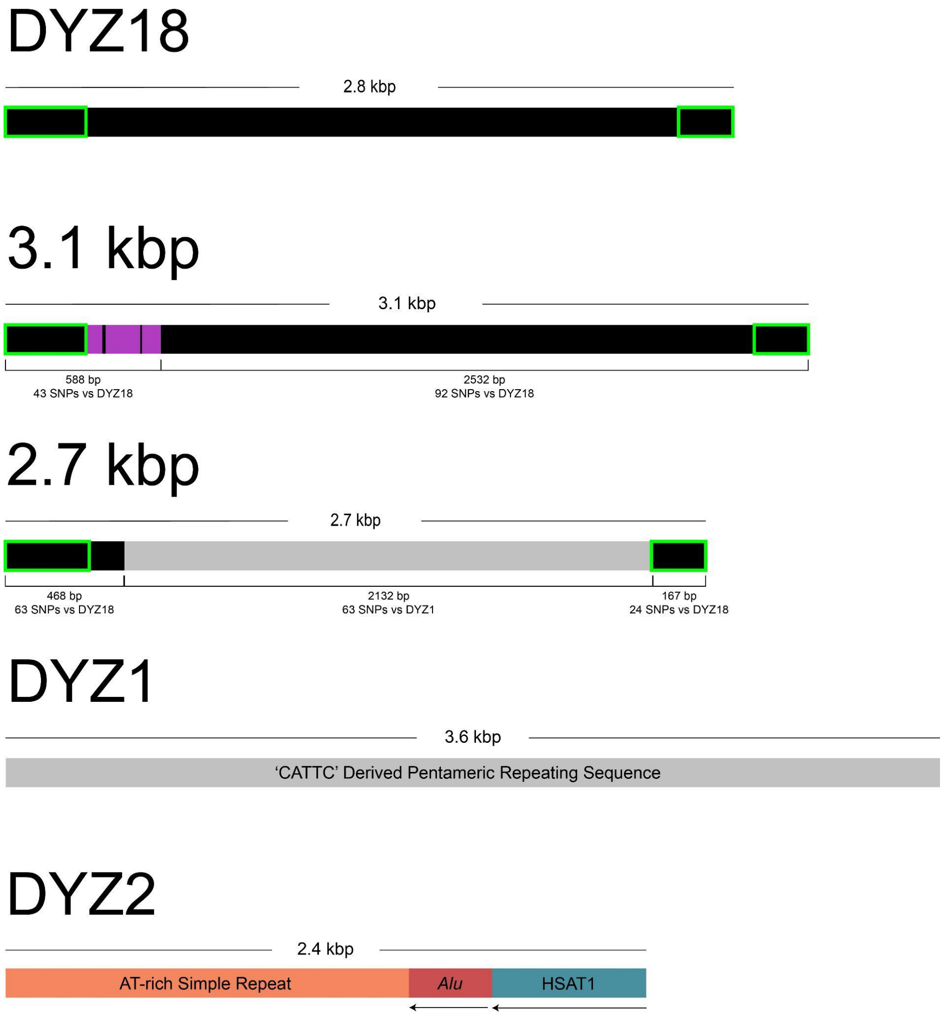


**Figure S62.** Composition and similarities of Yq12 heterochromatic repeat units. Green highlight - indicates regions with high sequence similarity to the *DYZ18* repeat unit. Light grey region in 2.7-kb repeat indicates a region of high sequence similarity to the *DYZ1* repeat unit. The purple region is a span of ~200-300 bases unique to the 3.1-kbp repeat.

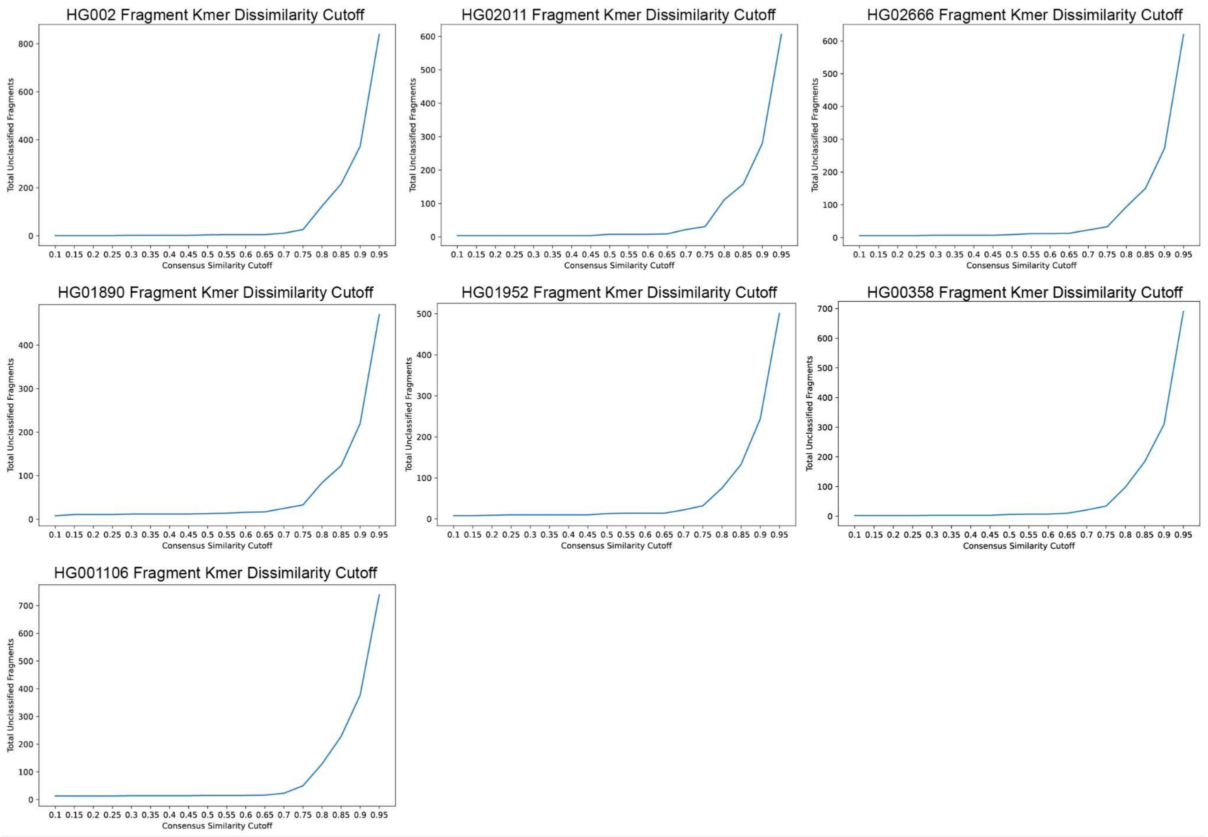


**Figure S63**. The line plots show the total HaeIII fragments (y-axis) that are unclassified at each k-mer abundance profile similarity cutoff (x-axis). Fragments were classified as either *DYZ18*, 3.1-kbp repeat, 2.7-kbp repeat, or *DYZ1* if their k-mer abundance profile was 75% or more similar. Each genome's plot exhibits an exponential growth in unclassified fragments above the 75% similarity cutoff.

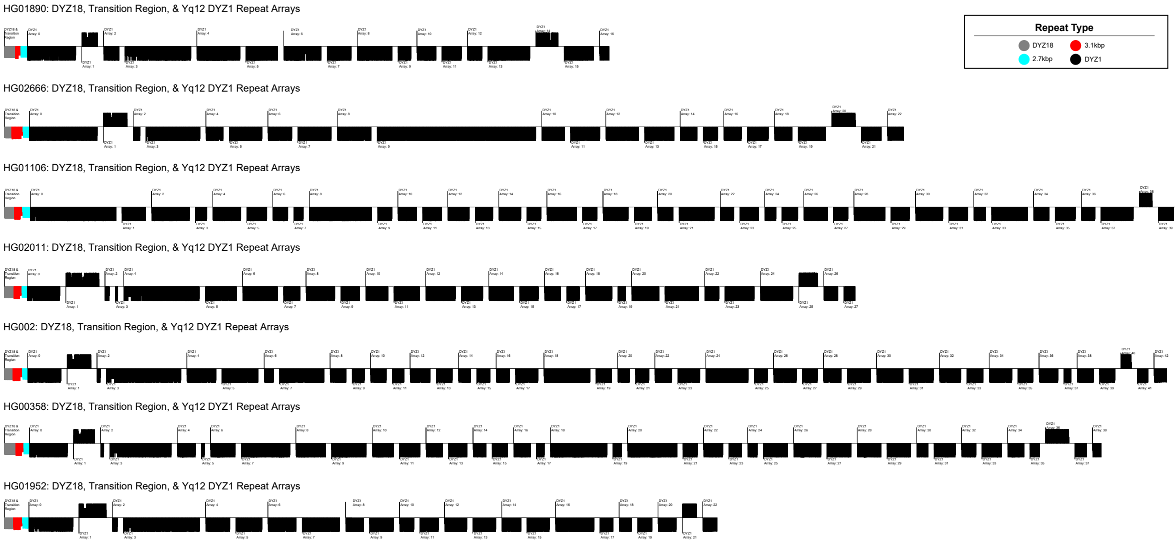


**Figure S64.** An overview of the *DYZ18* (gray), 3.1-kbp (red), 2.7-kbp (blue) and *DYZ1* (black) repeat arrays in the Yq11/transition region/Yq12 subregion within each of the seven samples with completely assembled Yq12 subregion. The length of individual lines is a function of the size of the repeat. The orientation (up = sense, down = antisense) was determined based on RepeatMasker annotations of satellite sequences within repeats.

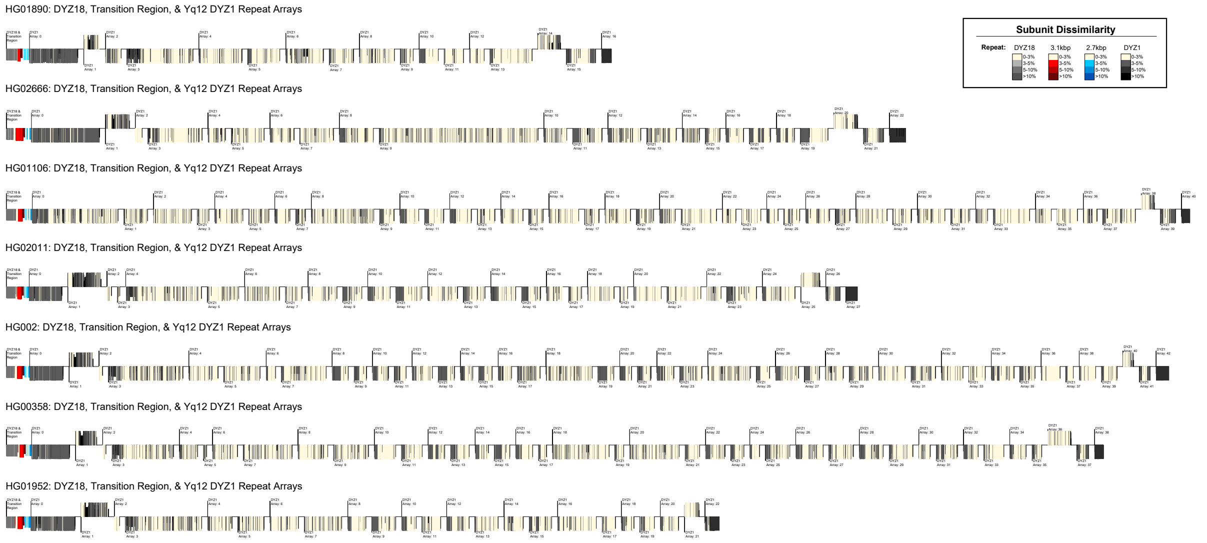


**Figure S65.** An overview of the Bray-Curtis distance/dissimilarity of k-mer abundance profiles for individual *DYZ18* (gray), 3.1-kbp (red), 2.7-kbp (blue) and *DYZ1* (black) repeats versus their consensus sequence. The Yq11/ transition region/Yq12 subregion are shown for each of the seven samples with a completely assembled Yq12 subregion. Lighter colors indicate less distance/dissimilarity (more similar) k-mer abundance profiles compared to their consensus sequence. Results indicate arrays located on the proximal and distal boundaries of the Yq12 region contain repeats with k-mer abundance compositions less similar to their consensus sequence (i.e., more diverged). The length of individual lines is a function of the size of the repeat. The orientation (up = sense, down = antisense) was determined based on RepeatMasker annotations of satellite sequences within repeats.

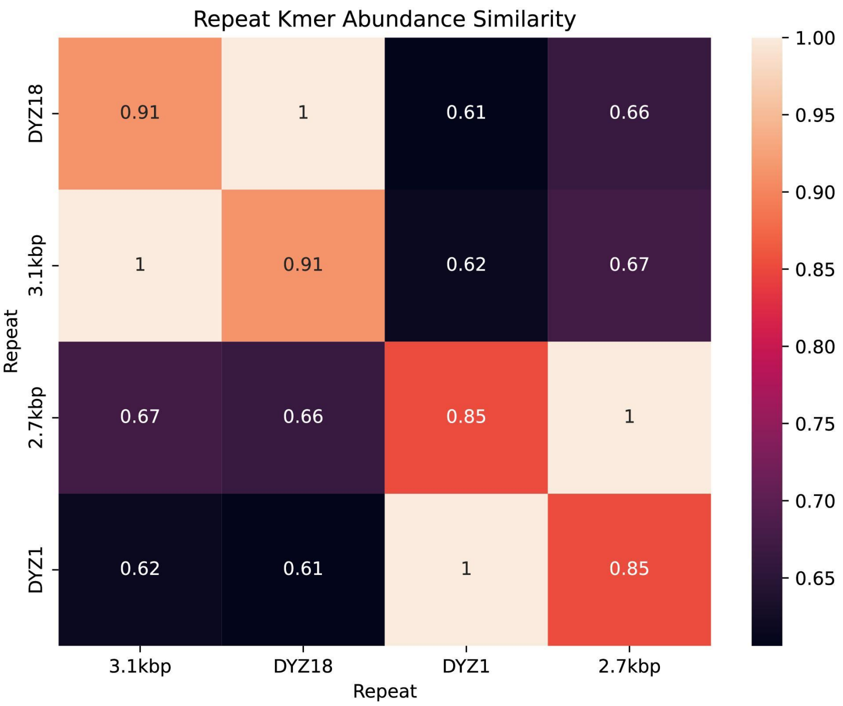


**Figure S66.** Heatmap of the complement of the Bray-Curtis distance/dissimilarity (i.e., 1-BC) between k-mer abundance profiles of *DYZ18*, 3.1-kbp, 2.7-kbp, and *DYZ1* consensus sequences is shown. The k-mer abundance profile of *DYZ1* was most similar to the 2.7-kbp repeat (85%), whereas the *DYZ18* and 3.1-kbp repeat sequences were more similar (91%) to each other.

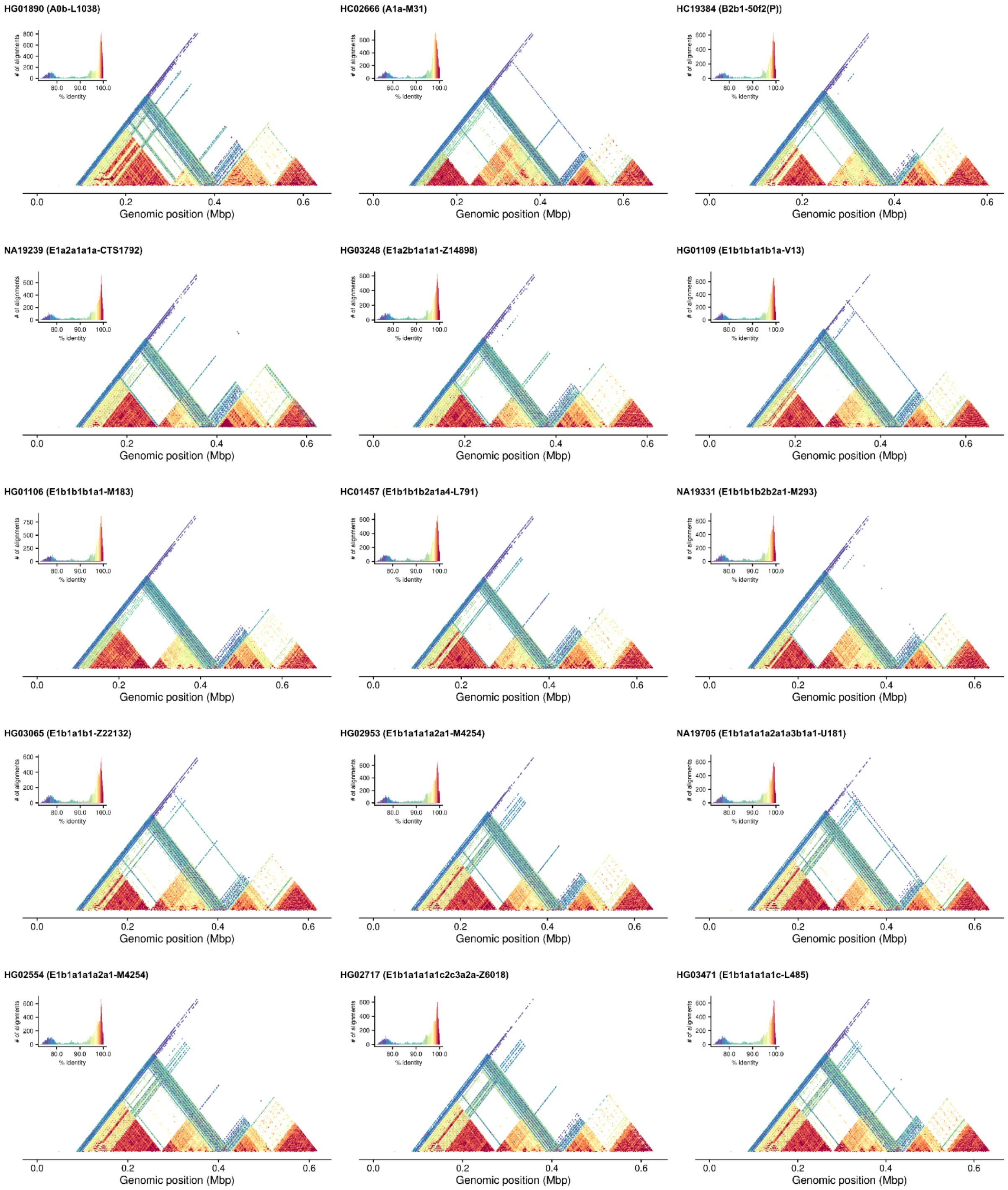


**Figure S67.** Sequence identity heat map of the Yq11/Yq12 transition region, including the *DYZ18*, 3.1-kbp, 2.7-kbp repeat arrays and 100 kbp of the first *DYZ1* repeat array generated using StainedGlass with 2 kbp window size. 100 kbp proximal to the *DYZ18* repeat array has also been included. Samples are ordered phylogenetically from the deepest-rooting sample (from left to right). The plot highlights higher sequence similarity between the *DYZ18* and 3.1-kbp repeat arrays, and between the 2.7-kbp and *DYZ1* repeat arrays, respectively.

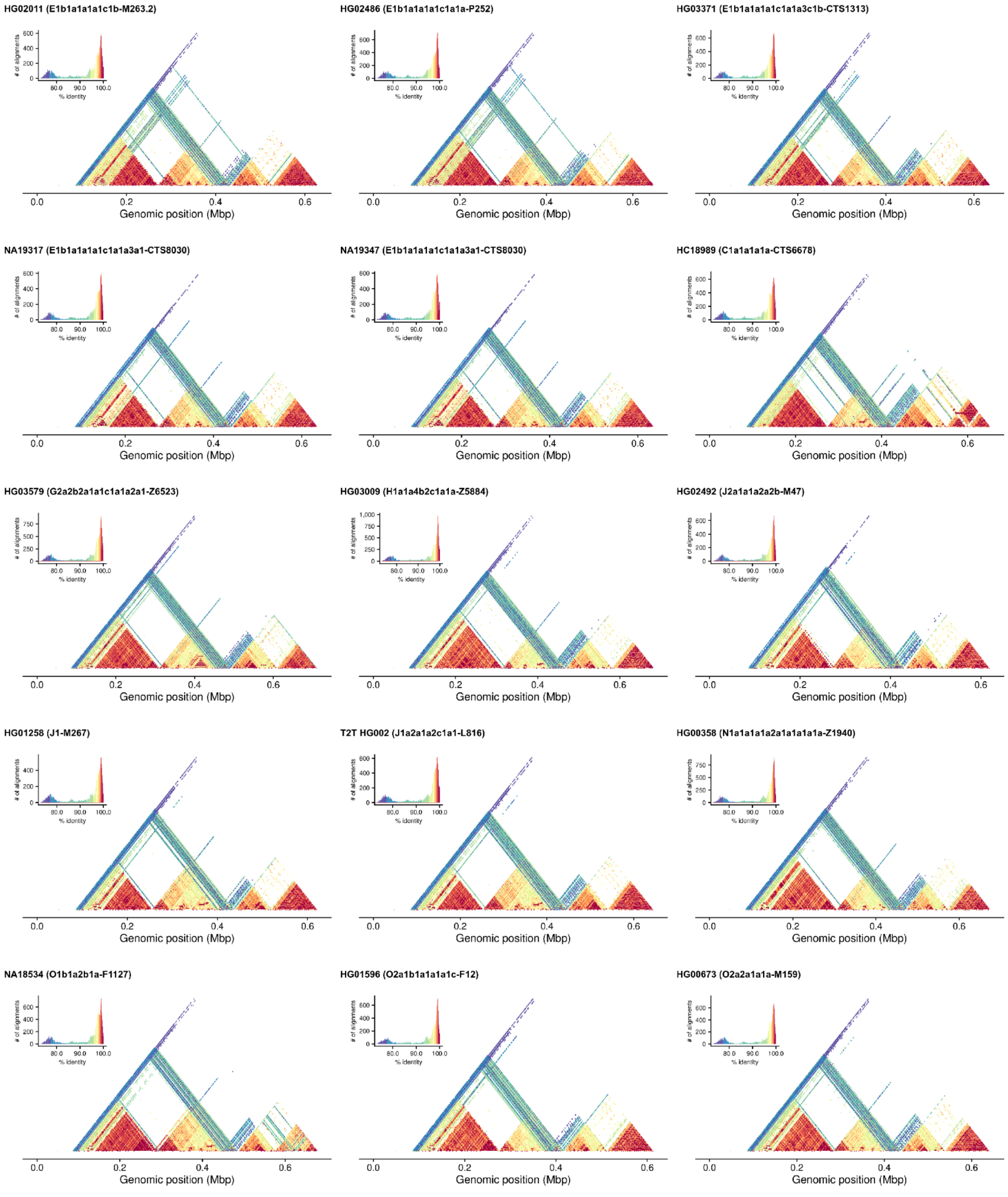


**Figure S68.** Sequence identity heat map of the Yq11/Yq12 transition region, including the *DYZ18*, 3.1-kbp, 2.7-kbp repeat arrays and 100 kbp of the first *DYZ1* repeat array generated using StainedGlass with 2 kbp window size. 100 kbp proximal to the *DYZ18* repeat array has also been included. Samples are ordered phylogenetically from the deepest-rooting sample (from left to right). The plot highlights higher sequence similarity between the *DYZ18* and 3.1-kbp repeat arrays, and between the 2.7-kbp and *DYZ1* repeat arrays, respectively.

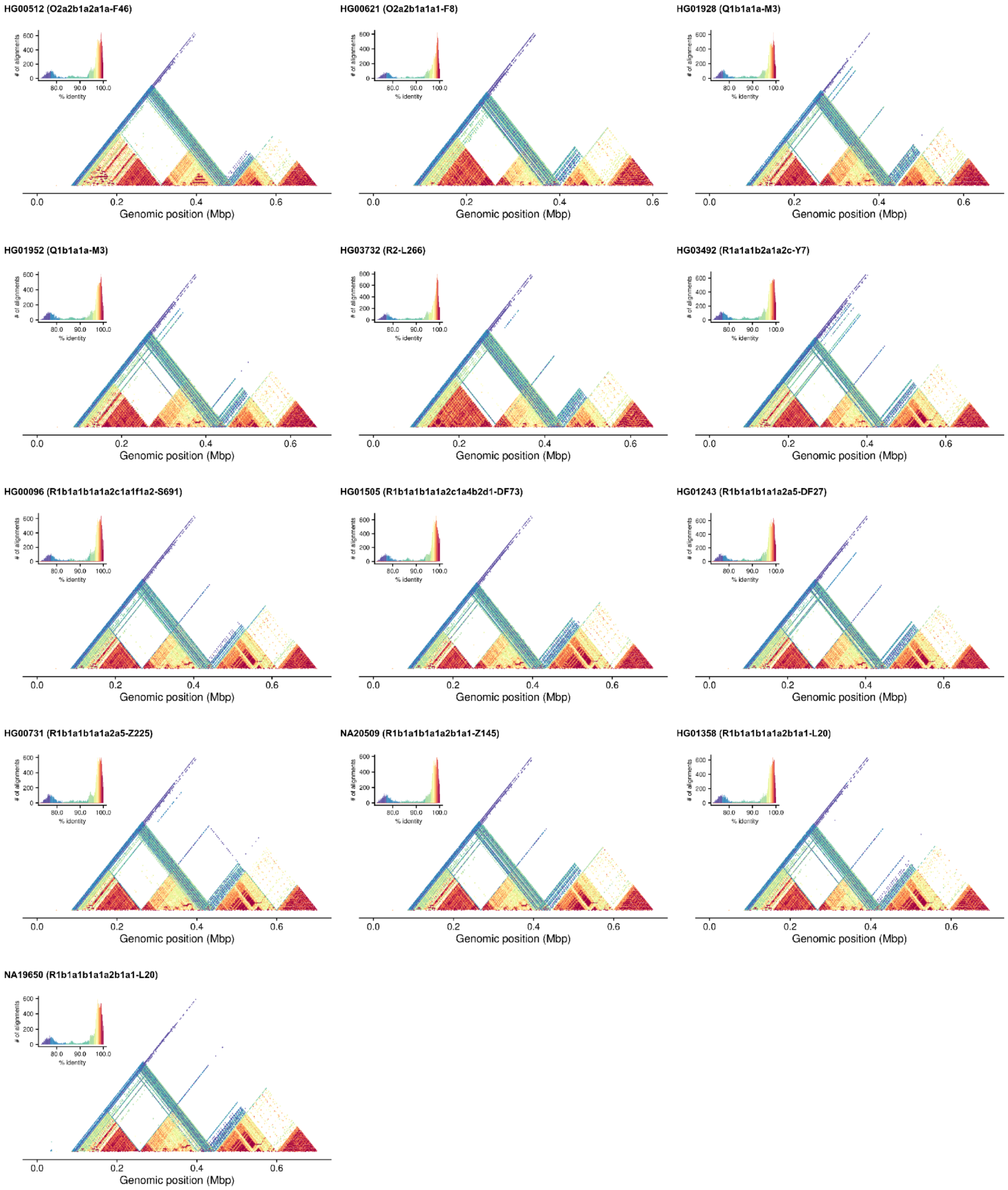


**Figure S69.** Sequence identity heat map of the Yq11/Yq12 transition region, including the *DYZ18*, 3.1-kbp, 2.7-kbp repeat arrays and 100 kbp of the first *DYZ1* repeat array generated using StainedGlass with 2 kbp window size. 100 kbp proximal to the *DYZ18* repeat array has also been included. Samples are ordered phylogenetically from the deepest-rooting sample (from left to right). The plot highlights higher sequence similarity between the *DYZ18* and 3.1-kbp repeat arrays, and between the 2.7-kbp and *DYZ1* repeat arrays, respectively.

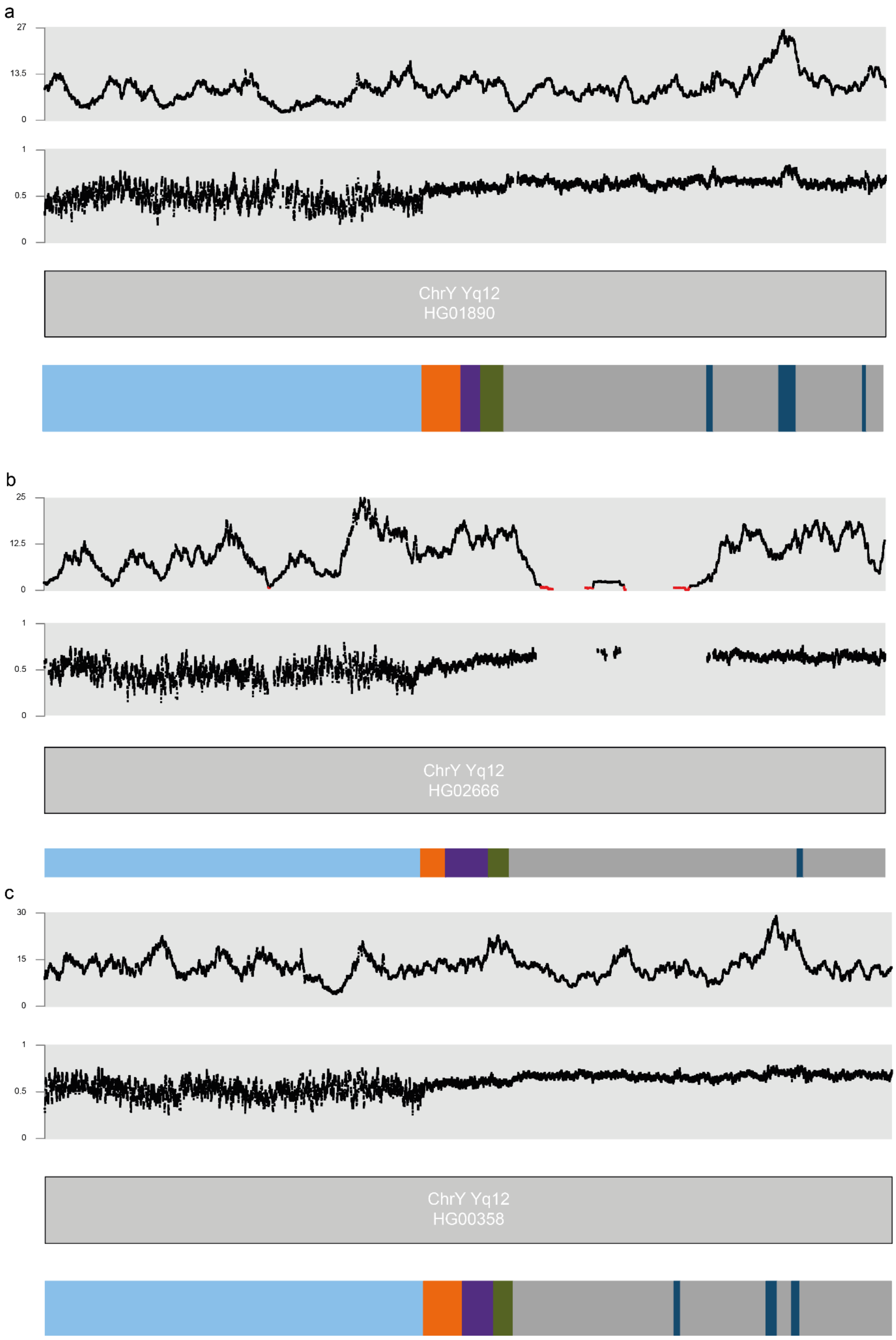


159

160 **Figure S70.** ONT read depth (top) and methylation patterns (below) around the boundary of Yq11 euchromatin and the
161 Yq12 heterochromatic subregion across the three contiguously assembled Y chromosomes, with the sequence annotations
162 shown below (light blue - ampliconic 7 subregion, orange - *DYZ18*, purple - 2.7-kb repeat, green - 3.1-kb repeat, grey -
163 *DYZ1*, dark blue - *DYZ2*).

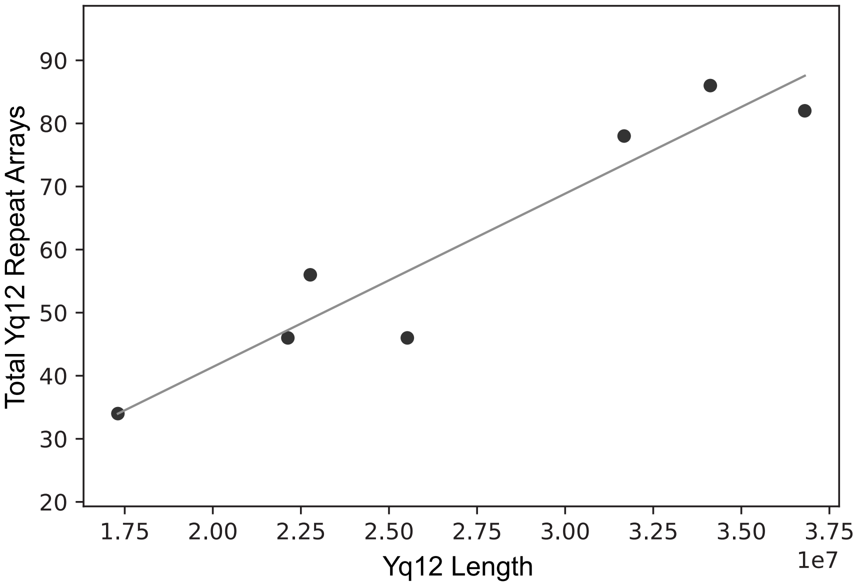


**Figure S46.** A scatter plot showing the total number of *DYZ1* and *DYZ2* arrays within the Yq12 subregions of each sample (n=7, samples with complete assembly plus the T2T Y) (y-axis) versus the total length of the Yq12 region (x-axis) is illustrated. This relationship was found to be significantly positively correlated (two-sided Spearman correlation=0.90; p-value $< 0.05$).

**a**

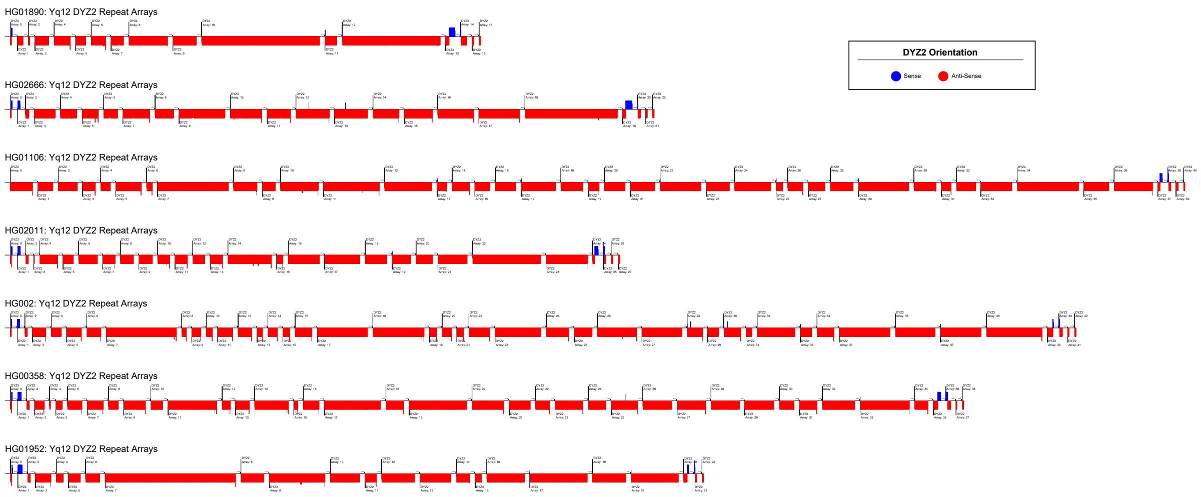


**b**

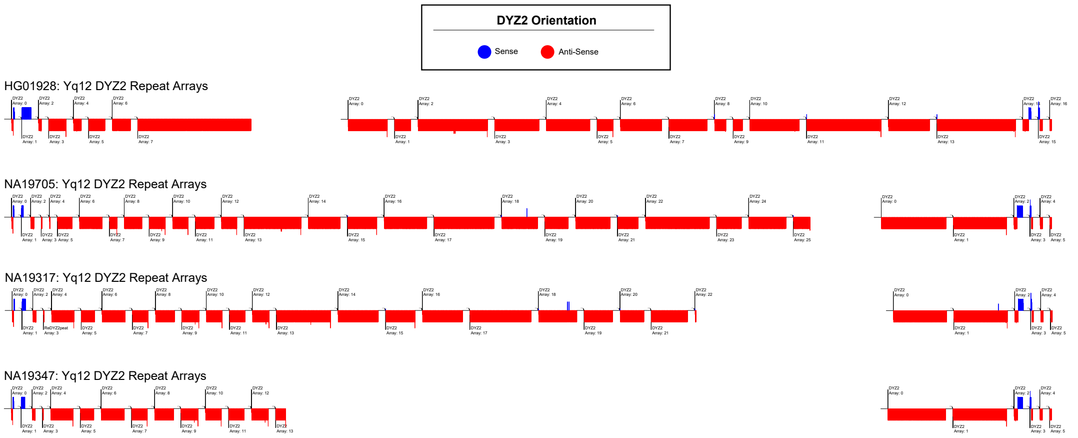


**Figure S72.** Overview of the *DYZ2* repeat array orientation and structure within the Yq12 subregion of each **a.** sample with completely assembled Yq12 subregion, and **b.** the four additional genomes (HG01928, NA19705, NA19317, NA19347) with incompletely assembled Yq12 regions. Red lines indicate individual *DYZ2* repeats in antisense orientation, blue lines indicate individual *DYZ2* repeats in sense orientation relative to the *DYZ2* consensus sequence. The length of each line is a function of the length of the repeat.

**a**

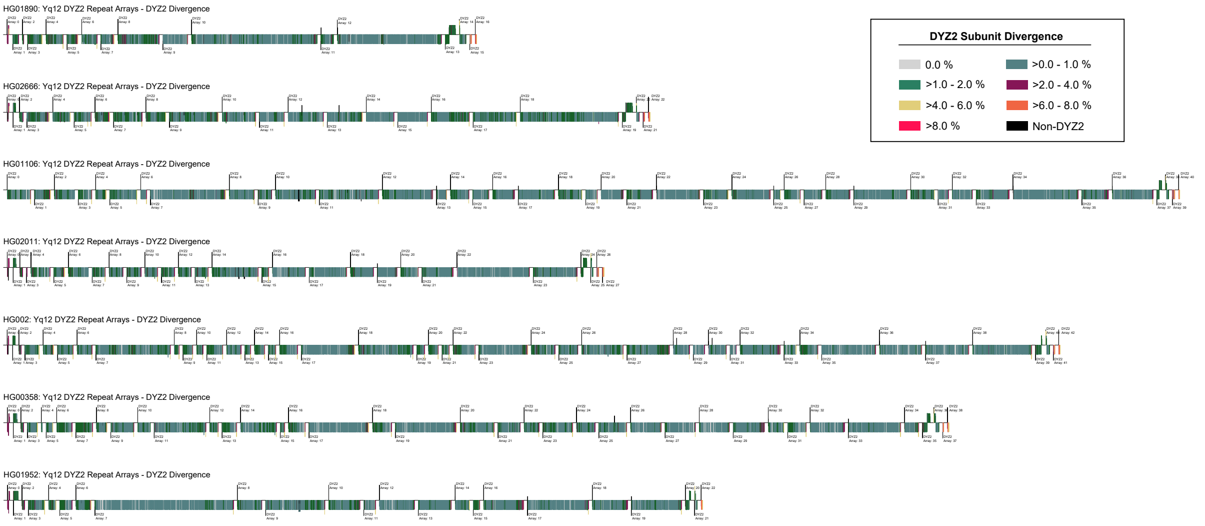


**b**

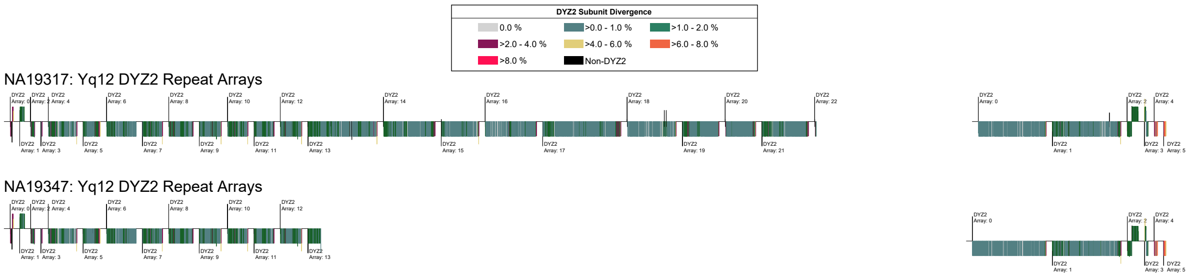


**Figure S73.** An overview of the divergence of individual *DYZ2* subunits for **a.** samples with completely assembled Yq12 subregion (HG01890, HG02666, HG01106, HG02011, T2T Y, HG00358, HG01952), and **b.** the two most closely related genomes (NA19317 and NA19347) with incompletely assembled Yq12 sunregions. A higher divergence was observed within the subunits located in arrays at the proximal and distal ends of the Yq12 subregion. Additionally, *DYZ2* subunits located near the boundaries of individual arrays tend to be more diverged than those located centrally. Between the closely related genomes, the divergence of *DYZ2* repeats within the shared *DYZ2* arrays are extremely similar.

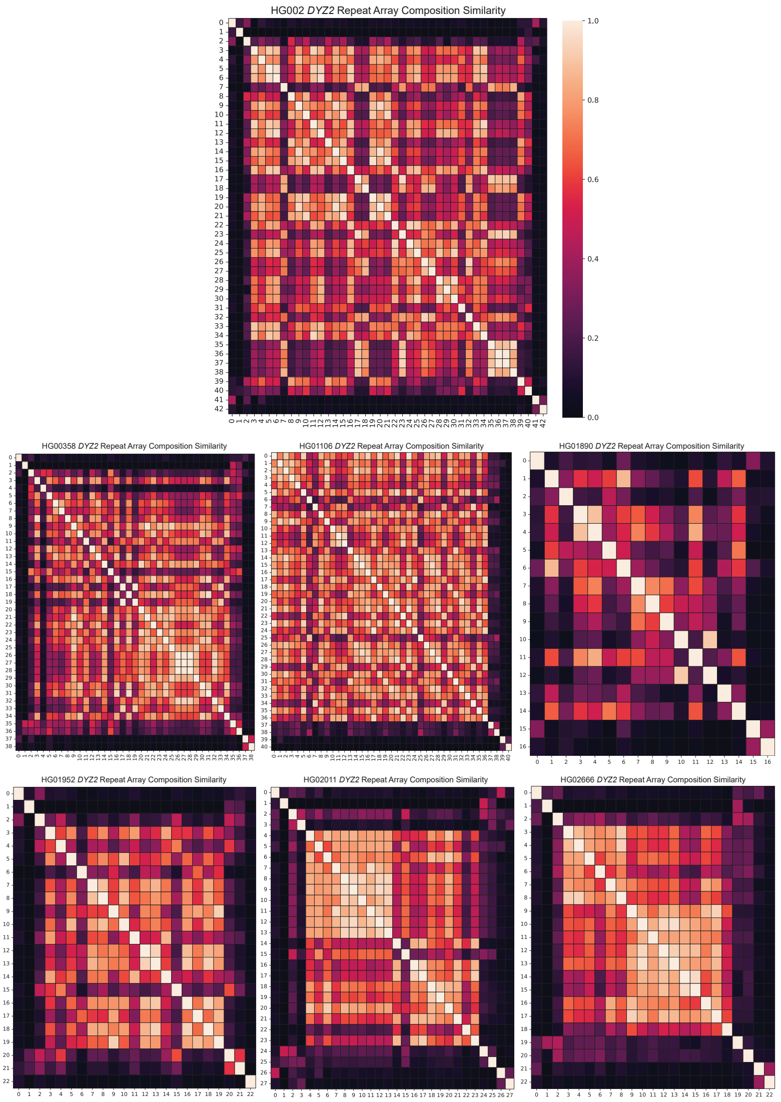


**Figure S74.** Heatmaps showing the complement of the Bray-Curtis (BC) distance/dissimilarity (i.e., 1-BC) for *DYZ2* repeat arrays within each genome with a completely assembled Yq12 subregion. Higher values (i.e., 1.00) indicate *DYZ2* arrays that contain exactly the same subunit composition whereas lower values (i.e., 0.0) suggest the opposite. Results show that the composition of arrays closer to one another tend to be more similar, except for the arrays located in the proximal and distal inversions, which tend to be more similar to each other than to surrounding arrays.

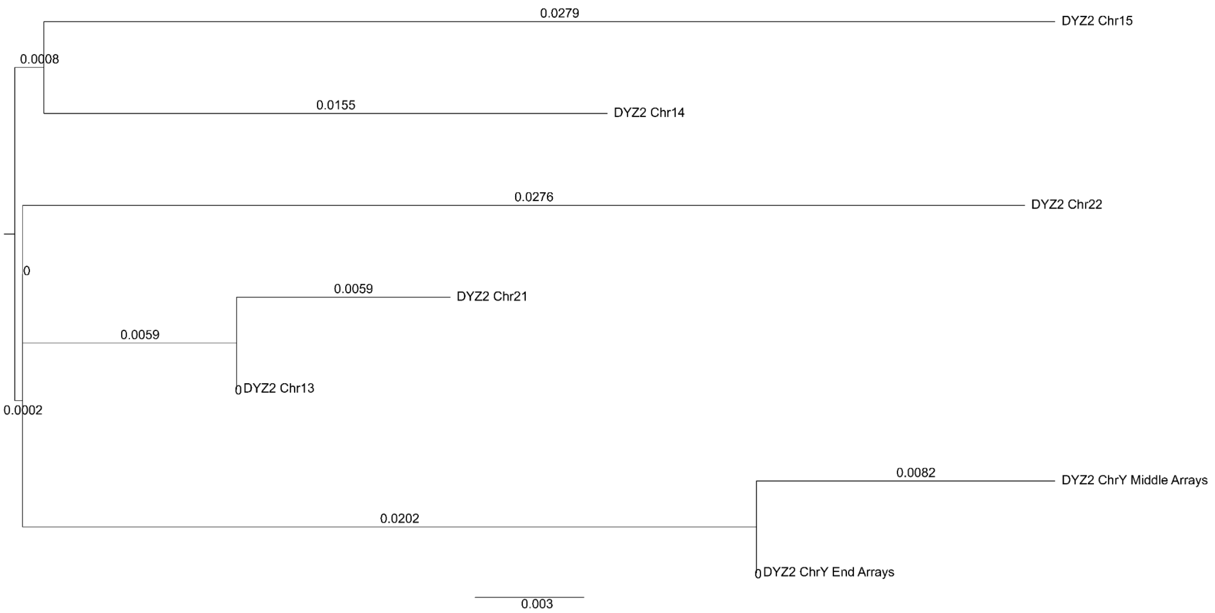


**Figure S75.** A phylogenetic tree of *DYZ2* consensus sequences from all human acrocentric chromosomes. Results show that the consensus sequence of *DYZ2* elements located in arrays outside the Yq12 inversion (ChrY End Arrays) is less divergent to the *DYZ2* consensus sequence of all other acrocentric chromosomes compared to the *DYZ2* consensus sequence of elements situated within arrays between the Yq12 inversion (ChrY Middle Arrays). Branch lengths are displayed. This is an unrooted maximum likelihood consensus tree created from 1000 bootstrap replicates and rooted on the midpoint (see **Methods**).

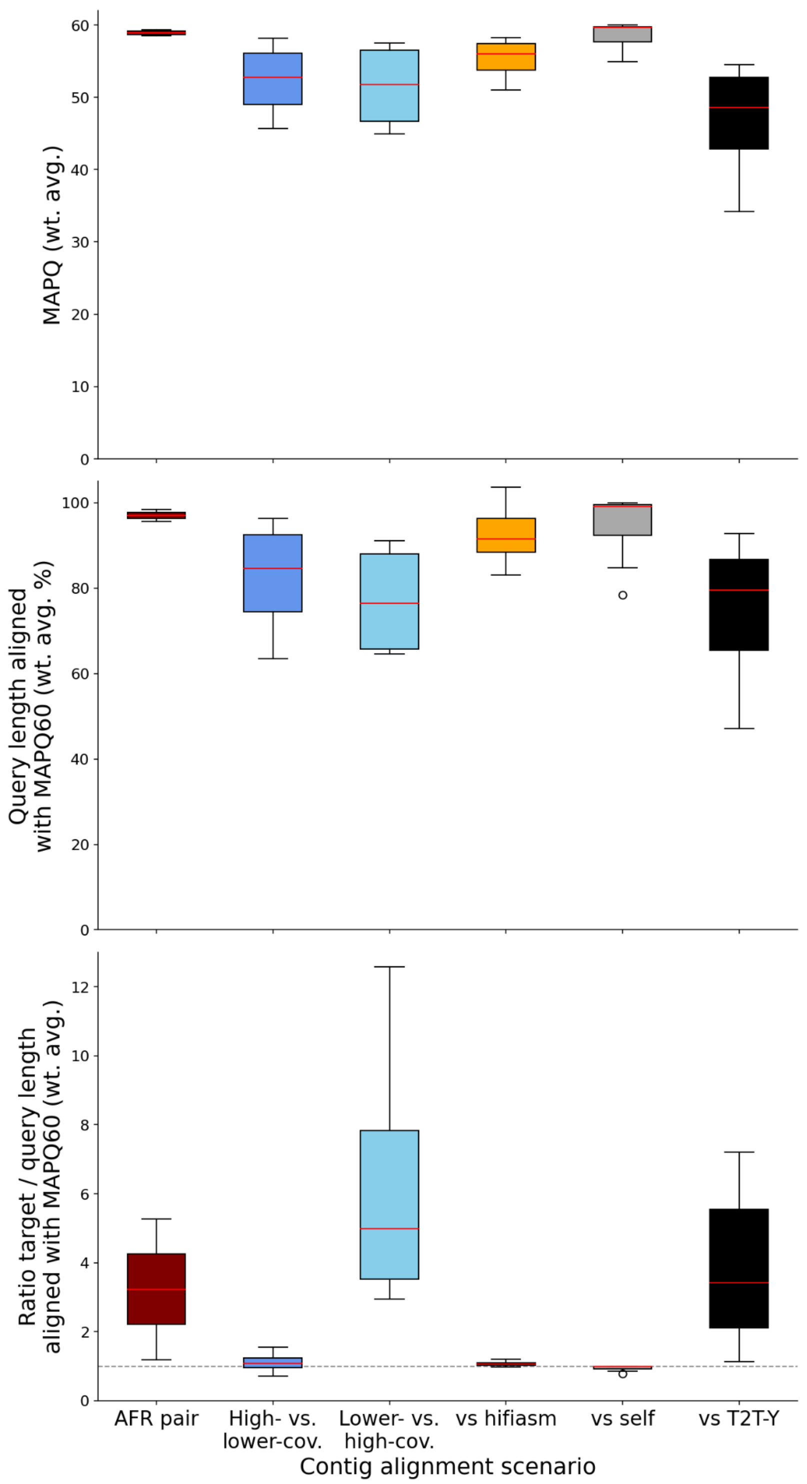


**Figure S76.** QC contig alignments for high-coverage samples in various scenarios: box plots depicting contig alignment statistics for (from left to right) the pair of closely related AFR samples (NA19317 and NA19347, dark red); the four selected high-coverage samples assembled with lower coverage for QC purposes, using the lower-coverage assembly as alignment target (dark blue) and vice versa the high-coverage assembly (light blue); the sample-matched alignment of Verkko- to hifiasm-assembled contigs (orange); the self-alignment of Verkko-assembled contigs (grey); contig-to-reference alignment using the T2T Y sequence as alignment target (black). Computed statistics per sample pair are (from top to bottom) average mapping quality (MAPQ) of the alignments weighted by alignment length (in bp); percent of the query sequence aligning with MAPQ 60, averaged over all contigs weighted by the contig length; ratio of target-to-query sequence lengths aligning with MAPQ 60, averaged over all contigs weighted by contig length.

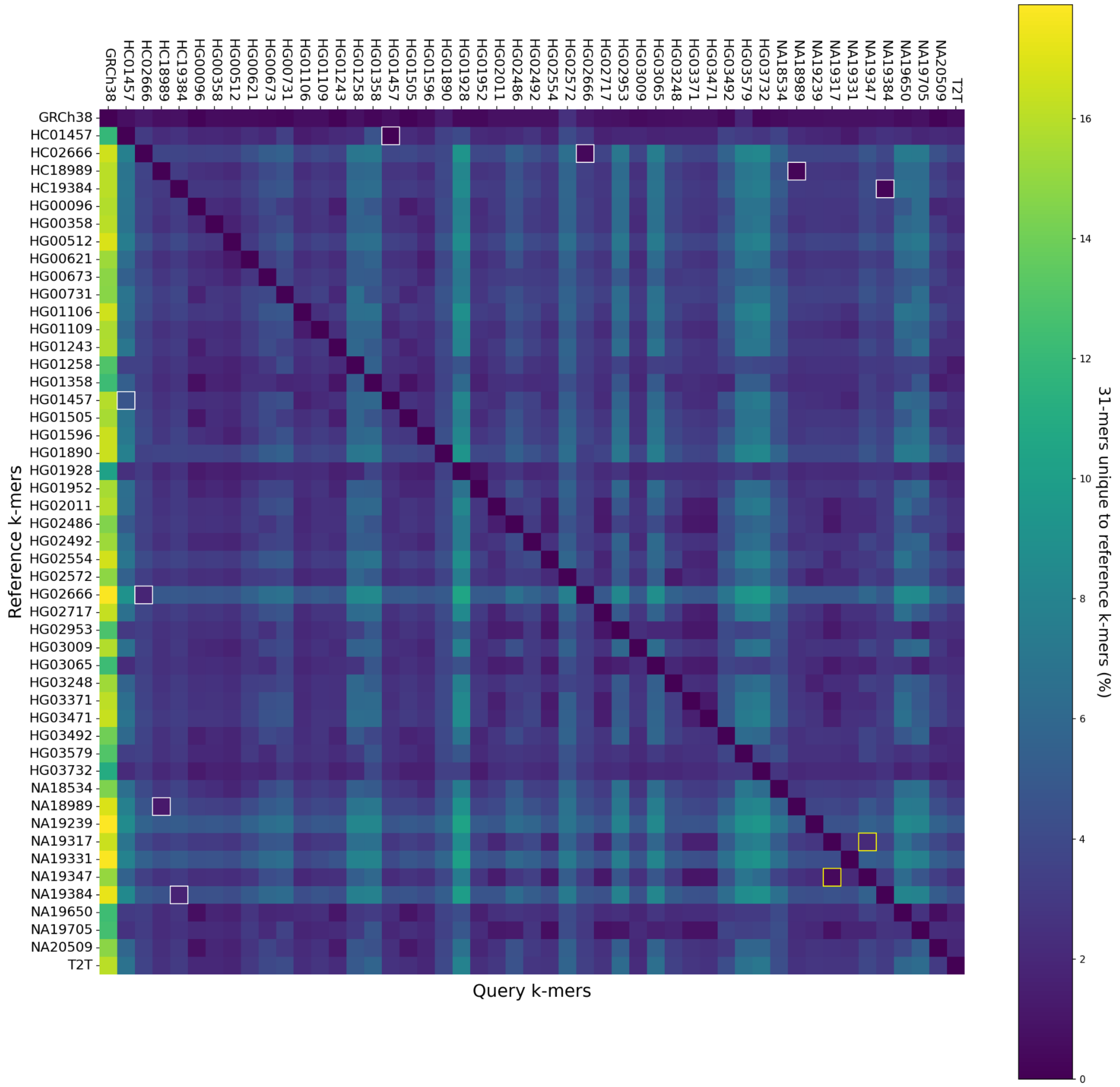


**Figure S77**. Unique sequence content in all assemblies expressed as percent of unique 31-mers relative to the respective query assembly. Comparisons of the high-/low-coverage pairs (HC02666/HG02666, HC01457/HG01457 etc.) are singled out by grey rectangles.

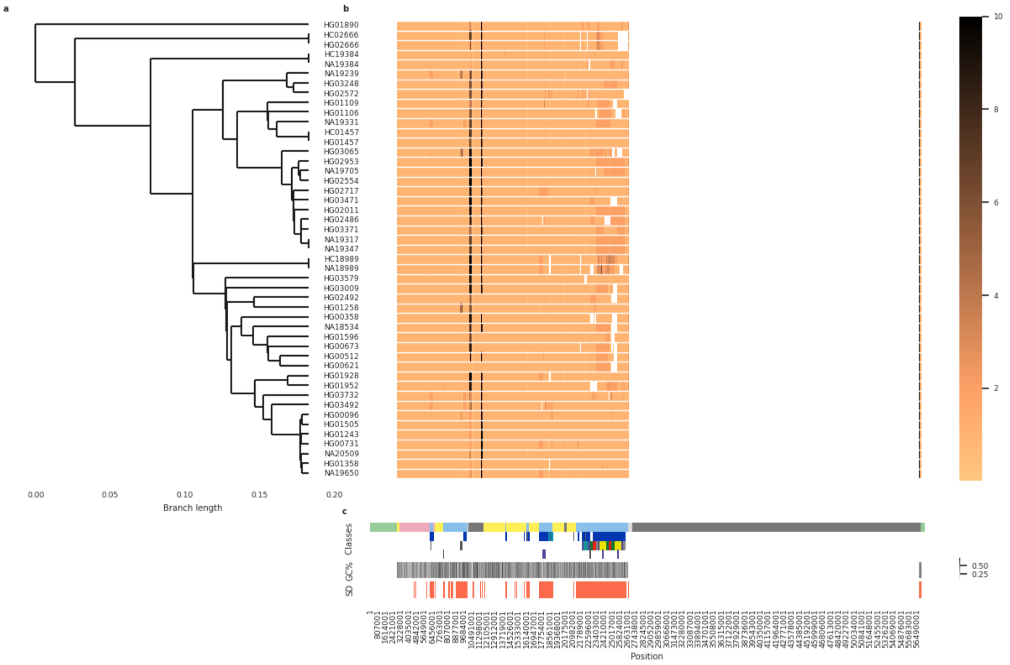


**Figure S78.** Composite plot depicting the Y contig alignments to the GRCh38 Y reference sequence across the whole Y chromosome span. **a.** Phylogenetic relationships of the samples (see **Fig. S1** for details). Note that two assemblies are visualized for 4 samples for which both high- and lower-coverage assemblies were generated (HG02666, NA19384, HG01457 and NA18989; HC - refers to the high-coverage assembly; see Methods). **b.** The coverage from Y contig alignments to reference sequence, with coverage=1 (light orange) in well-aligning regions. Darker shades indicate regions with multiple contig alignments, potentially indicating assembly errors or poor alignments, e.g., due to structural differences between the sample and reference or difficult sequence contexts such as high repeat content; white denotes regions with no coverage i.e., no contig to reference alignments (note - majority of the Yq12 subregion is not resolved in GRCh38, i.e., composed of 'Ns'). **c.** Y-chromosomal subregion locations as described in **Fig. 1a**, locations of inverted repeats (in dark blue) and *AZFc*/ampliconic subregion 7 segmental duplications as shown in **Fig. S39**, followed by GC% and segmental duplication locations (**Methods**).

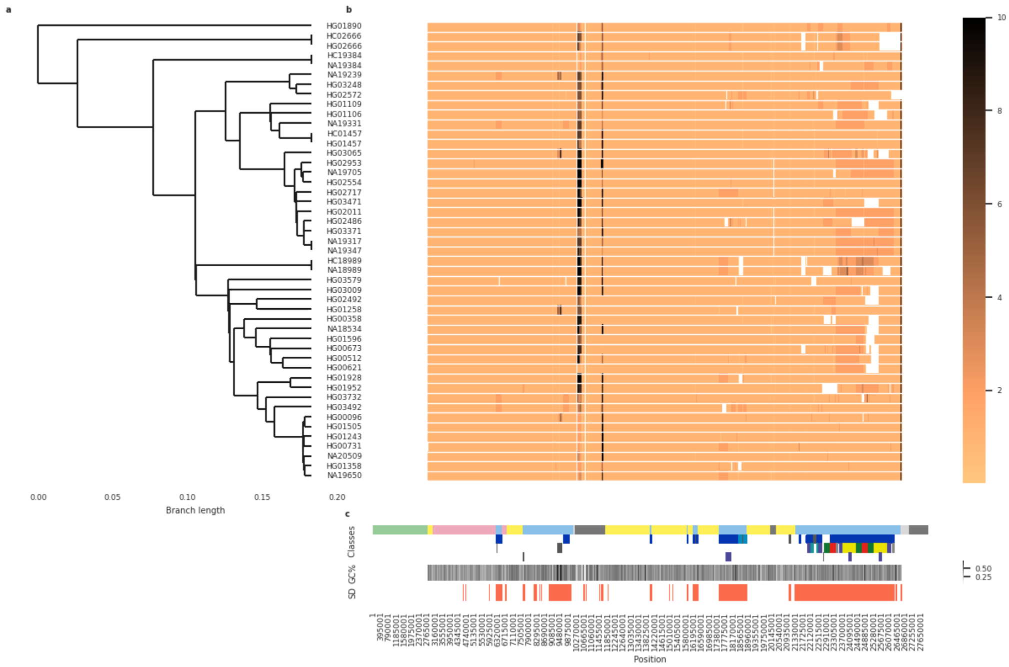


**Figure S79.** Composite plot depicting the Y contig alignments to the GRCh38 Y reference sequence excluding Yq12 and PAR2 subregions. **a.** Phylogenetic relationships of the samples (see **Fig. S1** for details). Note that two assemblies are visualized for 4 samples for which both high- and lower-coverage assemblies were generated (HG02666, NA19384, HG01457 and NA18989; HC - refers to the high-coverage assembly; see Methods). **b.** The coverage from Y contig alignments to reference sequence, with coverage=1 (light orange) in well-aligning regions. Darker shades indicate regions with multiple contig alignments, potentially indicating assembly errors or poor alignments, e.g., due to structural differences between the sample and reference or difficult sequence contexts such as high repeat content; white denotes regions with no coverage i.e., no contig to reference alignments (note - majority of the Yq12 subregion is not resolved in GRCh38, i.e., composed of 'Ns'). **c.** Y-chromosomal subregion locations as described in **Fig. 1a**, locations of inverted repeats (in dark blue) and *AZFc*/ampliconic subregion 7 segmental duplications as shown in **Fig. S39**, followed by GC% and segmental duplication locations (**Methods**).

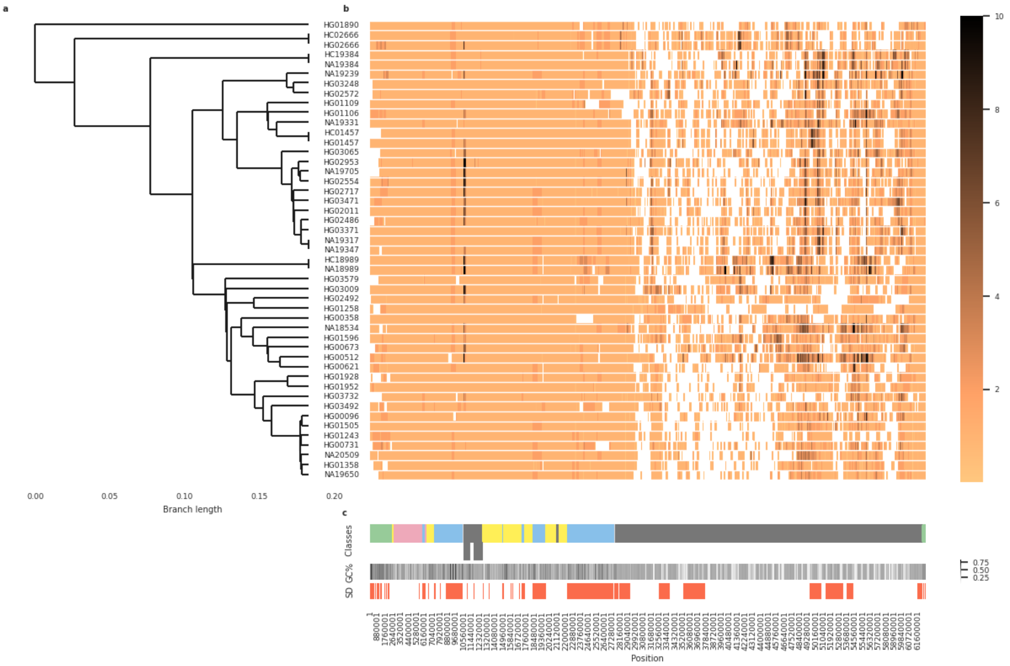


**Figure S80.** Composite plot depicting the Y contig alignments to the T2T Y reference sequence across the whole Y chromosome span. **a.** Phylogenetic relationships of the samples (see **Fig. S1** for details). Note that two assemblies are visualized for 4 samples for which both high- and lower-coverage assemblies were generated (HG02666, NA19384, HG01457 and NA18989; HC - refers to the high-coverage assembly; see Methods). **b.** The coverage from Y contig alignments to reference sequence, with coverage=1 (light orange) in well-aligning regions. Darker shades indicate regions with multiple contig alignments, potentially indicating assembly errors or poor alignments, e.g., due to structural differences between the sample and reference or difficult sequence contexts such as high repeat content; white denotes regions with no coverage i.e., no contig to reference alignments. **c.** Y-chromosomal subregion locations as described in **Fig. 1a**; below in grey locations of *DYZ3* (on the left) and *DYZ17* (on the right) repeat arrays, followed by GC% and segmental duplication locations (**Methods**).

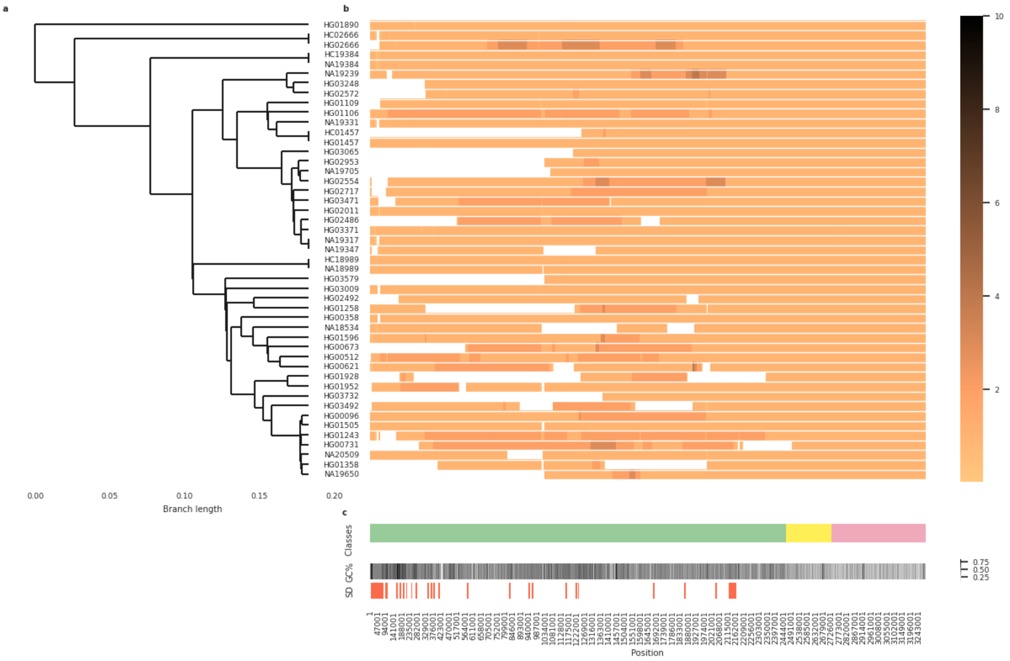


**Figure S81.** Composite plot depicting the Y contig alignments to the T2T Y reference sequence zooming into PAR1 subregion. **a.** Phylogenetic relationships of the samples (see **Fig. S1** for details). Note that two assemblies are visualized for 4 samples for which both high- and lower-coverage assemblies were generated (HG02666, NA19384, HG01457 and NA18989; HC - refers to the high-coverage assembly; see Methods). **b.** The coverage from Y contig alignments to reference sequence, with coverage=1 (light orange) in well-aligning regions. Darker shades indicate regions with multiple contig alignments, potentially indicating assembly errors or poor alignments, e.g., due to structural differences between the sample and reference or difficult sequence contexts such as high repeat content; white denotes regions with no coverage i.e., no contig to reference alignments. **c.** Y-chromosomal subregion locations as described in **Fig. 1a**, followed by GC% and segmental duplication locations (**Methods**).

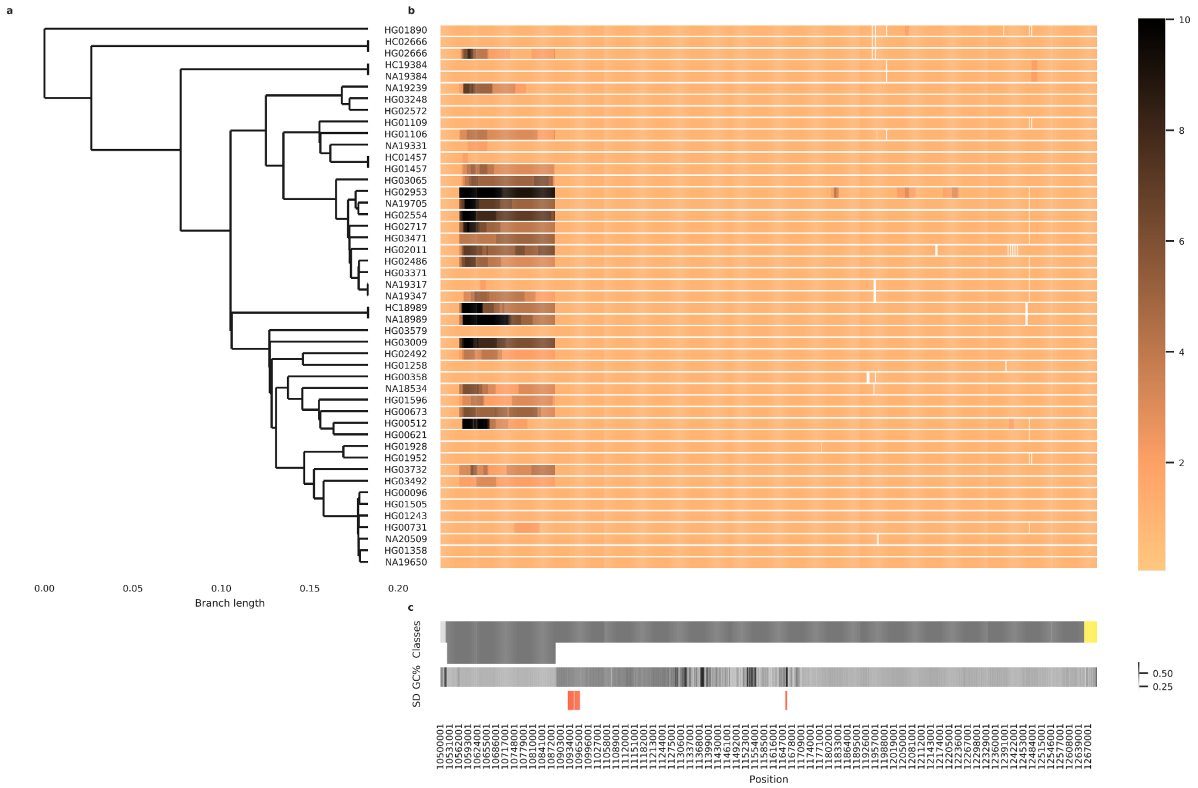


**Figure S82.** Composite plot depicting the Y contig alignments to the T2T Y reference sequence zooming into the (peri-)centromeric region. **a.** Phylogenetic relationships of the samples (see **Fig. S1** for details). Note that two assemblies are visualized for 4 samples for which both high- and lower-coverage assemblies were generated (HG02666, NA19384, HG01457 and NA18989; HC - refers to the high-coverage assembly; see Methods). **b.** The coverage from Y contig alignments to reference sequence, with coverage=1 (light orange) in well-aligning regions. Darker shades indicate regions with multiple contig alignments, potentially indicating assembly errors or poor alignments, e.g., due to structural differences between the sample and reference or difficult sequence contexts such as high repeat content; white denotes regions with no coverage i.e., no contig to reference alignments. **c.** Location of the (peri-)centromeric region (above) and the *DYZ3* α-satellite repeat array (below) shown in dark grey, followed by GC% and segmental duplication locations (**Methods**).

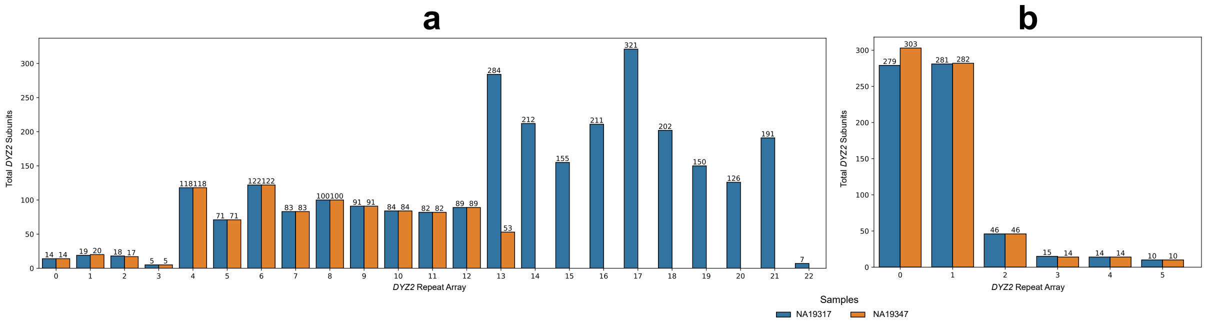


**Figure S83.** The bar plots show a comparison of the total *DYZ2* repeat copy numbers (y-axis) in each *DYZ2* array (x-axis) within the two most closely related genomes (NA19317 (blue) and NA19347 (orange)). **a.** *DYZ2* arrays within the proximally assembled contigs. **b.** *DYZ2* arrays within the distally assembled contigs. The analyses revealed an equal number of *DYZ2* repeats within 14 of 20 arrays.

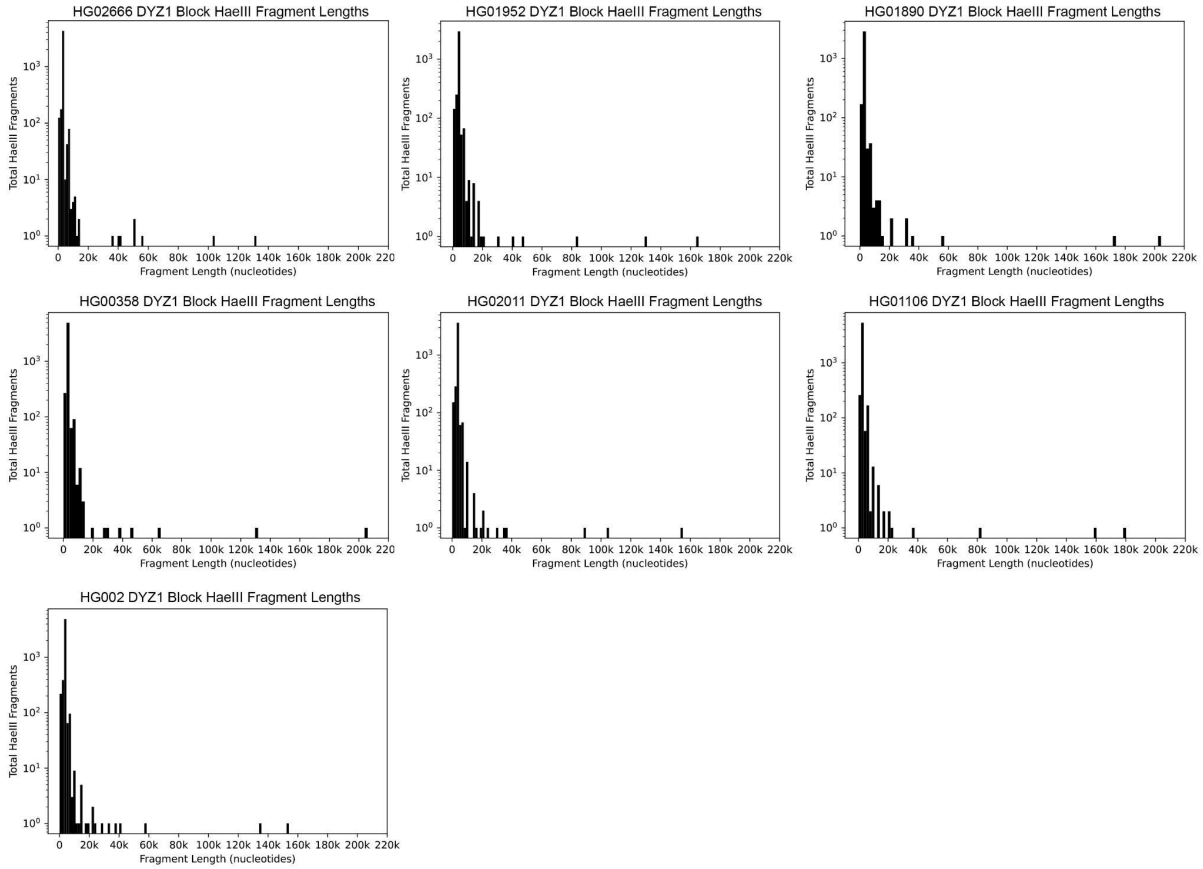


**Figure S84**. The distribution of total *DYZ1* array HaeIII virtual restriction digestion fragments (y-axis) and their lengths (x-axis) for each genome with a completely assembled Yq12 region is shown in the histograms. The majority of *DYZ1* repeat units were between 3-4 kbp in length within each genome.

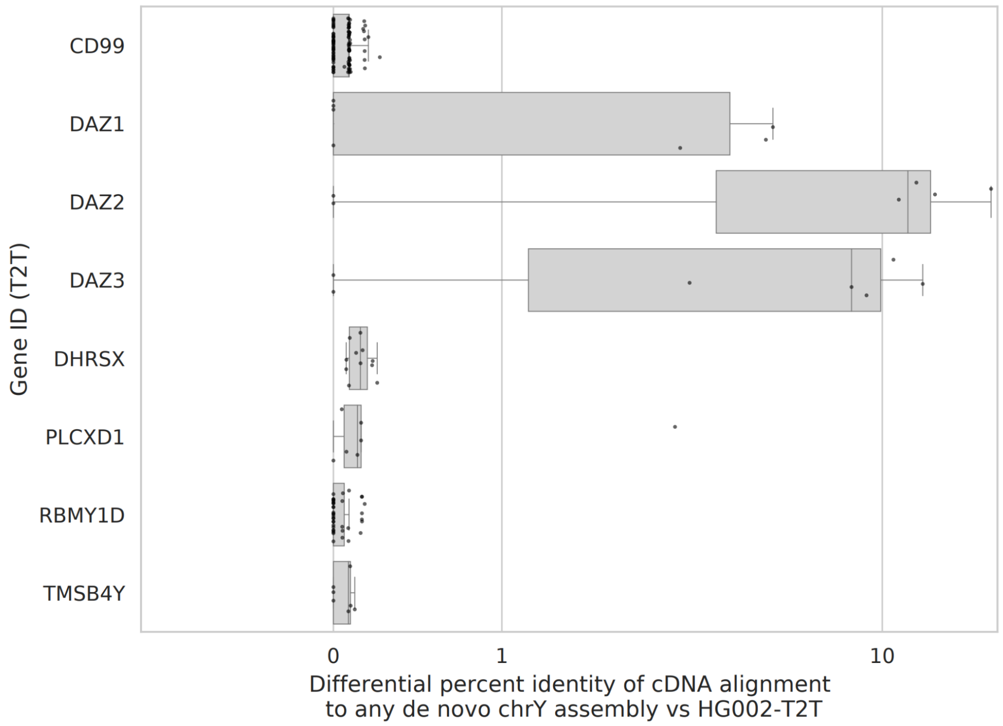


**Figure S85.** Testis Iso-seq percent identity to *de novo* assemblies compared to the T2T Y reference sequence. Each dot represents an individual cDNA read, and its position on the x-axis represents the numeric difference between percent identity of the read alignment to the T2T Y reference and the alignment to the best-matching *de novo* Y assembly. Gene IDs are based on alignment position to the reference.

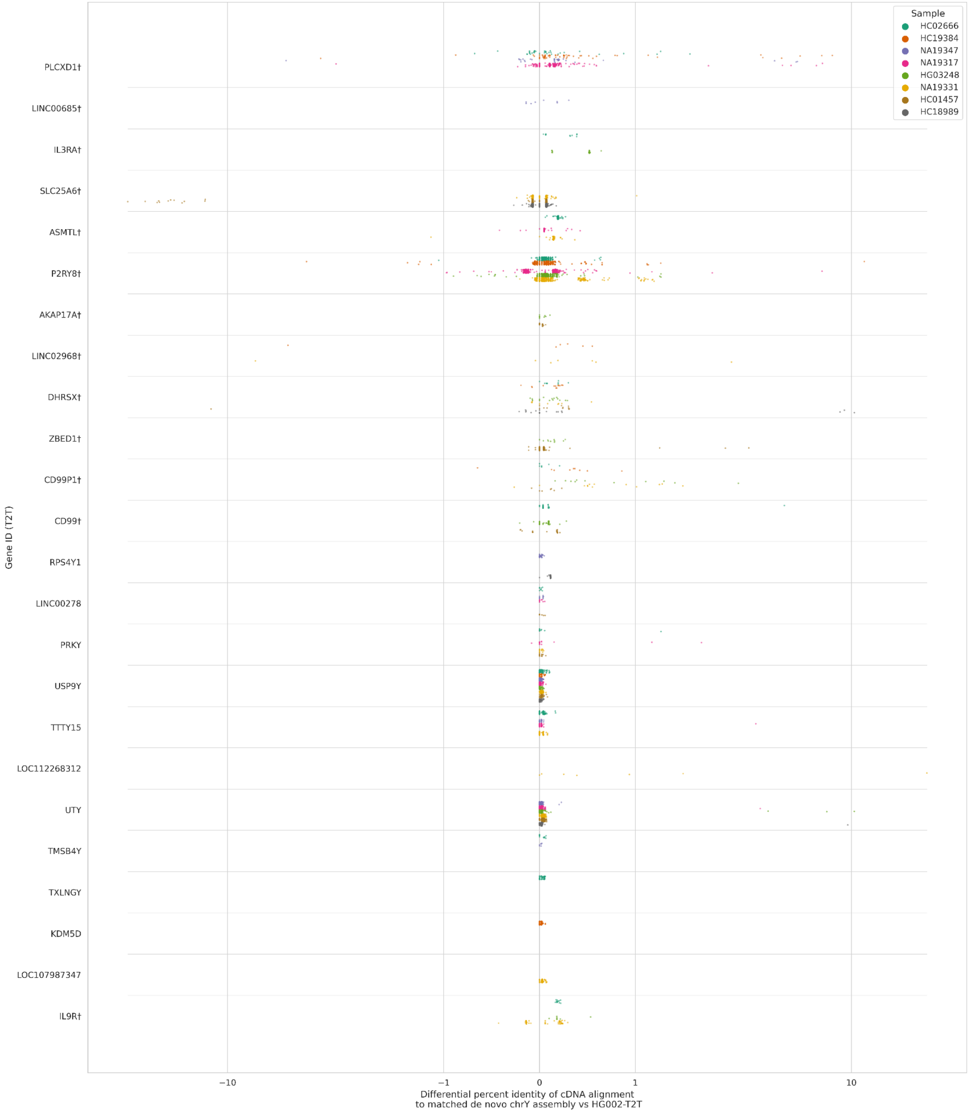


I

**Figure S86:** Iso-seq percent identity to matched *de novo* assemblies compared to the T2T Y reference sequence. Each dot represents an individual cDNA read, and its position on the x-axis represents the numeric difference between percent identity of the read alignment to the T2T Y reference and the alignment to its sample-matched *de novo* Y assembly. Gene IDs are based on alignment position to the reference, with † indicating genes located in either PAR. Colours specify the sample for both the Iso-Seq library and *de novo* assembly.

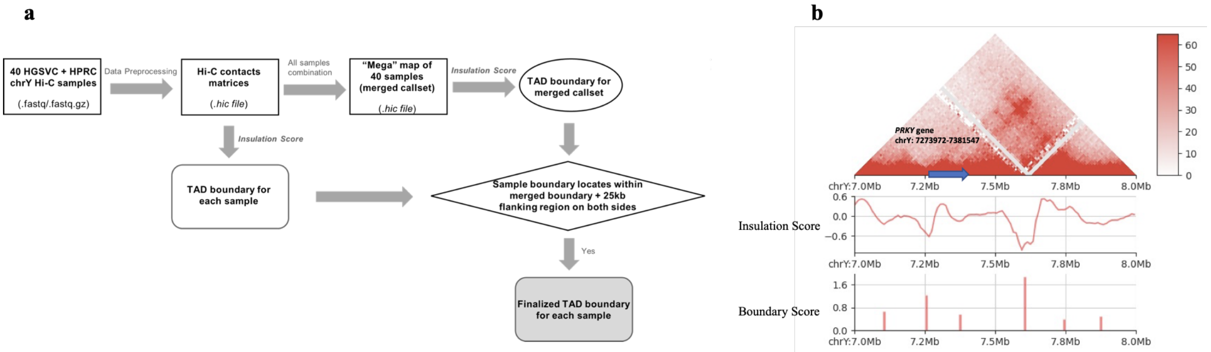


**Figure S87.** A step-by-step workflow to generate TAD boundaries in our chrY Hi-C analysis pipeline and a visualization of the chrY Hi-C merged callset calling results generated in our pipeline. **a.** 40 samples’ raw reads were used as an input in Juicer to do pre-processing and create Hi-C maps which were binned at multiple resolutions. Insulation score algorithm was applied to call TAD boundaries for each sample on each of those 40 .hic files separately. All 40 .hic files were then merged together to create a “mega” map and used as an input of insulation score algorithm to call TAD boundaries for the chrY merged callset. A finalized TAD boundary results for each sample were defined as those sample boundaries located within the merged boundary plus 25 kb flanking regions on the left side of the boundary start site and the right side of the boundary end site. **b.** The Hi-C contact map, the insulation score and the boundary strength for the merged callset over the region chrY: 7Mb-8Mb. The blue arrow shows where the *PRKY* gene is.

a

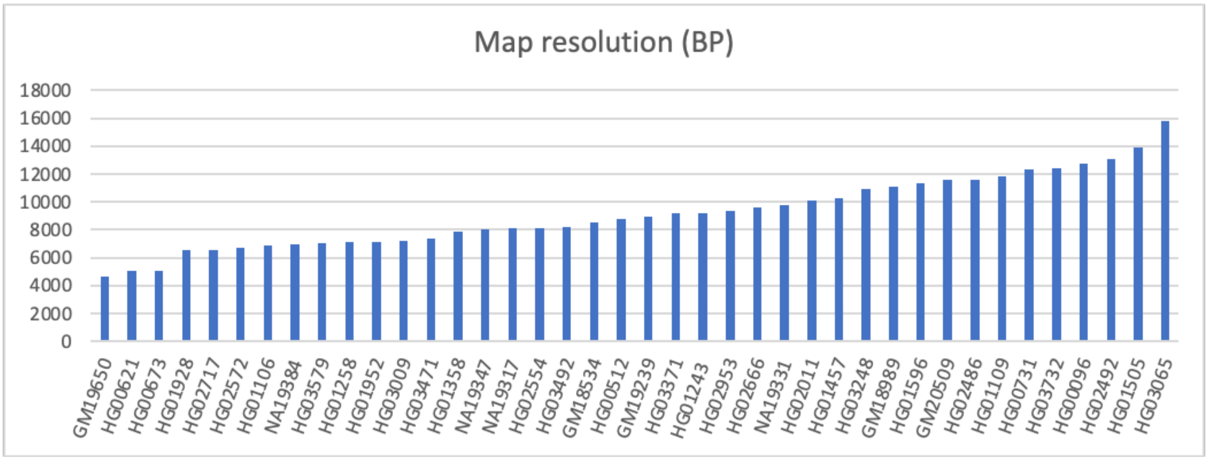


b

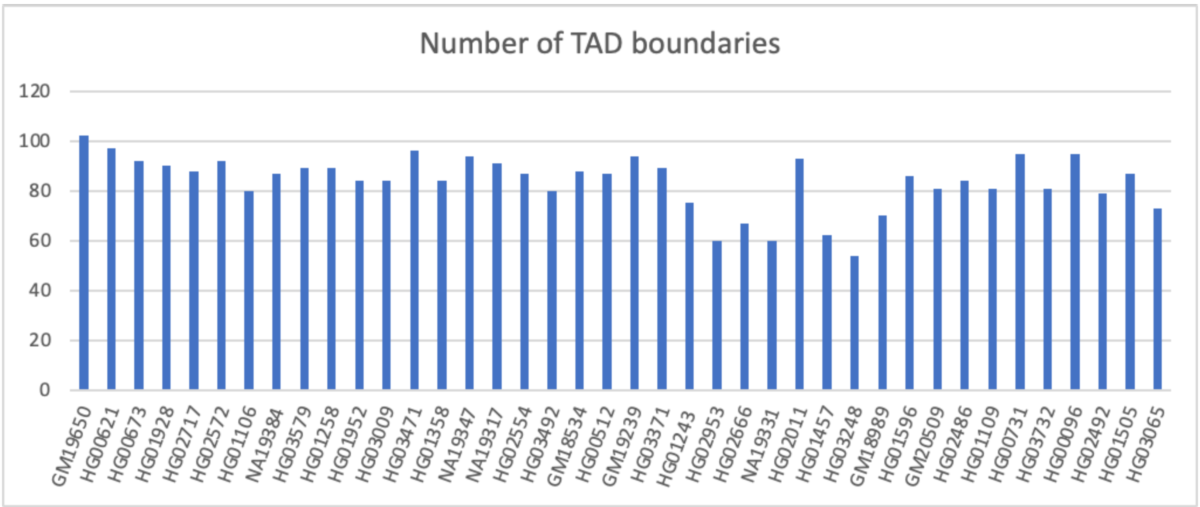


**Figure S88.** The map resolution (bp) for 40 Hi-C samples and the corresponding TAD boundaries detected by our strategy. **a.** As described in [46], the map resolution was calculated by calculate_map_resolution.sh script given by Juicer. The highest resolution is 4,650 bp while the lowest resolution is 15,800 bp. To average, 10 kbp resolution was chosen for the further analysis. **b.** The number the TAD boundaries for each sample which were redefined from the workflow shown in **Figure S87**.
